# Single-nuclei transcriptomes from human adrenal gland reveals distinct cellular identities of low and high-risk neuroblastoma tumors

**DOI:** 10.1101/2021.03.26.437162

**Authors:** O.C. Bedoya-Reina, W. Li, M. Arceo, M. Plescher, P Bullova, H. Pui, M. Kaucka, P. Kharchenko, T. Martinsson, J. Holmberg, I. Adameyko, Q. Deng, C. Larsson, C.C. Juhlin, P. Kogner, S. Schlisio

## Abstract

Childhood neuroblastoma has a remarkable variability in outcome. Age at diagnosis is one of the most important prognostic factors, with children less than 1 year old having favorable outcomes. We studied single-cell and single-nuclei transcriptomes of neuroblastoma with different clinical risk groups and stages, including healthy adrenal gland. We compared tumor cell populations with embryonic mouse sympatho-adrenal derivatives, and post-natal human adrenal gland. We provide evidence that low and high-risk neuroblastoma have different cell identities, representing two disease entities. Low-risk neuroblastoma presents a transcriptome that resembles sympatho- and chromaffin cells, whereas malignant cells enriched in high-risk neuroblastoma resembles an unknown subtype of TRKB+ cholinergic progenitor population identified in human post-natal gland. Analyses of these populations revealed different gene expression programs for worst and better survival in correlation with age at diagnosis. Our findings reveal two cellular identities and a composition of human neuroblastoma tumors reflecting clinical heterogeneity and outcome.

## Introduction

Neuroblastoma (NB) is a pediatric cancer arising from the sympathoadrenal cell lineage frequently originating in the adrenal glands (AG) [1]. This malignancy represents 8-10% of all childhood cancer cases, and is responsible for 15% of all pediatric oncology deaths worldwide [2]. A clinical hallmark of neuroblastoma is heterogeneity, featuring outcomes ranging from lethal progression to spontaneous regression. The risk classification predicting the clinical behavior of the malignancy and its response to treatment, utilizes the INRGSS criteria (i.e. International Neuroblastoma Risk Grouping Staging System) [1,3]. One of the most significant and clinically relevant factor for this risk classification is age. Children younger than 18 months at the time of diagnosis display better prognosis (i.e. low-risk) than children diagnosed at a later age, and aging is in turn associated a with poorer outcome (i.e. high-risk) [4,5]. Other prognostic markers are used to assign patients to specific risk groups, for example ploidy, chromosomal alterations, *MYCN* amplification, and expression of neurotrophin receptors, such as TRKB (encoded by *NTRK2*) associated with high-risk and poor outcome. In contrast, neurotrophin receptor TRKA expression (encoded by *NTRK1*) is associated with low-risk and favorable outcome [2]. The reason why the age of the patient at the time of diagnosis is one of the strongest predictor of risk and outcome is not understood.

Previously, it has been reported that the majority of mouse chromaffin cells forming the adrenal medulla originate from an embryonic neural crest progeny, specifically, from multipotent Schwann cell precursors (SCPs). SCPs are nerve associated cells that migrate along the visceral motor nerves to the vicinity of the developing adrenal gland, and form approximately 80% of the chromaffin cells [6]. The remaining 20% is directly derived from a migratory stream of neural crest cells (NCCs) that commit to a common sympathoadrenal lineage in proximity to the dorsal aorta [7,8], and is considered as the source of neuroblastoma [9,10]. SCPs also give rise to paraganglia during mouse embryonic development, such as chromaffin cells in Zuckerkandl’s organ (ZO) and to some sympathetic neurons [11]. In mouse, the ZO reaches a maximum cell number shortly before birth, in contrast to human where the peak of cell number is reached around the 3rd year of life, indicating species-specific developmental differences [12]. However, SCPs are retained for a rather short time during mouse embryonic development and disappear at around E15 [6]. Thus, it remains unknown how human chromaffin cells are produced and regenerated after birth and if any distinct population of cells serves as their progenitor pool.

We hypothesize that post-natal chromaffin cells are derived from a different source than their developmental origin (namely SCPs), and aberrations in this cell source could explain the age-dependent risk stratification of neuroblastoma. To understand the causes of clinical heterogeneity in neuroblastoma, we deep-sequenced full-length coverage RNA from single nuclei of tumors (*n*=11) across different risk groups. Further, cell clusters were identified, and their transcriptomes were cross-compared with those of cell clusters obtained from healthy tissue where the malignancy is commonly detected, specifically with human (*n*=3) and mouse (*n*=5) post-natal adrenal glands. For this comparison we also included recently published single-cell sequencing datasets from 10X single-sequenced NB tumors (*n*=8) [13], E12-E13 embryonic mouse adrenal anlagen [6], human fetal adrenal gland [14], and the transcriptional profiles of neuroblastoma mesenchymal-/NCC-like and (nor-)adrenergic cell lineages [15,16]. We identified a cluster of TRKB+ cholinergic cells in the human post-natal adrenal gland, that differ from previously described embryonic Schwann cell precursors (SCP). This TRKB+ population of cells shared a specific gene signature with a cluster of undifferentiated cells of mesenchymal nature enriched in high-risk neuroblastomas which was coupled with lower patient survival probability and older age-at-diagnosis when tested in a larger cohort of 498 neuroblastoma patients [17]. Conversely, more differentiated noradrenergic cells are over-represented in low-risk cases, and share specific gene signatures with adrenal human and mouse transcriptomes that resembles signature of sympatho- and chromaffin cells.

All together our results suggest that high-risk neuroblastomas are characterized by a population of progenitor cells that resemble a cell type in post-natal adrenal gland with migratory and mesenchymal signatures, while the low-risk neuroblastoma resemble post-natal and developing chromaffin cells and sympathoblasts.

## Results

To understand why high-risk neuroblastomas arises in children older than 18 months, we first cataloged normal cell populations in post-natal adrenal gland which is a common location for this pediatric malignancy. We define the identity of normal cell populations in the adrenal gland of both post-natal human (*n*=3, 1,536 single nuclei) and mouse (*n*=5, 1,920 single whole-cells) by single-nuclei/cell RNA-sequencing (SmartSeq2, see Methods) to an average depth of 485,000- and 669,000 reads per nuclei/cell, respectively (Supplementary Figure 1a-b, Supplementary Table 1). Technical and biological features of the transcription profile acquired from nuclei (human) or from whole cells (mouse) are summarized in Supplementary Figure 1c.

### Post-natal human and mouse adrenal glands share cell populations but exhibit differences in chromaffin cells

The cell populations in human and mouse adrenal glands were annotated under the expectation of recovering both adrenal cortex- and medulla-associated cells (Figure 1a for human, 1b for mouse, Supplementary Table 2). A reference guide of normal adrenal cell populations was generated by assigning an identity to each cluster, by cross-referencing significantly up-regulated transcripts with canonical markers curated from the literature (Supplementary Table 2). Adrenal medulla cells were identified by the expression of a panel of nor- and adrenergic markers, including *PNMT, TH, DBH, CHGA* and *CHGB* (Figure 1d, Supplementary Table 2). In human, hC4 was identified as the chromaffin cell cluster (“NOR” panel were significantly up-regulated, *FDR*<0.01, Welch’s *t*-test, Figure 1a,d), whereas in mouse two chromaffin cell clusters (i.e. mC11 and mC15) were identified (noradrenergic markers exemplified in “NOR” panel were significantly up-regulated, *FDR*<0.01, Welch’s *t*-test, Figure 1b,d, Supplementary Table 2). Nevertheless, the mouse chromaffin population mC15 shared a more significant specific gene signature with the human chromaffin cluster hC4 (*FDR*<0.01, Fisher’s exact test, Figure 1c, Supplementary Table 3). To understand the differences between the two mouse chromaffin populations (mC11 and mC15), we investigated the expression of genes that differed significantly between them. The expression of *CHGA, CHGB, PHOX2A*, and *PHOX2B* were significantly higher in mC15 than in mC11 (*FDR*<0.01, Welch’s */-* test), while the expressions of *PNMT* was higher in mC11 than in mC15 (*FDR*<0.01, Welch’s *t*-test, Figure 1d, Supplementary Table 2). Additionally, mC15 cluster exhibited a significantly higher expression of a different repertoire of cholinergic muscarinic and nicotinic receptors (mAChR and nAChR) than mC11 including *CHRM1, CHRNA3, CHRNA7*, and *CHRNB4* (*FDR*<0.01, Welch’s *t*-test, Supplementary Figure 2a-b, Supplementary Table 2). In contrast to mouse chromaffin cells, human postnatal *PNMT+* chromaffin cells (cluster hC4) showed a significant expression of the sympathoblast marker *PRPH* (*FDR*<0.01, Welch’s *t*-test, Figure 2a).

**Figure 1.**
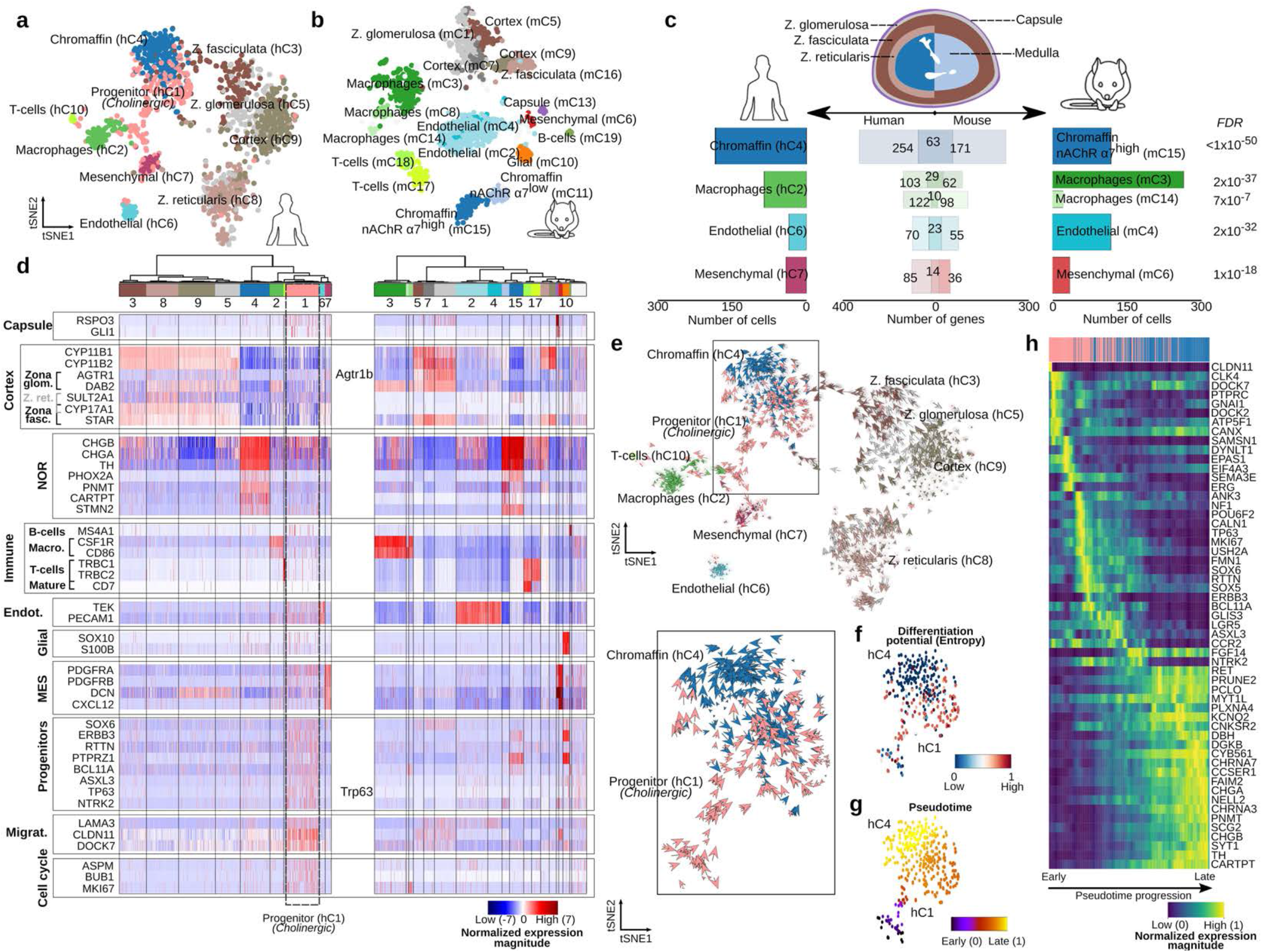
Anatomy of human and mouse adrenal gland (AG) revealed from single nuclei/cell analysis. **a**, 1,536 single nuclei of human AG from three different patients were sequenced with Smart-seq2 to an average depth of 485,000 reads per cell. Cells with high-quality (*n*=1,322) were selected and further processed with PAGODA. Cells were grouped into ten different clusters, including cortex (in brown and gray colors), chromaffin (blue hC4), mesenchymal (purple hC7), endothelial (light blue hC6) and immune cells (i.e. T-cells hC10 and Macrophages hC2 in green colors). **b**, 1,920 single cell of mouse AG from five different samples were sequenced with Smart-seq2 to an average depth of 670,000 reads per cell. High-quality reads were mapped to the mouse genome with STAR, and gene expression was estimated with HTSeq. Cells with high-quality (*n*=1,763) were selected and further processed with PAGODA. Cells were grouped into nineteen different clusters, including cortex (in brown and gray colors), chromaffin (blue mC15 and light blue mC11), mesenchymal (red mC6), capsule (purple mC13), endothelial (light blue mC2 and mC4), glial (orange mC10) and immune cells (i.e. T-cells mC17 and mC18, and Macrophages mC3, mC8 and mC14 in green colors). **c-d**, A comparison of the specific gene signature between human and mouse revealed a similar transcription signatures for mesenchymal, endothelial and immune, nevertheless the two different post-natal chromaffin cells can only be differentiated in mouse (i.e. by the expression of *CHRNA7).* **e-g**, Furthermore, a population of cells with progenitor markers (i.e. *SOX6+, ERBB3+, RTTN+*) and high differentiation potential found uniquely in human AG, is sourcing the chromaffin cells as indicated by velocity, entropy, and pseudotime analyses. **h**, In this process, gene expression elapses from an undifferentiated stem-like-(i.e. *RTTN+*) to an adrenergic signature (*PNMT+}*, passing by a noradrenergic stage (i.e. *DBH+)*, as indicated by the pseudotime of the underlying cellular processes.

**Figure 2.**
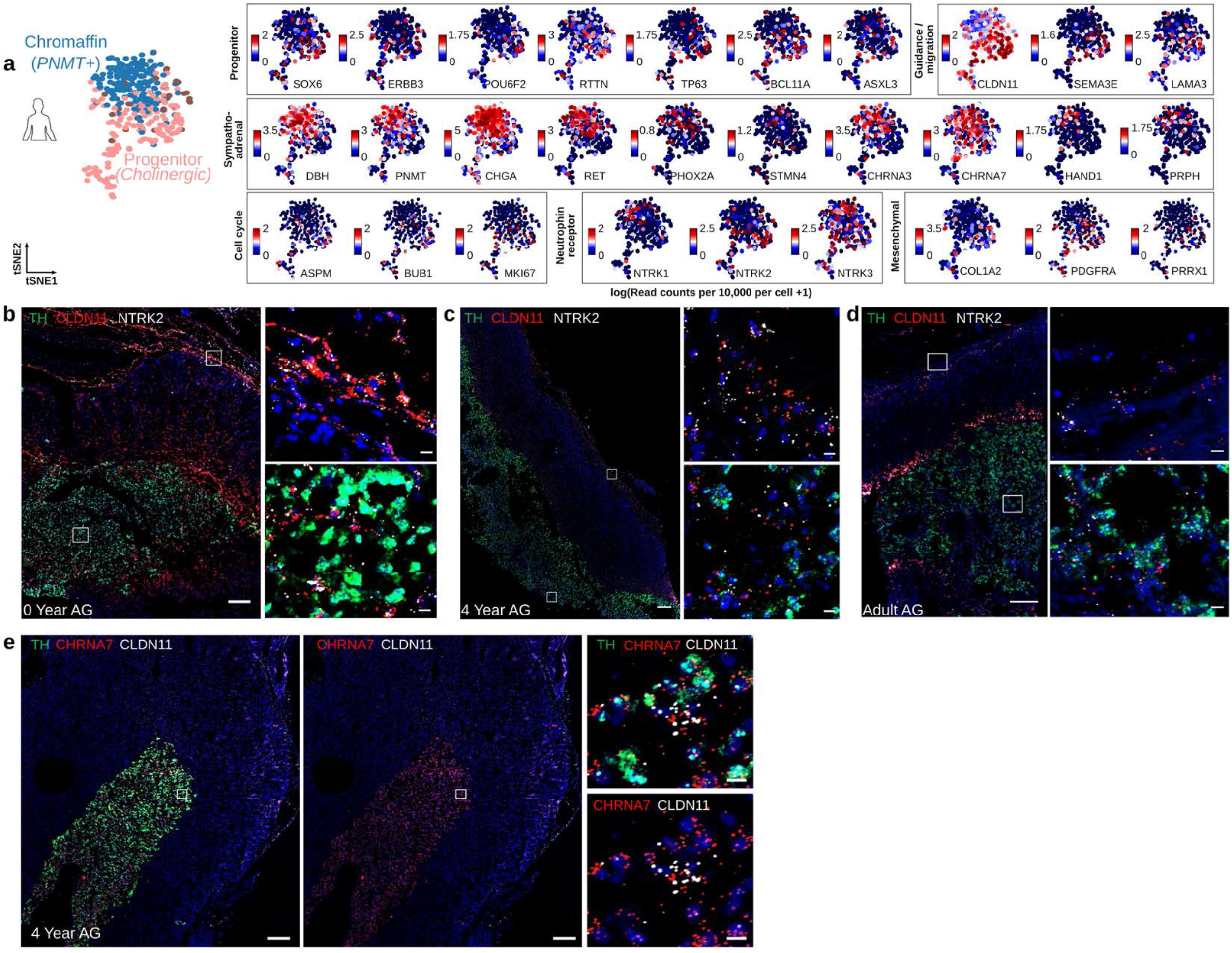
Location of human cholinergic progenitor (*NTRK2+CLDN11+*) and chromaffin (*TH+*) cells within the post-natal human adrenal gland (AG). **a**, tSNE representing the log(read counts per 10,000 per cell + 1) of indicated genes expressed in human cholinergic progenitors and chromaffin population. **b-f**, Overview of tile-scanned images (20x) of post-natal human adrenal glands (AG) at indicated age. Scalebar of overview: 200μm, zoom of boxed image: 10μm. **b-d**, RNAscope *in si/u* hybridization (ISH) for *TH* (green)*, CLDN11* (red) and *NTRK2* (white) mRNA and counter stained with DAPI (blue)*. NTRK2+ CLDN11+* double positive cells were found in adrenal capsule and medulla exclusive from *TH* positive cells. **e**, RNAscope ISH of a 4 year old AG labeled with for *TH* (green)*, CHRNA7* (red) and *CLDN11* (white) mRNA and nuclear counter-stain (DAPI) as indicated.

### Identification of a novel population of progenitor cells in the human adrenal gland

Unexpectedly, we found a population of cells unique to the human post-natal adrenal gland (hC1) which significantly expressed various progenitor and migratory markers, including *SOX6, BCL11A, ERBB3, NTRK2* (TRKB), *RTTN, PTPRZ1, TP63, ASXL3, POU6F2, SEMA3E, LAMA3, DOCK7* and *CLDN11* (*FDR*<0.01, Welch’s *t*-test, “Progenitors” and “Migrat.”, highlighted by a dotted line, Figure 1d and 2a, Supplementary Table S2 and Supplementary Dataset 1). Furthermore, hC1 progenitor population are of proliferating nature and significantly expressed cell cycling genes such as *MKI67, ASPM, BUB1* (“Cell Cycle” panel was significantly up-regulated *FDR*<0.01, Welch’s *t*-test, Figure 1d and 2a and Supplementary Dataset 1). These progenitors however did not express previously described multipotent Schwann cell precursors (SCPs) markers such as *SOX10, FOXD3* and *S100B* (Figure 1d, Supplementary Table 2). In support of this finding, previously described SCPs from mouse adrenal anlagen at embryonic day E12/E13 [6], and from human fetal adrenal glands at 8-14 post-conception weeks (PCW) [14] shared no significant specific gene signature with the human post-natal progenitor cluster hC1 (Fisher’s exact test, Supplementary Figure 2 c-d, Supplementary Table 3). No other progenitor cells than SCP, chromaffin and sympathoblast populations have been described for fetal adrenal gland [13,14]. In this regard, the only shared gene signatures with human fetal adrenal gland and mouse embryonic anlagen belonged to cell-cycle genes expressed in the cycling sympathoblast (mouse E13 and human 8-14 PCW) and cycling chromaffin (human 8-14 PCW, *FDR*<0.05, Fisher’s exact test, Supplementary Table 3). An estimation of cell velocities (i.e. computational reconstructions of cells trajectory and faiths that uses transcript splicing to calculate the direction and speed of differentiation) [18,19] suggest that this type of cells (i.e. hC1) repopulates chromaffin cells in post-natal human adrenal gland (Figure 1e). A velocity-driven gene trajectory analysis using pseudotime indicates that the progenitor cells transits from precursors cells with high differentiation potential to differentiated chromaffin cells (Figure 1f-h). Furthermore, both progenitor (hC1) and chromaffin population (hC4), express the nicotinic acetylcholine receptor nAChRα7 (*CHRNA7)*, suggesting that progenitor cells are cholinergic in nature (*FDR*<0.01, Welch’s *t*-test, Supplementary Table 2).

To validate the expression and spatial context of the human cholinergic progenitor cells (hC1), a series of RNA-scope *in situ* hybridizations (ISH) was performed in post-natal adrenal glands from 0 and 4 years old children, and adult (Figure 2b-e, Supplementary Figure 2e, Supplementary Figure 3). We first elucidated the anatomy of the glands in each section for medulla (*TH*), cortex (*CYP11B2*) and capsule (*RSPO3*) (Supplementary Figure 2e). To identify cells belonging to the hC1 progenitor population, we tested markers identified as significantly up-regulated in hC1 population: *NTRK2* and *CLDN11* (Figure 2b-d and Supplementary Figure 3a-c). *NTRK2+CLDN11+* double positive cells were found in human adrenal capsule and medulla exclusive from *TH+* cells at all ages, but most abundantly at 0 year of age, suggesting that these cells decline over age. To confirm that the post-natal progenitor *CLDN11+* cells are of cholinergic nature, we performed *in situ* RNA-hybridizations with the cholinergic nicotinic receptor *CHRNA7* (i.e. nAChR α7) together with *TH* and *CLDN11.* Both cell types, *TH+* chromaffin and *CLDN11+* progenitor cells, express nAChR α7 (Figure 2e). To further confirm that the post-natal progenitor cells in human post-natal gland are different from the previously described embryonic multipotent SCPs, we performed *in situ* RNA-hybridizations together with SCP/glia marker *SOX10* (Supplementary Figure 3d-f). *NTRK2+* cells were exclusive from *SOX10+* cells in human post-natal adrenal gland at all ages.

### Different neuroblastoma risk groups present differences in cell population composition

Next we used single-nuclei transcriptomics to characterize eleven neuroblastoma samples across different clinical risk groups, and genetic subsets (Figure 3a-b, Supplementary Table 1). Deep frozen samples obtained from surgical resections were used for nuc-Seq (SmartSeq2), yielding a total of 3,212 high-quality nuclei with an average of 709,676 high-quality reads per nuclei (see Methods and Supplementary Figure 1). Cluster analysis using PAGODA identified ten cell populations, classified as: undifferentiated (nC2, and nC3), noradrenergic (NOR clusters nC5, nC7, nC8, nC9), and stroma clusters: mesenchymal stroma (MSC nC1), endothelial (nC4), macrophages (nC6) and T-cells (nC10) (Figure 3a and Supplementary Dataset 3). The identity of each cluster was assigned by cross-referencing cluster-defining transcripts with canonical markers curated from the literature (Supplementary Table 2). Clusters nC2 and nC3 (referred as “undifferentiated”), presented a significant high expression of progenitor markers *PROM1, RTTN, ERBB3, POU6F2* and the migratory marker *CLDN11*. Remarkably, the undifferentiated nC3 clusters presented a significantly high expression of *MYCN, ALK, BRCA1* and *BRCA2* genes and progenitor markers *BCL11A, NTRK2, SOX5, SOX6, TP63, LGR5, USH2A* (*FDR*<0.01, Welch’s *t*-test, Figure 3c and Supplementary Table 2, Supplementary Dataset 3). Oppositely, the noradrenergic clusters (NOR, nC5, nC7, nC8, and nC9) expressed significantly high *NTRK1* (TRKA) and noradrenergic markers *TH, DBH, PHOX2A, PHOX2B*, and *ISL1* (*FDR*<0.01, Welch’s *t*-test, Figure 3c-d and Supplementary Table 2, Supplementary Dataset 3).

**Figure 3.**
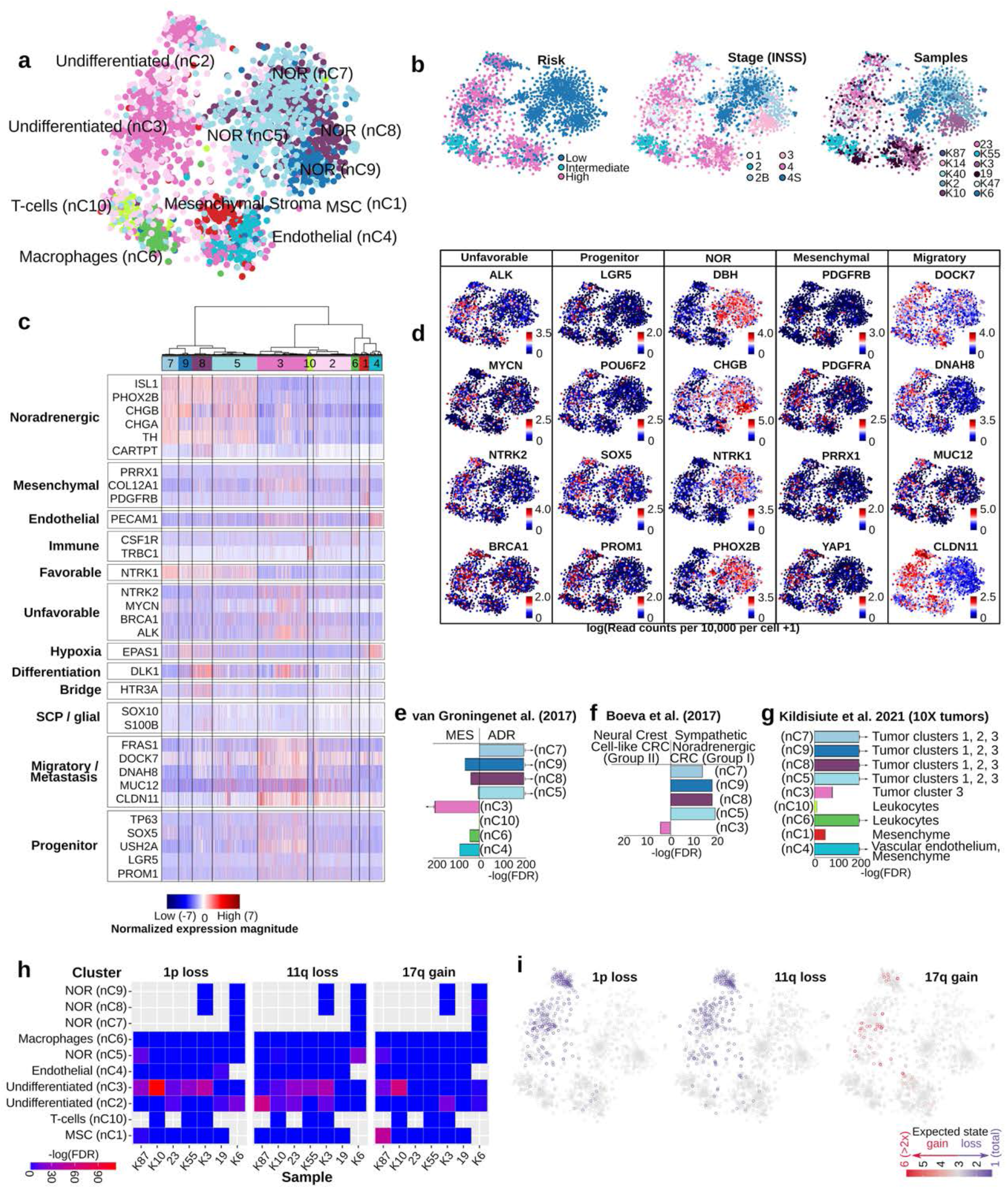
Intratumoral heterogeneity of childhood neuroblastoma (NB). 4,224 single nuclei of human NB tumors of different stages were sequenced with Smart-seq2 to an average depth of 676,000 reads per cell. Cells with high-quality (*n*=3,212 were selected and further processed with PAGODA. **a**, Cells were grouped into ten different clusters, broadly separated into two different sections (**b**): favorable NB (high *TrkA/*NTRK1, low *TrkB/*NTRK2), and non-favorable NB (low *TrkA*, high *TrkB).* Non-favorable NB included cells expressing stem cell-like genes and featured high *Mycn* expression. NB risk groups and stages are exemplified in **b**. **c**, Representation of the cell clusters: Undifferentiated (light pink nC2, and pink nC3), mesenchymal stroma (MSC red nC1), endothelial (light blue nC4), T-cells (light green nC10), macrophages (green nC6), and noradrenergic (“NOR”, blue nC5, nC7, nC8, and nC9); and various markers for cell types of interest: noradrenergic, mesenchymal, endothelial and immune, for favorable and unfavorable outcomes, and hypoxia-, and metastasis-associated (also in insert **d**). **e-f**, Using a gene enrichment-based approach, the noradrenergic clusters of NB (nC5, nC7, nC8 and nC9) proved to share a significant number of up-regulated genes with the sympathetic noradrenergic and the “ADR” (i.e. adrenergic) transcriptional signatures described by Boeva et al. [15] and van Groningen et al. [16], respectively. Oppositely, the undifferentiated cluster nC3 shared a significant number of up-regulated genes with the neural crest cell-like signature and the MES (i.e. mesenchymal) signatures described by Boeva et al. [15] and van Groningen et al. [16] (respectively).. **g**, A comparison of the gene-specific signature of the neuroblastoma clusters with the markers previously reported for six cell-clusters in 8 tumors sequenced with 10X (GOSH samples, Kildisiute et al. [13]), suggests a larger similarity of the noradrenergic clusters (nC5, nC7, nC8 and nC9) with tumor clusters 1, 2, and 3, and a stronger similarity of the undifferentiated nC3 cluster with tumor cluster 3 that exhibits a weaker sympathoblast signal [13]. Only clusters with the most significant similarities are displayed (Supplementary Table 4). **h**, An analysis of the CNVs in the different clusters indicates a significant number of cells with rearrangements (indicated by red, Fisher exact corrected with Benjamini-Hochberg for multiple tests) in the undifferentiated cluster nC3 for most samples, while a remarkable number of rearrangements was observed in fewer samples for the NOR (nC5), MSC (nC1), and undifferentiated nC2 clusters. **i**, A subset of the predicted CNVs was validated for each sample, and illustrated in the tSNE for cells with an expected rearrangement (i.e. expected state) □ overlapping the validated region.

Neuroblastoma samples of different clinical risk groups and stages contained different contribution of cells within these clusters reflecting the clinical heterogeneity of these clinical groups. High-risk cases had a significantly higher proportion of cells in stroma and undifferentiated clusters, whereas the low-risk ones including spontaneous regressing (4S) cases, consisted mostly of cells belonging to the noradrenergic (NOR) clusters (*FDR*<0.01, Chi-square test, Figure 3b, Supplementary Figure 4). In agreement with these observations, a deconvolution of 172 neuroblastoma bulk-sequenced samples (NB172 NCI TARGET project) indicated that a significantly higher proportion of cells in the stroma and undifferentiated nC3 clusters was present in high-risk patients (*n*=139), and a higher proportion of cells in the noradrenergic clusters nC7, nC8, and nC9 in low-risk cases (*n*=14, *FDR*<0.01, Chi-square test, Supplementary Figure 4). Some differences observed between single-nuclei and bulk-seq for nC2 and nC5 clusters could be a consequence of the clinical heterogeneity within patients of the same risk groups (Supplementary Figure 4).

Mesenchymal markers *PRRX1*, *YAP1*, and *PDGFRA* were significantly expressed in both clusters, undifferentiated nC3 and MSC nC1 clusters (*FDR*<0.01, Welch’s *t*-test, Figure 3c-d and Supplementary Table 2), in contrast to *PDGFRB* whose up-regulation was significantly higher in MSC cells (nC1, *FDR*<0.01, Welch’s *t*-test, Figure 3c-d and Supplementary Table 2). Consistently with this observation, the undifferentiated nC3 cluster shared a significant number of up-regulated genes with the neural crest cells-like signature (group II) and the mesenchymal (MES) gene expression signatures previously described in neuroblastoma cell lines [15,16], while the noradrenergic clusters nC5, nC7, nC8, and nC9 presented a significantly high expression of genes characterizing the sympathetic noradrenergic (group I) and ADR (adrenergic) signatures (*FDR*<0.01, Fisher’s exact test, Figure 3e-f, Supplementary Table 4). Additionally, specific gene signatures from each neuroblastoma cluster was compared with the recently described markers of six cell-clusters in eight GOSH-cohort tumors sequenced with 10X [13]. The noradrenergic clusters nC5, nC7, nC8, and nC9 resembled more significantly the GOSH sympathoblast-like tumors clusters 1, 2, and 3, whereas the undifferentiated cluster nC3 showed a higher significant resemblance with the less differentiated GOSH-tumor cluster 3 (*FDR*<0.01, Fisher’s exact test, Figure 3g, Supplementary Table 4).

To differentiate malignant-from stroma cell clusters, an analysis of the genome rearrangements was conducted, and further experimentally validated. Significant copy number changes were computed for each sample using as background the changes observed in immune cells (i.e. T-cells nC10 and macrophages nC6, only in the 7 samples with immune cells as detailed in Methods). For the majority of samples, a significantly higher number of cells presented gains in 17q, losses in 1p or in 11q, in the undifferentiated nC3 cluster (K87, K10, 23, K55, and K3). Fewer samples presented a remarkable number of cells with rearrangements in the MSC nC1 (K87), undifferentiated nC2 (23, K3, K6, and K87), and NOR nC5 clusters (K6 and K87, *FDR*<0.05, Fisher’s exact test, Figure 3h). 1p deletion was confirmed by microarrays for samples 23, K10, and K55, 11q loss for samples 23 and K87, and 17q gains for samples K10, K3, and K87. Cells with predicted significant gains and losses from these samples are illustrated in Figure 3i.

Next, we validated the expression of significantly enriched markers identified in neuroblastoma clusters (Figure 4, Supplementary Figure 5). RNA-scope *in situ* hybridizations in a *MYCN* amplified neuroblastoma case (K10, Supplementary Table 1) as independently confirmed by comparative genomic hybridization (CGH) [20], revealed a high intratumoral heterogeneity for *TH*, *MYCN* and *ALK* expression, with tumor regions with intense staining of *MYCN* and *ALK*, negative for *TH* (Figure 4a, zoom 2). Other tumor regions revealed some diffuse stainings of few *TH+* positive cells with enlarged nuclei positive for *MYCN*, but negative for *ALK* (Figure 4a, zoom 1), whereas other regions did not exhibited expression of *TH, MYCN* and *ALK* (zoom 4). Interestingly, *in situ* hybridizations for *NTRK1* and *NTRK2* (encoding TRKA and TRKB, respectively) revealed that the identified *TH+* cells with enlarged nuclei are *NTRK1+* but *NTRK2−*, in contrast to all surrounding *NTRK1-NTRK2+* cells with small nuclei (Figure 4b). It is possible that the identified *TH+* cells with enlarged nuclei (*TH+, MYCN+, NTRK1+, NTRK2−, ALK-*) might account for cells undergoing differentiation [21]. Expression of neurotrophin receptor *NTRK1* is characterized as a marker for favorable outcome and low-risk, while *NTRK2* expression is associated with poor outcome and high-risk [2]. In agreement with these findings, cells for the favorable low-risk neuroblastoma case (K6, stage 4S, Supplementary Table 1) were homogeneously *NTRK1+, TH+, NTRK2−* (Supplementary Figure 5a).

**Figure 4.**
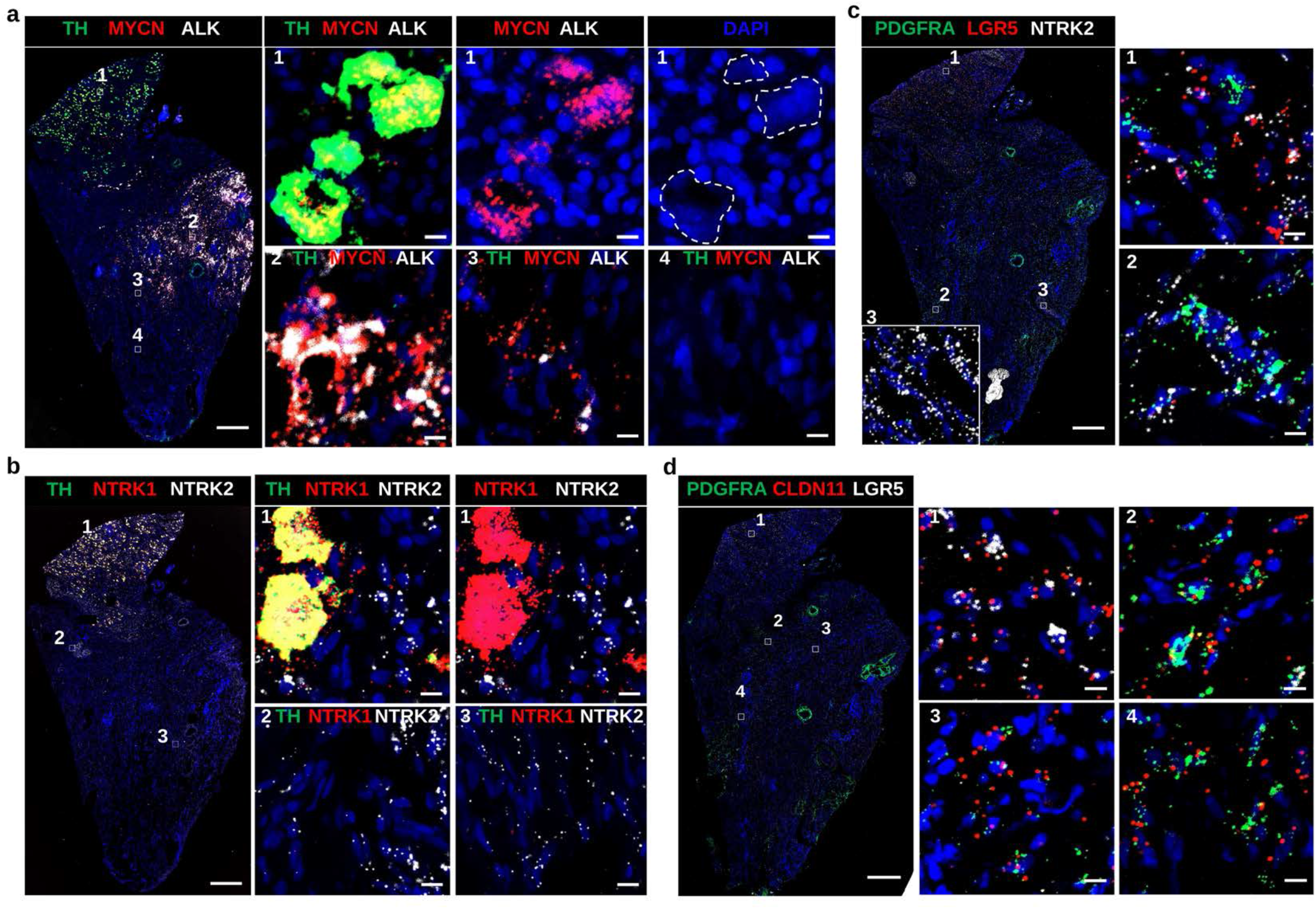
RNA *in situ* hybridization validating intratumoral heterogeneity in high-risk neuroblastoma stage 4. Overview of tile-scanned images (20x) high-risk neuroblastoma (K10, *MYCN* amplified) using RNAscope *in situ* hybridization. Scalebar of overview: 500μm; zoom of boxed image: 10μm. **a**, Tumor labeled with RNAscope ISH for *TH* (green), *MYCN* (red) and *ALK* (white) mRNA and counter stained with DAPI (blue). Dashed circles indicate cells with large nuclei in a region (box #1), that are *TH* and *MYCN* positive, but negative for *ALK.* Box #2 indicates part of the tumor with majority of cells double positive for *MYCN* and *ALK* but negative for *TH.* Box #4 indicates cells negative for all probes: *TH, MYCN* and *ALK.* **b**, Adjacent section from (a) labeled for *TH* (green), *NTRK1* (red) and *NTRK2* (white). Box #1 indicate cells with large nuclei in a region that are *TH+* and *NTRK1+*, and negative for *NTRK2.* Surrounding cells with small nuclei are positive for *NTRK2* only. Box #2,3 visualizes tumor region with majority of cells positive for *NTRK2* that are negative for *TH* and *NTRK1.* **c**, Adjacent section stained for *PDGFRA* (green), *LGR5* (red) and *NTRK2* (white) mRNA is highlighting cells double positive for *LGR5* and *NTRK2* (box #1) or double positive for *PDGRFA, NTRK2* (box #2). Box #3 indicates a region of the tumor that is positive for *NTRK2* only. **d**, Adjacent section stained for mesenchymal markers *PDGFRA* (green), *CLDN11* (red) and *LGR5* (white). Similar to **c**: some tumor regions (box #1) highlight cells double positive for *CLDN11* (red) and *LGR5* (white) that are negative for *PDGFRA* (box #1), as were other regions show cells that are double positive for *PDGFRA* (green) and *CLDN11* (red) (box #2,3 and 4).

In addition, we validated the expression of the mesenchymal marker *PDGFRA* that was identified to be significantly highly expressed in high-risk neuroblastoma cluster (nC3). *PDGFRA* expression in the high-risk *MYCN* amplified case (i.e. K10) was observed in both *NTRK2+*, (Figure 4c zoom 2), and also *in CLDN11+* cells (Figure 4d, zoom 2, 3, and 4), whereas in *LGR5+* cells showed no *PDGFRA* signal (Figure 4c zoom 1 and Figure 4d zoom 1). In contrast, the low-risk stage 4S case was entirely negative for *PDGFRA* and *PRRX1* expression, however homogeneously expressed in all cells noradrenergic markers *DBH* and *PHOX2B* (Supplementary Figure 5b,c). Stage 2B neuroblastoma was more heterogeneous in *DBH* and *PHOX2B* expression (Supplementary Figure 5d), with all cells lacking expression of *PRRX1* and very few cells positive for *PDGFRA* (Supplementary Figure 5e).

### A cell population enriched in high-risk neuroblastoma resembles the human post-natal adrenal progenitor population

High-risk neuroblastoma commonly arises in children older than 18 months and frequently originates in the adrenal glands [1]. To understand how the observed neuroblastoma populations of high and low-risk cases are related to normal post-natal and embryonic adrenal tissue, we performed a comparative analysis of their transcriptional profiles (Figures 5, 6 and 7). Significant expression of (nor-)adrenergic (“NOR”) markers (i.e. *CHGA, CHGB, DBH, TH, PHOX2A* and *PHOX2B*) is common for healthy sympatho-adrenal cells and tumor NOR populations identified in low-risk neuroblastoma cases (*FDR*<0.01, Welch’s *t*-test, Figure 5a, Supplementary Table 3). In agreement with this observation, specific gene signatures from neuroblastoma NOR clusters were found to be significantly shared with both, mouse embryonic (E12-E13) and human (8-14 PCW) chromaffin and sympathoblast clusters, and also with human and mouse post-natal chromaffin clusters (*FDR*<0.01, Fisher’s exact test, Figure 5b-e, Supplementary Table 5, Supplementary Figure 6a). A more significant resemblance was observed between the NOR nC7 and nC9 clusters with fetal (8-14 PCW) sympathoblast cells, while the NOR nC8 cluster presented a closer similitude to fetal (8-14 PCW) chromaffin and sympathoblast clusters (Figure 5e, Supplementary Table 5). A comparative analysis with the mouse embryonic transcriptional profiles showed similar significance of gene signatures shared between NOR nC7,and nC9 clusters with both, mouse embryonic chromaffin and sympathoblast populations (*FDR*<0.01, Fisher’s exact test, Figure 5b,c, Supplementary Figure 6a, Supplementary Table 5). In support of these findings, neuroblastoma NOR clusters presented a high signature score (measuring transcriptional resemblance, see Methods) for bridge, sympathoblast and chromaffin cells from in mouse and human developing adrenal glands, and for human post-natal chromaffin cells (Figure 6a-c). Furthermore, the significant specific gene signatures of NOR nC7 and nC8 clusters was significantly associated with a better outcome in a large neuroblastoma cohort (i.e. Bonferroni corrected *p*-value < 0.01, 498 cases in SEQC [17], see Methods, Figure 6d).

**Figure 5.**
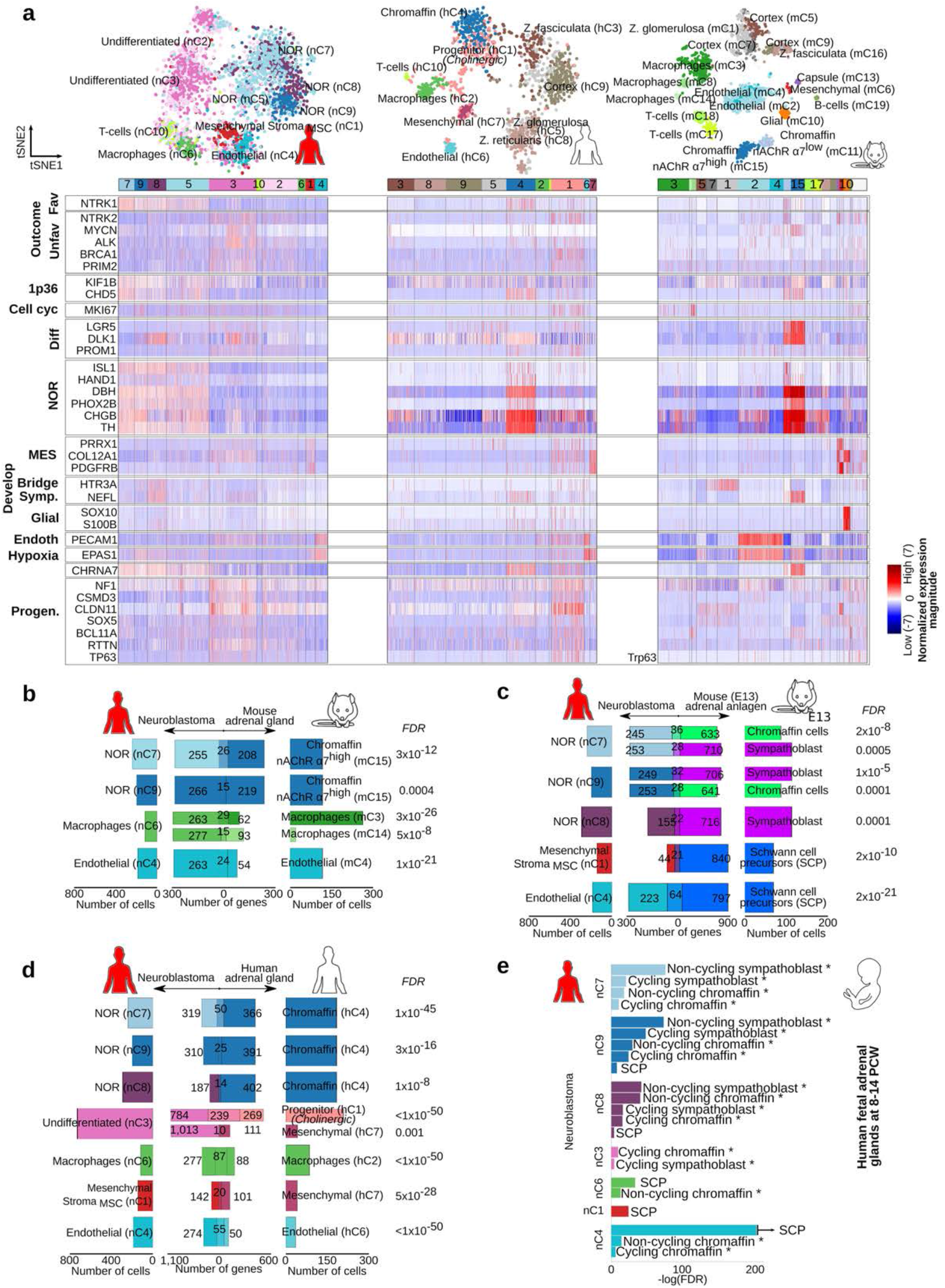
Cell clusters from neuroblastoma and healthy post-natal human and mouse adrenal glands share expression signatures. **a**, Heatmap illustrating the normalized expression magnitude for selected genes organized following the hierarchical clustering (PAGODA) as shown in Figures 1 and 3. **b-d**, Venn diagrams illustrating the significantly shared specific gene signature. Shared genes are listed in Supplementary Table 5. Indicated *FDR* were calculated with a Benjamini-Hochberg correction on *p*-values obtained with Welch’s *t*-tests as detailed in the Methods. **b**, Venn diagrams of human neuroblastoma clusters compared to adult mouse post-natal clusters, **c**, embryonic mouse clusters previously described [6], and **d**, human post-natal adrenal gland clusters. **e**, Comparison of the specific signature from the human neuroblastoma clusters and the reported transcriptional signal for human fetal adrenal glands cell clusters (Dong el al. [14]). * The original cluster annotations are currently debated and the included labels here correspond to those given by Kildisiute et al. [45], and Bedoya-Reina and Schlisio [46].

**Figure 6.**
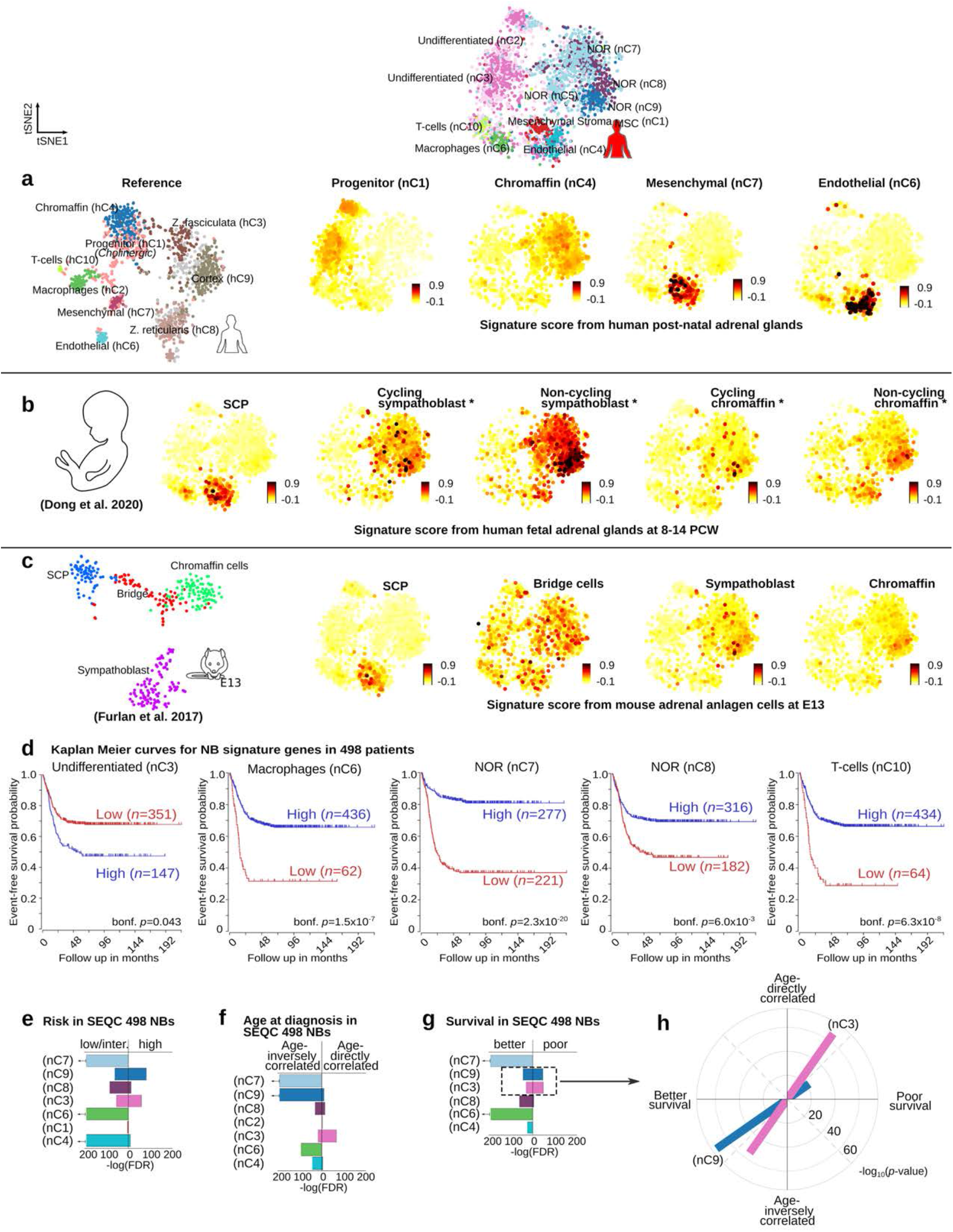
Cell clusters from neuroblastoma and developing human and mouse adrenal glands share distinct expression signatures, and their expression is associated with patient survival and age at diagnosis. tSNE of neuroblastoma clusters illustrating the signature score of genes characterizing **a**, human post-natal and **b**, embryonic [14] adrenal glands, and, **c**, mouse embryonic adrenal anlagen (E13) cell clusters [6]. Signature score = [(Reference gene set average read counts per 10,000 per cell + 1) − (Random gene set average read counts per 10,000 per cell + 1)]. SCP = refers to the signature score of genes significantly up-regulated in cluster of mouse multi-potent Schwann cell precursors at E13; Bridge = refers to the signature score of genes significantly up-regulated in cell cluster that defines transiting cells from SCP towards chromaffin population. **d**, Kaplan Meier curves for the signature genes from the neuroblastoma clusters with significant differences (Bonferroni Corrected, logrank tests) in survival for 498 SEQC neuroblastoma patients [17]. **e-g**, Using a gene enrichment-based approach (Benjamini-Hochberg corrected, Fisher’s exact tests), the specific signature genes for the noradrenergic nC7, nC8, and nC9, endothelial nC4, macrophages nC6, and undifferentiated nC3 clusters were found to be significantly up-regulated in low- and intermediate-risk patients and also in individuals with better survival. In contrast, the signature genes of noradrenergic nC9 and undifferentiated nC3 clusters presented a significantly enrichment in patients with high-risk and poor survival. The signature genes of undifferentiated nC3 cluster presented also a remarkable significant correlation with age at diagnosis. **h**, Signature genes associated with poor survival for noradrenergic nC9 and undifferentiated nC3 clusters are significantly correlated with age at diagnosis, and oppositely those associated with better survival are inversely correlated with age. Genes are included in Supplementary Table 6.

**Figure 7.**
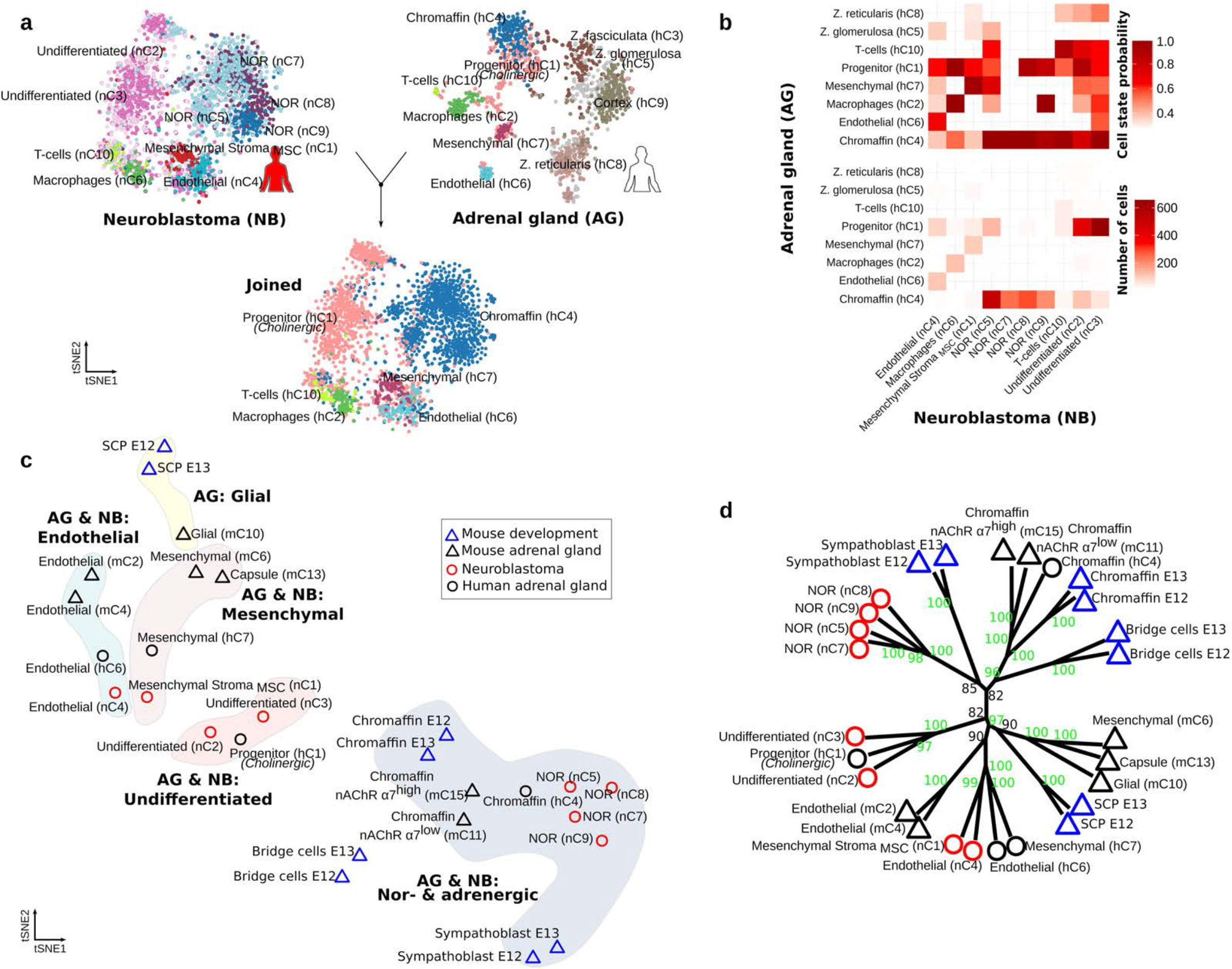
Commonalities in the transcriptional profile of cell population in post-natal and developing human and mouse adrenal gland, and neuroblastoma. **a**, tSNE plot of snRNA-seq clusters for the projection of human adrenal gland cell states (reference) onto neuroblastoma (query), based on the detection of mutual nearest neighbor cell anchors (Seurat v3). Joint visualization of same neuroblastoma cell clustering (bottom) after cell classification, colors corresponding to transferred adrenal gland cell states. **b**, Heatmap of the overall profiles of probability of best matching cell populations including number of cells between neuroblastoma and adrenal gland. **c**, tSNE using Euclidean distances in the quartile-normalized matrix of the average normalized expression from single-cell/nuclei sequenced cell populations: from (A) human i) neuroblastoma and ii) adrenal gland; and (B) mouse i) adrenal gland, and ii) derived adrenal anlagen at E12 and E13 (reported by Furlan et al. [6]). Mesenchymal, endothelial and neural-crest derived populations from human and mouse, adrenal gland, and neuroblastoma group accordingly to gene expression in i) adrenergic and noradrenergic cells, ii) undifferentiated cells, iv) mesenchymal, and v) endothelial cells. **d**, Hierarchical-clustering of transcriptional similarities between cell clusters (**c**) supported with unbiased *p*-values [41]. Green numbers in the branches indicates a high support (i.e. unbiased *p*-value >95) that has a lowest support at 0 and a highest at 100.

In contrast to the neuroblastoma NOR cells, the undifferentiated neuroblastoma clusters (nC3, nC2) did not share specific gene signatures with mouse embryonic and post-natal sympathoblast nor chromaffin populations (Figure 5b-e). The nC3 cluster however shared a marginally significant number of specific-signature genes with cycling populations in the human fetal gland (i.e. cycling sympathoblast and chromaffin, *FDR*<0.05, Fisher’s exact test, Figure 5e). These were nonetheless only cell-cycle related genes (i.e. *ASPM, BUB1, CENPE, CLSPN*, and *ESCO2*, Supplementary Table 5).

Importantly, and in contrast to the fetal adrenal gland comparison, the neuroblastoma nC3 cluster shared a specific gene signature with the human cholinergic progenitor population (hC1) identified in post-natal human adrenal gland (*FDR*<0.01, Fisher’s exact test, Figure 5d, Supplementary Table 5). Additionally, nC3 and hC1 shared a significantly high expression (*FDR*<0.01, Welch’s *t*-test) of *NTRK2*, mesenchymal (i.e. *COL1A2, COL6A3, COL12A1)*, migratory (i.e. *CLDN11, DOCK7)*, and progenitor genes (i.e. *BCL11A, ERBB3, RTTN, TP63, ASXL3, POU6F2* and *SOX6*, Supplementary Table 2, Supplementary Datasets 1 and 3). Furthermore, the undifferentiated neuroblastoma cluster (nC3) constituted a larger proportion of cells in high-risk samples in both the 11 nuc-Seq-samples and in a deconvolved cohort of 172 patients (*FDR*<0.01, Chi-square test, Supplementary Figure 4), and its specific-signature genes have a marginally higher expression average in patients with a lower survival probability in a larger cohort (i.e. Bonferroni corrected *p*-value < 0.05, 498 cases in SEQC [17], see Methods, Figure 6d).

Recently, embryonic multipotent schwann cell precursors (SCP) have been suggested to be a potential source of neuroblastoma origin [6]. However, human fetal and mouse embryonic SCPs shared only specific gene signatures with neuroblastoma stroma clusters MSC nC1 and endothelial nC4. (Figure 5c-d, Supplementary Figure 6a, Supplementary Table 5). A detailed look into their shared gene signatures suggest a high expression of genes related to cell adhesion and motility (i.e. GO term, *FDR*<0.05, Fisher’s exact test) and did not include SCP lineage markers (i.e. *SOX10, S100B, FABP7* or *PLP1*, Supplementary Table 5).

We further analyzed in detail how the transcriptional profiles of the different neuroblastoma clusters were associated with the survival and age at diagnosis in a larger cohort (i.e. 498 cases in SEQC [17], Figure 6e-h, Supplementary Table 6). We compared the specific gene signatures from each neuroblastoma cluster with genes in 498 patient samples with 1) differential expression by risk groups (Figure 6e), 2) significant correlation with age-at-diagnosis (Figure 6f,) and 3) differential expression by survival outcome (Figure 6g as detail in Methods). Genes in the specific signatures from endothelial nC4, macrophages nC6, undifferentiated nC3, and NOR clusters (nC7, nC8, and nC9) were significantly enriched in both low/intermediate risks and better survival cases (*FDR*<0.01, Fisher’s exact test, Figure 6e-g, Supplementary Table 6). Nevertheless, genes in the specific signatures from undifferentiated nC3 and NOR nC9 clusters were also significantly enriched in both high-risk and worst survival patients (*FDR*<0.01, Fisher’s exact test, Figure 6e-g, Supplementary Table 6). Remarkably, a more significant directly correlation was observed for the specific gene signature of the undifferentiated nC3 cluster with age at diagnosis (i.e. they are more likely to be expressed in later ages), while a more significant inversely correlation was found for the NOR nC9 cluster (*FDR*<0.01, Fisher’s exact test, Figure 6f, Supplementary Table 6).

A detailed analysis of the specific gene signatures for the undifferentiated nC3 and NOR nC9 clusters revealed that the two cell populations feature different gene programs that can be either 1) directly correlated with age at diagnosis and associated with poor survival (more significant in nC3 than in nC9), or otherwise 2) inversely correlated with age at diagnosis and resulting in better survival (more significant in nC9 than in nC3, *FDR*<0.01, Fisher’s exact test, Figure 6h, Supplementary Table 6). The genes inversely correlated with age at diagnosis, and also those resulting in better survival, are significantly associated with cell adhesion and differentiation for both nC3 and nC9 clusters (GO term, *FDR*<0.01, Fisher’s exact test, Supplementary Figure 6d-g, Supplementary Table 7). Oppositely, genes in NOR nC9 associated with a later age at diagnosis and a worst survival are significantly enriched in translation, RNA processing and splicing, while genes in the undifferentiated nC3 cluster associated with a worst survival are enriched in DNA damage and repair and cell cycle, and those associate with a later age of diagnosis are enriched in cell motility (GO term *FDR*<0.01, Fisher’s exact test, Supplementary Figure 6d-g, Supplementary Table 7).

We further investigated the commonalities of human post-natal adrenal gland and neuroblastoma clusters, and performed a comparative analysis based on the detection of matching mutual nearest neighbors [22,23] (Figure 7a-b). We used human adrenal gland for the projection of cell states onto neuroblastoma data (i.e. query). In agreement with the identified significant shared gene signatures (shown in Figure 5b), human post-natal chromaffin (hC4) population matched to the malignant NOR clusters (nC5, nC7, nC8, and nC9), whereas the adrenal gland progenitor population (hC1) matched the malignant undifferentiated neuroblastoma clusters nC2 and nC3 (Figure 6a-b). Populations of endothelial, macrophages and mesenchymal cells (i.e. stroma cells) in neuroblastoma have a large number of cells with a high state probability of (i.e. best transcriptional match using joined labeling) to similar populations in human adrenal gland.

Additionally, we interrogated the relationships between embryonic mouse adrenal anlagen, neuroblastoma and healthy human post-natal adrenal cell populations. We performed an Euclidean distance-based analysis of the transcriptional similarities between all populations, excluding cortex and immune cells (Figure 7c-d). This analysis suggested that these populations could be clustered into four groups, namely 1) (nor-)adrenergic, 2) undifferentiated, 3) mesenchymal, and 4) endothelial cells (Figure 7c). Consistently with the identified significant shared gene signatures (shown in Figure 5b-d), a statistical analysis of these Euclidean distances (Figure 7d) indicated that 1) SCPs, glial, endothelial and mesenchymal cells from human and mouse adult and embryonic

E12-E13 populations share a similar transcriptional profile; 2) human adrenal progenitor cluster hC1 resemble that of the neuroblastoma undifferentiated populations nC2 and nC3; and 3) chromaffin cells from human and mouse (post-natal and embryonic) have a similar transcriptional profile, that resemble the neuroblastoma noradrenergic clusters nC5, nC7, nC8 and nC9.

## Discussion

With the aim of understanding the transcriptional basis of the clinical heterogeneity in neuroblastoma, we analyzed 3,212 high-quality single cell/nuclei from eleven tumors from patients across different risk groups. We sequenced to an average of approximately 710,000 reads per nuclei/cell, and compared them to 3,456 human and mouse developing and post-natal AG cells sequenced to similar depths. While a larger number of cells could provide a more comprehensive view of the different cell types within the tumors, the large sequencing depth combined with full-length coverage offers higher sensitivity. A recent study compared the results obtained with Smart-Seq2 and 10X Genomics (10X) and concluded that the detection sensitivity was higher in Smart-seq2 than 10X. In particular, the gene capture rate of 20 cells sequenced with Smart-Seq2 is comparable to that of 1,000 cells with 10X [24]. Taking advantage of the enhanced sensitivity offered by Smart-Seq2, we consistently recovered the same populations of cells in neuroblastoma tumors of the same risk groups, and compared them with post-natal and developing mouse and human adrenal glands.

The way chromaffin cells are repopulated in the post-natal adrenal gland remains unknown. While during mouse embryonic development, multipotent Schwann cell precursors (i.e. SCPs) [6,11] and sympathoadrenal progenitors [7] can give rise to both chromaffin cells and sympathoblasts, their post-natal existence has not been demonstrated. In this study, we identified a population of cells unique to post-natal human adrenal gland that presented precursor features of chromaffin cells, namely 1) the expression of progenitor and migratory markers (i.e. *NTRK2, ERBB3, BCL11A, ASXL3, RTTN, SOX6, DOCK7, LAMA3, SEMA3E* and *CLDN11)*, 2) a high differentiation potential, and 3) a computed (by RNA velocity) directionality of the *in vivo* transitions from progenitor to chromaffin cells. Both human progenitor and chromaffin cells express the cholinergic receptor nAChRs α7 (encoded by the nicotinic acetylcholine receptor gene *CHRNA7)*, suggesting that progenitor cells are of cholinergic nature. nAChRα7 is a critical component for the cholinergic system mediating ligand-gated ion channel activation of calcium dependent signaling pathways [25], and is involved in the response to cortisol and oxidative stress [26,27]. As this post-natal progenitor population of chromaffin cells does not express the SCP markers *SOX10, S100B*, nor *FOXD3* and is not present in the mouse adrenal gland, it is possible that it is unique to humans. RNA *ISH* for markers characteristic of this progenitor population at different ages in human post-natal adrenal glands validated their existence and suggested that this population declines with age.

Comparative analysis of the transcriptional profiles of post-natal human adrenal glands and neuroblastoma tumors of different clinical risk groups revealed that the identified post-natal cholinergic progenitor population (hC1) had a remarkable transcriptional resemblance to the undifferentiated cluster (nC3) characteristic of high-risk neuroblastoma. Both populations, high-risk undifferentiated cluster (nC3) and progenitor population in post-natal gland (hC1) shared a significantly high expression of neurotrophic receptor *NTRK2* (encoding TRKB), mesenchymal (i.e. *COL1A2, COL6A3, COL12A1)*, migratory (i.e. *LAMA3, CLDN11, DOCK7)*, and progenitor genes (i.e. *BCL11A, ERBB3, RTTN, TP63, ASXL3, POU6F2* and *SOX6).* In contrast, the transcriptional profile of the clusters identified in low-risk neuroblastoma did not resemble the human progenitor population identified in post-natal gland, instead they showed a noradrenergic (NOR) transcriptional signature (i.e. *NTRK1, CHGA, CHGB, DBH, TH* and *PHOX2B*) matching the transcriptome of human and mouse post-natal chromaffin cells, as well as fetal human (8-14 PCW) and mouse embryonic (E13) sympathoblast and chromaffin populations.

Furthermore, our results indicate that the high-risk-associated cell cluster (nC3) has a transcriptional program that changes with age-at-diagnosis and is correlated with lower survival probabilities of patients. This program favors the expression of genes associated with cell motility and metastasis in older patients and with replication stress (i.e. DNA repair and cell cycle) in worst outcome cases. Oppositely, the low-risk-enriched NOR cell clusters presented transcriptional programs associated with greater survival probabilities and younger ages at diagnosis. Within these clusters, the NOR nC9 was an exception as it also showed a transcriptional program that presented a worst outcome in correlation with age. Remarkably, this program is associated with RNA-(mis-)splicing and not with replication stress. Our results suggest that at a younger age (at time of diagnosis) the expression of cell adhesion and differentiation genes from NOR-(and to less extent undifferentiated-) cells signals a better survival probability. Oppositely, a higher expression in older patients of RNA splicing genes from NOR cells, and to a larger extent of replication-stress genes in undifferentiated cells is likely to signal a worst outcome.

In this regard, low-risk neuroblastoma is common in younger children with less than 18 months of age at the time of diagnosis [1,5] and *NTRK1* expression (encoding TRKA) is a strong prognostic histological marker for favorable cases, whereas TRKB (encoded by *NTRK2*) is associated with poor outcome [2]. One possibility is that different progenitor populations source neuroblastomas at different ages, thereby accounting for the remarkable clinical heterogeneity that is coupled with age. Thus, age-at-diagnosis is a strong outcome predictor. Specifically, favorable neuroblastomas might originate from embryonic developmental errors when SCPs differentiate to chromaffin and neuroblast, whereas unfavorable neuroblastomas would arise from errors during post-natal development when TRKB+ cholinergic progenitors repopulate chromaffin cells post-natally.

## Acknowledgements

The single-cell transcriptome data was generated at the Eukaryotic Single-cell Genomics facility at Science for Life Laboratory in Stockholm, Sweden. The computations and data handling for this project were performed on resources provided by the Swedish National Infrastructure for Computing (SNIC) at sllstore2017016, sens2018122, SNIC 2019/35-3, SNIC 2019/3-462, SNIC 2020/5-457, SNIC 2020/6-171, and SNIC 2020/6-172, partially funded by the Swedish Research Council through grant agreements no. 2016-07213 and 2018-05973. We thank Lotta Elfman and PJ Svensson for help in providing neuroblastoma samples. We thank the NIH NeuroBioBank providing human adrenal glands. We thank Marie Arsenian-Henriksson and also members of the Schlisio and Holmberg groups for insightful comments on this research. We also thank Yao Shi for his support with the analysis of the 498 NB SEQC cohort. Funding: S.S. was funded by the Swedish Research Council, Swedish Childhood Cancer Fund, the Swedish Cancer Society, Knut and Alice Wallenberg Foundation, ParaDiff foundation and ERC Synergy grant (KILL-OR-DIFFERENTIAT). P.Ko. was funded by the Swedish Research Council, the Swedish Childhood Cancer Fund, and the Swedish Foundation for Strategic Research. I.A. was supported by ERC Consolidator grant (STEMMING-FROM-NERVE), Swedish Research Council, Paradifference Foundation, Bertil Hallsten Research Foundation, Cancer Foundation in Sweden, Knut and Alice Wallenberg Foundation. P.V.K. was founded by the NSF-14-532-CAREER grant.

## Author contributions

Conceptualization, S.S.; Investigation, O.C.B.-R., W.L., M.A., M.P., J.H., I.A., M.K. and S.S.; Validation, O.C.B.-R., W.L., M.A., M.P., S.S. Computational investigation and analysis, O.C.B.-R. and M.A.; Sample collection and characterization, C.L., C.C.J., T.M., P.Ko., M.K.; Writing O.C.B.-R. and S.S. Funding Acquisition, Resources & Supervision, O.C.B.-R. and S.S. Competing interests: none.

## Methodology

### Sample collection and single cell/nuclei sequencing

Eleven neuroblastoma samples were collected from patients at different ages (0-79 months at diagnosis) and their INSS stages determined following Shimada’s criteria. In particular, three samples were classified as favorable INSS stages (one stage 1, two stage 2/2B), two as favorable widespread stage (4S) and six in intermediate/high metastatic stages (two stage 3, and four stage 4) [4]. According to the INRG clinical and biological criteria five patients were classified as high-risk, one as intermediate risk and five as low-risk, and treated accordingly to local and international protocols [3]. Tumors were genotyped according to [20], five high-risk genotype *MYCN*-amplified and/or 11q-deleted, and six low-risk genotype Numerical Only or Other Structural (Supplementary Table 1). In addition, three human and five mouse adrenal glands were obtained. All human adrenal samples were collected in conjunction to disease unrelated to pheochromocytoma, paraganglioma and neuroblastoma. Further details about these samples are included in Supplementary Table 1.

Human samples were collected following surgical resection at the Karolinska University Hospital under the ethical permits from Stockholm Regional Ethical Review Board and the Karolinska University Hospital Research Ethics Committee (2009/1369-31/1 and 03-736) and KI 2007/069 and KI 2001/136, issued to Professor Per Kogner and Dr. C. Christofer Juhlin for neuroblastoma and adrenal gland samples, respectively. Additional post-mortem human adrenal glands for staining’s were obtained from the NIH Neurobiobank (University of Maryland, Baltimore, MD) under the same ethical permit from Stockholm Regional Ethical Review Board and the Karolinska University Hospital Research Ethics Committee. All samples were obtained following an informed patient consent. Samples were processed following the Nuc-Seq protocol [28] as illustrated in Supplementary Figure 1. Briefly, nuclei are obtained from deep frozen tissue after homogenization and filtration, and further FACs sorted in 384-wells plates where cDNA synthesis was conducted with Smart-seq2 [29]. Libraries were prepared using the Tn5 transposase tagmentation (Nextera XT), and their quality was assessed with fragment analyzer. High quality libraries were sequenced using Illumina HiSeq 2500, and further de-multiplexed with deindexer (https://github.com/ws6/deindexer) using the Nextera index adapters and the 384-well layout.

Mouse samples were collected via surgical resection after perfusion under the ethical permit N32/14 issued by Jordbruksverket in accordance with the declaration of Helsinki. The tissue from each sample was dissociated and sorted in 384-well plates. Sorted cells were lysed and their RNA obtained using the Smart-seq2 protocol [29], and further processed as described above.

### Read mapping, quality control, and cell clustering

We planned an experimental design that allowed us to obtain and analyze high quality data [30]. To select high quality reads we allowed for a hard-clipping of adaptors (using CutAdapt 1.13), and excluded reads with less than 20 bases. Additional diagnostics on the reads quality were conducted with FASTAQC (https://www.bioinformatics.babraham.ac.uk/projects/fastqc), cells with reads failing at three or more quality control (QC) tests (among eleven in total including per base- and sequence quality scores, frequency of each nucleotide, GC content, over-represented motifs, and number of duplicated reads) were excluded for further analysis. High quality reads were mapped with STAR [31] using 2-pass alignment to have improved performance of *de novo* splice junction reads.

High quality reads from human samples were mapped to the hg38 human genome version hg38/GRCh38.p12 (released 2019), annotated by the comprehensive annotation of GENCODE 28 [32] as obtained from the UCSC browser [33] on 25.02.2019. Reads from mouse were mapped to the mm10 mouse genome version mm10/GRCm38.p6 (released 2019), annotated by the comprehensive annotation of GENCODE 18 [32] obtained from the UCSC browser [33] on 14.01.2019. In both cases, alternative chromosomes were excluded from the annotations, and in the cases of human reads mapping to the mitochondrial genome were excluded (as mitochondrial reads in nuc-Seq can originate from different cells). Gene expression (i.e. read counts allowing ambiguity) was calculated using HTSeq [34].

To conduct a QC of the cells, different technical and biological features were calculated with QoRT [35] and Celloline [36]. After an inspection of the features, cell for human adrenal glands expressing at least 3,000 and at most 9,000 genes, and from mouse adrenals and neuroblastoma expressing at least 2,000 and at most 8,000 genes were selected for further analysis.

### Expression analysis

Gene expression of cells/nuclei was filtered, transformed, scaled and standardized accounting for sequence depth, using PAGODA [37]. In particular error models for individual cells were fitted with parameters robust for noisy data (i.e. min.count.threshold of 2 and at least 5 non-failed measurements per gene). These models were successfully generated for all high-quality cells from mouse (*n*=1,763) and human (*n*=1,322) adrenal glands, and (*n*=3,212) in neuroblastoma. PAGODA clusters cells based on non-redundant significant aspects of transcriptional heterogeneity. The resulting hierarchical clustering was cut by height initially to maximize the silhouette value [38], and further adjusted to increase the resolution of the defined clusters (i.e. cell/nuclei populations). The standardized expression for mouse E12 and E13 adrenal anlagen (i.e. expression magnitude) was obtained from the PAGODA rds application from [6] kindly provided by the authors.

The standardized gene expression from PAGODA (i.e. expression magnitude) was used to determine the genes (A) significantly up-regulated in each cluster, and (B) significantly and uniquely up-regulated in each cluster (i.e. specific gene signature). In particular, for a given population, the group of genes significantly up-regulated in each cluster (i.e. A) is determined as the genes whose average expression is significantly higher than the average expression of cells in all the other clusters. In contrast, the group of genes significantly and uniquely up-regulated in each cluster (i.e. B: specific gene signature) is defined as the group of genes whose average expression is significantly higher than each and all of the other clusters’ averages. This significance was calculated using Benjamini-Hochberg multiple test correction on Welch’s *t*-test generated *p*-values, with a *FDR* threshold of 0.01.

Gene expression in specific clusters was validated using RNA-scope. Fresh frozen samples were prepared following the manufacturer (ACD Bio) instructions, and the RNA from target genes labeled with the Multiplex Fluorescent Reagent Kit v2.

### Cell cluster analysis

We calculated the under- and over-representation of cells in clusters for neuroblastoma samples, INSS stages, and risk groups, with two-sided Chi-square tests using Yates adjustment and corrected with the Benjamini-Hochberg approach and a *FDR* threshold of 0.01. In particular, Chi-square tests evaluated the hypothesis that a given case study (risk group for instance) presented a significantly larger or lower proportion of cells in a given cell cluster.

The enrichment of gene ontology (GO) terms was calculated for the significantly up-regulated genes in each cluster using a Benjamini-Hochberg multiple test correction on Fisher’s exact test *p* - values. The same approach was taken to determine the enrichment of significantly up-regulated genes defining 1) mesenchymal (MES) and adrenergic (ADR) cell types by [16], and 2) Sympathetic Noradrenergic (Group 1) and Neural crest cell-like (Group 2) by [15]. In particular, Fisher’s exact tests evaluated the hypothesis that genes significantly up-regulated in a given cell cluster were significantly enriched in a gene set of interest. To conduct inter-species comparisons, only the 1:1 human:mouse orthologues annotated by ENSEMBL version GRCh38.p12 [39] were considered.

To compare the clusters among the various study cases, four different approaches were taken. In the first approach a comparison of specific gene signatures was conducted following the gene enrichment approach described above, using a *FDR* threshold of 0.01 and a minimum number of shared genes of 10 (marginally significant results with *FDR*<0.05 are included in the supplementary tables). In the second approach (as displayed in Figure 7), the average standardized expression (i.e. expression magnitude calculated with PAGODA) from each population was computed and further quantile-normalized using the limma package [40]. Further, Euclidean pairwise distances were calculated for cell populations using their quantile-normalized average gene expression values, and a hierarchical clustering was conducted using Ward’s-2 distance with hclust package and supported with the approximately unbiased *p*-value from the pvclust package [41]. In addition a tSNE was built with the Rtsne package (https://github.com/jkrijthe/Rtsne).

For the third and fourth approaches the raw gene counts were filtered, normalized and rescaled using Satija’s pipeline [42]. With these values, we conducted as a third approach an asymmetrical cluster imputation of NB cells using as reference human AG-cells with Seurat version 3 [43]. In particular, genes expressed in more than 5 cells, expressed in at least 2 genes were selected. Further processing was conducted using the top 2,000 most variable genes after variance transformation and dimension reduction. Finally, as a fourth approach we calculated a signature score for reference specific gene signatures (obtained as described above) using Scanpy [44]. This signature score calculates the average expression of the reference genes in the specific gene signature per 10,000 per cell + 1, minus the average of a (*n*) randomly selected set of genes per 10,000 per cell + 1 (where *n* = max [# reference genes, 50]). The signature score provides an estimation of the transcriptional resemblance of each cell to each reference specific gene signature.

The similarity between clusters of 1) post-natal and fetal [14] human adrenal glands, and 2) neuroblastoma in this study and from the GOSH cohort [13], were estimated by two different approaches: 1) computing the enrichment of genes in the specific signatures for genes in the reference clusters (i.e. fetal adrenal gland and GOSH cohort), and 2) computing the signature score for the same references. Gene enrichments were computed with Benjamini-Hochberg corrected Fisher’s exact tests. Reference gene sets for human fetal adrenal gland were obtained from the differentially expressed genes (with an adjusted *p* < 0.01) in five sympathoadrenal cell types in fetal adrenal glands (i.e. SCP, cycling and non-cycling sympathoblast/chomaffin cells) reported in [14]. The original cluster annotations are debated and the included labels in this study correspond to those given by Kildisiute et al. [45], and Bedoya-Reina and Schlisio [46]. Reference gene sets for the GOSH cohort were obtained from the algorithmc markers of different cell population in eight neuroblastoma tumors sequenced by 10X and reported in [13].

### Analysis of bulk RNA-seq neuroblastoma samples

For gene sets of interest (i.e. target genes), Kaplan-Meier curves were calculated using the 498 SEQC database [17] and the tools available in the R2 database (http://r2.amc.nl). Specifically, the average standardized expression (*z*-score) for target genes in the 498 SEQC database was calculated, and further classified into two patient groups using the “scan modus” in the KaplanScanner tool. The event free-survival probability was estimated with a logrank test between these two groups and corrected with Bonferroni for multiple tests. Only specific gene signatures with more than two genes were studied (i.e. specific gene signature from NOR nC5 cluster was excluded as *n*=2).

Sets of genes significantly 1) up-regulated in patients classified in high-(*n=*6,696) or non-high-(i.e. low/intermediate, *n*=8,158) risk groups, and 2) directly (*n*=4,572) or inversely (*n*=3,982) correlated with age at diagnosis, were obtained with the GeneSelector tool in R2 for the 498 SEQC database. RPMs were log2-transformed and the difference between risk groups was calculated for each gene using FDR-corrected ANOVAs. Pearson correlations were computed between gene expression and age at diagnosis with FDR-corrected *p*-values signaling the chances of obtaining the correlation coefficient in an uncorrelated dataset. Similarly, gene sets with a significantly higher expression in poor (i.e. worst, *n*=9,049) or better (*n*=5,842) survival cases were retrieved with the Kaplan Meier Scaner Pro tool, using FDR-correct *p*-values obtained with logrank tests between groups of patients with different event free-survival probabilities. Significant genes were selected with a *FDR* threshold of 0.01. Gene enrichment were further calculated with Fisher’s exact tests corrected with the Benjamini-Hochberg approach.

To calculate the proportion of cells of each neuroblastoma cluster in a larger cohort, 172 TARGET neuroblastoma samples (National Cancer Institute TARGET, dbGap Study Accession: phs000218.v16.p6.) were deconvolved using the reference-based decomposition model in BisqueRNA with the default settings [47]. In comparison with other popular methods, deconvolution by this BisqueRNA has been shown to produce the closest mean estimations to the cell proportions obtained by single nuclei-RNA [47]. Bulk expression from each sample was decomposed into the 10 neuroblastoma cell types using as reference 42,215 genes present in the annotation of both bulk and single-nuclei, filtering 1,676 zero-variance and 383 unexpressed genes. To make an accurate deconvolution, bulk-sequence raw data obtained for the 172 TARGET neuroblastoma samples were pre-processed and quantified in a similar way as the single-nuclei data. Briefly, reads were hard-clipped for adapters and low quality calls, and reads with less than 20 bases were excluded. High quality pair-reads were mapped with STAR [31] using 2-pass alignments to the hg38 human genome version hg38/GRCh38.p12 (released 2019) annotated with the comprehensive annotation of GENCODE 28 [32], and further quantified with HTSeq [34]. The expected (i.e. proportional) number of cells per cluster for each deconvolved sample was computed as the product of the predicted cluster percentage, and the proportional number of cells (*c*) in the TARGET NB samples (*c*=[average number of nuclei from 11 nuc-Seq neuroblastomas in the same risk group x number of TARGET NB samples in the risk group]). Significance of the expected (i.e. proportional) number of cells was computed with Benjamini-Hochberg corrected Chi-squares tests with Yates’ adjustment.

### Genome rearrangement analysis

Genome rearrangements of cells (i.e. copy number variants) were determined using inferCNV v1.4.0 of the Trinity CTAT Project (https://github.com/broadinstitute/inferCNV). InferCNV was computed with the gene expression for 7 samples (out of the initial 11: K87, K10, 23, K55, K3, 19, and K6) that presented cells in the immune clusters: macrophages (nC6) and/or T-cells (nC10). Arrangements in these cells were used as a non-malignant reference. Copy number variations (CNVs) were calculated from moving averages of 101-genes’ windows with the default hidden Markov model. Tumor cells were simulated to have CNVs with probabilities (*p*) that matched six possible states, from a chromosomal segment complete loss to a gain of more than 2 copies (1=complete loss, 2=loss of one copy, 3=neutral, 4=addition of one copy, 5=addition of 2 copies, 6=addition of more than 2 copies). These probabilities were used to estimate the likelihood of observing a rearrangement (total sum of *p*) and to select the most likely state (maximum *p*). CNVs with a (Bayesian) likelihood value of 0.1 (probability of being true rearrangements greater than 0.9) were selected for further analysis.

For each sample independently, rearrangements (i.e. expected states) in each region of interest (i.e. 1p loss, 11q loss, and 17q gain) were computed for each cell as the average of the most likely state (as either gain or loss) for the overlapping CNVs. The significance for gains or losses in each cluster was calculated by testing the hypothesis that the number of cells with the rearrangement was significantly higher than expected, using a Benjamini-Hochberg corrected Fisher’s exact test. Rearrangements for tumor samples were confirmed using Affymetrix 250K microarrays as previous indicated in [20]. In particular, microarray data was processed using GDAS software (Affymetrix) and CNVs were estimated using CNAG v3.0 (http://www.genome.umin.jp).

### Cell velocities and differentiation potential analysis

To calculate the differentiation trajectory of cells, the percentage of splicing and unspliced genes per cell was estimated with velocyto [18]. Further, the cell velocities were calculated using scVelo [18,19]. To obtain high confidence velocities (as confirmed by visual inspection of confidence scores) for human adrenal gland we selected genes with a minimum number of shared read counts of 40, and built the velocities using the top 600 genes. For mouse adrenal gland, these parameters were 3,000 and 1,000, respectively. Further, the gene transition over the pseudotime (representing the cell’s internal clock, and the approximate time of cell differentiation) was determined using a multiple kinetic regimes, and illustrated for the selected populations of interest. The differentiation potential of cells was determined with Palantir [48] using the default parameters, and as starting cell the one with the highest expression of the precursor marker *ERBB3*. This approach uses entropy to estimate cell plasticity in modeled trajectories of cells fates, so that cell plasticity increases with entropy.

All data generated during and/or analyzed during the current study are available in the manuscript and/or the Supplementary Materials, or otherwise available from the corresponding authors on reasonable request. Raw sequences for human adrenal gland were stored in the Synapse ID projectsyn22301662 (https://www.synapse.org/#!Synapse:syn22302430), with the following ID numbers: syn22302285 and syn25189163 (sample 6657), syn22301836 (sample 6435), and syn22301667 (sample 16-D). Raw sequences for neuroblastoma were stored in the Synapse ID projectsyn22302605 (https://www.synapse.org/#!Synapse:syn22310692), with the following ID numbers: syn22307346 (sample K87), syn22306928 (sample K6), syn22306349(sample K55), syn22305928 (sample K47), syn22305408 (sample K40), syn22304935(sample K3), syn22304482 (sample K2), syn22304038 (sample K14), syn22303649(sample K10), syn22303256 (sample 23), and syn22302820 (sample 19). Raw sequences for mouse adrenal gland were stored in the Synapse ID project syn22308005(https://www.synapse.org/#!Synapse:syn22310690), with the following ID numbers: syn22308008 (sample 6801), syn22308616 (sample 6802), syn22309301 (sample 3431), syn22310231 (sample 3432), and syn22309843 (sample I1). RNA-sequences from neuroblastoma available at: National Cancer Institute TARGET, dbGap Study Accession: phs000218.v16.p6. Similarly, the code used for the current study is available in https://github.com/oscarcbr/single_cell_tools or otherwise available from the corresponding authors on reasonable request to use, redistribute it and/or modify under the terms of the GNU General Public License as published by the Free Software Foundation; either version 2 of the License, or any later version.

---

**Supplementary Figure 1. a-b**, Overview of the pipeline used to sample, sequence and analyze single nuclei/cell from mouse and human adrenal glands (AG) and neuroblastoma (NB). **a**, Samples were collected and processed following sc-Seq [29] and the nuc-Seq protocols [28]. Nuclei (for fresh frozen human samples) and whole cells (for mouse samples) were obtained and FACs-sorted in 384-wells plates. Libraries were prepared with Smart-Seq2 and sequenced with Illumina HiSeq 2500. **b**, High-quality reads and cells were selected for further analysis with PAGODA. After clustering, gene expression for each cluster was determined and further compared with other case studies and reference databases using four different approaches detailed in Methods. Different technical and biological features characterize single(-whole cell, i.e. sc-Seq), and single(-nucleus, i.e. nuc-Seq) sequencing. **c**, Features associated to partial transcript splicing and transcription noise are higher in nuc-Seq than in sc-Seq, including the percentage of intronic, indels in read alignments, and intergenic reads. Oppositely, features associated to (cytoplasmic) mature transcripts are higher in sc-Seq than in nuc-Seq, including percentage of intragenic and exonic reads.

**Supplementary Figure 2. a-b**, Chromaffin cells in mouse are grouped in two different clusters characterized by the low and high expression of nAChR α7 (encoded by *Chrna7*). **a**, Velocity analysis suggests that each group has a different faith and do not inter-convert between them. **b**, A panel of noradrenergic markers is shared between the two population, and others characterize each cluster, particularly the expression in *Pnmt* is higher in mC11 in comparison to mC15 (*FDR*<0.01, Welch’s *t*-test). Other genes that have a higher (non-significant) average expression include *Cart/t, Isl1*, and *Hand1.* Oppositely, the expression of *Epas1, Cox8b, Ndufa4l2, Phox2b* and *Neurod4* is significantly higher (*FDR*<0.01) in mC15 than in mC11. The expression of a repertoire of Muscarinic and Nicotinic cholinergic receptors is higher in mC15 than mC11 (*FDR*<0.01, Welch’s *t*-test). **c-d**, Specific signatures significantly shared between human adrenal gland and mouse adrenal anlagen at (**c**) E13 and (**d**) E12 (*FDR*<0.01, Welch’s *t*-test, marginally significant results are included in Supplementary Table 3). **e**, Overview of tile-scanned images (20x) of post-natal human adrenal glands (AG) at indicated age. Scalebar of overview: 200μm, zoom of boxed image (indicating capsule): 10μm. RNAscope *in situ* hybridization for *TH* (green) labeling adrenal medulla, *RSPO3* (red) labeling adrenal capsule and *CYP11B2* (white) labeling adrenal cortex. Nuclei were counter-stained with DAPI (blue).

**Supplementary Figure 3. a-c**, RNAscope ISH zoom images of AG of indicated age as shown in Figure 2 without *TH* (green) channel. *NTRK2* mRNA is shown in white and *CLDN11* mRNA is shown in red. Scalebar: 10μm. **d-f**, Overview images of tile-scanned (20x) post-natal human AG at indicated age. Scalebar of overview: 200μm, zoom of boxed image: 10μm. Adrenal medulla labeled with RNAscope *in situ* hybridization for *TH* (green), *SOX10* (red) and *NTRK2* (white) mRNA and counter stained with DAPI (blue). *NTRK2+* positive cells were exclusive from *SOX10+* cells.

**Supplementary Figure 4.** Different cell populations are differently represented in neuroblastoma risks groups and INSS stages. In particular, clusters of undifferentiated and stroma cell clusters in NB (i.e. nC1, nC3 and nC4) represent a larger proportion of cells in high-risk neuroblastoma, while noradrenergic clusters (i.e. nC7, nC8, and nC9) represent a larger proportion of cells in low-risk neuroblastoma (top right). 172 TARGET NB bulk-sequenced samples (NB172 NCI TARGET project) in different risks groups were deconvolved to estimate the expected number of cells from each NB cell clusters (top left). A significant higher number of cells (Benjamini-Hochberg corrected Chi-square tests) was recapitulated in high-risks samples for clusters MSC nC1, Undifferentiated nC3, and Endothelial nC4; and in low-risk for NOR clusters nC7, nC8, and nC9. Cell numbers in deconvolution correspond to the product of the predicted proportion of cells for each cluster, and the proportional number of cells in the TARGET NB samples ([average number of single-nuclei in risk group] x [number of TARGET NB samples in the risk group]).

**Supplementary Figure 5. *In situ* transcriptomic images of favorable neuroblastoma 4S and stage 2B.** Overview of tile-scanned (20x) favorable neuroblastoma after RNAscope ISH. Scalebar of overview: 200μm; zoom of boxed image: 10μm. **a**-**c**, RNAscope ISH of low-risk INSS stage 4S neuroblastoma (**a**) for *NTRK1* (red), *NTRK2* (white) and *TH* (green) revealing homogeneous expression of *NTRK1* and *TH* mRNA in the of entire tumor section with no evidence of *NTRK2* positive cells. **b**, RNAscope ISH for *PDGFRA* (green), *CLDN11*(red) and *LGR5* (white) reveals no expression of these mRNAs. **c**, RNAscope ISH for *PRRX1* (green), *DBH* (red) and *PHOX2B* (white) showing homogeneous expression of *DBH* and *PHOX2B* mRNA in the entire tumor with no evidence of *PRRX1* positive cells. **d**, RNAscope ISH in low-risk INSS stage 2B neuroblastoma revealing a more heterogeneous pattern of *DBH* (red) and *PHOX2B* (white) positive tumor region with no evidence of *PRRX1 (green*) labeled cells. **e**, RNAscope ISH for *PDGFRA* (green), *CLDN11* (red) and *LGR5* (white) in adjacent section of (**d**) with no evidence of their expression. Only one small region of the entire tumor (box #4) showed positively labeled cells.

**Supplementary Figure 6. a**, Specific genes signatures significantly shared between neuroblastoma and mouse adrenal anlagen at E12. Kaplan Meier curves for genes with significant differences (Bonferroni corrected, logrank tests) in survival for 498 SEQC neuroblastoma patients [17] for **b-c**, signature genes from the neuroblastoma undifferentiated nC3 and NOR nC9 clusters, directly and inversely correlated with age at diagnosis in 498 SEQC neuroblastoma patients [17]. A gene enrichment-based approach (Benjamini-Hochberg corrected, Fisher’s exact tests) of the specific signature genes for the NOR nC9, and undifferentiated nC3 clusters in 498 SEQC neuroblastoma patients [17], indicates different biological processes (GO) associated with **d-e**, survival and **f-g**, age-at-diagnosis. At most the top ten GO terms with *FDR<0.01* are shown. ⋂ symbol signifies the intersection between the two gene sets.

**Supplementary Table 1.** Samples, cells and reads details and statistics for each case study, before and after quality controls.

**Supplementary Table 2.** List of cured markers from literature used to annotate cell populations in mouse and human adrenal glands (AG). Only genes with a significantly high expression in any cluster “*x*” (i.e. FDR(*s*C*x*>*s*Co)≤0.01) are included in the table. FDR(*s*C*x*>*s*Co) and FDR(*s*C*x*>*s*C*y*) test the hypotheses that the expression of a gene in cell cluster “*x*” is higher than in cells from all other clusters (“Each cluster in *s*: FDR” column), and that in cluster “*y*” (“Each cluster in pairwise comparison *s*: FDR” column), respectively, for a given case study “*s*”. FDRs were calculated with a Benjamini-Hochberg correction on Welch’s *t*-Tests (as detailed in Methods). Clusters with FDR(*s*C*x*>*s*Co)≤0.01 are included in the column “Evidence for high expression in cluster”. All clusters *y* in this column that do not present in pairwise comparisons any FDR(*s*C*x*>*s*C*y*)≤0.01 are included in the column “Evidence for higher expression in cluster”. List of specific gene signatures for clusters in human and mouse post-natal adrenal gland, and neuroblastoma calculated as detailed in Methods.

**Supplementary Table 3.** Cell populations in human post-natal adrenal glands sharing a significant specific gene signature with 1) mouse post-natal-, 2) human fetal adrenal glands [14], and 3) developing (E13) mouse [6], as detailed in Methods (*FDR*<0.05, Fisher’s exact test).

**Supplementary Table 4.** Gene enrichment analysis for neuroblastoma and selected study cases, namely van Groningen et al. [16], and Boeva et al. [15], and cell populations in neuroblastoma sharing a significant gene specific signature with markers in GOSH neuroblastoma (10X-sequenced) cell clusters [13], as detailed in methods (*FDR*<0.05, Fisher’s exact test).

**Supplementary Table 5.** Cell populations in neuroblastoma sharing a significant specific gene signature with 1) mouse post-natal-, 2) developing (E13) mouse [6], 3) human post-natal-, and 4) human fetal-[14] adrenal glands, as detailed in Methods (*FDR*<0.05, Fisher’s exact test).

**Supplementary Table 6.** Genes in the specific signature of neuroblastoma cell populations significantly enriched in different 1) risk- and 2) survival groups, and 3) significantly correlated with age at diagnosis, as detailed in methods (*FDR*<0.05, Fisher’s exact test).

**Supplementary Table 7.** Biological processes (GO) enrichment for genes in the specific signature of neuroblastoma cell populations 1) significantly enriched in different survival groups, and 2) significantly correlated with age at diagnosis, as detailed in methods (*FDR*<0.05, Fisher’s exact test).

