## Supplementary figures and images for "Single-nuclei transcriptomes from human adrenal gland reveals distinct cellular identities of low and high-risk neuroblastoma tumors"

### Supplementary Figures

Suppl. figure 1

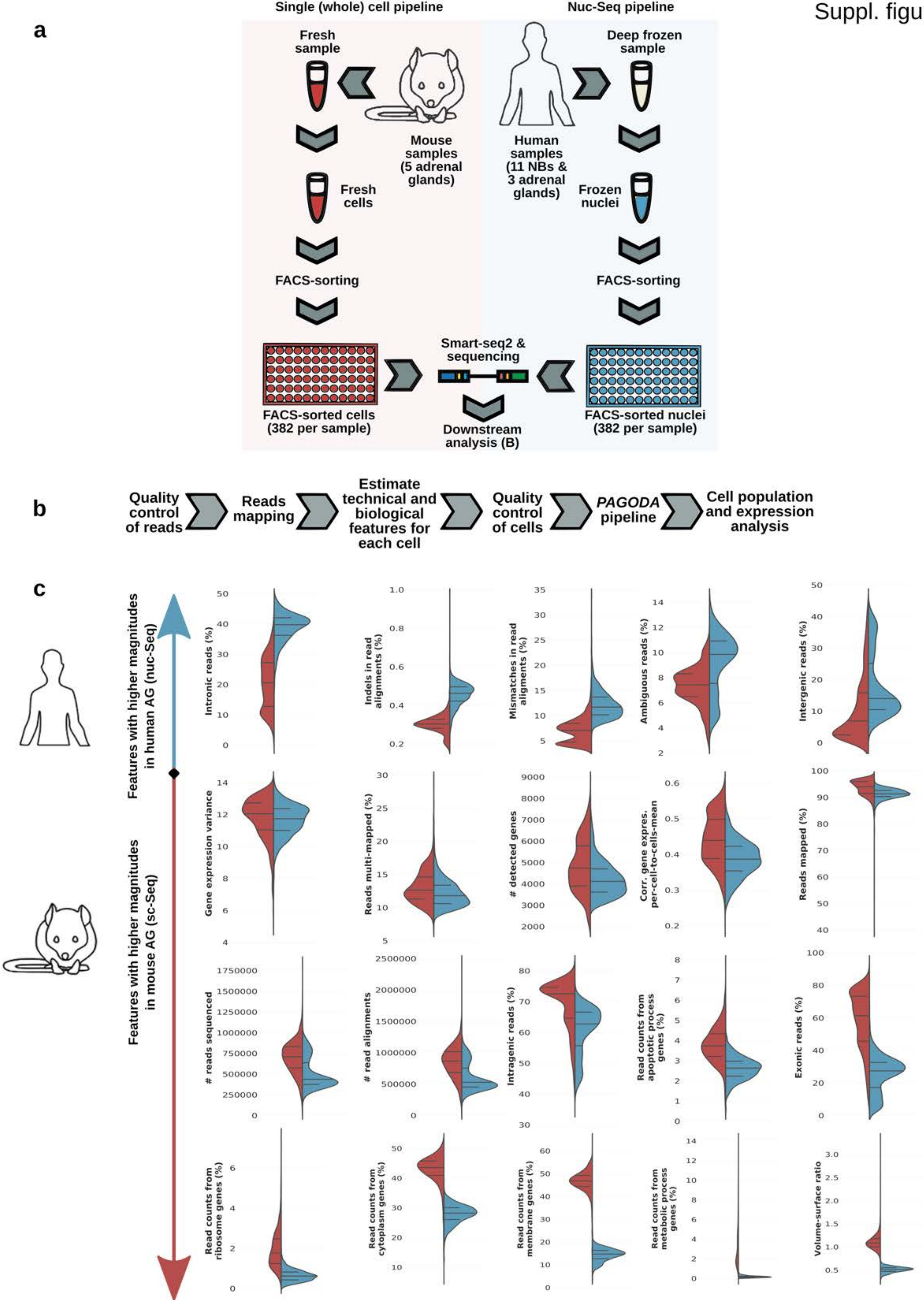

Suppl. figure 2

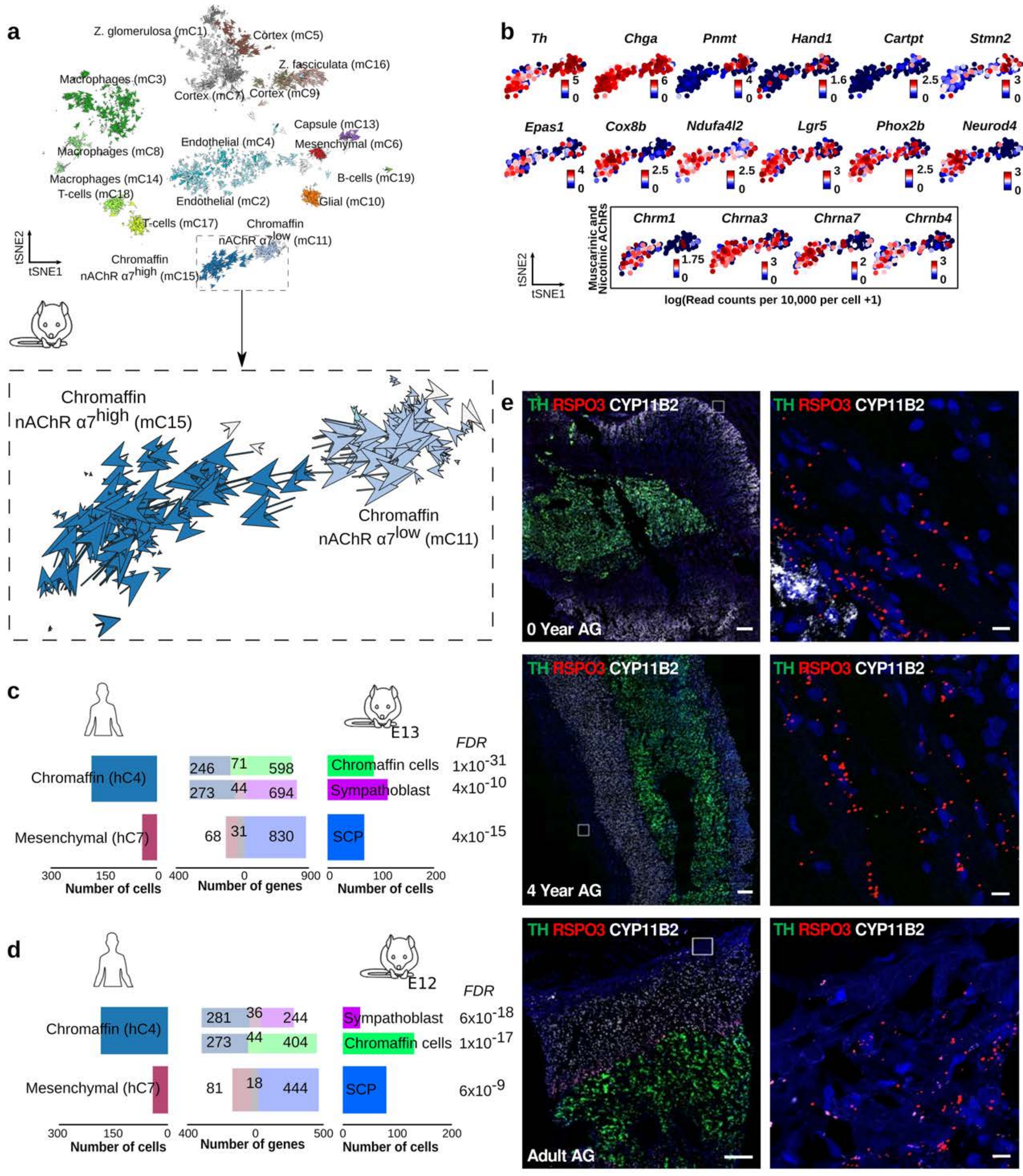

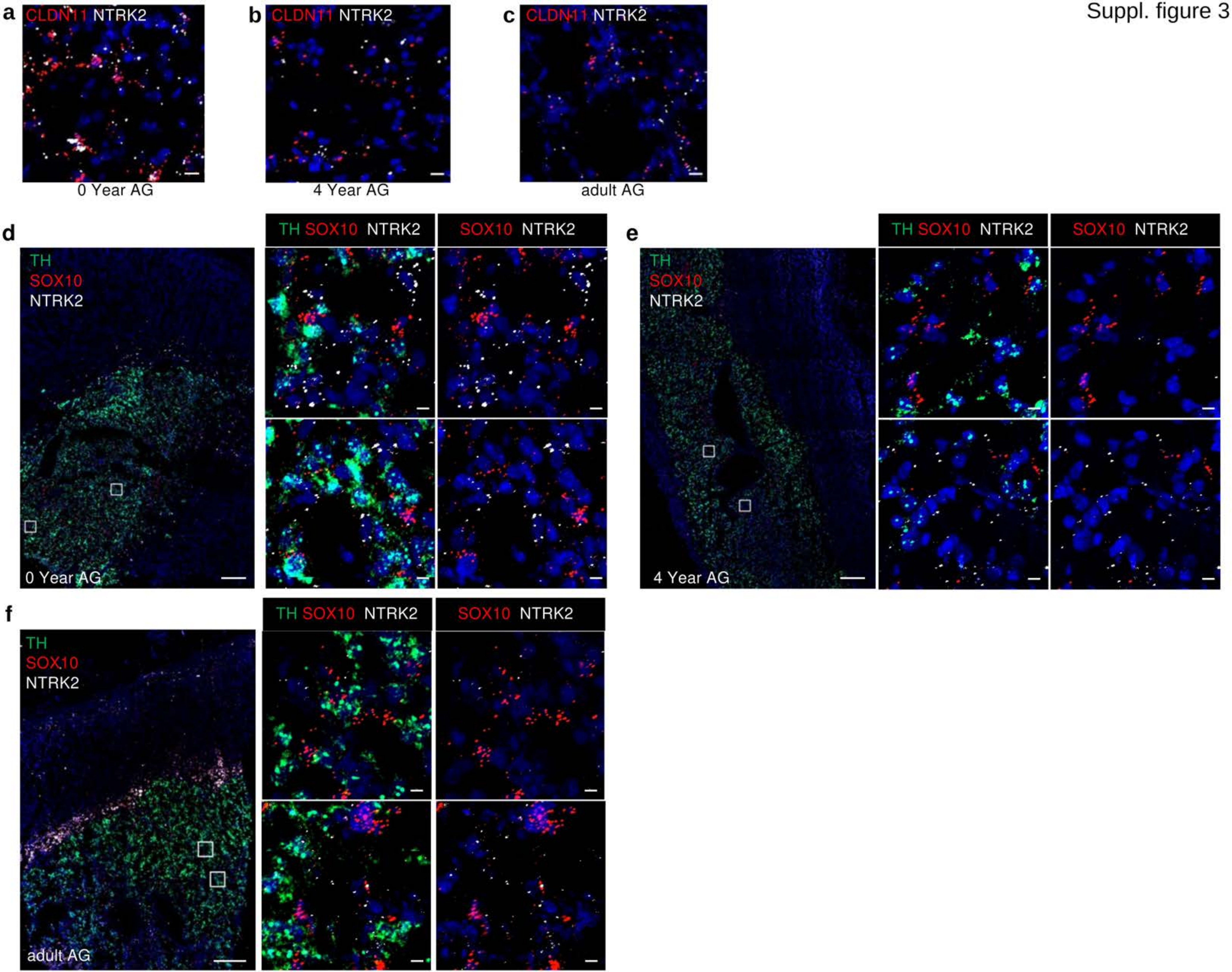

Suppl. figure 4

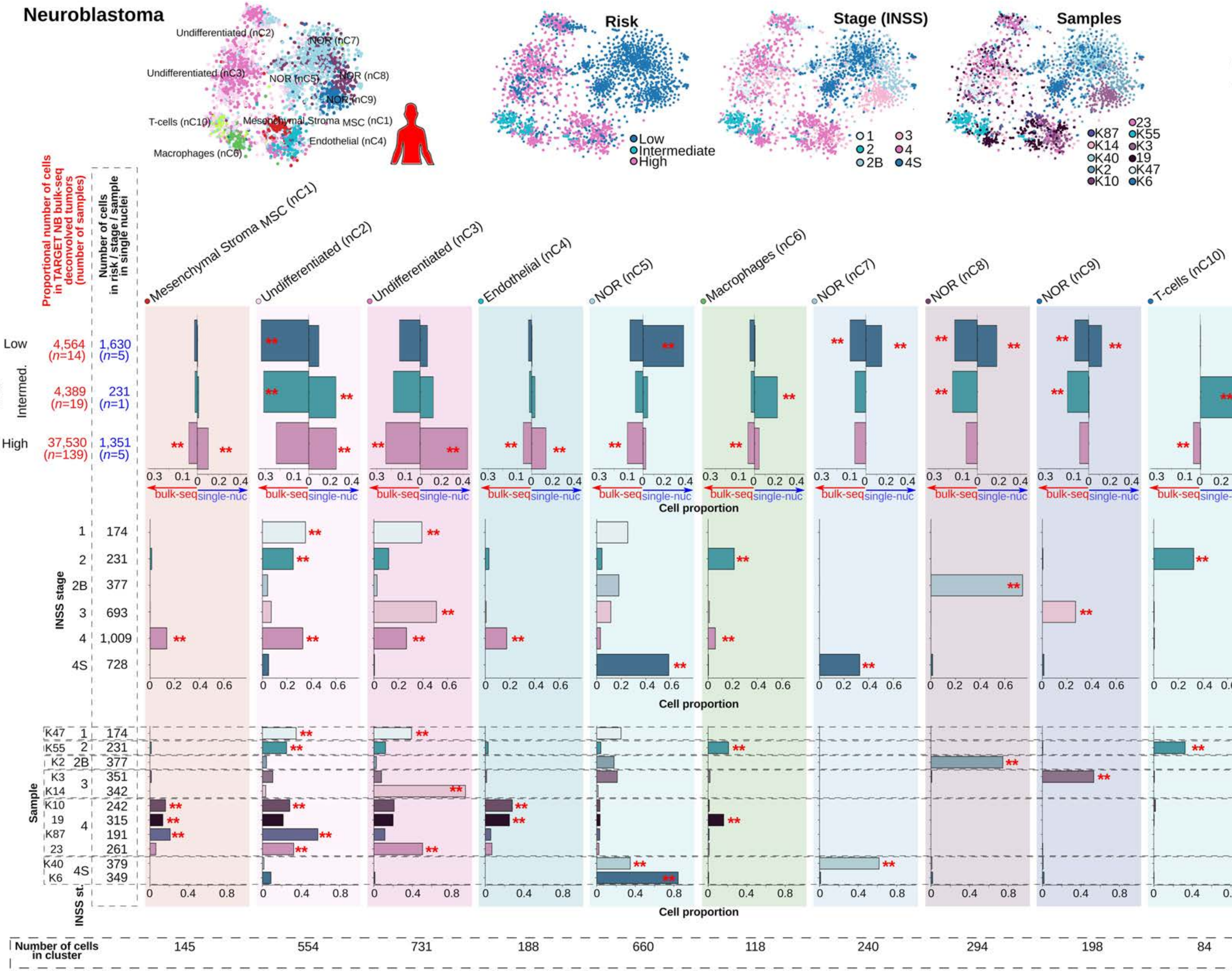

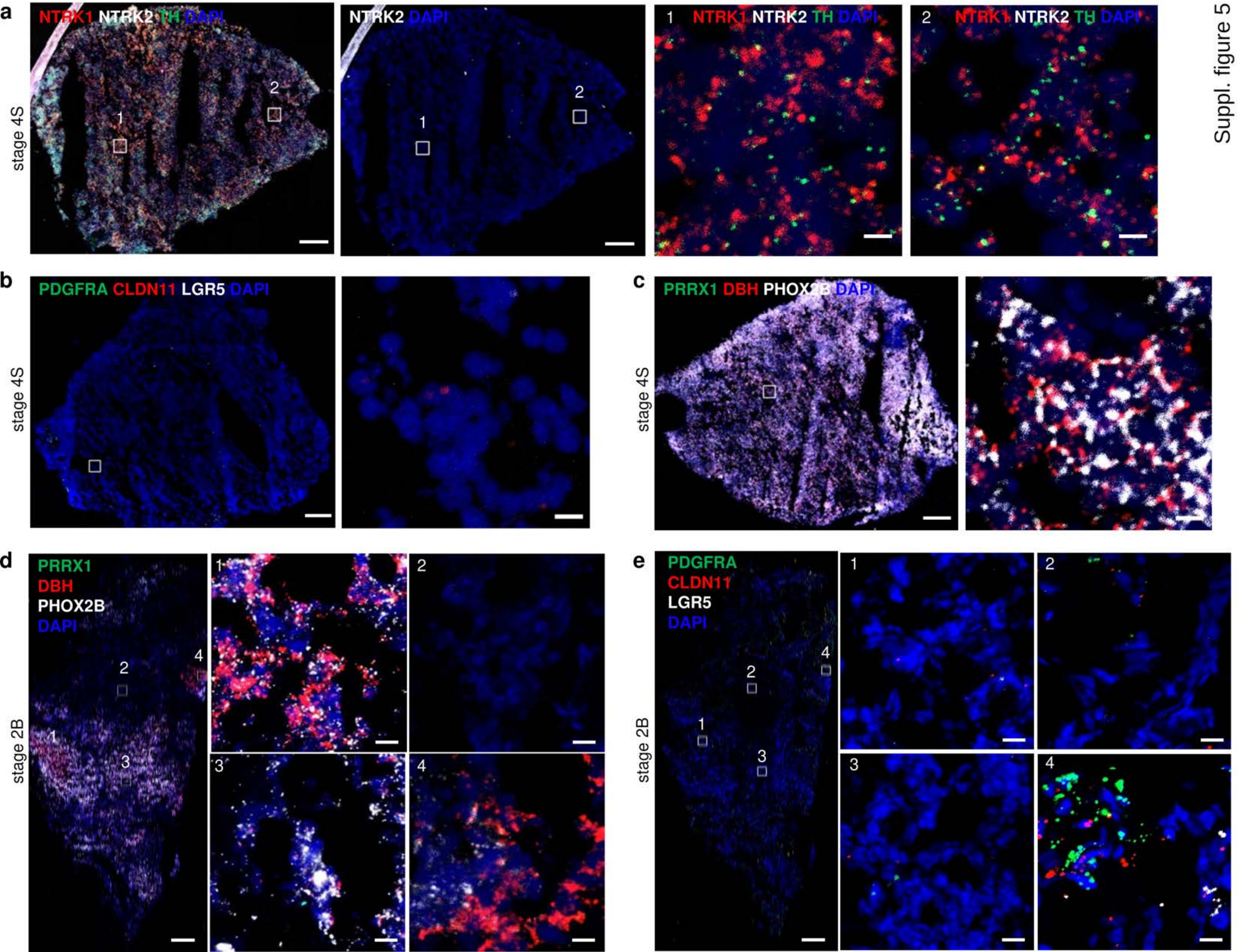

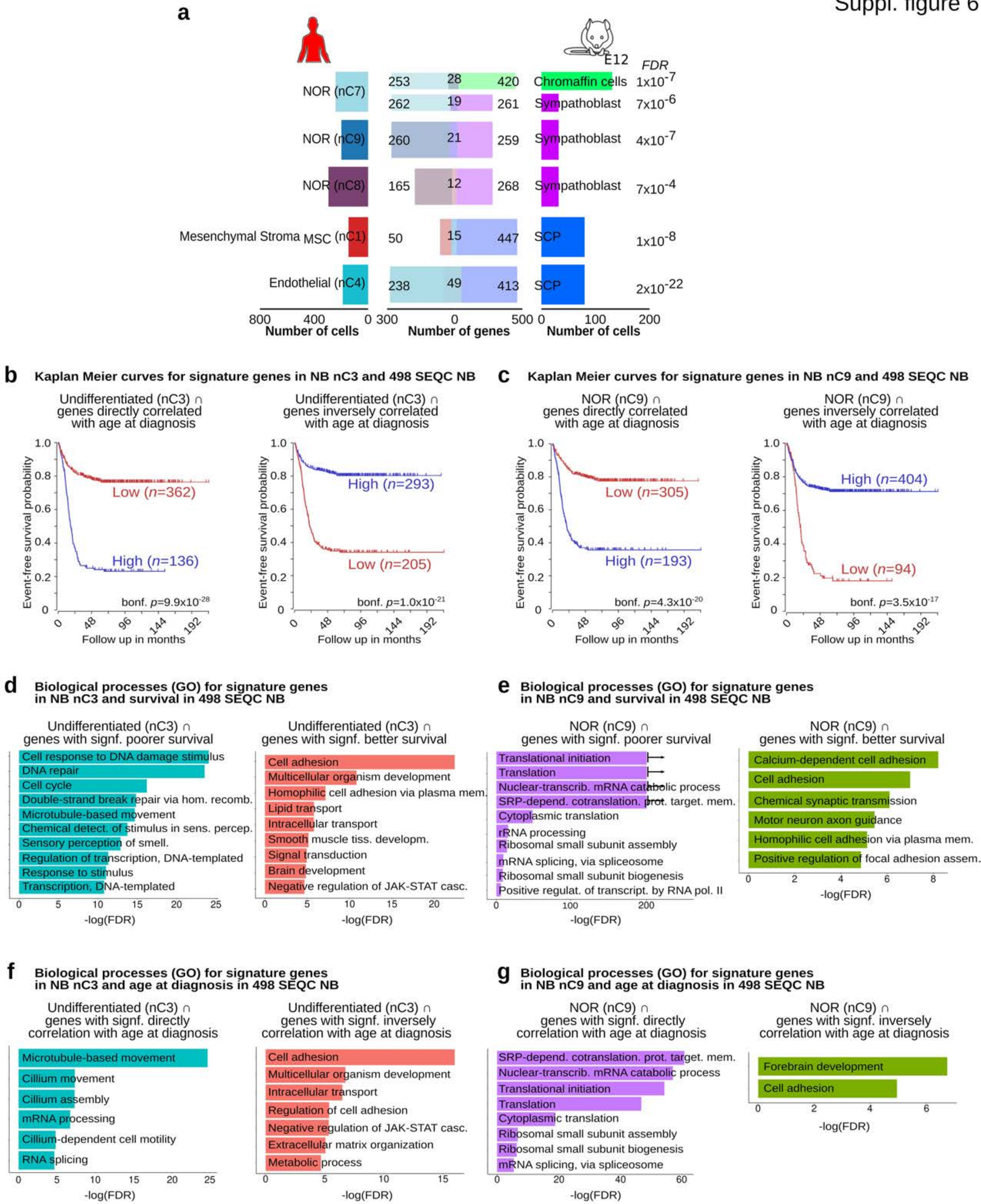
