## Supplementary Table 1 for "Single-nuclei transcriptomes from human adrenal gland reveals distinct cellular identities of low and high-risk neuroblastoma tumors"

| Sample | Case study | Total number of cells/nuclei sequenced | Total number of reads sequenced | Number of high-quality cells/nuclei | Total number of high-quality reads | Number of mapped high-quality reads | Age (weeks/months/years) |
|---|---|---|---|---|---|---|---|
| 16-D | Human adrenal gland | 384 | 282,900,000 | 372 | 275,963,970 | 255,587,957 | Adult |
| 6657 | Human adrenal gland | 768 | 301,300,000 | 622 | 255,465,926 | 232,007,098 | 50 years |
| 6435 | Human adrenal gland | 384 | 160,600,000 | 328 | 139,869,057 | 126,741,253 | 44 years |
| 6801 | Mouse adrenal gland | 384 | 276,000,000 | 370 | 271,605,447 | 261,029,317 | 44 weeks |
| 6802 | Mouse adrenal gland | 384 | 270,900,000 | 353 | 264,299,115 | 249,793,782 | 44 weeks |
| 3431 | Mouse adrenal gland | 384 | 255,500,000 | 363 | 248,666,531 | 228,411,355 | 42 weeks |
| I1 | Mouse adrenal gland | 384 | 239,000,000 | 308 | 211,053,288 | 196,335,893 | ~30 weeks |
| 3432 | Mouse adrenal gland | 384 | 243,200,000 | 369 | 236,618,245 | 218,409,975 | 42 weeks |
| 19 | Neuroblastoma | 384 | 223,000,000 | 315 | 195,130,950 | 169,973,268 | 30 months |
| 23 | Neuroblastoma | 384 | 270,300,000 | 261 | 189,958,512 | 170,423,470 | 27 months |
| K10 | Neuroblastoma | 384 | 238,200,000 | 242 | 166,689,065 | 149,398,264 | 14 months |
| K14 | Neuroblastoma | 384 | 281,000,000 | 342 | 252,164,823 | 230,215,012 | 79 months |
| K2 | Neuroblastoma | 384 | 255,000,000 | 377 | 253,718,019 | 233,377,167 | 31 months |
| K3 | Neuroblastoma | 384 | 258,100,000 | 351 | 239,337,436 | 215,787,879 | 6 months |
| K40 | Neuroblastoma | 384 | 254,700,000 | 379 | 253,338,912 | 231,285,195 | 0 months |
| K47 | Neuroblastoma | 384 | 271,600,000 | 174 | 134,358,061 | 120,026,522 | 13 months |
| K55 | Neuroblastoma | 384 | 280,800,000 | 231 | 197,447,878 | 176,063,121 | 62 months |
| K6 | Neuroblastoma | 384 | 272,200,000 | 349 | 255,883,674 | 224,429,878 | 0 months |
| K87 | Neuroblastoma | 384 | 251,800,000 | 191 | 141,451,299 | 126,303,929 | 21 months |

**Neuroblastoma sample details**

| Sample | Genotype | Gender | Risk | INSS stage | INRGSS Ada | MYCN ampl | 1p deletion | 1p ab. Ext. (mb fr 1 pter) | Ploidy | CGH class | Outcome | Survival (Months) | Pretreated | 17q-gain |
|---|---|---|---|---|---|---|---|---|---|---|---|---|---|---|
| 19 | nd | M | High | 4 | M | no | ? | ? | - | NMA | DOD | 13 | ? | ? |
| 23 | 1p36+/- | M | High | 4 | M | yes | yes | 0-32.9 | - | NMA;11q- | AWD | 39+' | ? | 39.6-qter |
| K10 | 1p36+/- | M | High | 4 | M | yes | yes | 0-54.9 | 2n | NMA | DOD | 13 | yes | 29.4-qter |
| K14 | 1p36+/+ | M | High | 3 | L | yes | gain | 0-227.3 | 3n | NMA | NED | 179+' | yes | 34.8-qter |
| K2 | 1p36+/+ | M | Low | 2B | L | no | no | - | - | NO | NED | 271+' | no | ? |
| K3 | nd | M | Low | 3 | L | no | nd | - | - | NO | NED | 288+' | no | whole 17 gain |
| K40 | 1p36+/+ | M | Low | 4S | MS | no | no | - | - | NO | NED | 269+' | ? | ? |
| K47 | 1p36+/+ | M | Low | 1 | L | no | no | - | - | OS | NED | 120+' | no | whole 17 gain |
| K55 | 1p36+/- | F | Intermediate | 2 | L | no | yes | 0-28.4 | - | OS | NED | 77+' | yes | no |
| K6 | 1p36+/+ | M | Low | 4S | MS | no | no | - | - | NO | NED | 248+' | ? | ? |
| K87 | 1p36+/+ | F | High | 4 | M | no | no | - | - | 11q- | AWD | 60+' | ? | 39.9-qter |

**CGH class:**

OS=Other segmental

17q=17q+ without NMA/11q-'

NMA=MYCN-amplification

**INRGSS stage:**

L=Localized, L1/L2 (INSS 1,2,3)
