## Supplementary Table 2 for "Single-nuclei transcriptomes from human adrenal gland reveals distinct cellular identities of low and high-risk neuroblastoma tumors"

**Markers in human post-natal AG**

| Human adrenal gland marker | Cell type | References | Evidence for high expression in human AG cluster | Evidence for higher expression in human AG cluster | Each cluster in human AG: FDR | Cluster pairwise comparison in human AG: FDR |
|---|---|---|---|---|---|---|
| CHGA | Adrenergic (Adrenal medulla) | [54,63] | hC4 | hC4 | FDR(hC4>hCo): <1E-50 | |
| CHGB | Adrenergic (Adrenal medulla) | [54,63] | hC1, hC3, hC4 | hC4 | FDR(hC1>hCo): 1.7E-4, FDR(hC3>hCo): 6.8E-4, FDR(hC4>hCo): <1E-50 | FDR(hC4>hC1): 1.7E-29, FDR(hC4>hC3): 5.9E-37 |
| DBH | Adrenergic (Adrenal medulla) | [54,64] | hC4 | hC4 | FDR(hC4>hCo): <1E-50 | |
| PNMT | Adrenergic (Adrenal medulla) | [54,62] | hC4 | hC4 | FDR(hC4>hCo): <1E-50 | |
| TH | Adrenergic (Adrenal medulla) | [54,62] | hC4 | hC4 | FDR(hC4>hCo): <1E-50 | |
| CHRNA3 | Chromaffin | [54,62] | hC4 | hC4 | FDR(hC4>hCo): 1.0E-36 | |
| CHRNA7 | Chromaffin NE-secreting | [65,66,67,68] | hC1, hC4 | hC4 | FDR(hC1>hCo): 2.2E-9, FDR(hC4>hCo): <1E-50 | FDR(hC4>hC1): 7.3E-9 |
| DLK1 | Differentiation | [2] | hC3, hC4 | hC3, hC4 | FDR(hC3>hCo): 6.9E-8, FDR(hC4>hCo): <1E-50 | |
| RET | Differentiation | [106] | hC4 | hC4 | FDR(hC4>hCo): <1E-50 | |
| TP63 | Differentiation | [105] | hC1 | hC1 | FDR(hC1>hCo): 2.5E-9 | |
| PECAM1 | Endothelial | [78] | hC1, hC6 | hC6 | FDR(hC1>hCo): 1.4E-8, FDR(hC6>hCo): 3.8E-11 | FDR(hC6>hC1): 7.2E-8 |
| SOX18 | Endothelial | [80] | hC6 | hC6 | FDR(hC6>hCo): 1.5E-5 | |
| TEK | Endothelial | [78] | hC1, hC6 | hC6 | FDR(hC1>hCo): 3.1E-4, FDR(hC6>hCo): 5.3E-6 | FDR(hC6>hC1): 2.1E-4 |
| TIE1 | Endothelial | [79] | hC1, hC6 | hC6 | FDR(hC1>hCo): 4.0E-3, FDR(hC6>hCo): 1.1E-5 | FDR(hC6>hC1): 1.9E-4 |
| NTRK1 | Favourable outcome | [2] | hC4 | hC4 | FDR(hC4>hCo): 1.1E-8 | |
| EPAS1 | Hypoxia | [103] | hC6, hC7 | hC6, hC7 | FDR(hC6>hCo): 1.1E-11, FDR(hC7>hCo): 1.0E-5 | |
| CD14 | Macrophages | [85] | hC2 | hC2 | FDR(hC2>hCo): 2.9E-18 | |
| CD163 | Macrophages | [82] | hC2 | hC2 | FDR(hC2>hCo): <1E-50 | |
| CD300A | Macrophages | [85] | hC2 | hC2 | FDR(hC2>hCo): 6.3E-4 | |
| CD84 | Macrophages | [84] | hC1, hC2 | hC2 | FDR(hC1>hCo): 6.0E-5, FDR(hC2>hCo): 4.9E-10 | FDR(hC2>hC1): 3.0E-4 |
| CD86 | Macrophages | [83] | hC2 | hC2 | FDR(hC2>hCo): 7.6E-8 | |
| CSF1R | Macrophages | [81] | hC2 | hC2 | FDR(hC2>hCo): 7.5E-32 | |
| COL12A1 | Mesenchymal | [77] | hC1, hC7 | hC7 | FDR(hC1>hCo): 6.7E-13, FDR(hC7>hCo): 1.2E-9 | FDR(hC7>hC1): 6.8E-5 |
| COL1A2 | Mesenchymal | [75,76] | hC1, hC7 | hC7 | FDR(hC1>hCo): 7.9E-12, FDR(hC7>hCo): 4.3E-10 | FDR(hC7>hC1): 1.3E-5 |
| COL6A3 | Mesenchymal | [76] | hC1, hC7 | hC7 | FDR(hC1>hCo): 2.7E-10, FDR(hC7>hCo): 3.6E-6 | FDR(hC7>hC1): 9.0E-3 |
| CXCL12 | Mesenchymal | [74] | hC7 | hC7 | FDR(hC7>hCo): 6.9E-9 | |
| DCN | Mesenchymal | [73] | hC5, hC7, hC9 | hC7 | FDR(hC5>hCo): 1.4E-6, FDR(hC7>hCo): 4.6E-16, FDR(hC9>hCo): 1.8E-30 | FDR(hC7>hC5): 3.4E-8, FDR(hC7>hC9): 7.3E-5 |
| PDGFRA | Mesenchymal | [70,71] | hC1, hC7 | hC7 | FDR(hC1>hCo): 2.8E-11, FDR(hC7>hCo): 2.9E-16 | FDR(hC7>hC1): 1.8E-11 |
| PDGFRB | Mesenchymal | [72] | hC1, hC7 | hC7 | FDR(hC1>hCo): 6.4E-4, FDR(hC7>hCo): 1.5E-8 | FDR(hC7>hC1): 7.8E-7 |
| PRRX1 | Mesenchymal | [16] | hC1, hC5 | hC1, hC5 | FDR(hC1>hCo): 2.1E-4, FDR(hC5>hCo): 9.5E-3 | |
| CLDN11 | Migratory | [100] | hC1 | hC1 | FDR(hC1>hCo): 1.7E-36 | |
| DOCK7 | Migratory | [101] | hC1, hC2 | hC1 | FDR(hC1>hCo): 5.9E-20, FDR(hC2>hCo): 1.3E-4 | FDR(hC1>hC2): 6.4E-3 |

| | | | | | | |
|---|---|---|---|---|---|---|
| LAMA3 | Migratory | [102] | hC1 | hC1 | FDR(hC1>hCo): 1.1E-18 | |
| SEMA3E | Migratory | [99] | hC1 | hC1 | FDR(hC1>hCo): 9.8E-6 | |
| NTRK3 | Neuroblastoma | [2] | hC1, hC4 | hC1, hC4 | FDR(hC1>hCo): 6.7E-18, FDR(hC4>hCo): 1.0E-9 | |
| ASXL3 | Progenitors | [95] | hC1 | hC1 | FDR(hC1>hCo): 1.1E-7 | |
| BCL11A | Progenitors | [94] | hC1 | hC1 | FDR(hC1>hCo): 5.1E-10 | |
| ERBB3 | Progenitors | [91] | hC1 | hC1 | FDR(hC1>hCo): 4.4E-10 | |
| LGR5 | Progenitors | [96] | hC5 | hC5 | FDR(hC5>hCo): 1.3E-3 | |
| NF1 | Progenitors | [98] | hC1 | hC1 | FDR(hC1>hCo): 6.6E-12 | |
| POU6F2 | Progenitors | [93] | hC1 | hC1 | FDR(hC1>hCo): 1.1E-8 | |
| RTTN | Progenitors | [92] | hC1 | hC1 | FDR(hC1>hCo): 7.2E-11 | |
| SOX5 | Progenitors | [97] | hC1, hC4 | hC1, hC4 | FDR(hC1>hCo): 2.4E-7, FDR(hC4>hCo): 4.2E-7 | |
| SOX6 | Progenitors | [90] | hC1 | hC1 | FDR(hC1>hCo): 2.8E-10 | |
| CARTPT | Sympathoadrenal lineage | [6] | hC4 | hC4 | FDR(hC4>hCo): 2.6E-22 | |
| GATA3 | Sympathoadrenal lineage | [6] | hC4 | hC4 | FDR(hC4>hCo): 1.5E-17 | |
| HAND1 | Sympathoadrenal lineage | [6] | hC4 | hC4 | FDR(hC4>hCo): 1.5E-6 | |
| PHOX2A | Sympathoadrenal lineage | [69] | hC4 | hC4 | FDR(hC4>hCo): 1.9E-13 | |
| PHOX2B | Sympathoadrenal lineage | [69] | hC4 | hC4 | FDR(hC4>hCo): 6.1E-9 | |
| STMN2 | Sympathoadrenal lineage | [6] | hC4 | hC4 | FDR(hC4>hCo): 5.4E-18 | |
| PRPH | Sympathoblast | [6] | hC4 | hC4 | FDR(hC4>hCo): 7.4E-11 | |
| CD3E | T-cells | [86] | hC10 | hC10 | FDR(hC10>hCo): 3.7E-4 | |
| HCST | T-cells | [88] | hC10 | hC10 | FDR(hC10>hCo): 1.9E-4 | |
| TRAC | T-cells | [89] | hC10 | hC10 | FDR(hC10>hCo): 2.5E-3 | |
| TRBC1 | T-cells | [89] | hC10 | hC10 | FDR(hC10>hCo): 1.4E-5 | |
| TRBC2 | T-cells | [89] | hC10 | hC10 | FDR(hC10>hCo): 5.6E-5 | |
| BRCA1 | Unfavourable outcome | [104] | hC1 | hC1 | FDR(hC1>hCo): 6.5E-12 | |
| NTRK2 | Unfavourable outcome | [2] | hC1 | hC1 | FDR(hC1>hCo): 6.7E-9 | |
| CYP11A1 | Zona fasciculata (Cortex) | [49] | hC3, hC8, hC9 | hC3 | FDR(hC3>hCo): <1E-50, FDR(hC8>hCo): 1.3E-29, FDR(hC9>hCo): 7.0E-24 | FDR(hC3>hC8): 9.9E-4, FDR(hC3>hC9): 4.9E-4 |
| CYP11B1 | Zona fasciculata (Cortex) | [50,51,52] | hC3, hC5, hC8, hC9 | hC3 | FDR(hC3>hCo): <1E-50, FDR(hC5>hCo): 2.5E-20, FDR(hC8>hCo): <1E-50, FDR(hC9>hCo): <1E-50 | FDR(hC3>hC5): 6.2E-23, FDR(hC3>hC8): 1.3E-11, FDR(hC3>hC9): 2.3E-26, FDR(hC8>hC5): 2.0E-12, FDR(hC8>hC9): 1.3E-5, FDR(hC9>hC5): 6.0E-6 |
| STAR | Zona fasciculata (Cortex) & Zona reticularis (Cortex) | [53,54] | hC3, hC5, hC8, hC9 | hC3 | FDR(hC3>hCo): <1E-50, FDR(hC5>hCo): 3.3E-27, FDR(hC8>hCo): 8.3E-24, FDR(hC9>hCo): 6.7E-33 | FDR(hC3>hC5): 4.9E-5, FDR(hC3>hC8): 1.0E-12, FDR(hC3>hC9): 1.4E-6 |
| AGTR1 | Zona glomerulosa (Cortex) | [57,58,59] | hC5, hC7 | hC5, hC7 | FDR(hC5>hCo): 4.5E-3, FDR(hC7>hCo): 6.1E-3 | |
| CYP11B2 | Zona glomerulosa (Cortex) | [54,50,51] | hC3, hC5, hC8, hC9 | hC3, hC5, hC8 | FDR(hC3>hCo): <1E-50, FDR(hC5>hCo): <1E-50, FDR(hC8>hCo): <1E-50, FDR(hC9>hCo): <1E-50 | FDR(hC3>hC9): 3.3E-24, FDR(hC5>hC9): 1.7E-4, FDR(hC8>hC9): 1.4E-24 |
| DAB2 | Zona glomerulosa (Cortex) | [55,56] | hC2, hC5, hC8, hC9 | hC2, hC5, hC9 | FDR(hC2>hCo): 1.7E-12, FDR(hC5>hCo): 2.1E-9, FDR(hC8>hCo): 1.5E-10, FDR(hC9>hCo): 1.0E-27 | FDR(hC2>hC8): 7.6E-3 |
| AKR1B1 | Zona reticularis (Cortex) | [60] | hC3, hC5, hC8, hC9 | hC3, hC8, hC9 | FDR(hC3>hCo): 2.4E-18, FDR(hC5>hCo): 5.8E-5, FDR(hC8>hCo): <1E-50, FDR(hC9>hCo): <1E-50 | FDR(hC3>hC5): 6.2E-3, FDR(hC8>hC5): 1.9E-6, FDR(hC9>hC5): 7.2E-10 |
| SULT2A1 | Zona reticularis (Cortex) | [61] | hC8 | hC8 | FDR(hC8>hCo): 6.4E-24 | |
| TSPAN12 | Zona reticularis (Cortex) | [49] | hC8 | hC8 | FDR(hC8>hCo): 1.3E-6 | |

| | | | | | |
|---|---|---|---|---|---|
| CYP17A1 | Zona reticularis (Cortex) & Zona fasiculata (Cortex) | [50,49,54] | hC3, hC8, hC9 | hC3, hC8, hC9 | FDR(hC3>hCo): 2.7E-16, FDR(hC8>hCo): <1E-50, FDR(hC9>hCo): 5.0E-35 |

**Markers in mouse post-natal AG**

| Mouse adrenal gland marker | Cell type | References | Evidence for high expression in mouse AG cluster | Evidence for higher expression in mouse AG cluster | Each cluster in mouse AG: FDR | Cluster pairwise comparison in mosue AG: FDR |
|---|---|---|---|---|---|---|
| Chga | Adrenergic (Adrenal medulla) | [6,63] | mC2, mC11, mC15, mC16, mC17 | mC15 | FDR(mC2>mCo): 3.7E-3, FDR(mC11>mCo): <1E-50, FDR(mC15>mCo): <1E-50, FDR(mC16>mCo): 1.0E-9, FDR(mC17>mCo): 1.4E-3 | FDR(mC11>mC2): <1E-50, FDR(mC11>mC16): 9.3E-29, FDR(mC11>mC17): 1.4E-25, FDR(mC15>mC2): <1E-50, FDR(mC15>mC11): 4.1E-11, FDR(mC15>mC16): 1.4E-35, FDR(mC15>mC17): 3.3E-31, FDR(mC16>mC2): 1.7E-4 |
| Chgb | Adrenergic (Adrenal medulla) | [6,63] | mC2, mC11, mC15, mC16, mC17, mC18 | mC15 | FDR(mC2>mCo): 2.9E-5, FDR(mC11>mCo): <1E-50, FDR(mC15>mCo): <1E-50, FDR(mC16>mCo): 4.2E-12, FDR(mC17>mCo): 4.4E-8, FDR(mC18>mCo): 5.4E-4 | FDR(mC11>mC2): <1E-50, FDR(mC11>mC16): 3.6E-25, FDR(mC11>mC17): 3.3E-26, FDR(mC11>mC18): 6.9E-23, FDR(mC15>mC2): <1E-50, FDR(mC15>mC11): 2.1E-13, FDR(mC15>mC16): 4.8E-32, FDR(mC15>mC17): 9.0E-33, FDR(mC15>mC18): 7.9E-28, FDR(mC16>mC2): 2.2E-5, FDR(mC17>mC2): 5.4E-3 |
| Dbh | Adrenergic (Adrenal medulla) | [6,64] | mC11, mC15, mC16 | mC15 | FDR(mC11>mCo): <1E-50, FDR(mC15>mCo): <1E-50, FDR(mC16>mCo): 3.1E-3 | FDR(mC11>mC16): 3.8E-29, FDR(mC15>mC11): 1.2E-10, FDR(mC15>mC16): 3.1E-36 |
| Pnmt | Adrenergic (Adrenal medulla) | [6,62] | mC11, mC15, mC16 | mC11, mC16 | FDR(mC11>mCo): 2.3E-6, FDR(mC15>mCo): 1.4E-8, FDR(mC16>mCo): 8.0E-9 | FDR(mC11>mC15): 9.9E-3 |
| Th | Adrenergic (Adrenal medulla) | [6,62] | mC11, mC15, mC16, mC17 | mC11, mC15 | FDR(mC11>mCo): 8.3E-40, FDR(mC15>mCo): <1E-50, FDR(mC16>mCo): 9.6E-5, FDR(mC17>mCo): 9.5E-3 | FDR(mC11>mC16): 5.4E-23, FDR(mC11>mC17): 1.2E-22, FDR(mC15>mC16): 9.5E-22, FDR(mC15>mC17): 1.7E-21 |
| Cd19 | B-cells | [115] | mC1, mC5, mC15, mC16, mC19 | mC19 | FDR(mC1>mCo): 1.6E-6, FDR(mC5>mCo): 6.8E-22, FDR(mC15>mCo): 3.6E-7, FDR(mC16>mCo): 1.0E-8, FDR(mC19>mCo): 1.8E-5 | FDR(mC5>mC1): 1.0E-13, FDR(mC5>mC15): 5.7E-4, FDR(mC16>mC1): 7.1E-6, FDR(mC16>mC5): 3.2E-3, FDR(mC16>mC15): 6.8E-5, FDR(mC19>mC1): 8.1E-6, FDR(mC19>mC5): 1.0E-5, FDR(mC19>mC15): 8.6E-6, FDR(mC19>mC16): 1.5E-5 |
| Ighm | B-cells | [116] | mC3, mC5, mC8, mC18, mC19 | mC19 | FDR(mC3>mCo): 7.2E-23, FDR(mC5>mCo): 1.9E-11, FDR(mC8>mCo): 4.0E-5, FDR(mC18>mCo): 3.8E-4, FDR(mC19>mCo): 3.4E-31 | FDR(mC19>mC3): <1E-50, FDR(mC19>mC5): 4.2E-27, FDR(mC19>mC8): 3.3E-11, FDR(mC19>mC18): 5.3E-20 |
| Ms4a1 | B-cells | [114,115] | mC1, mC5, mC15, mC16, mC19 | mC19 | FDR(mC1>mCo): 3.7E-4, FDR(mC5>mCo): 5.1E-17, FDR(mC15>mCo): 1.5E-3, FDR(mC16>mCo): 8.6E-6, FDR(mC19>mCo): 4.5E-6 | FDR(mC5>mC1): 1.0E-13, FDR(mC16>mC1): 2.5E-4, FDR(mC19>mC1): 1.9E-6, FDR(mC19>mC5): 2.5E-6, FDR(mC19>mC15): 2.4E-6, FDR(mC19>mC16): 3.8E-6 |
| Htr3a | Bridge | [6] | mC1, mC7, mC11 | mC1 | FDR(mC1>mCo): 6.2E-25, FDR(mC7>mCo): 4.9E-3, FDR(mC11>mCo): 1.8E-3 | FDR(mC1>mC7): 8.1E-5, FDR(mC1>mC11): 4.9E-6 |
| Gli1 | Capsule | [109] | mC5, mC13 | mC13 | FDR(mC5>mCo): 3.3E-4, FDR(mC13>mCo): 2.1E-3 | FDR(mC13>mC5): 1.1E-3 |
| Rspo3 | Capsule | [109] | mC1, mC13 | mC13 | FDR(mC1>mCo): 4.4E-5, FDR(mC13>mCo): 1.2E-12 | FDR(mC13>mC1): 1.3E-13 |
| Mki67 | Cell cycle | | mC8 | mC8 | FDR(mC8>mCo): 7.5E-4 | |

| | | | | | | |
|---|---|---|---|---|---|---|
| Chrm1 | Chromaffin | [107] | mC11, mC15, mC17, mC19 | mC15 | FDR(mC11>mCo): 1.4E-4, FDR(mC15>mCo): 1.7E-16, FDR(mC17>mCo): 7.1E-4, FDR(mC19>mCo): 8.7E-3 | FDR(mC11>mC17): 1.1E-3, FDR(mC15>mC11): 7.3E-6, FDR(mC15>mC17): 3.4E-14, FDR(mC15>mC19): 8.7E-10 |
| Chrna3 | Chromaffin | [65,67] | mC11, mC15 | mC15 | FDR(mC11>mCo): 3.6E-8, FDR(mC15>mCo): <1E-50 | FDR(mC15>mC11): 2.4E-9 |
| Chrnb4 | Chromaffin | [67] | mC11, mC15 | mC15 | FDR(mC11>mCo): 1.5E-5, FDR(mC15>mCo): 5.7E-29 | FDR(mC15>mC11): 1.6E-4 |
| Npy | Chromaffin | [63] | mC11, mC15, mC16 | mC11, mC15 | FDR(mC11>mCo): <1E-50, FDR(mC15>mCo): <1E-50, FDR(mC16>mCo): 1.3E-15 | FDR(mC11>mC16): 5.5E-21, FDR(mC15>mC16): 4.0E-20 |
| Chrna7 | Chromaffin NE-secreting | [65,66,67,68] | mC11, mC15, mC19 | mC15 | FDR(mC11>mCo): 1.2E-5, FDR(mC15>mCo): <1E-50, FDR(mC19>mCo): 3.2E-5 | FDR(mC11>mC19): 2.5E-4, FDR(mC15>mC11): 1.3E-13, FDR(mC15>mC19): 1.6E-42 |
| Ret | Differentiation | [106] | mC15, mC19 | mC15 | FDR(mC15>mCo): 6.4E-4, FDR(mC19>mCo): 2.5E-4 | FDR(mC15>mC19): 3.9E-3 |
| Pecam1 | Endothelial | [78] | mC2, mC4 | mC2, mC4 | FDR(mC2>mCo): <1E-50, FDR(mC4>mCo): <1E-50 | |
| Sox18 | Endothelial | [80] | mC2, mC4, mC5, mC16 | mC2, mC4 | FDR(mC2>mCo): <1E-50, FDR(mC4>mCo): 2.8E-41, FDR(mC5>mCo): 9.1E-5, FDR(mC16>mCo): 2.6E-14 | FDR(mC2>mC5): <1E-50, FDR(mC2>mC16): 4.1E-41, FDR(mC4>mC5): 4.8E-35, FDR(mC4>mC16): 2.2E-30, FDR(mC16>mC5): 1.1E-5 |
| Sox4 | Endothelial | [111] | mC4 | mC4 | FDR(mC4>mCo): 4.2E-9 | |
| Tek | Endothelial | [78] | mC2, mC4, mC16 | mC2, mC4 | FDR(mC2>mCo): <1E-50, FDR(mC4>mCo): <1E-50, FDR(mC16>mCo): 7.7E-4 | FDR(mC2>mC16): <1E-50, FDR(mC4>mC16): <1E-50 |
| Tie1 | Endothelial | [79] | mC2, mC4, mC16 | mC2, mC4 | FDR(mC2>mCo): <1E-50, FDR(mC4>mCo): <1E-50, FDR(mC16>mCo): 5.0E-6 | FDR(mC2>mC16): <1E-50, FDR(mC4>mC16): <1E-50 |
| Sox17 | Endothelial (haemogenic) | [110] | mC2, mC4, mC5, mC16 | mC4 | FDR(mC2>mCo): 7.6E-8, FDR(mC4>mCo): 5.7E-14, FDR(mC5>mCo): 1.2E-10, FDR(mC16>mCo): 1.6E-14 | FDR(mC4>mC2): 3.2E-6, FDR(mC4>mC5): 1.7E-10, FDR(mC4>mC16): 8.5E-9, FDR(mC16>mC5): 2.2E-3 |
| S100b | Glial | [117,6] | mC10 | mC10 | FDR(mC10>mCo): 1.2E-27 | |
| Sox10 | Glial | [117,6] | mC10 | mC10 | FDR(mC10>mCo): <1E-50 | |
| Cox8b | Hypoxia | [103] | mC11, mC15 | mC15 | FDR(mC11>mCo): 3.0E-5, FDR(mC15>mCo): <1E-50 | FDR(mC15>mC11): 2.1E-15 |
| Epas1 | Hypoxia | [103] | mC2, mC4, mC5, mC15 | mC2, mC4 | FDR(mC2>mCo): <1E-50, FDR(mC4>mCo): <1E-50, FDR(mC5>mCo): 1.7E-3, FDR(mC15>mCo): 7.7E-11 | FDR(mC2>mC5): 3.3E-14, FDR(mC2>mC15): 4.0E-6, FDR(mC4>mC5): 3.0E-16, FDR(mC4>mC15): 7.4E-8 |
| Ndufa4l2 | Hypoxia | [103] | mC11, mC15 | mC15 | FDR(mC11>mCo): 2.5E-20, FDR(mC15>mCo): <1E-50 | FDR(mC15>mC11): 1.5E-6 |
| Cd14 | Macrophages | [85] | mC3, mC8 | mC3, mC8 | FDR(mC3>mCo): <1E-50, FDR(mC8>mCo): 1.4E-11 | |
| Cd163 | Macrophages | [82] | mC3, mC5, mC8, mC16 | mC8 | FDR(mC3>mCo): 1.6E-4, FDR(mC5>mCo): 4.9E-3, FDR(mC8>mCo): 5.4E-4, FDR(mC16>mCo): 1.8E-3 | FDR(mC3>mC5): 9.6E-3, FDR(mC8>mC3): 9.8E-3, FDR(mC8>mC5): 5.6E-4, FDR(mC8>mC16): 2.7E-3 |
| Cd300a | Macrophages | [85] | mC3, mC8 | mC3, mC8 | FDR(mC3>mCo): <1E-50, FDR(mC8>mCo): 8.6E-12 | |
| Cd300d | Macrophages | [85] | mC3, mC8, mC14 | mC3 | FDR(mC3>mCo): <1E-50, FDR(mC8>mCo): 4.0E-10, FDR(mC14>mCo): 8.8E-4 | FDR(mC3>mC8): 2.4E-5, FDR(mC3>mC14): 1.4E-3 |
| Cd84 | Macrophages | [84] | mC3 | mC3 | FDR(mC3>mCo): <1E-50 | |
| Cd86 | Macrophages | [83] | mC3, mC8, mC14 | mC3, mC8, mC14 | FDR(mC3>mCo): <1E-50, FDR(mC8>mCo): 3.2E-12, FDR(mC14>mCo): 3.4E-7 | |
| Csf1r | Macrophages | [81] | mC3, mC8 | mC3 | FDR(mC3>mCo): <1E-50, FDR(mC8>mCo): 1.3E-16 | FDR(mC3>mC8): 4.8E-3 |
| Cd68 | Macrophages (Intermediary phenotype) | [85] | mC3, mC8, mC14 | mC3, mC14 | FDR(mC3>mCo): <1E-50, FDR(mC8>mCo): 2.6E-30, FDR(mC14>mCo): 6.9E-7 | FDR(mC3>mC8): 3.2E-4 |
| Cd209a | Macrophages (M2 phenotype, anti-inflammatory) | [112,85] | mC8 | mC8 | FDR(mC8>mCo): 4.0E-24 | |
| Cd209c | Macrophages (M2 phenotype, anti-inflammatory) | [112,85] | mC8 | mC8 | FDR(mC8>mCo): 4.1E-6 | |

| | | | | | | |
|---|---|---|---|---|---|---|
| Col12a1 | Mesenchymal | [77] | mC10, mC12, mC13 | mC10, mC13 | FDR(mC10>mCo): 1.2E-17, FDR(mC12>mCo): 1.0E-3, FDR(mC13>mCo): 4.4E-6 | FDR(mC10>mC12): 3.7E-15, FDR(mC13>mC12): 1.4E-5 |
| Col1a2 | Mesenchymal | [75,76] | mC6, mC10, mC12, mC13 | mC6, mC13 | FDR(mC6>mCo): 2.6E-25, FDR(mC10>mCo): 2.9E-15, FDR(mC12>mCo): 8.6E-3, FDR(mC13>mCo): 3.5E-11 | FDR(mC6>mC10): 5.4E-7, FDR(mC6>mC12): 1.4E-34, FDR(mC10>mC12): 1.3E-11, FDR(mC13>mC12): 1.2E-12 |
| Col6a3 | Mesenchymal | [76] | mC6, mC10, mC12 | mC6 | FDR(mC6>mCo): 1.9E-8, FDR(mC10>mCo): 6.4E-3, FDR(mC12>mCo): 9.1E-5 | FDR(mC6>mC10): 1.3E-5, FDR(mC6>mC12): 1.4E-6 |
| Cxcl12 | Mesenchymal | [74] | mC6, mC12, mC16 | mC6 | FDR(mC6>mCo): 4.5E-18, FDR(mC12>mCo): 1.3E-3, FDR(mC16>mCo): 9.2E-3 | FDR(mC6>mC12): 1.6E-19, FDR(mC6>mC16): 2.8E-19 |
| Dcn | Mesenchymal | [73] | mC6, mC13 | mC6, mC13 | FDR(mC6>mCo): 6.2E-13, FDR(mC13>mCo): 8.4E-15 | |
| Pdgfra | Mesenchymal | [70,71] | mC6, mC13 | mC6, mC13 | FDR(mC6>mCo): 4.2E-18, FDR(mC13>mCo): 1.2E-4 | |
| Pdgfrb | Mesenchymal | [72] | mC6, mC10 | mC6 | FDR(mC6>mCo): 5.3E-15, FDR(mC10>mCo): 5.8E-3 | FDR(mC6>mC10): 1.5E-12 |
| Prrx1 | Mesenchymal | [16] | mC6, mC13 | mC6, mC13 | FDR(mC6>mCo): 5.8E-9, FDR(mC13>mCo): 7.6E-4 | |
| Yap1 | Mesenchymal | [108] | mC1, mC2, mC7, mC16 | mC1, mC2, mC7, mC16 | FDR(mC1>mCo): 7.5E-8, FDR(mC2>mCo): 4.5E-4, FDR(mC7>mCo): 5.8E-3, FDR(mC16>mCo): 1.5E-3 | |
| Cldn11 | Migratory | [100] | mC1, mC6, mC7 | mC6 | FDR(mC1>mCo): 6.0E-9, FDR(mC6>mCo): 1.6E-8, FDR(mC7>mCo): 2.3E-3 | FDR(mC6>mC1): 3.2E-6, FDR(mC6>mC7): 1.3E-6 |
| Dock7 | Migratory | [101] | mC1, mC10 | mC10 | FDR(mC1>mCo): 4.3E-3, FDR(mC10>mCo): 3.0E-7 | FDR(mC10>mC1): 2.8E-4 |
| Lama3 | Migratory | [102] | mC1, mC2, mC4 | mC1, mC4 | FDR(mC1>mCo): 1.7E-10, FDR(mC2>mCo): 7.3E-3, FDR(mC4>mCo): 3.6E-6 | FDR(mC1>mC2): 6.6E-4, FDR(mC4>mC2): 5.0E-3 |
| Sema3e | Migratory | [99] | mC10 | mC10 | FDR(mC10>mCo): 1.5E-7 | |
| Erbb3 | Progenitors | [91] | mC10, mC15 | mC10 | FDR(mC10>mCo): 4.5E-37, FDR(mC15>mCo): 1.1E-7 | FDR(mC10>mC15): 3.4E-33 |
| Lgr5 | Progenitors | [96] | mC11, mC15 | mC15 | FDR(mC11>mCo): 4.4E-7, FDR(mC15>mCo): <1E-50 | FDR(mC15>mC11): 1.4E-7 |
| Neurod4 | Progenitors | [118] | mC11, mC15, mC17, mC19 | mC15 | FDR(mC11>mCo): 1.2E-5, FDR(mC15>mCo): 1.4E-30, FDR(mC17>mCo): 3.7E-3, FDR(mC19>mCo): 9.5E-9 | FDR(mC11>mC17): 1.4E-4, FDR(mC11>mC19): 6.9E-3, FDR(mC15>mC11): 2.4E-15, FDR(mC15>mC17): 1.7E-27, FDR(mC15>mC19): 2.7E-25, FDR(mC19>mC17): 9.5E-5 |
| Nf1 | Progenitors | [98] | mC2, mC10 | mC2, mC10 | FDR(mC2>mCo): 1.5E-6, FDR(mC10>mCo): 3.7E-5 | |
| Sox6 | Progenitors | [90] | mC1, mC7, mC10 | mC1, mC7, mC10 | FDR(mC1>mCo): 8.0E-12, FDR(mC7>mCo): 1.4E-3, FDR(mC10>mCo): 6.3E-6 | |
| Cartpt | Sympathoadrenal lineage | [6] | mC1, mC5, mC11, mC15, mC16 | mC11, mC15, mC16 | FDR(mC1>mCo): 4.9E-5, FDR(mC5>mCo): 1.0E-19, FDR(mC11>mCo): 6.1E-3, FDR(mC15>mCo): 3.2E-5, FDR(mC16>mCo): 1.4E-8 | FDR(mC5>mC1): 1.0E-13, FDR(mC11>mC1): 7.8E-3, FDR(mC15>mC1): 5.3E-4, FDR(mC15>mC5): 7.9E-3, FDR(mC16>mC1): 1.1E-5 |
| Gata3 | Sympathoadrenal lineage | [6] | mC11, mC15, mC17, mC18 | mC15 | FDR(mC11>mCo): 2.7E-9, FDR(mC15>mCo): 3.6E-39, FDR(mC17>mCo): 4.0E-9, FDR(mC18>mCo): 1.5E-3 | FDR(mC15>mC11): 4.7E-3, FDR(mC15>mC17): 1.3E-5, FDR(mC15>mC18): 5.6E-10 |
| Hand1 | Sympathoadrenal lineage | [6] | mC1, mC5, mC11, mC15, mC16 | mC11, mC15, mC16 | FDR(mC1>mCo): 1.1E-4, FDR(mC5>mCo): 1.5E-19, FDR(mC11>mCo): 3.1E-3, FDR(mC15>mCo): 1.5E-5, FDR(mC16>mCo): 3.7E-10 | FDR(mC5>mC1): 1.0E-13, FDR(mC11>mC1): 4.4E-3, FDR(mC15>mC1): 1.9E-4, FDR(mC15>mC5): 2.7E-3, FDR(mC16>mC1): 1.1E-6 |
| Phox2a | Sympathoadrenal lineage | [6] | mC11, mC15 | mC15 | FDR(mC11>mCo): 5.1E-9, FDR(mC15>mCo): 1.4E-38 | FDR(mC15>mC11): 5.3E-4 |
| Phox2b | Sympathoadrenal lineage | [6] | mC10, mC11, mC15 | mC15 | FDR(mC10>mCo): 2.0E-4, FDR(mC11>mCo): 1.2E-8, FDR(mC15>mCo): <1E-50 | FDR(mC15>mC10): 5.4E-11, FDR(mC15>mC11): 2.1E-6 |
| Stmn2 | Sympathoadrenal lineage | [6] | mC11, mC15 | mC11, mC15 | FDR(mC11>mCo): 9.2E-16, FDR(mC15>mCo): 1.5E-37 | |

| | | | | | | |
|---|---|---|---|---|---|---|
| Cd3e | T-cells | [86] | mC1, mC5, mC15, mC16, mC17 | mC17 | FDR(mC1>mCo): 4.4E-10, FDR(mC5>mCo): 5.2E-33, FDR(mC15>mCo): 1.2E-6, FDR(mC16>mCo): 1.5E-14, FDR(mC17>mCo): 1.9E-32 | FDR(mC5>mC1): 4.9E-11, FDR(mC16>mC1): 1.1E-8, FDR(mC16>mC5): 1.2E-3, FDR(mC17>mC1): 2.8E-30, FDR(mC17>mC5): 1.7E-28, FDR(mC17>mC15): 2.5E-30, FDR(mC17>mC16): 4.1E-27 |
| Hcst | T-cells | [88] | mC3, mC5, mC15, mC16, mC17, mC18 | mC18 | FDR(mC3>mCo): 5.6E-6, FDR(mC5>mCo): 6.1E-19, FDR(mC15>mCo): 3.4E-3, FDR(mC16>mCo): 1.3E-32, FDR(mC17>mCo): 5.3E-11, FDR(mC18>mCo): 3.3E-25 | FDR(mC16>mC5): 3.7E-6, FDR(mC16>mC15): 1.9E-6, FDR(mC17>mC3): 2.8E-6, FDR(mC17>mC5): 3.0E-7, FDR(mC17>mC15): 2.2E-8, FDR(mC17>mC16): 2.5E-5, FDR(mC18>mC3): 1.8E-21, FDR(mC18>mC5): 3.7E-21, FDR(mC18>mC15): 1.5E-23, FDR(mC18>mC16): 6.4E-19, FDR(mC18>mC17): 5.0E-4 |
| Trac | T-cells | [89] | mC1, mC5, mC15, mC16, mC17 | mC17 | FDR(mC1>mCo): 8.2E-15, FDR(mC5>mCo): 3.4E-39, FDR(mC15>mCo): 1.4E-10, FDR(mC16>mCo): <1E-50, FDR(mC17>mCo): 9.4E-30 | FDR(mC5>mC1): 2.2E-11, FDR(mC16>mC1): 1.3E-23, FDR(mC16>mC5): 3.7E-6, FDR(mC16>mC15): 2.2E-8, FDR(mC17>mC1): 2.8E-27, FDR(mC17>mC5): 2.1E-25, FDR(mC17>mC15): 5.0E-27, FDR(mC17>mC16): 6.8E-24 |
| Trbc1 | T-cells | [89] | mC1, mC5, mC15, mC16, mC17, mC18 | mC17, mC18 | FDR(mC1>mCo): 4.4E-13, FDR(mC5>mCo): 8.2E-37, FDR(mC15>mCo): 1.3E-9, FDR(mC16>mCo): 2.4E-42, FDR(mC17>mCo): <1E-50, FDR(mC18>mCo): 2.5E-37 | FDR(mC5>mC1): 1.0E-13, FDR(mC16>mC1): 4.6E-24, FDR(mC16>mC5): 3.2E-6, FDR(mC16>mC15): 2.3E-8, FDR(mC17>mC1): <1E-50, FDR(mC17>mC5): 2.4E-41, FDR(mC17>mC15): <1E-50, FDR(mC17>mC16): 1.1E-38, FDR(mC18>mC1): 3.3E-34, FDR(mC18>mC5): 2.3E-31, FDR(mC18>mC15): 2.5E-36, FDR(mC18>mC16): 5.6E-29 |
| Trbc2 | T-cells | [89] | mC1, mC5, mC15, mC16, mC17, mC18 | mC17 | FDR(mC1>mCo): 9.1E-14, FDR(mC5>mCo): 1.7E-37, FDR(mC15>mCo): 2.0E-9, FDR(mC16>mCo): 7.1E-40, FDR(mC17>mCo): <1E-50, FDR(mC18>mCo): 9.9E-41 | FDR(mC5>mC1): 1.0E-13, FDR(mC5>mC15): 2.0E-3, FDR(mC16>mC1): 7.7E-23, FDR(mC16>mC5): 3.1E-6, FDR(mC16>mC15): 1.2E-10, FDR(mC17>mC1): <1E-50, FDR(mC17>mC5): <1E-50, FDR(mC17>mC15): <1E-50, FDR(mC17>mC16): <1E-50, FDR(mC17>mC18): 5.2E-4, FDR(mC18>mC1): 7.3E-37, FDR(mC18>mC5): 9.7E-34, FDR(mC18>mC15): 3.7E-40, FDR(mC18>mC16): 6.2E-31 |
| Cd7 | T-cells (active) | [113] | mC1, mC5, mC15, mC16, mC18 | mC18 | FDR(mC1>mCo): 4.5E-12, FDR(mC5>mCo): 1.4E-35, FDR(mC15>mCo): 1.5E-9, FDR(mC16>mCo): 1.6E-29, FDR(mC18>mCo): <1E-50 | FDR(mC5>mC1): 1.0E-13, FDR(mC5>mC15): 5.7E-4, FDR(mC16>mC1): 1.0E-16, FDR(mC16>mC5): 7.3E-5, FDR(mC16>mC15): 6.6E-10, FDR(mC18>mC1): 3.1E-42, FDR(mC18>mC5): 6.3E-41, FDR(mC18>mC15): <1E-50, FDR(mC18>mC16): 1.2E-41 |
| Mycn | Unfavourable outcome | [2,104] | mC15 | mC15 | FDR(mC15>mCo): 3.8E-7 | |

| | | | | | | |
|---|---|---|---|---|---|---|
| Cyp11a1 | Zona fasciculata (Cortex) | [49] | mC1, mC5, mC7, mC9, mC16 | mC7, mC9 | FDR(mC1>mCo): <1E-50, FDR(mC5>mCo): <1E-50, FDR(mC7>mCo): <1E-50, FDR(mC9>mCo): <1E-50, FDR(mC16>mCo): <1E-50 | FDR(mC1>mC5): 8.6E-3, FDR(mC1>mC16): 6.9E-9, FDR(mC5>mC16): 5.3E-3, FDR(mC7>mC1): 5.5E-36, FDR(mC7>mC5): 1.4E-37, FDR(mC7>mC16): 2.7E-42, FDR(mC9>mC1): 1.1E-16, FDR(mC9>mC5): 7.5E-20, FDR(mC9>mC16): 1.0E-23 |
| Cyp11b1 | Zona fasciculata (Cortex) | [50,51,52] | mC1, mC5, mC7, mC9, mC16 | mC9 | FDR(mC1>mCo): 1.6E-15, FDR(mC5>mCo): 5.4E-26, FDR(mC7>mCo): 2.1E-25, FDR(mC9>mCo): <1E-50, FDR(mC16>mCo): <1E-50 | FDR(mC5>mC1): 2.3E-7, FDR(mC7>mC1): 1.3E-9, FDR(mC9>mC1): 2.8E-40, FDR(mC9>mC5): 3.0E-16, FDR(mC9>mC7): 1.1E-9, FDR(mC9>mC16): 5.2E-20, FDR(mC16>mC1): 1.4E-18 |
| Star | Zona fasciculata (Cortex) & Zona reticularis (Cortex) | [53] | mC1, mC5, mC7, mC9, mC16 | mC7, mC9 | FDR(mC1>mCo): <1E-50, FDR(mC5>mCo): <1E-50, FDR(mC7>mCo): <1E-50, FDR(mC9>mCo): 1.4E-39, FDR(mC16>mCo): <1E-50 | FDR(mC1>mC16): 5.3E-4, FDR(mC7>mC1): 5.5E-36, FDR(mC7>mC5): 1.0E-37, FDR(mC7>mC16): 4.1E-36, FDR(mC9>mC1): 4.0E-15, FDR(mC9>mC5): 1.9E-17, FDR(mC9>mC16): 2.3E-19 |
| Agtr1b | Zona glomerulosa (Cortex) | [57,58,59] | mC1, mC5, mC7 | mC1,mC7 | FDR(mC1>mCo): <1E-50, FDR(mC5>mCo): 1.3E-14, FDR(mC7>mCo): 1.1E-24 | FDR(mC1>mC5): 1.8E-34, FDR(mC7>mC5): 6.7E-11 |
| Cyp11b2 | Zona glomerulosa (Cortex) | [50,51] | mC1, mC5, mC7, mC9, mC16 | mC7, mC9 | FDR(mC1>mCo): 2.0E-29, FDR(mC5>mCo): 3.1E-25, FDR(mC7>mCo): 2.1E-21, FDR(mC9>mCo): <1E-50, FDR(mC16>mCo): <1E-50 | FDR(mC7>mC1): 8.1E-3, FDR(mC7>mC5): 2.2E-4, FDR(mC7>mC16): 1.1E-4, FDR(mC9>mC1): 5.7E-11, FDR(mC9>mC5): 1.3E-16, FDR(mC9>mC16): 7.2E-21 |
| Dab2 | Zona glomerulosa (Cortex) | [55,56] | mC1, mC3, mC5, mC7, mC8 | mC1,mC7 | FDR(mC1>mCo): <1E-50, FDR(mC3>mCo): <1E-50, FDR(mC5>mCo): 5.9E-16, FDR(mC7>mCo): <1E-50, FDR(mC8>mCo): 1.4E-11 | FDR(mC1>mC3): 5.3E-18, FDR(mC1>mC5): 1.5E-16, FDR(mC1>mC8): 1.3E-3, FDR(mC3>mC5): 1.4E-6, FDR(mC7>mC3): 7.9E-11, FDR(mC7>mC5): 7.7E-17, FDR(mC7>mC8): 4.6E-4 |

**Markers in NB**

| Neuroblastoma (NB) marker | Cell type | References | Evidence for high expression in NB cluster | Evidence for higher expression in NB cluster | Each cluster in NB: FDR | Cluster pairwise comparison in NB: FDR |
|---|---|---|---|---|---|---|
| CHGA | Adrenergic (Adrenal medulla) | [54,63] | nC5,nC7,nC8 | nC7 | FDR(nC5>nCo): 2.6E-18, FDR(nC7>nCo): <1E-50, FDR(nC8>nCo): 7.4E-15 | FDR(nC7>nC5): 6.6E-16, FDR(nC7>nC8): 2.3E-7 |
| CHGB | Adrenergic (Adrenal medulla) | [54,63] | nC5,nC7,nC9 | nC9 | FDR(nC5>nCo): <1E-50, FDR(nC7>nCo): <1E-50, FDR(nC9>nCo): <1E-50 | FDR(nC7>nC5): 7.6E-3, FDR(nC9>nC5): 7.4E-19, FDR(nC9>nC7): 3.7E-8 |
| DBH | Adrenergic (Adrenal medulla) | [54,64] | nC5,nC7,nC8,nC9 | nC7 | FDR(nC5>nCo): <1E-50, FDR(nC7>nCo): <1E-50, FDR(nC8>nCo): <1E-50, FDR(nC9>nCo): <1E-50 | FDR(nC7>nC5): 2.8E-7, FDR(nC7>nC8): 5.0E-8, FDR(nC7>nC9): 3.7E-4 |
| TH | Adrenergic (Adrenal medulla) | [54,62] | nC5,nC7,nC8,nC9 | nC7 | FDR(nC5>nCo): <1E-50, FDR(nC7>nCo): <1E-50, FDR(nC8>nCo): 2.0E-42, FDR(nC9>nCo): 1.3E-42 | FDR(nC7>nC5): 5.9E-13, FDR(nC7>nC8): 2.1E-6, FDR(nC7>nC9): 4.9E-7 |
| IGHM | B-cells | [116] | nC10 | nC10 | FDR(nC10>nCo): 3.2E-3 | |
| MS4A1 | B-cells | [114,115] | nC2,nC3 | nC2,nC3 | FDR(nC2>nCo): 4.1E-3, FDR(nC3>nCo): 8.2E-9 | |
| HTR3A | Bridge | [6] | nC8 | nC8 | FDR(nC8>nCo): 2.1E-16 | |
| GLI1 | Capsule | [109] | nC1,nC2 | nC1,nC2 | FDR(nC1>nCo): 3.5E-3, FDR(nC2>nCo): 1.1E-7 | |
| RSPO3 | Capsule | [109] | nC1,nC2,nC4 | nC1,nC2,nC4 | FDR(nC1>nCo): 3.4E-4, FDR(nC2>nCo): 1.3E-3, FDR(nC4>nCo): 1.3E-4 | |
| MKI67 | Cell cycle | | nC5,nC9 | nC5,nC9 | FDR(nC5>nCo): 9.1E-3, FDR(nC9>nCo): 1.4E-4 | |

| | | | | | | |
|---|---|---|---|---|---|---|
| CHRNA3 | Chromaffin | [65,67] | nC5,nC7,nC8,nC9 | nC8 | FDR(nC5>nCo): 2.5E-18, FDR(nC7>nCo): 3.2E-10, FDR(nC8>nCo): <1E-50, FDR(nC9>nCo): 4.8E-20 | FDR(nC8>nC5): 9.4E-28, FDR(nC8>nC7): 2.4E-10, FDR(nC8>nC9): 1.4E-6, FDR(nC9>nC5): 2.7E-4 |
| CHRNB4 | Chromaffin | [65,67] | nC8 | nC8 | FDR(nC8>nCo): 4.5E-3 | |
| NPY | Chromaffin | [63] | nC5,nC7,nC8,nC9 | nC7 | FDR(nC5>nCo): 3.9E-12, FDR(nC7>nCo): <1E-50, FDR(nC8>nCo): 8.6E-32, FDR(nC9>nCo): 2.6E-16 | FDR(nC7>nC5): <1E-50, FDR(nC7>nC8): 3.1E-17, FDR(nC7>nC9): 7.5E-15, FDR(nC8>nC5): 2.1E-8, FDR(nC9>nC5): 1.1E-4 |
| CHRNA7 | Chromaffin NE-secreting | [65,66,67,68] | nC5,nC7,nC8,nC9 | nC7, nC8 | FDR(nC5>nCo): 1.4E-35, FDR(nC7>nCo): <1E-50, FDR(nC8>nCo): <1E-50, FDR(nC9>nCo): 1.1E-25 | FDR(nC7>nC5): 4.0E-9, FDR(nC7>nC9): 1.2E-4, FDR(nC8>nC5): 7.4E-12, FDR(nC8>nC9): 5.7E-6 |
| DLK1 | Differentiation | [2] | nC3,nC8 | nC8 | FDR(nC3>nCo): 1.0E-3, FDR(nC8>nCo): <1E-50 | FDR(nC8>nC3): 6.9E-28 |
| PROM1 | Differentiation | [106] | nC2,nC3 | nC2,nC3 | FDR(nC2>nCo): 2.0E-5, FDR(nC3>nCo): 1.1E-18 | |
| RET | Differentiation | [106] | nC2 | nC2 | FDR(nC2>nCo): 7.6E-4 | |
| STMN4 | Differentiation | [120] | nC5,nC7,nC8,nC9 | nC9 | FDR(nC5>nCo): 8.1E-16, FDR(nC7>nCo): 2.1E-15, FDR(nC8>nCo): 2.9E-7, FDR(nC9>nCo): 9.9E-31 | FDR(nC7>nC5): 8.1E-3, FDR(nC9>nC5): 8.3E-11, FDR(nC9>nC7): 2.4E-3, FDR(nC9>nC8): 1.7E-8 |
| TP63 | Differentiation | [105] | nC3 | nC3 | FDR(nC3>nCo): 1.8E-24 | |
| PECAM1 | Endothelial | [78] | nC3,nC4 | nC4 | FDR(nC3>nCo): 5.9E-19, FDR(nC4>nCo): 4.3E-29 | FDR(nC4>nC3): 5.9E-14 |
| SOX17 | Endothelial | | nC3,nC4,nC8 | nC4 | FDR(nC3>nCo): 1.3E-3, FDR(nC4>nCo): 4.9E-8, FDR(nC8>nCo): 1.2E-3 | FDR(nC4>nC3): 3.1E-6, FDR(nC4>nC8): 2.9E-6 |
| SOX18 | Endothelial | [80] | nC4 | nC4 | FDR(nC4>nCo): 1.9E-4 | |
| TEK | Endothelial | [78] | nC3,nC4 | nC3,nC4 | FDR(nC3>nCo): 1.4E-13, FDR(nC4>nCo): 2.3E-4 | |
| TIE1 | Endothelial | [79] | nC4 | nC4 | FDR(nC4>nCo): 6.7E-21 | |
| SOX4 | Endothelial (tumor-induced angiogenesis) | [119] | nC5,nC7,nC8,nC9 | nC7 | FDR(nC5>nCo): <1E-50, FDR(nC7>nCo): <1E-50, FDR(nC8>nCo): <1E-50, FDR(nC9>nCo): <1E-50 | FDR(nC7>nC5): 1.0E-15, FDR(nC7>nC8): 2.2E-32, FDR(nC7>nC9): 3.0E-22 |
| NTRK1 | Favourable outcome | [2] | nC5,nC7,nC8,nC9 | nC7 | FDR(nC5>nCo): 4.7E-32, FDR(nC7>nCo): <1E-50, FDR(nC8>nCo): 2.5E-29, FDR(nC9>nCo): 1.7E-15 | FDR(nC7>nC5): 1.4E-32, FDR(nC7>nC8): 3.3E-16, FDR(nC7>nC9): 4.0E-15 |
| COX4I2 | Hypoxia | [103] | nC1,nC2,nC4 | nC1, nC2 | FDR(nC1>nCo): 9.5E-6, FDR(nC2>nCo): 2.4E-5, FDR(nC4>nCo): 9.0E-6 | FDR(nC1>nC4): 3.3E-3 |
| EPAS1 | Hypoxia | [103] | nC4,nC8,nC9 | nC4 | FDR(nC4>nCo): 6.7E-43, FDR(nC8>nCo): 7.0E-7, FDR(nC9>nCo): 9.0E-19 | FDR(nC4>nC8): 3.9E-19, FDR(nC4>nC9): 2.4E-6, FDR(nC9>nC8): 1.9E-4 |
| NDUFA4L2 | Hypoxia | [103] | nC5,nC7 | nC7 | FDR(nC5>nCo): 9.7E-3, FDR(nC7>nCo): 2.0E-22 | FDR(nC7>nC5): 3.3E-15 |
| CD14 | Macrophages | [85] | nC6 | nC6 | FDR(nC6>nCo): 1.3E-9 | |
| CD163 | Macrophages | [82] | nC3,nC6 | nC6 | FDR(nC3>nCo): 6.2E-13, FDR(nC6>nCo): 1.7E-11 | FDR(nC6>nC3): 7.2E-6 |
| CD300A | Macrophages | [85] | nC2,nC6 | nC6 | FDR(nC2>nCo): 7.4E-4, FDR(nC6>nCo): 2.6E-10 | FDR(nC6>nC2): 7.9E-8 |
| CD84 | Macrophages | [84] | nC3,nC6 | nC6 | FDR(nC3>nCo): 1.2E-12, FDR(nC6>nCo): 3.2E-14 | FDR(nC6>nC3): 6.7E-8 |
| CD86 | Macrophages | [83] | nC3,nC6 | nC6 | FDR(nC3>nCo): 2.0E-3, FDR(nC6>nCo): 4.2E-10 | FDR(nC6>nC3): 1.6E-7 |
| CSF1R | Macrophages | [81] | nC2,nC6 | nC6 | FDR(nC2>nCo): 4.5E-3, FDR(nC6>nCo): 6.8E-7 | FDR(nC6>nC2): 1.0E-4 |
| COL12A1 | Mesenchymal | [77] | nC1,nC3 | nC1,nC3 | FDR(nC1>nCo): 2.5E-3, FDR(nC3>nCo): <1E-50 | |
| COL1A2 | Mesenchymal | [75,76] | nC1,nC3,nC9 | nC1 | FDR(nC1>nCo): 3.6E-13, FDR(nC3>nCo): 2.2E-12, FDR(nC9>nCo): 1.2E-4 | FDR(nC1>nC3): 3.8E-6, FDR(nC1>nC9): 6.5E-5 |
| COL6A3 | Mesenchymal | [76] | nC1,nC3 | nC1 | FDR(nC1>nCo): 8.1E-14, FDR(nC3>nCo): 1.3E-13 | FDR(nC1>nC3): 1.9E-6 |
| PDGFRA | Mesenchymal | [70,71] | nC1,nC3 | nC1,nC3 | FDR(nC1>nCo): 8.7E-4, FDR(nC3>nCo): 1.0E-19 | |
| PDGFRB | Mesenchymal | [72] | nC1 | nC1 | FDR(nC1>nCo): 7.0E-8 | |
| PRRX1 | Mesenchymal | [16] | nC1,nC3 | nC1 | FDR(nC1>nCo): 8.4E-9, FDR(nC3>nCo): 4.5E-8 | FDR(nC1>nC3): 3.0E-4 |
| YAP1 | Mesenchymal | [108] | nC1,nC3 | nC1,nC3 | FDR(nC1>nCo): 2.7E-3, FDR(nC3>nCo): 1.2E-17 | |

| | | | | | | |
|---|---|---|---|---|---|---|
| CLDN11 | Migratory | [100] | nC1,nC2,nC3 | nC3 | FDR(nC1>nCo): 6.3E-3, FDR(nC2>nCo): 7.1E-19, FDR(nC3>nCo): <1E-50 | FDR(nC3>nC1): 3.6E-9, FDR(nC3>nC2): 2.0E-5 |
| DOCK7 | Migratory | [101] | nC3 | nC3 | FDR(nC3>nCo): 4.1E-34 | |
| LAMA3 | Migratory | [102] | nC3 | nC3 | FDR(nC3>nCo): 1.2E-41 | |
| SEMA3E | Migratory | [99] | nC3 | nC3 | FDR(nC3>nCo): 4.4E-22 | |
| NTRK3 | Neuroblastoma | [2] | nC3,nC5,nC7,nC8 | nC3,nC5,nC7,nC8 | FDR(nC3>nCo): 8.6E-9, FDR(nC5>nCo): 5.1E-3, FDR(nC7>nCo): 2.2E-4, FDR(nC8>nCo): 3.3E-7 | |
| ASXL3 | Progenitors | [95] | nC3,nC7,nC8 | nC3,nC7,nC8 | FDR(nC3>nCo): 2.3E-4, FDR(nC7>nCo): 1.6E-3, FDR(nC8>nCo): 2.9E-4 | |
| BCL11A | Progenitors | [94] | nC3 | nC3 | FDR(nC3>nCo): 7.2E-9 | |
| ERBB3 | Progenitors | [91] | nC2,nC3 | nC2,nC3 | FDR(nC2>nCo): 1.1E-6, FDR(nC3>nCo): 4.5E-5 | |
| LGR5 | Progenitors | [96] | nC3 | nC3 | FDR(nC3>nCo): 1.0E-10 | |
| NF1 | Progenitors | [98] | nC3 | nC3 | FDR(nC3>nCo): 9.7E-28 | |
| POU6F2 | Progenitors | [93] | nC2,nC3 | nC3 | FDR(nC2>nCo): 1.0E-3, FDR(nC3>nCo): 1.2E-17 | FDR(nC3>nC2): 6.2E-3 |
| RTTN | Progenitors | [92] | nC2,nC3 | nC3 | FDR(nC2>nCo): 3.0E-3, FDR(nC3>nCo): <1E-50 | FDR(nC3>nC2): 3.3E-12 |
| SOX5 | Progenitors | [97] | nC3 | nC3 | FDR(nC3>nCo): 7.2E-28 | |
| SOX6 | Progenitors | [90] | nC3,nC9 | nC3,nC9 | FDR(nC3>nCo): 1.8E-23, FDR(nC9>nCo): 3.8E-3 | |
| CARTPT | Sympathoadrenal lineage | [6] | nC1,nC2,nC4,nC8 | nC1, nC8 | FDR(nC1>nCo): 9.9E-32, FDR(nC2>nCo): 5.0E-7, FDR(nC4>nCo): 2.3E-9, FDR(nC8>nCo): 1.4E-3 | FDR(nC1>nC2): 1.9E-3, FDR(nC1>nC4): 5.7E-6 |
| GATA3 | Sympathoadrenal lineage | [6] | nC5,nC7,nC8,nC9 | nC7 | FDR(nC5>nCo): <1E-50, FDR(nC7>nCo): <1E-50, FDR(nC8>nCo): <1E-50, FDR(nC9>nCo): 1.6E-24 | FDR(nC7>nC5): 3.1E-14, FDR(nC7>nC8): 2.7E-12, FDR(nC7>nC9): 3.2E-14 |
| HAND1 | Sympathoadrenal lineage | [6] | nC5,nC8,nC9 | nC5,nC8,nC9 | FDR(nC5>nCo): 2.1E-5, FDR(nC8>nCo): 2.6E-3, FDR(nC9>nCo): 3.2E-4 | |
| ISL1 | Sympathoadrenal lineage | [6] | nC5,nC7,nC8,nC9 | nC7, nC8, nC9 | FDR(nC5>nCo): 6.7E-43, FDR(nC7>nCo): 3.7E-27, FDR(nC8>nCo): <1E-50, FDR(nC9>nCo): 7.1E-38 | FDR(nC8>nC5): 1.2E-11, FDR(nC9>nC5): 9.5E-5 |
| PHOX2A | Sympathoadrenal lineage | [69] | nC5,nC7,nC8,nC9,nC10 | nC7, nC9, nC10 | FDR(nC5>nCo): 3.2E-10, FDR(nC7>nCo): 2.3E-21, FDR(nC8>nCo): 4.4E-3, FDR(nC9>nCo): 1.4E-20, FDR(nC10>nCo): 3.0E-4 | FDR(nC7>nC5): 2.8E-7, FDR(nC7>nC8): 4.8E-7, FDR(nC9>nC5): 1.1E-7, FDR(nC9>nC8): 1.9E-7 |
| PHOX2B | Sympathoadrenal lineage | [69] | nC5,nC7,nC8,nC9 | nC7, nC8, nC9 | FDR(nC5>nCo): 1.3E-42, FDR(nC7>nCo): 1.0E-31, FDR(nC8>nCo): 1.9E-35, FDR(nC9>nCo): 2.4E-16 | FDR(nC7>nC5): 5.1E-3 |
| STMN2 | Sympathoadrenal lineage | [6] | nC5,nC7,nC8,nC9 | nC7 | FDR(nC5>nCo): <1E-50, FDR(nC7>nCo): <1E-50, FDR(nC8>nCo): <1E-50, FDR(nC9>nCo): <1E-50 | FDR(nC5>nC8): 5.1E-7, FDR(nC5>nC9): 9.0E-6, FDR(nC7>nC5): 8.0E-6, FDR(nC7>nC8): 3.5E-31, FDR(nC7>nC9): 4.5E-24 |
| PRPH | Sympathoblast | [6] | nC5,nC7,nC8,nC9 | nC7,nC9 | FDR(nC5>nCo): 2.7E-6, FDR(nC7>nCo): <1E-50, FDR(nC8>nCo): <1E-50, FDR(nC9>nCo): 2.0E-42 | FDR(nC7>nC5): 5.7E-38, FDR(nC7>nC8): 5.4E-9, FDR(nC8>nC5): 3.9E-16, FDR(nC9>nC5): 1.1E-19 |
| CD69 | T-cells | [113] | nC10 | nC10 | FDR(nC10>nCo): 1.8E-4 | |
| IL32 | T-cells | [87] | nC10 | nC10 | FDR(nC10>nCo): 3.0E-4 | |
| TRAC | T-cells | [89] | nC10 | nC10 | FDR(nC10>nCo): 7.4E-4 | |
| TRBC1 | T-cells | [89] | nC10 | nC10 | FDR(nC10>nCo): 1.3E-6 | |
| TRBC2 | T-cells | [89] | nC10 | nC10 | FDR(nC10>nCo): 6.0E-8 | |
| ALK | Unfavourable outcome | [2] | nC3 | nC3 | FDR(nC3>nCo): <1E-50 | |
| BRCA1 | Unfavourable outcome | [104] | nC3 | nC3 | FDR(nC3>nCo): 1.5E-20 | |
| MYCN | Unfavourable outcome | [2,104] | nC1,nC3 | nC1,nC3 | FDR(nC1>nCo): 2.7E-5, FDR(nC3>nCo): 1.3E-12 | |
| NTRK2 | Unfavourable outcome | [2] | nC3 | nC3 | FDR(nC3>nCo): 4.6E-30 | |
| CYP11B1 | Zona fasciculata (Cortex) | [50,51,52] | nC2 | nC2 | FDR(nC2>nCo): 1.0E-6 | |
| STAR | Zona fasciculata (Cortex) & Zona reticularis (Cortex) | [53,54] | nC2 | nC2 | FDR(nC2>nCo): 4.4E-7 | |

| | | | | | |
|---|---|---|---|---|---|
| CYP11B2 | Zona glomerulosa (Cortex) | [54,50,51] | nC2 | nC2 | FDR(nC2>nCo): 2.3E-3 |

**Specific gene signature in human post-natal AG**

<table>
<tr><th>Human adrenal gland cluster</th><th>Number of genes in the specific signature</th><th>Specific gene signature</th></tr>
<tr><td>Progenitor (hC1)</td><td>508</td><td>A1CF|A2ML1|ABCA13|ABCA4|ABCB5|ABCC12|ABCC2|ABCG8|AC003664.1|AC004862.6|AC006994.2|AC007064.24|AC007405.4|AC007731.1|AC008079.9|AC008103.3|AC008103.4|AC008132.15|AC008697.1|AC008993.3|AC009264.1|AC011718.2|AC012368.2|AC074093.1|AC074391.1|AC079807.4|AC099057.8|AC108025.2|AC139099.5|AC144521.1|AC145110.1|ACOXL|ACSM2A|ACSM2B|ADAM18|ADAM32|ADAM7|ADAMTS20|ADAMTS6|ADCY10|ADGRF1|AFF2|AJ006998.2|AL035610.2|AMER2|ANKRD30A|ANKRD30BL|ANLN|AP000235.2|AP000432.3|AP003900.6|ARHGAP28|ARL14EPL|ARMC3|ARMC4|ARPP21|ASB15|ASPM|ASXL3|ATP13A5|ATP6V0A4|B4GALNT2|BPIFC|BRCA1|BUB1|BUB1B|C12orf42|C12orf45|C14orf23|C1orf141|C1orf87|C2orf58|C2orf78|C4BPA|C5orf64|CACNG8|CASQ1|CCDC150|CCDC168|CCDC26|CCDC83|CECR2|CENPI|CFAP54|CFAP57|CFAP58|CFTR|CH17-264L24.1|CH507-154B10.1|CH507-154B10.2|CH507-513H4.1|CH507-528H12.1|CHRM2|CHST9|CLDN11|CLNK|CMYA5|CNKSR3|CNTN5|COBL|COL19A1|COL21A1|COL24A1|COL25A1|COL6A5|COL9A1|COLEC10|CORIN|CR2|CSMD2|CSMD3|CT45A1|CT45A10|CT45A2|CT45A3|CT45A5|CT45A6|CT45A7|CT45A8|CT45A9|CTA-212A2.3|CTA-243E7.4|CTC-312O10.3|CTD-2036A2.1|CTD-2144E22.9|CTD-2147F2.2|CTD-2311B13.5|CTD-2311B13.7|CTD-2626G11.2|CTD-3088G3.8|CTNNA3|CUBN|CYP8B1|DCC|DCDC1|DDX4|DEC1|DEFB108B|DEFB119|DIAPH3|DLX6-AS1|DNAH10|DNAH11|DNAH2|DNAH3|DNAH9|DNTT|DOCK7|DRAIC|DRP2|DSC1|DSG2|DSG3|EFCAB5|EFCAB8|EFL1|EGF|EGFL6|ELF3|ENAM|EPHA6|EPM2A|ERBB3|ERVW-1|ESCO2|ESRG|FAM129C|FAM230A|FAM230B|FGF5|FREM2|FRG2|FRG2B|FRK|FRMPD4|GABRA2|GABRA3|GABRG1|GBP6|GC|GDNF-AS1|GFAP|GOLGA6L1|GOLGA6L22|GOLGA6L6|GRM1|GS1-122H1.2|GSDMC|GYPA|HAVCR1|HCG23|HELLPAR|HMCN1|HMGA2|HNF4G|HTN1|HULC|IGFN1|ITGB6|ITIH5|KB-1183D5.13|KCNU1|KIAA1257|KIF14|KIF15|KIF4A|KIF6|KLHL33|KNTC1|KRT39|LA16c-23H5.4|LA16c-3G11.5|LA16c-60D12.1|LA16c-60D12.2|LAMA1|LAMA3|LINC00466|LINC00506|LINC00710|LINC00923|LOXHD1|LPA|LRAT|LRP1B|LVRN|MALRD1|MCM10|MEP1A|MGAM|MME|MMS22L|MPP4|MTUS2|MUC16|MUC19|MUC3A|MYH15|MYLK3|MYLK4|MYO18B|MYO1H|MYO3A|MYO5C|MYOCD|MYPN|NAALADL2|NANOG|NCAN|NCAPH|NEB|NEK10|NEK2|NELL1|NF1|NLN|NPFFR1|NPSR1-AS1|OR10A2|OR10K1|OR14J1|OR1A1|OR1D2|OR2M3|OR2T12|OR2T33|OR4F17|OR4F4|OR4F5|OR51L1|OR56A4|OR5A1|OR5AS1|OR7A5|OR7C1|OR7D2|OR7D4|OR8A1|OTOA|OTOGL|PAPPA2|PCAT1|PCAT14|PCSK5|PDE3A|PDE6A|PGM2L1|PHEX|PHGDH|PIEZO2|PIWIL3|PKD1L1|PKD1L2|PKHD1|PLCH1|PLEKHD1|POLQ|POTEB|POTEB3|POTED|POU2F3|PPEF1|PRAMEF11|PRAMEF13|PRAMEF15|PRAMEF2|PRAMEF25|PRAMEF26|PRAMEF27|PRAMEF4|PRAMEF5|PRAMEF6|PRKG2|PSG1|PSG11|PSG2|PSG3|PSG4|PSG6|PSG7|PSG8|PSG9|PTH2R|PTPRD|PTPRQ|PTPRT|PTPRZ1|RASSF6|RBM20|REG4|RF00012|RF00017|RF00019|RGR|RHPN2|RN7SKP247|ROPN1|ROS1|RP1-101G11.3|RP1-14N1.2|RP1-60O19.1|RP11-1038A11.2|RP11-114G22.1|RP11-115D19.1|RP11-118H15.1|RP11-120D5.1|RP11-139E24.1|RP11-142I20.1|RP11-146E13.4|RP11-146I2.1|RP11-153K16.1|RP11-15E1.5|RP11-161I6.2|RP11-168K9.1|RP11-169F17.1|RP11-210M15.2|RP11-215N21.1|RP11-21F16.1|RP11-244H18.1|RP11-24J23.2|RP11-255B23.3|RP11-267C16.1|RP11-280K24.4|RP11-2A4.3|RP11-2A4.4|RP11-306O1.2|RP11-30J20.1|RP11-33H15.1|RP11-341D18.3|RP11-344E13.3|RP11-34P13.13|RP11-34P13.7|RP11-374A4.1|RP11-385D13.1|RP11-395L14.18|RP11-402N8.1|RP11-407B7.1|RP11-410C4.5|RP11-416O18.1|RP11-430C1.1|RP11-438L7.1|RP11-458B24.2|RP11-459O16.8|RP11-461F11.2|RP11-482G13.1|RP11-495L19.1|RP11-4L24.3|RP11-506N2.1|RP11-515E23.1|RP11-532N4.2|RP11-536C10.7|RP11-541P9.3|RP11-544I20.2|RP11-54H7.4|RP11-553E24.2|RP11-573D15.8|RP11-586K12.1|RP11-597A11.2|RP11-624L4.1|RP11-631B21.1|RP11-632K5.2|RP11-65B7.1|RP11-673E1.1|RP11-69H14.6|RP11-69I8.2|RP11-728G15.1|RP11-733C7.2|RP11-735B13.1|RP11-739N10.1|RP11-73M7.10|RP11-774D14.1|RP11-776H12.1|RP11-886D15.1|RP11-89M16.1|RP11-941H19.3|RP11-96K19.4|RP13-262C2.4|RP3-454G6.2|RP4-671O14.7|RP4-734C18.1|RP4-760C5.5|RP4-809F18.1|RP5-1014D13.2|RP5-898I4.1|RP5-916L7.1|RP5-947P14.1|RTKN2|SAG|SALRNA2|SCLY|SCN11A|SEMA3E|SFRP4|SGO2|SH2D4B|SH3GL3|SH3RF2|SI|SLC13A1|SLC14A1|SLC17A4|SLC1A6|SLC30A8|SLC44A5|SLC4A4|SLC6A11|SLC6A4|SLC7A11|SLC9A4|SLC9B1|SLC9C1|SLC9C2|SNX31|SOCS7|SOX2-OT|SOX9-AS1|SPATA17|SPINK5|SPTA1|ST3GAL3|STK31|SULT1B1|SV2B|SYT16|TAT|TDRD12|TENM2|TF|TMC5|TMEM132B|TNFSF11|TNN|TOP2A|TOPAZ1|TP63|TPH2|TRIM49|TRIM49B|TRIM49L2|TRIM51|TRIM64|TRIM64B|TRIML2|TRPM3|TRPM8|TSIX|TTC29|TYR|UGT2B10|USH2A|UTS2B|VWA3B|WDR64|WDR87|WDR93|WNT3|XIRP2|XKR4|XXbac-B33L19.6|XXbac-BPG55C20.7|ZAN|ZNF385D|ZNF549|ZNF658|ZNF705B|ZNF717|ZP2|ZPLD1|bP-21264C1.2</td></tr>
<tr><td>Macrophages (hC2)</td><td>175</td><td>AC074289.1|ACSL1|ADAM28|ADAP2|ADGRG6|ADORA3|ALOX15B|ALOX5|ARHGAP24|ARRB2|ARRDC2|ASAH1|ATP8B4|BMP2K|BTK|C1QA|C1QB|C1QC|C1orf162|C5AR1|CARD9|CD14|CD163|CD33|CD86|CEP170|CH507-9B2.1|CLEC4E|CLEC7A|CPM|CSF1R|CSF2RA|CSF3R|CTD-2282P23.2|CTD-2353F22.2|CTSB|CTSH|CYBB|DENND4B|DOK3|ELMO1|EMILIN2|F13A1|FCER1G|FCGR1A|FCGR2A|FCGR2B|FCGR2C|FCGR3A|FCGRT|FCHO2|FGD2|FGD4|FGL2|FGR|FMN1|FMNL2|FNIP2|FOLR2|FOLR3|FPR1|FPR3|FRMD4A|FRMD4B|GAA|GAL3ST4|GLT25D1|GPR183|GPR34|GRN|GSAP|HAVCR2|HBEGF|HCK|HCLS1|HLA-DMB|HSPA6|ICOSLG|IFI30|IL18|IRF5|ITGAX|IVNS1ABP|JDP2|KCTD12|LAIR1|LAT2|LGALS9B|LILRA6|LILRB1|LILRB2|LILRB3|LILRB4|LILRB5|LPAR5|LST1|LY86|MAF|MAFB|MARCH1|MEF2A|MIR181A1HG|MKNK1|MPEG1|MRC1|MS4A14|MS4A4A|MS4A4E|MS4A6A|MS4A7|MSR1|NCF4|NPL|OLFML2B|PDGFB|PLEKHG5|PLXDC2|PRKAG2|PRSS36|PSTPIP2|PTGS1|PTPN18|RASSF2|RASSF4|RCN3|REL|RHBDF2|RIN3|RN7SL138P|RN7SL368P|RNA5SP498|RNASE6|RNASET2|RNF144B|RNF149|RP11-12O16.1|RP11-796I2.2|RP11-848P1.3|RP6-159A1.4|SAT1|SCIMP|SH2B3|SIGLEC1|SIGLEC10|SIGLEC8|SIGLEC9|SLC11A1|SLC1A3|SLC43A2|SLCO2B1|SMAP2|SPI1|SRGAP2|SRGAP2B|SRGAP2C|STAB1|SYK|TBC1D12|TFEC|TGFBI|THEMIS2|TLR1|TLR2|TM6SF1|TMEM106A|TMIGD3|TNFAIP2|TREM2|TREML1|TRPM2|TYMP|TYROBP|VSIG4|VSIR|WAS</td></tr>
<tr><td>Z. fasciculata (hC3)</td><td>151</td><td>ABTB2|AC012065.4|AC013394.2|AC087590.3|ADAMTS9|ADCK5|ALAS1|ALDH3A2|AP001476.2|APEX1|ATP1B3|AXL|C18orf8|C1orf54|C2orf66|CASP9|CCDC138|CDH4|CH507-42P11.5|CH507-42P11.8|CHD1|CPEB4|CTB-118N6.3|CTB-32O4.2|CTB-50E14.6|CTD-2311M21.3|CTH|CUL4A|CXorf40A|CXorf40B|CYP11B1|DAZAP1|DLG5|DNAJC12|EEF2K|ELL2|ELOVL5|EPOR|ESYT2|FLNB|FN3K|FNDC4|FOXK2|FZR1|GALC|GALNT2|GCKR|GLUD1|GNL3|GRAMD1B|GRAMD4|GTPBP4|GUK1|HEATR1|HOXA5|HSCB|HSPE1|IDI1|JAZF1|LARP4|LARP4B|LATS2|LDLR|LINC00313|LONRF2|MC2R|MEG8|MIDN|MIR202|MIR22HG|MIR370|MTCH1|MTSS1|NDEL1|NEK1|NGLY1|NMB|NPC1|PABPC4|PAPSS2|PC|PGAP1|PHLDB2|PIGA|PMEPA1|PPA1|PRMT9|PRR26|PTPN14|QRICH1|RAB20|RANBP2|RGPD1|RGPD2|RGPD3|RGPD4|RGPD5|RGPD6|RGPD8|RGS2|RP1-266L20.2|RP11-271K21.7|RP11-385F5.4|RP11-388C12.8|RP11-430C17.1|RP11-44N11.1|RP11-456N14.2|RP13-49I15.5|RP3-445N2.1|RTL1|SCAP|SEMA6A|SH3BP5|SIK1|SLC16A9|SLC19A2|SLC25A28|SLC26A2|SMYD4|SNORD113-2|SNORD113-3|SNORD113-7|SNORD114-13|SNORD114-19|SNORD114-2|SNORD114-28|SNORD114-29|SNORD114-3|SNORD114-4|SNORD114-6|SOGA1|STAR|THAP2|TIPARP|TLE1|TOR3A|USP36|USP37|UVRAG|VEGFA|WDR55|YBX3|ZBED1|ZC3H12A|ZDBF2|ZFAND5|ZNF141|ZNF331|ZNF503|ZNF536|ZNF571</td></tr>
</table>

<table>
<tr><td>Chromaffin (hC4)</td><td>416</td><td>ABCC8|AC006019.4|AC006042.6|AC008268.3|AC010082.2|AC012309.5|AC016716.2|AC066606.1|AC087886.1|AC138969.4|AC159540.1|ACE|ACOT11|ACSL6|ACVR1C|ADAMTS19|ADCYAP1R1|ADGRA1|ADGRB3|ADM|AHI1|AKAP6|AKR1C1|AKR1C2|ALCAM|AMPD2|ANK2|ANKRD20A1|ANKRD20A2|ANKRD20A4|AP3B2|APC2|ARC|ARFGEF3|ARHGAP33|ARHGAP36|ARHGEF4|ASIC3|ATCAY|ATP1A3|ATP1B1|BAIAP2|BAIAP3|BEX4|BICD1|BMP8B|BRINP1|BSCL2|BSN|C10orf113|C1orf226|CACNA1B|CACNA1D|CACNA1E|CACNA2D3|CADM1|CADM2|CADPS|CALY|CAMK2B|CARTPT|CCDC148|CCDC154|CCP110|CCSER1|CD24|CDH18|CELF3|CELF4|CELSR3|CEP126|CERCAM|CFAP46|CHD5|CHGA|CHGB|CHRFAM7A|CHRNA3|CHRNA7|CLASP2|CNKSR2|CNTN1|CNTN3|COL22A1|CRMP1|CXorf57|CYB561|DBH|DDC|DGKB|DLX1|DNALI1|DPP6|DPY19L2|DPYSL3|DRD2|DRGX|DUSP26|DYNC1I1|DYNC2H1|DZIP1|DZIP3|EFHC1|EFS|EGFLAM|ELAVL4|EML5|ENO2|EPHA8|ERICH3|FAIM2|FAM133A|FAM162B|FAM171B|FAM221A|FBXW7|FHOD3|FLRT1|FLRT3|FMN2|FRRS1L|FSTL5|FUT8|GALR1|GCGR|GCH1|GDAP1|GJD2|GNB3|GPR176|GPRASP1|GPRASP2|GRIA2|GRIA4|GRIK1|GRIK2|GSE1|HAGHL|HAND1|HAND2|HAND2-AS1|HDAC9|HMP19|HSF4|HTATSF1|ICA1|INA|INPP5F|INS|IQCA1|JAKMIP3|JPH4|KCNH2|KCNH5|KCNH8|KCNK9|KCNQ2|KHDRBS3|KIF1A|KIF3A|KIF5A|KIFAP3|KNDC1|KRT19|KSR2|L1CAM|LINC00632|LRRC4B|LRRN2|LRRTM3|MAGED4|MAGED4B|MAGI2|MAP1A|MAP2|MAP2K5|MAP7|MAPK8IP2|MAPRE3|MAPT|MARK1|MAST1|MDGA1|MDH1B|MGAT4C|MIAT|MIR137HG|MIR7-3HG|MON2|MPP2|MTCL1|MUM1|MYEF2|MYT1|MYT1L|NACAD|NALCN|NBEA|NCAM1|NCAM2|NCKAP5|NDRG4|NEBL|NEFM|NELL2|NOS1AP|NPHP1|NPIPA1|NPIPA2|NPIPA3|NPIPA5|NPIPA7|NPIPA8|NRIP3|NRSN1|NT5DC2|NTNG1|OCRL|OGDHL|OSBPL6|PAK3|PAMR1|PCBP4|PCLO|PCSK1N|PCSK2|PENK|PER3|PHC1|PHOX2A|PHOX2B|PIANP|PKD1|PKDCC|PKIA|PLCB4|PLEKHA6|PM20D1|PNMT|PPFIA2|PPP1R1B|PRCD|PREPL|PRKAR1B|PRKCE|PRPH|PRR36|PRUNE2|PSD|PTPRN|PWRN1|RAB3C|RAMP1|RAP1GAP2|REPS1|RET|RFX3|RFX6|RGS11|RGS4|RGS5|RIC3|RIMBP2|RIMS1|RMST|RN7SL508P|RNF112|ROBO1|RP11-117L5.4|RP11-1186N24.5|RP11-1191J2.2|RP11-1212A22.4|RP11-121G22.3|RP11-134G8.7|RP11-145P16.2|RP11-148L24.1|RP11-177H2.1|RP11-192H23.8|RP11-216L13.19|RP11-259P6.2|RP11-275E10.1|RP11-346C4.2|RP11-379F12.7|RP11-416N2.4|RP11-457P14.4|RP11-479O17.12|RP11-481F24.3|RP11-490G2.2|RP11-54O7.3|RP11-567N4.2|RP11-627G23.1|RP11-650L12.2|RP11-666A20.4|RP11-701H24.3|RP11-710C12.1|RP11-728K20.2|RP11-734K23.9|RP11-74E22.8|RP11-761E20.1|RP11-763F8.1|RP11-798M19.6|RP13-20L14.1|RP13-487P22.1|RP3-388E23.2|RP3-403A15.5|RP3-525N10.2|RP4-795A5.1|RSPH1|RTN1|RUNDC3A|RUSC1|SAMD11|SAMD5|SCG2|SCG3|SCG5|SCN2A|SCN3B|SCN9A|SCRT1|SEC31B|SESN3|SEZ6L|SEZ6L2|SGIP1|SGK494|SGPL1|SLC18A1|SLC18A2|SLC22A17|SLC24A2|SLC25A12|SLC25A27|SLC25A36|SLC29A4|SLC31A1|SLC37A1|SLC38A11|SLC4A8|SLC6A17|SLC6A2|SLC8A2|SMPD4|SNAP25|SNAP91|SNHG14|SNORD116-14|SNORD116-15|SNORD116-17|SNORD116-19|SNORD116-29|SNRNP40|SPATA46|SPEG|SPOCK3|SRCIN1|SRRM3|SRRM4|SST|ST8SIA3|STMN2|STX1A|STXBP1|SULT4A1|SUSD4|SYT1|SYT11|SYT4|SYT5|TACC2|TAGLN3|TBX20|TEAD1|TENM1|TFAP2B|TH|TIAM1|TLX2|TMEM130|TMEM179|TMEM59L|TMOD1|TMX4|TNKS|TPPP|TPPP3|TRIM9|TRO|TTC21A|TTC8|TTLL7|TUB|UCHL1|UNC13A|UNC5A|UNC5B|UNC5C|UNC79|UNC80|VAT1L|VSTM2A|VSTM2L|VWA5A|VWA5B2|VWDE|WASF1|WDR17|YPEL4|ZC3H14|ZDHHC11|ZDHHC11B|ZFR2|ZMAT1|ZNF334|ZNF483|ZNF704</td></tr>
<tr><td>Z. glomerulosa (hC5)</td><td>18</td><td>AC078941.1|AC097662.2|ARRDC3|BBS2|CALN1|DACH1|ETV1|HDAC5|IRS1|MCOLN3|NTM|PHACTR2|RIMS2|RP11-1113L8.1|SACS|UGCG|VCAN|VSNL1</td></tr>
<tr><td>Endothelial (hC6)</td><td>105</td><td>AC007952.5|ADAM15|ADCY4|ADGRL4|APOLD1|ARHGAP29|ARHGEF15|BTNL9|CALCRL|CASKIN2|CCM2L|CD34|CDH5|CEACAM1|CLDN5|CLEC14A|COL8A1|CPLX1|CRHBP|CX3CL1|CXorf36|DCHS1|DHRS3|DNASE1L3|DOCK6|DYSF|EFNA1|EFNB2|EGFL7|ELK3|EMCN|EPHB4|ESAM|EXOC3L2|F8|FAM167B|FCN2|FLT1|FLT4|FZD4|GAS6|GJA4|GPM6A|GSN|HECW2|HSPG2|HYAL2|ID1|IFI27|IGFBP4|IGFBP5|IRX3|ITPRIP|KCNIP4|KDR|LDB2|LIMS2|LMO2|MIR7978|MMRN2|MYCT1|NOS3|NOSTRIN|NOTCH4|NOVA2|NPDC1|PCDH12|PCDH17|PECAM1|PLAT|PLK2|PLPP1|PLPP3|PLVAP|PODXL|PRSS23|PTPRB|RAMP2|RAMP3|RARB|RASIP1|ROBO4|RP11-286H15.1|S1PR1|SCARF1|SDPR|SEC14L1|SEPN1|SERPINB6|SH2D3C|SHANK3|SHE|SLC9A3R2|SLCO2A1|SOX18|TAL1|TEK|TFF3|TGM2|TIE1|TM4SF1|TM4SF18|TMEM233|TMEM88|VWA1</td></tr>
<tr><td>Mesenchymal (hC7)</td><td>121</td><td>ABCA6|ABCA9|ADAMTSL2|ADH1A|ADH1B|ADH1C|AEBP1|AMOTL2|AP000892.6|ARHGAP10|ATF3|BGN|C10orf10|C11orf96|C1R|C1S|C7|CALD1|CCL2|CFH|CFHR1|CNN1|COL12A1|COL16A1|COL1A1|COL1A2|COL27A1|COL3A1|COL6A1|COL6A2|COL6A3|COLEC11|CRISPLD2|CSRP1|CTD-2033D15.2|CTD-2033D15.3|CTGF|CXCL12|CYP1B1|CYR61|DCN|DSE|ECM2|EFEMP1|EFEMP2|EMILIN1|EPS8|ERRFI1|F3|FAM153A|FAM153C|FAM180A|FAM65C|FBLIM1|FBLN1|FBLN5|FERMT2|FMO2|GGT5|GPC3|GPX3|HGF|IGFBP6|LAMA2|LHFP|LTBP4|LUM|MAMDC2|MATN2|MFGE8|MGP|MIR143HG|MIR145|MMP19|MMP2|MXRA8|MYADM|MYLK|NEURL1B|NFASC|NGFR|NID1|NNMT|NPY1R|NR2F1|PALLD|PDGFA|PDGFRA|PDGFRB|PDLIM5|PEAR1|PLA2R1|PLCXD3|PLEKHG2|PRELP|PRPS2|PRR16|PTH1R|RBMS3|RBPMS|RN7SL625P|RP11-1101H11.1|RP11-327J17.3|RP11-517I3.2|RP11-613D13.8|RP11-635L1.2|RP11-826N14.7|RP11-844P9.1|RXFP1|S1PR3|SCN7A|SERPINE1|SETBP1|SH3D19|STEAP4|TAGLN|TCF21|THBS1|TIMP1|VCL|WISP1</td></tr>
<tr><td>Z. reticularis (hC8)</td><td>39</td><td>ABCC3|ABO|AC009410.1|AMHR2|ANKRA2|COX19|CSDC2|CYP2W1|EPHX2|ERV3-1|FBN2|GSTA4|HDHD3|HELQ|KCNN2|METTL17|MYO7A|MYZAP|NEIL1|NR0B2|NR1H4|PALMD|PLEKHA5|PRLR|RP11-12L8.1|RP11-290F5.2|RP11-334C17.5|RP11-713M15.1|RP11-806O11.1|RP11-818F20.5|SNAP47|SNORD14A|STRADB|TKFC|TPCN1|UBAP2|WISP2|ZMYM3|ZNF608</td></tr>
<tr><td>Cortex (hC9)</td><td>18</td><td>C10orf11|KIAA1024|MT1B|NUDT4|RF00100|RN7SKP185|RN7SKP203|RN7SKP230|RN7SKP255|RN7SKP291|RN7SKP36|RN7SKP37|RN7SKP48|RN7SKP71|RN7SKP80|RN7SKP9|RP11-399K21.12|TFPI2</td></tr>
<tr><td>T-cells (hC10)</td><td>33</td><td>AC092580.4|CCL5|CCND3|CD2|CD3D|CD3E|CD6|CD8A|CXCR4|EMB|GPR174|GZMH|HCST|HLA-F|IKZF3|IL32|ITGAL|KIAA1551|LCK|PARP8|PTPRC|PYHIN1|RP11-264B14.2|RP11-353N4.6|RP11-3P17.5|RP11-473M20.7|RP11-622O11.2|TBC1D10C|TMC8|TRAC|TRBC1|TRBC2|TRG-AS1</td></tr>
</table>

**Specific gene signature in mouse post-natal AG**

<table>
<tr><th>Mouse adrenal gland cluster</th><th>Number of genes in the specific signature</th><th>Specific gene signature</th></tr>
<tr><td>Z. glomerulosa (mC1)</td><td>14</td><td>Cdk18|Cpb1|Eno3|Galnt18|Htr3a|Lepr|Lrrc55|Neurl1b|Nkd1|Nxn|Pde2a|Qsox1|RP23-89M14.3|Vsnl1</td></tr>
<tr><td>Endothelial (mC2)</td><td>15</td><td>Adgrl4|Car8|Cyp4b1|F8|Fam117b|Gpr182|Ipo11|Jam2|Kcnb1|Lrg1|Ly6i|Oit3|Ptprb|RP24-374N16.7|Rapgef5</td></tr>
<tr><td>Macrophages (mC3)</td><td>128</td><td>AC164004.2|Abcc3|Acp5|Adam17|Adgre1|Aoah|Apobec1|Apoe|Arhgap19|Blvrb|C1qa|C1qb|C1qc|C3ar1|C5ar1|C5ar2|Camk1|Ccl3|Ccl4|Cd300d|Cd300lb|Cd300ld|Cd300lh|Cd84|Cebpa|Ch25h|Clec4a3|Clec4d|Clec4n|Clec7a|Cln8|Cmklr1|Cndp2|Col14a1|Creb5|Csf1r|Csf3r|Ctsa|Ctsb|Ctss|Ctsz|Cx3cr1|Cxcl2|Cxcl3|Dok3|F13a1|Fcgr1|Fcgr3|Fcgr4|Fmn1|Frmd4b|Ftl1|Fyb|Gabbr2|Gm22816|Gpr31b|H2-M2|Hacd4|Hexb|Hpgds|Ifi207|Itgb5|Klf6|Klra2|Lair1|Lgmn|Lilr4b|Lilra5|Lilrb4a|Lpcat2|Lst1|Lyz1|Lyz2|Maf|Mafb|Mertk|Mmp13|Mrc1|Ms4a14|Ms4a4a|Ms4a6b|Ms4a6d|Ms4a7|Msr1|Npl|Ntpcr|Olfr110|Olfr111|P2ry13|Pf4|Pla2g15|Pla2g7|Plau|Pparg|RP23-131O4.4|RP23-331L12.5|RP23-331L12.6|RP23-331L12.7|RP23-44G15.5|RP24-167A12.1|RP24-92F8.3|Rab20|Rab3il1|Rin2|Sat1|Sdc4|Sdcbp|Siglec1|Slc11a1|Slc13a3|Stab1|Tgfbr1|Tlr1|Tlr6|Tlr7|Tlr8|Tmem106a|Tmem132a|Tnf|Tnfrsf11a|Tpd52|Trem2|Trpv4|Vcam1|Vsir|Zeb2os|Zfp691|Zmynd15</td></tr>
</table>

<table>
<tr><td>Endothelial (mC4)</td><td>99</td><td>Apbb2|Aplnr|Arhgef15|Bcl6b|Btnl9|Bvht|Car7|Cdc42ep1|Chrm3|Cldn5|Clic5|Cmtm8|Dkk2|Dll4|Dock6|Dok4|Dram1|Dusp3|Dysf|Ecscr|Efna1|Efnb2|Egfl7|Enpep|Eogt|Esm1|Fam65a|Fam71a|Flt1|Flt4|Fscn1|Gipc3|Gm22918|Gm23616|Gm25255|Gm25287|Gm25871|Gm26067|Gm27438|Gm27514|Gng11|Gpihbp1|Hbegf|Hdac7|Hlx|Hspa1a|Hspa1b|Hspg2|Igfbp5|Jak3|Kank3|Kdr|Kitl|Lmo2|Lrrc3b|Mapk11|Meis2|Meox1|Mir680-2|Mpzl1|N4bp3|Nos3|Npr3|Olfr1396|Palm|Pbx1|Plat|Plk2|Plpp3|Plvap|Plxnd1|Pola2|RP23-162G11.1|RP23-192A6.2|RP23-410L16.5|RP24-171J16.5|RP24-293E17.2|RP24-337D17.1|RP24-425B15.4|Rapgef3|Rasip1|Rsad2|Scarf1|Sema3g|Sema7a|Shank3|Sigirr|Slc9a3r2|Sox17|Stap2|Stc1|Tbxa2r|Tcf4|Tmem204|Tmem88|Tnfaip1|Ushbp1|Vamp5|Zfp69</td></tr>
<tr><td>Cortex (mC5)</td><td>2</td><td>Pla2g5|Rab4a</td></tr>
<tr><td>Mesenchymal (mC6)</td><td>62</td><td>Abcc9|Adamtsl2|Ahsg|Angpt1|Art4|Bgn|Bicc1|Bmp4|Bmp6|C6|C7|Ccdc68|Ccl19|Cfh|Cfhr1|Cfhr2|Cldn11|Colec11|Crispld2|Cxcl12|Cygb|Cyp1b1|Dkk3|Dpep1|Dpyd|Elmod1|Errfi1|Fam205c|Fbln7|Fbn1|Fhl1|Gdf10|Gja4|Gm2564|Gpx3|Grem1|H6pd|Hsd11b1|Lox|Loxl1|Lum|Mfge8|Ngf|Osmr|Pdgfrb|Pir|Prss23|Ptges|RP23-180A8.3|RP23-304I1.4|RP23-388P16.8|RP23-411B7.2|RP24-145O12.1|Sdc1|Selenbp1|Selenbp2|Sntg1|Steap4|Svep1|Tmeff2|Wisp1|Wnt5a</td></tr>
<tr><td>Cortex (mC7)</td><td>25</td><td>Alas1|Cacna2d1|Chchd10|Cpe|Ctsf|Cttnbp2|Cyb5b|Cyp21a1|Efhd1|Fgfr2|Grhpr|Kcnk3|Klhdc8a|Nr5a1|Oxr1|Pam|Prdx3|Rerg|Slc27a3|Soat1|Sorbs2|Tbx3os1|Tfdp2|Vegfa|Wnt4</td></tr>
<tr><td>Macrophages (mC8)</td><td>44</td><td>Alox5|Ank|Bach1|Batf|Bhlhe40|Ccl9|Ccr1|Ccr3|Ccrl2|Cd163|Cd209a|Cd209c|Cd33|Cfp|Clec10a|Clec4b1|Emb|Emilin2|Fgr|Gadd45b|Gpr141|Gpx1|Ifitm6|Il1rl1|Itgam|Krt80|Ltb4r1|Mcemp1|Mcl1|Mcub|Mgl2|Nfkbia|Nfkbie|Nfkbiz|Ptgs2|Ptpro|Pvr|RP23-101F14.3|S100a4|Socs3|Tnfaip3|Tnfsf9|Tnip3|Trem1</td></tr>
<tr><td>Cortex (mC9)</td><td>17</td><td>Abhd2|Akr1c18|Cyp11b1|Dhx58|Fads2|Gstm7|Idh1|Me1|Mrpl33|Prss35|Pyroxd2|RP23-216C6.8|RP23-388H13.4|Rap1gap|Sdsl|Srd5a2|Tmem205</td></tr>
<tr><td>Glial (mC10)</td><td>208</td><td>AC110496.3|Aatk|Abca8b|Adam11|Adam23|Adgrg6|Agrn|Akap12|Aldoc|Anks1b|Apod|Arhgef10|Arhgef16|Arhgef26|Art3|Aspa|Bche|Cadm2|Cadm4|Car12|Car5b|Catsperz|Cd59a|Cd59b|Cdh10|Cdh19|Cdh2|Celsr2|Chl1|Chst5|Cmtm5|Cntf|Cobl|Col11a1|Col18a1|Col20a1|Col27a1|Col28a1|Col5a2|Col9a2|Crtac1|Ctnnal1|Cyp2j12|Cyp2j13|Cyp2j6|Cyp2j7|Cyp2j8|Cyp2j9|Dagla|Dbi|Dhh|Dst|Dtna|Edil3|Egfl8|Elovl2|Enpp1|Entpd2|Ephb3|Erbb3|Fabp5|Fabp7|Fam19a5|Farp2|Fbln2|Fbln5|Fbxo7|Fez1|Flrt1|Foxd3|Fxyd3|Gal3st1|Galnt16|Gas7|Gfap|Gfra1|Gfra2|Gfra3|Gjc3|Gpc1|Gpm6b|Gpr37l1|Gpsm2|Gpt2|Grb14|Grik3|Gulp1|Hapln1|Hepacam|Hey1|Hey2|Igsf11|Il1rap|Insc|Itga2|Itm2a|Itpk1|Kcna1|Kcna2|Kcna6|Kcnj10|Kcnk5|Kif21a|L1cam|Lgi4|Lhfpl2|Lpar1|Lrrc4c|Lrrtm1|Luzp2|Map7d2|Matn4|Mboat2|Mcam|Megf10|Megf6|Megf9|Mfap3l|Mfsd2a|Mmp15|Mmp17|Mocos|Ncam1|Ncan|Ncdn|Ngfr|Nid1|Nkain2|Nkain4|Nlgn3|Nrn1|Olfml2a|Pcbp4|Phgdh|Phldb1|Plce1|Plekhb1|Plp1|Plxdc2|Plxnb3|Pmepa1|Ppp1r1b|Prkca|Prnp|Ptgfrn|Ptprz1|Qk|RP23-199B2.4|RP23-28O11.1|RP23-35D18.1|RP23-95H5.1|RP24-147J7.1|RP24-209I7.2|RP24-222G3.1|RP24-222G3.2|RP24-278L20.2|Rasgef1c|Reep6|Rhbdf1|Rhobtb3|Rtkn|Rxrg|S100b|S1pr3|Scd2|Schip1|Scn7a|Scrg1|Sema3b|Sema3c|Sema3e|Serpine2|Sfrp1|Shc4|Shroom2|Slc35f1|Slc44a1|Slitrk6|Snca|Sntb1|Sorbs1|Sorcs1|Sort1|Sostdc1|Sox10|Sox2|Sox8|Sparc|Stard13|Sv2c|Syt11|Tenm3|Tgfb2|Thbs2|Tmem254a|Tmem254b|Tmem254c|Tmod2|Tmprss5|Trim2|Ttyh1|Tubb4a|Ugt1a6a|Ugt1a6b|Vcan|Vwa1|Zdhhc2|Zfp536</td></tr>
<tr><td>Chromaffin nAChR $\alpha7^{low}$ (mC11)</td><td>13</td><td>Atp1b1|Atxn10|Cfap61|Cplx2|Cyb561|Map1b|Pcp4|RP23-182N12.4|RP23-410K18.1|Rab3b|Slc31a1|Slc6a4|Uchl1</td></tr>
<tr><td>Ambiguous (mC12)</td><td>4</td><td>Gm10717|Gm11168|Gm21738|RP23-416G20.2</td></tr>
<tr><td>Capsule (mC13)</td><td>55</td><td>Adamtsl1|Adgrg2|Adra1b|Ano1|Bdh2|Bhmt|Col4a3|Col4a4|Cpxm2|Csrp1|Dock9|Fam20c|Fmod|Foxp2|Gas1|Gas6|Gm5096|Gpc3|Grip2|Igfbp6|Inhba|Islr|Klf5|Lamb2|Lypd1|Myl9|Nbl1|Npnt|Nr2f2|Ntn1|Ntng1|Olfr1033|Omd|Pcdh9|Pdk1|Pdpn|Penk|Perp|Pkdcc|Ptch1|Ptrf|RP23-130L21.1|Rspo3|Selm|Slc7a2|Smo|Smoc2|Stxbp6|Sulf2|Tjp1|Tnfrsf11b|Twsg1|Vldlr|Wfdc1|Wnt10a</td></tr>
<tr><td>Macrophages (mC14)</td><td>148</td><td>2310031A07Rik|AC092855.1|Adam19|Adam8|Adrb2|Ahr|Alox5ap|Amica1|Anpep|Apobec3|Arhgap30|Arid4a|Arpc2|Batf3|Bloc1s2|Cd180|Cd24a|Cd36|Cd48|Cd52|Cdk14|Clec12a|Clec9a|Coro1a|Cst3|Cyp8b1|Dnase1l3|Dock5|Dpp4|Efr3a|Esyt3|Evi2a|Ffar4|Fgd2|Fgl2|Flt3|Fmnl2|Fnbp1|Fndc7|Gcsam|Gm6133|Gna15|Gng10|Gpr33|Gsap|H2-Ab1|H2-DMb1|Hcls1|Hmgb2|Hps4|Ifi205|Impa2|Irf5|Irf8|Itgae|Itgb7|Itpripl1|Kctd12|Kit|Laptm5|Lcp1|Lrrc18|Lsp1|Ly75|Man2b1|Mctp1|Met|Mnda|Mndal|Mpeg1|Mycl|Myo1f|Myo9a|Naaa|Naga|Nckap1l|Ncoa7|Nek6|Olfr54|Parp8|Parvg|Pde1b|Phf11a|Phf11b|Phf11c|Phf11d|Pi4k2a|Pkib|Plbd1|Plekho2|Ppm1m|Ppp1r11|Ppt1|Pqlc1|Proser2|Prr5l|Psap|Psmb8|Psmb9|Ptpn18|Qpct|RP23-182K18.1|RP23-250E1.9|RP23-381B19.2|RP23-423B21.7|RP23-458B6.22|RP23-46L5.1|RP23-4F16.11|RP23-76L21.3|RP23-83N10.1|RP24-104I20.2|RP24-158L15.1|Rab11a|Rab32|Rab43|Rab7b|Rnase6|Rnf157|Rnf19b|Sept3|Sept6|Sh3bp1|Shtn1|Slc36a3os|Slc8b1|Snx22|Strip2|Sult1a1|Tap1|Tap2|Tbc1d8|Tcf19|Tlr11|Tlr3|Tmsb10|Tmsb4x|Tnni2|Treml4|Trpm2|Trpv2|Ttc39a|Ubash3b|Ucp2|Unc119|Unc119b|Wdfy4|Xcr1|Zdhhc23</td></tr>
<tr><td>Chromaffin nAChR $\alpha7^{high}$ (mC15)</td><td>266</td><td>AC122733.3|AC157555.1|AC157667.1|Abca7|Ace|Ache|Adcy1|Adcyap1r1|Adrbk2|Agtpbp1|Ak5|Aldh1a3|Amer2|Ap3b2|Apbb1|Arfgef3|Arhgef28|Arnt2|Asic4|Astn1|Atp2b2|Atp6v0d1|B3galnt1|B3galt2|Braf|Btbd8|C1ql1|Cacna1a|Cacna1b|Cacna1d|Cadps|Camk2b|Casz1|Cd9|Cdh12|Cdh22|Cdon|Celf3|Celf4|Chga|Chgb|Chic1|Chrm1|Chrna3|Chrna7|Chrnb4|Cidea|Ckmt1|Cnksr2|Cntnap5a|Coro2b|Cox8b|Cpeb4|Cplx1|Crmp1|Cxxc4|Cxxc5|Dbh|Dgkb|Dgkg|Dgkz|Diablo|Disp2|Dixdc1|Dlk1|Dmxl2|Dnm1|Dopey2|Dpysl3|Dqx1|Dusp26|Elavl4|Eml5|Eml6|Enah|Epha5|Erc1|Erc2|Exph5|Eya1|Fam155a|Fam163a|Fam171b|Fam19a1|Fndc3a|Fstl5|Gabrb3|Gal|Garnl3|Gata3|Gcgr|Gdap1l1|Ggt7|Gnao1|Gnas|Gper1|Gpr155|Gpr19|Gpr22|Gprasp1|Gprasp2|Grb10|Gria2|Grm7|Hid1|Hlf|Hpca|Hrh3|Hspa2|Hspb8|Il20ra|Jph3|Kcnc4|Kcnn3|Kcnq2|Kcns2|Kif5c|Klhl32|Lgi1|Lgr5|Lix1|Lrrc3|Lsamp|Ltk|Mab21l2|Mapk10|Mapt|March8|Mbd5|Megf11|Mfsd6|Mmd|Mtcl1|Myl1|Myt1|Myt1l|Nap1l5|Ncald|Ndufa4l2|Necab2|Nefl|Neurl1a|Neurod4|Nfasc|Nlrp6|Nnat|Npy2r|Nrg3|Nsg1|Nsg2|Nt5dc2|Nxph4|Nyap2|Osbpl2|Pacsin1|Parp6|Pcbp3|Pcdh15|Pcgf1|Pclo|Phactr1|Phactr3|Phox2a|Phox2b|Phyhipl|Pirt|Plekha6|Plppr2|Pnmal1|Ppfia2|Ppp2r2c|Prrt1|Prune2|Ptger3|Ptprf|Ptprn|Ptprr|RP23-121F17.2|RP23-121N17.2|RP23-122E16.2|RP23-122E16.3|RP23-146D13.2|RP23-14P23.2|RP23-14P23.9|RP23-177J21.10|RP23-263O11.1|RP23-316B4.2|RP23-363K7.4|RP23-469N16.1|RP23-52J10.3|RP23-52J10.5|RP23-52J10.6|RP23-54G8.1|RP23-77H12.1|RP24-189K3.2|RP24-274B19.1|RP24-68E20.11|Rab27a|Rab40b|Ralgds|Rap1gap2|Rgs9|Ripply2|Rnf165|Rnf217|Robo2|Rpap1|Rps6kl1|Ryr2|Scg2|Scn3a|Scn3b|Scn9a|Sgip1|Sgpp2|Sgsm1|Shank2|Sidt1|Slc10a4|Slc18a1|Slc18a2|Slc26a4|Slc35d3|Slc7a14|Slco3a1|Slit3|Smpd3|Snap25|Snap91|Socs2|Soga3|Srpk2|Srrm3|Ssbp2|Sspo|St18|St8sia3|Stac2|Stk32a|Stxbp1|Syn2|Syt1|Syt17|Syt4|Sytl4|Tacc2|Tbx20|Tle1|Tlx2|Tmem64|Tppp3|Tspyl4|Tuba4a|Tubb3|Unc5a|Unc5c|Unc79|Unc80|Vamp1|Ypel4|Zcchc18|Zdbf2|Zfhx2|Zfp423|Zfp467|Zswim5</td></tr>
<tr><td>Z. fasciculata (mC16)</td><td>15</td><td>Aif1l|Deptor|Gabrg3|Hs6st1|Ildr2|Itga9|Mgat4b|Pcyt1a|Ppl|RP23-256J14.2|Runx2|Slc25a30|Spred1|Stx11|mt-Te</td></tr>
<tr><td>T-cells (mC17)</td><td>32</td><td>Bcl11b|Cd247|Cd28|Cd3d|Cd3e|Cd3g|Cd40lg|Cd5|Cd6|Icos|Il4|Lat|Sh2d1a|Sit1|Tcrg-C1|Tcrg-C2|Tcrg-C3|Themis|Thy1|Trac|Traj12|Traj13|Traj18|Traj9|Trav11|Trav11d|Trbc2|Trbv12-2|Trbv13-2|Trbv13-3|Trgj1|Trgj3</td></tr>
<tr><td>T-cells (mC18)</td><td>82</td><td>Adamts14|Apol7a|Apol7b|Apol7c|Apol7e|Ccl5|Cd160|Cd200r2|Cd200r4|Cd7|Cracr2a|Ctsw|Fam189b|Fasl|Gzma|Gzmb|Gzmc|H2-K1|H2-Q7|Hcst|Igsf5|Il2rb|Klra10|Klra3|Klra5|Klra6|Klra8|Klra9|Klrb1a|Klrb1c|Klrb1f|Klrc1|Klrc2|Klrc3|Klrd1|Klre1|Klri2|Klrk1|Mctp2|Mxd1|Ncr1|Nkg7|Padi2|Pim2|Ppp3cc|Prf1|RP23-152I6.3|RP23-20K24.9|RP23-41L21.4|RP23-52N2.1|RP24-335J14.1|RP24-95J4.3|Samd3|Serpina3f|Serpina3g|Serpina3i|Serpinb6b|Serpinb9|Spry2|Stat4|Styk1|Sytl3|Tbx21|Tcrg-C4|Tigit|Tmem106b|Tnfrsf9|Trbd1|Trbj1-1|Trbj1-2|Trbj1-3|Trbj1-4|Trbj1-5|Trbj1-6|Trbj1-7|Trdv2-1|Trdv2-2|Trdv4|Txk|Ugcg|Vgll4|Xcl1</td></tr>
<tr><td>B-cells (mC19)</td><td>27</td><td>Bank1|Blk|Ccr7|Cd19|Cd22|Cd55|Cd55b|Cd79a|Cd79b|Cxcr5|Fcer2a|Fcmr|Foxp1|Ighd|Ighj4|Ighm|Igkc|Igkj5|Iglc2|Iglc3|Iglj3|Ms4a1|Pax5|Pgap1|Pou2af1|Sell|Spib</td></tr>
</table>

**Specific gene signature in neuroblastoma**

| Neuroblastoma cluster | Number of genes in the specific signature | Specific gene signature |
|---|---|---|
| MSC (nC1) | 162 | AC002064.4\|AC006076.1\|AC008154.4\|AC010090.1\|AC068491.1\|AC079807.2\|AC107057.2\|AC116035.2\|AC132807.1\|ADAM33\|AEBP1\|ALDH1A3\|AP000560.3\|ARL2BP\|AXL\|C11orf96\|C18orf21\|C1R\|C1S\|CALD1\|CD248\|CFHR1\|COL16A1\|COL1A2\|COL27A1\|COL5A1\|COL6A2\|COL6A3\|CRISPLD2\|CRYBA2\|CTB-187M2.1\|CTC-336P14.1\|CTC-340D7.1\|CTD-2033D15.2\|CTD-2033D15.3\|CTD-2526M8.3\|CYTOR\|ERRFI1\|F3\|FAM46A\|FGFR1\|FN1\|FSTL3\|GEM\|GFPT2\|GS1-72M22.1\|HOXD13\|ID4\|INHBA\|LIF\|LIMA1\|LMCD1\|MAGEA12\|MALAT1\|MAZ\|MICALL2\|MIR3685\|MIR4282\|MRC2\|MT1E\|MYL9\|NFATC4\|NOTCH3\|NR4A1\|OR6C74\|PALM2\|PDGFRB\|PKDCC\|PLEKHA4\|PRRX1\|PTGIR\|RAB34\|RAD51AP2\|RELL2\|RIMS4\|RMRPP5\|RN7SKP5\|RN7SL140P\|RN7SL35P\|RNA5SP210\|RNA5SP290\|RNA5SP443\|RNU5E-7P\|RNU6-1146P\|RNU6-444P\|RNU6-448P\|RNU6-67P\|RP1-228P16.5\|RP1-309F20.4\|RP1-79C4.4\|RP11-111J6.2\|RP11-120K24.3\|RP11-122A8.4\|RP11-1275H24.2\|RP11-129B22.7\|RP11-152N13.5\|RP11-195B17.1\|RP11-234K24.6\|RP11-252M21.4\|RP11-272B17.2\|RP11-27G14.4\|RP11-290H9.5\|RP11-293A21.1\|RP11-295M18.7\|RP11-298J23.8\|RP11-304F15.7\|RP11-328C8.5\|RP11-341N2.1\|RP11-350N15.4\|RP11-35O15.1\|RP11-363J20.2\|RP11-365H22.2\|RP11-378J18.8\|RP11-402P6.15\|RP11-40E6.2\|RP11-438D14.3\|RP11-455J20.3\|RP11-499A10.3\|RP11-531F16.4\|RP11-579A4.1\|RP11-57H12.2\|RP11-628O18.1\|RP11-637C24.5\|RP11-644K8.1\|RP11-666A8.12\|RP11-667K14.3\|RP11-6B19.1\|RP11-702A23.1\|RP11-704J17.5\|RP11-727F15.11\|RP11-78C3.1\|RP11-78O9.1\|RP11-803D5.1\|RP11-861L17.4\|RP11-95H11.1\|RP3-412A9.11\|RP3-453P22.1\|RP3-495K2.3\|RP4-593H12.1\|RP4-742N3.1\|RP5-1068E13.7\|RP5-856G1.1\|RP5-897D18.1\|RP5-908M14.9\|SERPINE1\|SNAI2\|SNORD92\|SPRY2\|THBS1\|THBS2\|TMEFF1\|TMEM56-RWDD3\|TPM2\|TRAV3\|TRBV24-1\|UAP1\|VASN\|VCAN\|VPS37D\|WISP1\|XXyac-YX65C7_A.2\|ZNF703 |
| Undifferentiated (nC2) | 23 | ALOXE3\|C2orf61\|CDC45\|CH507-513H4.1\|CH507-528H12.1\|CTD-3074O7.5\|ELF3\|FCGBP\|GPA33\|MROH2A\|OTOP2\|PITPNM3\|PYGM\|RP11-165M6.1\|RP11-23E10.2\|RP11-552D8.1\|RP13-262C2.4\|SCNN1A\|SEC14L3\|SPATA31A3\|SPATA31A5\|UMODL1\|VAX1 |

| | | |
|---|---|---|
| Undifferentiated (nC3) | 1023 | AC005301.9\|AC005537.2\|AC007064.24\|AC007405.4\|AC007731.1\|AC008067.2\|AC008079.9\|AC008103.3\|AC008103.4\|AC008132.15\|AC008691.1\|AC008697.1\|AC009120.4\|AC009264.1\|AC010096.2\|AC011718.2\|AC012593.1\|AC013400.2\|AC015922.6\|AC016907.3\|AC064846.1\|AC073130.1\|AC073464.10\|AC074093.1\|AC092415.1\|AC092594.1\|AC097467.2\|AC099057.8\|AC104820.2\|AC108025.2\|AC144521.1\|ACOXL\|ACSM2A\|ACSM2B\|ACSM3\|ACSS3\|ACTR3B\|ACTR3C\|ADAM29\|ADAM32\|ADAM7\|ADAMTS12\|ADAMTS19\|ADAMTS20\|ADAMTS6\|ADAMTSL3\|ADCY10\|ADCY2\|ADGB\|ADGRF1\|ADGRF2\|ADGRL2\|ADGRV1\|AFF2\|AGBL1\|AGBL3\|AJ006998.2\|AK5\|AKAP7\|AL035610.1\|AL035610.2\|AL078471.5\|AL449209.1\|ALDH1L2\|ALG10B\|ALG14\|AMY1A\|AMY1B\|AMY1C\|AMY2A\|ANAPC1\|ANKEF1\|ANKRD20A1\|ANKRD20A2\|ANKRD20A3\|ANKRD20A4\|ANKRD30A\|ANKRD30B\|ANKRD30BL\|ANKRD31\|ANKRD45\|ANKRD7\|ANKS1B\|ANKUB1\|ANO3\|ANO4\|ANO5\|ANTXR2\|AOX1\|AP000432.3\|AP000476.1\|AP5M1\|APAF1\|APOB\|APOH\|ARHGAP24\|ARHGAP28\|ARHGAP42\|ARHGEF12\|ARHGEF28\|ARHGEF38\|ARMC10\|ARMC3\|ASAH2\|ASPA\|ASPM\|ATAD2\|ATL3\|ATP10B\|ATP13A4\|ATP8B4\|ATRNL1\|AXDND1\|B3GLCT\|B4GALNT2\|BARD1\|BBOF1\|BCO2\|BEST3\|BMS1\|BRCA1\|BRCA2\|BRIP1\|BRMS1L\|BUB1\|C10orf67\|C12orf42\|C12orf45\|C12orf48\|C14orf145\|C14orf37\|C18orf25\|C1orf112\|C1orf141\|C1orf168\|C1orf87\|C2CD3\|C5\|C6\|C6orf10\|C8A\|C8orf34\|C9orf85\|CACHD1\|CACNB4\|CADPS2\|CASC1\|CATSPERB\|CBWD5\|CC2D2A\|CC2D2B\|CCDC122\|CCDC129\|CCDC146\|CCDC15\|CCDC150\|CCDC168\|CCDC169\|CCDC178\|CCDC26\|CCDC30\|CCSER1\|CD109\|CD226\|CDC14B\|CDC27\|CDH10\|CDH13\|CDH18\|CDH7\|CDH9\|CDK14\|CDKN2B-AS1\|CDON\|CECR2\|CENPE\|CEP112\|CEP152\|CEPT1\|CFAP221\|CFAP43\|CFAP47\|CFAP52\|CFAP54\|CFAP58\|CFAP61\|CFAP69\|CFAP70\|CFTR\|CH17-132F21.5\|CH17-189H20.1\|CH17-264L24.1\|CH507-145C22.1\|CH507-154B10.1\|CH507-254M2.2\|CHEK2\|CHPT1\|CISD2\|CLDN11\|CLEC1B\|CLN5\|CLNK\|CLSPN\|CMC2\|CMYA5\|CNTN3\|CNTN4\|CNTN5\|CNTN6\|CNTNAP3\|CNTNAP5\|COG6\|COL11A1\|COL14A1\|COL19A1\|COL21A1\|COL24A1\|COL28A1\|COL4A3\|COL4A4\|COL4A5\|COL6A5\|COL6A6\|COL9A1\|COLGALT2\|CORIN\|COX10-AS1\|CPD\|CPS1\|CPSF2\|CPT2\|CR1\|CR2\|CRB1\|CRISPLD1\|CSMD1\|CSMD3\|CSRP2\|CTA-941F9.10\|CTC-228N24.3\|CTC-340A15.2\|CTC-454M9.1\|CTC-490G23.6\|CTCFL\|CTD-2144E22.9\|CTD-2230D16.1\|CTD-2311B13.5\|CTD-2536I1.1\|CTNNA3\|CUBN\|CYP2C9\|CYP3A5\|DAB1\|DAW1\|DAZ1\|DAZ2\|DAZ3\|DAZ4\|DBT\|DCBLD1\|DCDC1\|DCHS2\|DDX20\|DDX4\|DDX60\|DENND2C\|DEPDC1\|DEPDC5\|DET1\|DGKE\|DGKI\|DIAPH3\|DIO2\|DLEU1\|DLGAP1\|DLX6-AS1\|DMGDH\|DNAH10\|DNAH11\|DNAH12\|DNAH14\|DNAH3\|DNAH5\|DNAH6\|DNAH7\|DNAH8\|DNAH9\|DNAJC16\|DOCK7\|DOCK9\|DPP10\|DPY19L3\|DR1\|DSC1\|DSC2\|DSC3\|DSG2\|DSG3\|DTHD1\|DYNC1I1\|DYNC2H1\|E2F6\|EFCAB3\|EFCAB5\|EFCAB6\|EGF\|EHF\|ELK4\|ELMOD1\|ELOVL7\|ELP4\|EML1\|ENPP1\|ENPP2\|ENPP3\|ENTPD5\|EPHA6\|EPM2A\|ERBB4\|ERCC6\|ERVW-1\|ESCO2\|ESRP1\|ESRRG\|ETV1\|EXOC5\|EXOG\|EXPH5\|EYS\|F5\|FAM114A2\|FAM135A\|FAM135B\|FAM13A\|FAM153A\|FAM153B\|FAM153C\|FAM197Y5\|FAM19A2\|FAM227A\|FAM230A\|FAM230B\|FAM35A\|FANCD2\|FANCI\|FANCM\|FAT3\|FBN2\|FBXO15\|FBXO3\|FCRL5\|FGF7\|FGGY\|FLRT2\|FLT3\|FNDC1\|FOLH1\|FOXP2\|FRAS1\|FREM1\|FREM2\|FRK\|FSIP2\|FUT9\|G2E3\|GABRA1\|GABRA2\|GABRA6\|GABRB2\|GABRG2\|GABRG3\|GALK2\|GBP6\|GDA\|GDPD4\|GEMIN5\|GEN1\|GK5\|GLDC\|GLI3\|GLIDR\|GLYATL1\|GNAI3\|GOLGA6L22\|GPAM\|GRAMD1C\|GRAMD3\|GREB1L\|GRID2\|GRIN2A\|GRIN2B\|GRIP1\|GRM5\|GTPBP10\|GUCY1A2\|GUCY2C\|GULP1\|GYPA\|HAPLN1\|HCG23\|HCN1\|HEATR4\|HELLPAR\|HERC5\|HERC6\|HGF\|HHIP\|HMCN1\|HMGA2\|HSD17B4\|HSFY2\|HULC\|HYDIN\|IDE\|IGFN1\|IL1RAP\|ILDR2\|IMPG1\|INPP4B\|INTS2\|INTS6\|INTS7\|INTU\|IPCEF1\|IPO11\|IQUB\|ITGA8\|IYD\|KAT7\|KB-1183D5.13\|KCNU1\|KDM7A\|KIAA0586\|KIAA0825\|KIAA1257\|KIAA1328\|KIAA1468\|KIF13A\|KIF14\|KIF15\|KIF6\|KLHL3\|KLRD1\|KNG1\|KNTC1\|L2HGDH\|L3MBTL3\|LA16c-23H5.4\|LA16c-3G11.5\|LA16c-60D12.1\|LA16c-60D12.2\|LACE1\|LAMA1\|LAMA2\|LAMA3\|LAMB4\|LARGE\|LEPR\|LGI1\|LGR4\|LGR5\|LINC00466\|LINC00467\|LINC00470\|LINC00923\|LINC00960\|LINC01138\|LL22NC03-N64E9.1\|LMBR1\|LMO3\|LMO7\|LNPEP\|LPA\|LRMP\|LRP1B\|LRP2\|LRRC16A\|LRRC7\|LRRC9\|LRRIQ3\|LRRK1\|LRRK2\|LTBP1\|LVRN\|MACROD2\|MALRD1\|MAP2K6\|MAP3K20\|MATN3\|MBOAT1\|MCCC2\|MCM9\|MCTP2\|MDGA2\|MDN1\|MED12L\|MED17\|MEGF10\|MEP1B\|MET\|METTL15\|MGAM\|MGAM2\|MGST1\|MIA2\|MIPOL1\|MIR99AHG\|MKLN1\|MME\|MMP7\|MMS22L\|MR1\|MRE11A\|MROH2B\|MROH8\|MROH9\|MTHFD2L\|MUC12\|MUC16\|MUC17\|MUC19\|MUC3A\|MUSK\|MYB\|MYBPC1\|MYH1\|MYH13\|MYH15\|MYH2\|MYH4\|MYO16\|MYO1H\|MYO3A\|MYO3B\|MYO5B\|MYOM1\|MYPN\|NAALADL2\|NANOG\|NAV3\|NCAPG2\|NEB\|NECAB1\|NEDD1\|NEK10\|NEK5\|NF1\|NIPAL3\|NLGN1\|NLRP14\|NMNAT3\|NNT\|NOX4\|NPAS3\|NPHP1\|NRAP\|NRXN3\|NSF\|NT5DC3\|NTRK2\|NUBPL\|NUP133\|NUP155\|NUP210L\|NUP37\|NUP93\|NXPE1\|OLMALINC\|OR10K1\|OR1A1\|OR2AG1\|OR2AT4\|OR2M3\|OR2T12\|OR4D9\|OR4F17\|OR4F4\|OR4F5\|OR4N2\|OR52A1\|OR56A1\|OR56A4\|OR5AS1\|OR7C1\|OR8D1\|OSBPL10\|OSBPL1A\|OTOA\|OTOGL\|OVCH1\|PARP11\|PART1\|PATJ\|PCAT1\|PCCA\|PCDH15\|PCDHB1\|PCSK5\|PDE1A\|PDE4D\|PDE4DIP\|PDZD2\|PEX5L\|PGGT1B\|PGM5\|PGM5P4-AS1\|PGR\|PHLPP1\|PHLPP2\|PHTF2\|PIAS2\|PIEZO2\|PIGK\|PIGN\|PIK3C2G\|PIK3R4\|PIP5K1B\|PIWIL1\|PIWIL2\|PKD1L3\|PKHD1\|PKHD1L1\|PLA2G4A\|PLA2R1\|PLCE1\|PLCH1\|PLD5\|PLG\|POLQ\|POLR3B\|POLR3G\|POTEC\|POU2F1\|POU5F2\|POU6F2\|PPARGC1A\|PPAT\|PPIAL4G\|PRAMEF1\|PRAMEF13\|PRAMEF14\|PRAMEF25\|PRDM5\|PRDM7\|PRIM2\|PRKAA2\|PRKD1\|PRLR\|PRRC1\|PRTG\|PSG11\|PTCD2\|PTGFR\|PTPN13\|PTPN20\|PTPRG\|PTPRQ\|PTPRT\|PTPRZ1\|PWRN1\|PZP\|RAD51B\|RANBP17\|RASSF6\|RBBP8\|RBM44\|RBMY1A1\|RBMY1B\|RBMY1D\|RBMY1E\|RBMY1F\|RBMY1J\|RELN\|RF00012\|RF00017\|RF00026\|RF00096\|RF00322\|RFX6\|RGPD1\|RGPD2\|RGPD3\|RGPD4\|RGPD5\|RGPD6\|RGPD8\|RGS22\|RGSL1\|RMST\|RN7SL52P\|RN7SL544P\|RN7SL565P\|RN7SL722P\|RN7SL763P\|RN7SL787P\|RNASEH2B\|RNF217\|RORB\|ROS1\|RP1\|RP1-101G11.3\|RP1-14N1.2\|RP1-158P9.2\|RP11-100L22.2\|RP11-108M21.1\|RP11-111F5.8\|RP11-114G22.1\|RP11-115D19.1\|RP11-119D9.1\|RP11-136K7.2\|RP11-139E24.1\|RP11-142C4.6\|RP11-142I20.1\|RP11-143M1.2\|RP11-146D12.2\|RP11-146E13.4\|RP11-152L7.2\|RP11-154D6.1\|RP11-15E1.5\|RP11-168K9.1\|RP11-169F17.1\|RP11-175I6.5\|RP11-195E11.2\|RP11-211N8.7\|RP11-215N21.1\|RP11-217E22.2\|RP11-21F16.1\|RP11-222A11.1\|RP11-239L20.6\|RP11-244H18.1\|RP11-248G5.8\|RP11-255B23.3\|RP11-262H14.1\|RP11-262H14.3\|RP11-267C16.1\|RP11-275I4.2\|RP11-327J17.3\|RP11-341D18.3\|RP11-344E13.4\|RP11-345M22.2\|RP11-34P13.13\|RP11-366L20.2\|RP11-373D17.1\|RP11-374A4.1\|RP11-379B18.8\|RP11-38L15.2\|RP11-395L14.18\|RP11-3B12.1\|RP11-40A8.3\|RP11-416O18.1\|RP11-419I17.1\|RP11-422J15.1\|RP11-42F12.1\|RP11-430C1.1\|RP11-436D23.1\|RP11-458B24.2\|RP11-459O16.8\|RP11-461F11.2\|RP11-467L19.14\|RP11-470M1.2\|RP11-475I24.8\|RP11-478J18.1\|RP11-480C16.1\|RP11-482G13.1\|RP11-4L24.3\|RP11-4L24.4\|RP11-506B6.6\|RP11-510H23.1\|RP11-532N4.2\|RP11-536C10.7\|RP11-53O19.1\|RP11-541P9.3\|RP11-542H15.1\|RP11-550E22.3\|RP11-556O5.7\|RP11-561N12.2\|RP11-573D15.8\|RP11-575L7.4\|RP11-586K12.1\|RP11-588H23.3\|RP11-58E21.3\|RP11-597A11.2\|RP11-609N14.4\|RP11-626K17.3\|RP11-673E1.1\|RP11-67H24.3\|RP11-696N14.1\|RP11-704M14.1\|RP11-735B13.1\|RP11-747H7.3\|RP11-756H20.1\|RP11-764K9.1\|RP11-767I20.1\|RP11-76N22.2\|RP11-774D14.1\|RP11-776H12.1\|RP11-826N14.7\|RP11-82L2.1\|RP11-87H9.5\|RP11-90D4.2\|RP11-90D4.3\|RP11-96C23.13\|RP3-394A18.1\|RP4-565E6.1\|RP4-760C5.5\|RP4-809F18.1\|RP5-1101C3.1\|RP5-1198O20.4\|RP5-828K20.2\|RP5-916L7.1\|RP5-921G16.1\|RP5-947P14.1\|RPAP2\|RPGRIP1\|RPS6KA5\|RPS6KB1\|RTTN\|RXFP2\|RYR3\|SAMMSON\|SAR1B\|SATB2\|SCEL\|SCIN\|SCN11A\|SCO1\|SEC22B\|SEC23A\|SEMA3A\|SEMA3D\|SEMA3E\|SEMA5A\|SEPSECS-AS1\|SERPINB5\|SERPINB8\|SH3TC2\|SHC4\|SHQ1\|SI\|SLC12A1\|SLC12A2\|SLC13A1\|SLC15A2\|SLC1A2\|SLC26A3\|SLC26A5\|SLC26A8\|SLC2A13\|SLC35A3\|SLC35F4\|SLC35G1\|SLC38A9\|SLC44A5\|SLC4A4\|SLC5A12\|SLC5A8\|SLC9B1\|SLC9C1\|SLC9C2\|SLCO1A2\|SLCO4C1\|SLCO6A1\|SLMAP\|SMC1B\|SMC6\|SNAP25-AS1\|SNTG1\|SOCS7\|SOX2-OT\|SOX5\|SOX9-AS1\|SPAG17\|SPATA17\|SPDYA\|SPECC1\|SPEF2\|SPHKAP\|SPINK5\|SPTA1\|SPTLC3\|SRGAP2B\|SRGAP2C\|SSX2IP\|ST18\|STAB2\|STEAP4\|STIL\|STK3\|STK31\|STRIP2\|STXBP6\|STYK1\|SULT1B1\|SUMF1\|SVEP1\|SVOPL\|SYNE1\|SYTL2\|TAF4B\|TAOK1\|TARID\|TAT\|TBC1D31\|TBC1D8B\|TCEANC2\|TEX14\|TF\|THEMIS\|THRB\|THSD7B\|TMC1\|TMEM132B\|TMEM213\|TMEM232\|TMPRSS15\|TMPRSS7\|TOGARAM2\|TOR1AIP1\|TP63\|TPD52L1\|TPH1\|TPH2\|TPK1\|TPRG1\|TPTE2\|TRDMT1\|TRDN\|TRHDE\|TRIM43\|TRIM43B\|TRIM48\|TRIM49\|TRIM49B\|TRIM49D1\|TRIM49D2\|TRIM49L2\|TRIM51\|TRIM64\|TRIM64B\|TRPM3\|TRPM6\|TSHR\|TSIX\|TTC26\|TTC39B\|TTC5\|TTC6\|TTLL5\|TTN\|TYR\|TYRP1\|TYW5\|ULK4\|UNC13C\|USH2A\|USP32\|USP37\|USP54\|UTP20\|UTS2B\|VEPH1\|VGLL3\|VPS13D\|VWA8\|WDHD1\|WDR3\|WDR35\|WDR49\|WDR66\|WDR7\|WDR72\|WNT2B\|WRN\|XKR4\|XPNPEP3\|XX-C2158C12.1\|XXbac-B33L19.4\|XXbac-BPG55C20.7\|ZBBX\|ZBED3-AS1\|ZDHHC13\|ZFAND4\|ZGRF1\|ZNF169\|ZNF215\|ZNF277\|ZNF285\|ZNF385B\|ZNF426\|ZNF479\|ZNF519\|ZNF527\|ZNF546\|ZNF578\|ZNF595\|ZNF678\|ZNF679\|ZNF705B\|ZNF705D\|ZNF705G\| |

| | | |
|---|---|---|
| Endothelial (nC4) | 329 | [illegible] ANXA2\|APLNR\|APOLD1\|APP\|AQP1\|ARHGEF15\|ARL2\|ATG9B\|ATP11C\|ATP1A1\|BAG3\|BAG5\|BAZ1A\|BCAR1\|BCL3\|BCL6B\|BHLHE40\|BMP2\|BMP6\|BST2\|C10orf10\|C12orf71\|C2CD4B\|C6orf208\|CA2\|CALCRL\|CAV1\|CBLB\|CCDC69\|CCDC85B\|CD34\|CD55\|CD59\|CD93\|CD99\|CDC37\|CDC42EP2\|CDC42EP3\|CDH5\|CDK17\|CDKN1A\|CEACAM19\|CENPB\|CH507-513H4.3\|CH507-513H4.4\|CH507-513H4.5\|CH507-513H4.6\|CHD7\|CLEC14A\|CLIC4\|CMIP\|CNKSR3\|COL15A1\|COL4A1\|CPLX1\|CRIM1\|CRIP2\|CSF3\|CTD-3157E16.2\|CXorf36\|DDX21\|DIAPH1\|DLL4\|DOCK6\|DUSP5\|DUSP6\|ECE1\|ECSCR\|EDARADD\|EFHD2\|EFNA1\|EFNB1\|EGFL7\|EHD2\|EHD4\|EIF5B\|ELK3\|EML3\|EMP1\|EMP2\|ENDOD1\|ENG\|ENO1\|EPAS1\|EPHA2\|EPHB4\|ERG\|ESAM\|ETS1\|ETS2\|EVA1C\|F2R\|FAM110D\|FAM129B\|FAM198B\|FBL\|FJX1\|FKBP1A\|FKBP1C\|FLNB\|FLT1\|FLT4\|FOSL1\|FOXC1\|FOXO1\|FUT11\|FXYD5\|FZD4\|FZD6\|GABRE\|GALNT1\|GAR1\|GAS5\|GJA1\|GNAI2\|GNG11\|GPR4\|GRASP\|GRB10\|GRPEL1\|HAPLN3\|HES1\|HIPK3\|HLA-E\|HSPA5\|HSPG2\|HYAL2\|ID1\|ID3\|IER3\|IFI27\|IFITM2\|IFITM3\|IGFBP3\|IGFBP4\|IGFBP7\|IL33\|IL6\|IL6ST\|INHBB\|INSR\|ITGA5\|ITPKC\|ITPRIP\|JMJD1C\|KCNJ2\|KCTD12\|KIAA0355\|KIAA1462\|KLF2\|LAMA5\|LCN6\|LDB2\|LDLR\|LIMS2\|LMNA\|LMO2\|LRP10\|LRRC32\|LUZP1\|MALL\|MAN1A1\|MARCH3\|MAST4\|MCAM\|MESDC1\|MIDN\|MIR126\|MMRN1\|MMRN2\|MPZL2\|MRTO4\|MSN\|MTA2\|MYADM\|MYCT1\|MYH9\|NDRG1\|NECTIN2\|NEDD9\|NFIC\|NFKBIA\|NIP7\|NOS3\|NOTCH1\|NPR1\|NRGN\|PABPC1\|PABPC3\|PALMD\|PARVB\|PCDH12\|PCDH17\|PDGFB\|PDLIM1\|PDLIM4\|PECAM1\|PEX5\|PIEZO1\|PIM3\|PKIG\|PKN3\|PKP4\|PLAC4\|PLAT\|PLEC\|PLEKHG2\|PLPP3\|PLS3\|PLVAP\|PLXNA2\|PNP\|PODXL\|PPM1F\|PRAG1\|PRNP\|PTRF\|PVR\|PXN\|RAB13\|RAI14\|RALGAPA2\|RAMP2\|RAMP3\|RAPGEF3\|RASIP1\|RBPMS\|RCAN1\|RFLNB\|RHOC\|RHOJ\|RNASE1\|RND1\|RNF19B\|ROBO4\|RP11-1149O23.2\|RP11-14D22.5\|RP11-314N13.4\|RP11-360N9.2\|RPGR\|RPL36A\|RRP12\|S100A10\|S100A13\|S100A16\|S100A6\|S1PR1\|SASH1\|SBNO2\|SCARF1\|SDPR\|SELE\|SELP\|SEMA3F\|SEPN1\|SEPW1\|SERTAD1\|SHANK3\|SHB\|SHROOM4\|SKI\|SLC25A37\|SLC35E4\|SLC39A14\|SLC4A7\|SLC9A3R2\|SLCO2A1\|SLCO4A1\|SLK\|SMAD3\|SNAI1\|SNRK\|SOCS2\|SOCS3\|SOX17\|SOX18\|SOX7\|SPARCL1\|SPNS2\|SPRY1\|SSH1\|STC1\|STC2\|STOM\|SWAP70\|SYNPO\|TEAD4\|TGFB1\|TGFBR2\|TGM2\|THBD\|TIE1\|TINAGL1\|TM4SF1\|TMEM173\|TMEM2\|TMEM204\|TMEM255B\|TMEM70\|TMOD3\|TNFRSF10B\|TNFRSF10D\|TNKS1BP1\|TPM3\|TRIB1\|UPP1\|VEGFC\|VSIG2\|VWA1\|VWF\|WWTR1\| [illegible] |
| NOR (nC5) | 2 | AC006042.6\|ACTBL2 |
| Macrophages (nC6) | 364 | ABCA1\|ABCC3\|AC006547.13\|ACCS\|ACOT9\|ACP5\|ACSL1\|ADAM8\|ADAM9\|ADAP2\|ADPGK\|ALOX5\|AMDHD2\|AMPD2\|AMPD3\|AMZ1\|ANKRD9\|ANPEP\|APBB1IP\|APBB3\|APOE\|AQP9\|ARAP1\|ARHGAP18\|ARHGAP4\|ARL4C\|ARL8B\|ARRB2\|ASAH1\|ATG16L2\|ATP1B3\|AVPI1\|BCL2A1\|BHLHE41\|BMF\|BMP2K\|BTK\|C15orf48\|C1orf162\|C5AR1\|C9orf72\|CCL3L3\|CD14\|CD163\|CD300A\|CD300LF\|CD58\|CD72\|CD74\|CD83\|CD84\|CD86\|CDCP1\|CELF2\|CHMP1B\|CHST15\|CIITA\|CLEC7A\|CLN8\|CPVL\|CREG1\|CSAR2\|CSF1R\|CSF2RA\|CSF3R\|CSTB\|CTSB\|CTSD\|CTSH\|CTSL\|CTSS\|CTSZ\|CXCL1\|CXCL16\|CXCL2\|CXCL3\|CXCL8\|CYBB\|CYP27A1\|CYTH4\|DAB2\|DENND3\|DMXL2\|DOK3\|DPP7\|EEPD1\|ELL\|ELL2\|EMILIN2\|ENOSF1\|FAM20A\|FCER1G\|FCGR2C\|FCGRT\|FCHO2\|FERMT3\|FGD2\|FGR\|FKBP15\|FNIP2\|FOSL2\|FPR3\|FYB1\|GAB3\|GABARAPL1\|GGA1\|GGA2\|GK\|GLA\|GLIPR2\|GLUL\|GNA13\|GNA15\|GPCPD1\|GPNMB\|GPR137B\|GPR183\|GRAMD4\|GRINA\|GRN\|GS1-484O17.5\|GSAP\|GSTO1\|HAVCR2\|HCK\|HCLS1\|HERPUD1\|HEXA\|HEXB\|HIF1A\|HK2\|HK3\|HLA-DMA\|HLA-DMB\|HLA-DOA\|HLA-DPA1\|HLA-DPB1\|HLA-DQA1\|HLA-DQA2\|HLA-DQB1\|HLA-DQB2\|HLA-DRA\|HLA-DRB1\|HLA-DRB5\|HMOX1\|HPS1\|HPS3\|HS3ST2\|HSPA6\|HTRA4\|IFFO1\|IFI30\|IFNGR1\|IL18\|IL2RA\|IL4I1\|IL6R\|IRAK1\|IRF5\|IRF8\|ITGAD\|ITGAM\|ITGAX\|KCNAB2\|KLHL6\|KMO\|LACTB\|LAIR1\|LAPTM5\|LCP2\|LGALS3\|LGALS9\|LGMN\|LILRA6\|LILRB1\|LILRB2\|LILRB3\|LILRB4\|LILRB5\|LITAF\|LPCAT1\|LRP1\|MAFB\|MANBA\|MAP3K8\|MCL1\|MCOLN1\|MERTK\|METRNL\|MFSD1\|MFSD12\|MGAT1\|MGLL\|MIR181A1HG\|MKNK1\|MPEG1\|MPHOSPH6\|MPP1\|MS4A4A\|MS4A6A\|MS4A7\|MSR1\|MXD1\|MYO1F\|MYO9B\|NABP1\|NAMPT\|NBPF10\|NBPF19\|NCF2\|NCF4\|NEAT1\|NFKBID\|NPL\|NR1H3\|NR4A2\|ODF3B\|OLR1\|OSCAR\|OSM\|P2RX7\|PADI2\|PAPD4\|PAPSS2\|PARP14\|PARVG\|PDE4A\|PER1\|PFKFB3\|PGK1\|PIK3R5\|PLA2G7\|PLAUR\|PLCB2\|PLEK\|PLIN2\|PLSCR1\|PLXDC2\|PPARD\|PPIF\|PREX1\|PRKAG2\|PSAP\|PTK2B\|PTPN6\|RAB20\|RAB31\|RAB7B\|RAP2B\|RAPGEF1\|RASGEF1B\|RASSF2\|RASSF4\|RASSF5\|RBM47\|REL\|RGS1\|RGS2\|RHBDF2\|RIN3\|RIPK2\|RN7SL368P\|RNA5SP498\|RNASET2\|RNF13\|RNF135\|RNF144B\|RNF149\|RP11-106M3.3\|RP11-147L13.15\|RP11-202P11.1\|RP11-212I21.2\|RP11-213H15.3\|RP11-242C19.2\|RP11-325K4.2\|RP11-373D23.2\|RP11-473M20.9\|RP11-4B16.1\|RP11-52J3.2\|RP11-618G20.2\|RP11-668G10.2\|RP5-1171I10.5\|RREB1\|RXRA\|SAMHD1\|SAMSN1\|SAT1\|SCARB1\|SCARNA9\|SDC3\|SDS\|SEMA4A\|SETDB2\|SGK1\|SIGLEC1\|SLC11A1\|SLC11A2\|SLC15A3\|SLC16A10\|SLC16A3\|SLC16A6\|SLC17A9\|SLC18B1\|SLC1A3\|SLC2A14\|SLC31A2\|SLC37A2\|SLC43A2\|SLC7A8\|SLC8B1\|SLCO2B1\|SMAP2\|SMPDL3A\|SNAPC1\|SNHG12\|SNORD99\|SNX29\|SOAT1\|SPI1\|SRA1\|SRGN\|STK10\|STX4\|STX7\|STXBP2\|SYNGR2\|TBC1D12\|TBC1D2\|TCIRG1\|TFEC\|TFRC\|TGFBI\|TGIF1\|THEMIS2\|TIPARP\|TLR2\|TMEM52B\|TMEM63A\|TNFAIP2\|TNFRSF14\|TNFRSF1B\|TNS3\|TOM1\|TPCN1\|TPP1\|TREM1\|TRIM38\|TRPM2\|TUBGCP2\|TYMP\|TYROBP\|UAP1L1\|UBXN11\|VAC14\|VAMP8\|WDR91\|WIPI1\|XXbac-BPG254F23.5\|YTHDF3\|ZEB2\|ZFP36\|ZFP36L1\|ZMYND15\|ZNF331 |
| NOR (nC7) | 369 | AC002044.1\|AC004878.2\|AC006019.4\|AC012506.4\|AC109326.1\|AC138969.4\|AC159540.2\|ACAD10\|ACAP3\|ACTG1\|ADCY1\|ADGRB2\|ADGRL1\|AGAP2\|AGAP3\|AGER\|AGRN\|AHI1\|AKAP6\|AKAP9\|AMIGO2\|ANAPC2\|ANKRD12\|ANKRD13B\|ANKRD36\|ANKRD36B\|ANKRD36C\|ANKS1A\|APC2\|ARHGEF10L\|ARID1B\|ASIC3\|ASIC4\|ATAD5\|ATF6B\|ATG14\|B4GALNT4\|BICD1\|BMPR1B\|BPTF\|BRSK2\|BRWD1\|BTBD3\|C14orf132\|C1QTNF2\|C1orf226\|C2CD4A\|CALCOCO1\|CAMK2B\|CASP8AP2\|CASZ1\|CC2D1A\|CCDC136\|CCDC144NL\|CCNL2\|CDC42SE2\|CDH24\|CDK10\|CDK5RAP3\|CEP162\|CEP95\|CHD5\|CHD9\|CHGA\|CIC\|CNTFR\|CORO6\|COX11\|CRAMP1\|CRLF3\|CTC-378H22.2\|CTD-2349P21.10\|CTD-2349P21.5\|CTD-3014M21.4\|CYB561\|DAAM1\|DBH\|DBN1\|DCHS1\|DDC\|DDX17\|DDX39B\|DIRAS3\|DLC1\|DMTF1\|DNAJB2\|DNM1\|DOC2B\|DOCK4\|DOPEY1\|DPP6\|DST\|EBF1\|ECEL1\|EEF1A1\|EHMT2\|EIF1AY\|ELAVL2\|EML5\|EPB41L4A\|EPHB3\|ERC2\|ERV3-1\|EVL\|EXOC4\|EYA4\|EZH2\|FAM120B\|FAM184A\|FAM193B\|FASTK\|FNDC5\|FTX\|GABPB1-AS1\|GAS8\|GATA2\|GATA3\|GATS\|GCGR\|GFRA3\|GIGYF1\|GINM1\|GLG1\|GNAS\|GNB3\|GPATCH8\|GPR137C\|GPRASP1\|GRIK2\|GS1-18A18.1\|GSE1\|GTF2I\|GTF2IRD1\|GTF2IRD2\|GTF2IRD2B\|HBA1\|HBA2\|HBG1\|HECTD4\|HEXDC\|HID1\|HIP1R\|HMX1\|HS3ST5\|ICE1\|ING4\|INSRR\|INTS11\|ISYNA1\|KAT8\|KB-1572G7.1\|KCNAB3\|KCNQ1OT1\|KCNQ2\|KCNQ5\|KDM1A\|KDM5B\|KDM5D\|KIAA1683\|KLC4\|KLRK1\|KMT2E\|KRIT1\|LAMA4\|LINC00340\|LMBR1L\|LMO1\|LRRC16B\|LUC7L\|LUC7L2\|LUC7L3\|MAP1B\|MAP3K1\|MAPK8IP3\|MAPT\|MAST1\|MBTD1\|MC1R\|MCM7\|MDGA1\|MED23\|MEG3\|MEG8\|MEIS3\|MGAT4B\|MLH1\|MLLT4\|MLXIP\|MNAT1\|MTA1\|MTRNR2L1\|MTRNR2L12\|MTRNR2L5\|MTRNR2L8\|MUC20\|NBPF20\|NCAM1\|NCRNA00107\|NCRNA00185\|NDRG4\|NDUFA4L2\|NINL\|NISCH\|NKTR\|NOP56\|NOS1AP\|NPIPB15\|NPY\|NRCAM\|NRDE2\|NTRK1\|NXPH4\|OBSL1\|OCLN\|PABPN1\|PACS2\|PAXBP1\|PCDHA9\|PCLO\|PDE9A\|PEX6\|PHC1\|PHF14\|PHF19\|PHIP\|PI4KA\|PIGQ\|PKD1\|PLAGL1\|PLXNA4\|PLXNB1\|PNISR\|PNN\|POLR2J2\|POU2F2\|PPIP5K2\|PRPF4B\|PRPSAP2\|PRRC2B\|PRRT2\|PSD2\|PSMG4\|PTBP2\|PTPRF\|PXDN\|R3HDM2\|RABL6\|RADIL\|RAI1\|RBBP6\|RBCK1\|RBFOX1\|RBM33\|RBM6\|REC8\|RERE\|RGMB\|RGS7\|RHOT2\|RIMKLB\|RNF44\|RNU6-457P\|RP11-117L5.1\|RP11-1212A22.4\|RP11-124K4.1\|RP11-156P1.2\|RP11-166B2.1\|RP11-216L13.19\|RP11-274B21.14\|RP11-349A22.5\|RP11-416N2.4\|RP11-421E14.2\|RP11-439E19.3\|RP11-490G2.2\|RP11-499P20.2\|RP11-509E10.1\|RP11-509E10.3\|RP11-54O7.3\|RP11-56D16.8\|RP11-615I2.1\|RP11-640N20.4\|RP11-71L14.3\|RP11-728K20.2\|RP13-104F24.2\|RP13-152O15.2\|RP3-406A7.7\|RPS4Y1\|RPS6KL1\|RSBN1L\|RUNDC3A\|SDCCAG8\|SEMA6C\|SFPQ\|SFSWAP\|SGK494\|SGSM2\|SHC2\|SHPRH\|SLC18A1\|SLC19A1\|SLC25A29\|SLC38A1\|SMARCD1\|SMPD3\|SNAP91\|SOBP\|SOGA3\|SOX4\|SPG7\|SPPL2B\|SPTAN1\|SRGAP3\|SRRM1\|SRRM3\|SRRM4\|SRSF11\|STAG3\|STK38\|STMN1\|STMN2\|STXBP1\|TAF1C\|TBC1D24\|TBKBP1\|TCERG1\|TFAP2B\|TH\|TIA1\|TLE4\|TMEM191C\|TRIM33\|TRIP11\|TSC2\|TSTD2\|TTC3\|TUBA1A\|TXNDC16\|TXNIP\|UBAP1L\|UCHL1\|ULK1\|UNC5A\|USP9Y\|UTY\|VARS2\|VPS9D1\|WDR27\|WDR6\|WDR86\|WSB1\|XXbac-BPG283O16.9\|YJEFN3\|YLPM1\|ZC3H14\|ZCCHC11\|ZCCHC14\|ZFHX3\|ZFP90\|ZFY\|ZKSCAN1\|ZNF117\|ZNF138\|ZNF253\|ZNF292\|ZNF33A\|ZNF429\|ZNF512B\|ZNF608\|ZRANB2 |

<table>
<tr><td></td><td></td><td></td></tr>
<tr><td>NOR (nC8)</td><td>201</td><td>ACLY|AKR1B1|ANKRD54|ANKRD6|ARF5|ARL16|ASB6|ASIC1|ATP1B1|ATP5J2|ATP6V0E2|ATP6V1G1|AZIN1|BASP1|C11orf87|C9orf142|C9orf16|C9orf78|CARD19|CAT|CBX6|CCDC28A|CCDC68|CDYL2|CERK|CHMP5|CHRFAM7A|CHRNA3|CLTA|CNOT7|COPS6|COX6C|CRABP1|CREB5|CRNDE|CS|CUTA|CXADR|CYC1|CYCS|DACH1|DCLK1|DCLK3|DKK3|DLK1|DPYSL4|DUSP26|EBF3|EBP|EDF1|ELMO2|ENC1|EPB41L1|EPB41L4B|EYA1|FAM219A|FBP1|FECH|FRMD3|FRRS1L|FSD1L|GAL|GNAQ|GNB1|GPC6|HACD3|HDAC9|HMGN3|HNRNPDL|HTR3A|IFT20|ISCA1|ISCU|ITGB8|ITM2C|JUP|KBTBD6|KHDRBS3|KIF3B|KIT|KLHL35|LRRC42|MAPRE2|MMD|MRFAP1L1|MRPS21|MRPS26|NDRG3|NDUFA12|NDUFB9|NEFL|NEFM|NELL1|NFIB|NGRN|NHLH2|NORAD|NPTX2|NXPH1|OGFRL1|OLFM1|ONECUT2|OPRM1|OVCH1-AS1|PCMT1|PCSK1N|PCSK2|PDE10A|PDZD8|PHF1|PNMA1|POLR3F|PPP4R1|PSMA7|PSMC2|PSMD11|PTN|PTPN3|PTS|QPCT|RAB14|RAB15|RAB2A|RBM18|RGS3|RGS4|RHBDD2|RIN2|RNF10|RNF170|RP11-227F19.1|RP11-531A24.3|RP11-650L12.2|RP5-1085F17.3|RPRD1A|RRAGA|RUNX1T1|RUSC2|SAR1A|SCD5|SCOC|SCP2|SCYL2|SEC61B|SEL1L3|SERINC3|SERPINE2|SETBP1|SH3GLB2|SLC10A4|SLC25A4|SMAD9|SMU1|SNRPN|SNX18|SORCS1|SPIN1|SPTLC1|SRI|SSBP1|ST8SIA3|STMN3|STRAP|SURF4|SV2B|SYBU|THRA|THSD7A|TM2D3|TMBIM4|TMED2|TMED4|TMEFF2|TMEM163|TMEM176A|TMX2|TPI1|TRPA1|TSPAN11|TSPAN13|TUB|TUBA1B|TUBA1C|TUBB2A|TUBB4A|TUBB4B|TUBG1|TXN|UBQLN1|ULK3|VDAC3|VLDLR|VPS4B|VSNL1|WDR47|WDR61|WRB|WSB2|YWHAZ|ZBTB18|ZNF770</td></tr>
<tr><td>NOR (nC9)</td><td>335</td><td>AASDHPPT|ABHD2|AC073283.4|ACVR2B|ADD2|AFAP1|AL022344.4|ALCAM|ALKBH7|ANKRD17|ARHGAP35|ARL6IP5|ASTN1|ATF2|ATP11A|ATP2C1|ATP5D|ATP5I|ATRX|BCL2|BEX2|BMP7|BRINP2|BRK1|BSG|BTF3|C19orf12|C1orf21|C4orf33|CAMK2N1|CAMK4|CAMLG|CAMSAP2|CANX|CCDC186|CCDC82|CCT4|CCT7|CD47|CENPJ|CHGB|CHN1|CHST1|CKS1B|CLCN3|CLGN|CLIP3|CNRIP1|CNTN1|CNTNAP4|COMMD1|COPE|COX6B1|COX7C|COX8A|CPE|CPNE8|CREG2|CSRNP3|CTD-2192J16.22|CTD-2314G24.2|CXXC4|CYGB|DCX|DDAH1|DDX6|DLEU2|DPY30|DPYSL3|DSEL|DUSP8|DUT|EEF1B2|EEF2|EHBP1|EIF1B|EIF2AK2|EIF3L|EML6|EPM2AIP1|EPRS|ESD|FAM134A|FAM171A1|FAM63B|FANCL|FASTKD1|FAU|FBXO30|FIGN|FMN1|FMO1|FOXP1|FSTL5|FXR1|GAB2|GABARAPL2|GCSH|GIGYF2|GNAO1|GPR22|GSK3B|GSTP1|H1F0|H2AFZ|HADHA|HDGF|HDLBP|HIBCH|HINT1|HIST1H4C|HIST1H4E|HLTF|HMGA1|HMGB1|HMGB2|HNRNPA3|HNRNPL|HSD17B11|HSD17B12|HSPA8|HTATSF1|IGFBP5|INO80D|INPP5F|IRS2|ITGA1|JARID2|JPH3|JTB|JUN|KCNG1|KCNK3|KCNQ3|KCTD16|KIAA1549L|KIAA1715|KIDINS220|KIF26A|KIF26B|KIF2A|KLHL23|LANCL1|LDHB|LIFR|LL22NC03-2H8.5|LONRF2|LRRC75A-AS1|LRRN3|LRRTM2|MAP4|MARCH4|MARCH7|MBNL2|MEIS2|METTL9|MIR3654|MOB1B|MRPS5|MST1|MTHFD2|MTMR6|MXRA7|N4BP2|NAALAD2|NCK2|NDUFA1|NDUFB11|NDUFS1|NMNAT2|NRP1|NTNG1|OIP5-AS1|OPTN|OST4|OXCT1|PARD3|PBRM1|PCDH10|PCDH7|PCDHB10|PCDHB16|PCDHB6|PCDHB9|PCNP|PFN2|PGAP1|PGM2L1|PHYHIPL|PITPNB|PJA2|PKIB|PLCXD3|PMP22|PODXL2|POLR1D|PPP2R1A|PRDX2|PREPL|PROX1|RAB11B|RAB3C|RACK1|RAMP1|RBMS1|RBMS3|RGS5|RMND5A|RND3|RNF144A|RP11-118B22.2|RP11-133K1.6|RP11-264B14.1|RP11-264B14.2|RP11-346C4.2|RP11-3P17.4|RP11-3P17.5|RP11-727A23.7|RP11-92K2.2|RP4-555D20.2|RP5-827C21.2|RPL14|RPL15|RPL18|RPL18A|RPL23A|RPL24|RPL26|RPL27|RPL29|RPL31|RPL32|RPL34|RPL35A|RPL39L|RPL4|RPS10|RPS11|RPS13|RPS15|RPS15A|RPS19|RPS21|RPS24|RPS27A|RPS3|RPS5|RPS7|RPS9|RPSA|RSF1|RSL24D1|RUFY3|SACS|SAE1|SCARB2|SCG2|SCG3|SCG5|SCN7A|SCRN1|SEC62|SEMA6D|SEPT11|SEPT6|SESTD1|SETD2|SEZ6|SEZ6L|SF3B2|SFRP1|SLC1A4|SLC25A12|SLC39A10|SLC6A2|SLIT3|SMARCA1|SMARCA5|SMARCC1|SMC1A|SNORD114-20|SNORD114-23|SNRNP27|SNRPD2|SPATS2L|SSB|ST3GAL6|ST6GALNAC5|STAC|STMN4|SUMO1|SV2C|SYNPO2|SYP|SYT1|SYT13|SYT4|TAGLN3|TARS|TBCB|TCF12|TEAD1|TET3|TFDP2|TIAL1|TKT|TLE3|TMA7|TMCO3|TMEM255A|TMEM59L|TMOD2|TMSB15A|TNRC6B|TOMM22|TOP2B|TRIM13|TSHZ2|TSPAN7|TTBK2|U2SURP|UBA52|UBE2E3|UQCRC1|VAT1L|VPS36|WDR82|XRCC5|XRN1|YBX1|ZBTB38|ZC3H13|ZC3H15|ZDBF2|ZNF148</td></tr>
<tr><td>T-cells (nC10)</td><td>110</td><td>AL022344.7|CD2|CD69|CLSTN2|CSAG2|CSAG3|FAM19A3|ICOS|IGHA1|IGHA2|IGHG1|IGHG2|IGHG3|IGHG4|IGHJ3|IGHJ4|IGHJ5|IGHJ6|IGHV1-24|IGHV1-46|IGHV1-69|IGHV1-69-2|IGHV1-69D|IGHV3-11|IGHV3-13|IGHV3-15|IGHV3-16|IGHV3-20|IGHV3-21|IGHV3-23|IGHV3-30|IGHV3-33|IGHV3-35|IGHV3-38|IGHV3-43|IGHV3-48|IGHV3-53|IGHV3-66|IGHV3-7|IGHV3-72|IGHV3-74|IGHV3OR15-7|IGHV3OR16-10|IGHV3OR16-13|IGHV3OR16-8|IGHV3OR16-9|IGKC|IGKJ1|IGKV1-12|IGKV1-17|IGKV1-33|IGKV1-37|IGKV1-39|IGKV1-5|IGKV1-6|IGKV1-8|IGKV1-9|IGKV1D-12|IGKV1D-13|IGKV1D-17|IGKV1D-33|IGKV1D-37|IGKV1D-39|IGKV1D-42|IGKV1D-8|IGKV2-28|IGKV2-30|IGKV2-40|IGKV2D-28|IGKV2D-30|IGKV2D-40|IGKV3-11|IGKV3-15|IGKV3-20|IGKV3-7|IGKV3D-11|IGKV3D-15|IGKV3D-20|IGKV3D-7|IGKV3OR2-268|IGKV4-1|IGLC2|IGLC3|IGLC7|IGLJ2|IGLJ3|IGLL1|IGLV1-47|IGLV2-11|IGLV2-14|IGLV2-18|IGLV2-23|IGLV2-8|IGLV3-1|IGLV3-16|IGLV3-21|IGLV3-25|IGLV3-9|LL22NC03-N14H11.1|LTB|ODZ4|PITX1|PNMA5|PPM1J|RP11-812E19.9|SH2D2A|TRAC|TRBC1|TRBC2|VWA5B2</td></tr>
</table>
