## Supplementary Table 3 for "Single-nuclei transcriptomes from human adrenal gland reveals distinct cellular identities of low and high-risk neuroblastoma tumors"

**Human post-natal AG to mouse post-natal AG (Figure 1)**

| Cell cluster in human AG | Cell cluster in mouse AG (reference) | Number of genes in the specific signature of human AG shared with the specific signature of mouse AG | Genes in the specific signature of human AG shared with the specific signature of mouse AG | FDR | Significance |
|---|---|---|---|---|---|
| Chromaffin (hC4) | Chromaffin nAChR α7$^{high}$ (mC15) | 63 | ACE\|ADCYAP1R1\|AP3B2\|ARFGEF3\|CACNA1B\|CACNA1D\|CAMK2B\|CELF3\|CELF4\|CHGA\|CHGB\|CHRNA3\|CNKSR2\|CRMP1\|DBH\|DGKB\|DPYSL3\|DUSP26\|ELAVL4\|EML5\|FAM171B\|FSTL5\|GCGR\|GPRASP1\|GPRASP2\|GRIA2\|HMP19\|KCNQ2\|MAPT\|MTCL1\|MYT1\|MYT1L\|NT5DC2\|PCLO\|PHOX2A\|PHOX2B\|PLEKHA6\|PPFIA2\|PRUNE2\|PTPRN\|RAP1GAP2\|SCG2\|SCN3B\|SCN9A\|SGIP1\|SLC18A1\|SLC18A2\|SNAP25\|SNAP91\|SRRM3\|ST8SIA3\|STXBP1\|SYT1\|SYT4\|TACC2\|TBX20\|TLX2\|TPPP3\|UNC5A\|UNC5C\|UNC79\|UNC80\|YPEL4 | <1E-50 | ** |
| Chromaffin (hC4) | Chromaffin nAChR α7$^{low}$ (mC11) | 4 | ATP1B1\|CYB561\|SLC31A1\|UCHL1 | 1.26E-03 | ** |
| Endothelial (hC6) | Endothelial (mC2) | 3 | ADGRL4\|F8\|PTPRB | 1.38E-03 | ** |
| Endothelial (hC6) | Endothelial (mC4) | 23 | ARHGEF15\|BTNL9\|CLDN5\|DOCK6\|DYSF\|EFNA1\|EFNB2\|EGFL7\|FLT1\|FLT4\|HSPG2\|IGFBP5\|KDR\|LMO2\|NOS3\|PLAT\|PLK2\|PLVAP\|RASIP1\|SCARF1\|SHANK3\|SLC9A3R2\|TMEM88 | 2.39E-32 | ** |
| Macrophages (hC2) | Macrophages (mC14) | 10 | FGD2\|FGL2\|FMNL2\|GSAP\|HCLS1\|IRF5\|KCTD12\|PTPN18\|RNASE6\|TRPM2 | 6.58E-07 | ** |
| Macrophages (hC2) | Macrophages (mC3) | 29 | C1QA\|C1QB\|C1QC\|C5AR1\|CLEC7A\|CSF1R\|CSF3R\|CTSB\|DOK3\|F13A1\|FMN1\|FRMD4B\|LAIR1\|LST1\|MAF\|MAFB\|MRC1\|MS4A14\|MS4A7\|MSR1\|NPL\|SAT1\|SIGLEC1\|SLC11A1\|STAB1\|TLR1\|TMEM106A\|TREM2\|VSIR | 1.98E-37 | ** |
| Macrophages (hC2) | Macrophages (mC8) | 4 | ALOX5\|CD163\|EMILIN2\|FGR | 3.38E-03 | ** |
| Mesenchymal (hC7) | Glial (mC10) | 6 | COL27A1\|FBLN5\|NGFR\|NID1\|S1PR3\|SCN7A | 1.61E-02 | * |
| Mesenchymal (hC7) | Mesenchymal (mC6) | 14 | ADAMTSL2\|BGN\|C7\|COLEC11\|CRISPLD2\|CXCL12\|CYP1B1\|ERRFI1\|GPX3\|LUM\|MFGE8\|PDGFRB\|STEAP4\|WISP1 | 1.46E-18 | ** |
| T-cells (hC10) | T-cells (mC17) | 3 | CD3D\|CD3E\|CD6 | 2.68E-05 | ** |
| T-cells (hC10) | T-cells (mC18) | 2 | CCL5\|HCST | 1.61E-02 | * |

**Human post-natal AG to human fetal AG (Dong et al. 2020)**

| Cell cluster in human AG | Cell cluster in human fetal AG (Dong et al. 2020, reference) | Number of genes in the specific signature of human AG shared with the signature of human fetal AG (Dong et al. 2020) | Genes in the specific signature of human AG shared with the signature of human fetal AG (Dong et al. 2020) | FDR | Significance |
|---|---|---|---|---|---|

| | | | | | |
|---|---|---|---|---|---|
| Chromaffin (hC4) | Non-cycling chromaffin cells* | 60 | ADM\|AMPD2\|ARC\|ARFGEF3\|ATP1B1\|BAIAP2\|CALY\|CARTPT\|CHGA\|CHGB\|CHRNA3\|CYB561\|DBH\|DDC\|DNALI1\|DPP6\|EFHC1\|FAIM2\|FAM162B\|GCGR\|GCH1\|GRIA2\|HTATSF1\|JPH4\|KCNQ2\|KHDRBS3\|LINC00632\|LRRN2\|MGAT4C\|MIR7-3HG\|NDRG4\|NT5DC2\|PCSK1N\|PCSK2\|PENK\|PNMT\|PPP1R1B\|PRCD\|PTPRN\|RAB3C\|RAMP1\|RGS4\|RSPH1\|SCG2\|SCG5\|SEZ6L2\|SLC18A1\|SLC18A2\|SLC22A17\|SLC24A2\|SLC31A1\|SPOCK3\|STXBP1\|SYT1\|SYT4\|SYT5\|TH\|TIAM1\|VAT1L\|VWA5B2 | <1E-50 | ** |
| Chromaffin (hC4) | Non-cycling sympathoblast* | 43 | AKR1C1\|AKR1C2\|ALCAM\|ATP1A3\|CADPS\|CAMK2B\|CD24\|CELF4\|CRMP1\|DPYSL3\|DUSP26\|ELAVL4\|ENO2\|HAND1\|INA\|KRT19\|MAP1A\|MAP7\|MAPT\|MIAT\|NEFM\|PAK3\|PCBP4\|PHOX2A\|PHOX2B\|PRPH\|RGS4\|RGS5\|RTN1\|SCG3\|SCG5\|SCN3B\|SCN9A\|SLC6A2\|SNAP25\|STMN2\|SULT4A1\|SYT11\|TAGLN3\|TFAP2B\|TMOD1\|UCHL1\|VSTM2L | 7.38E-41 | ** |
| Chromaffin (hC4) | Cycling chromaffin cells* | 41 | ADGRA1\|ADM\|ARC\|ARFGEF3\|BAIAP2\|CADM2\|CARTPT\|CCP110\|CHGA\|CHGB\|CNTN1\|FAM162B\|FLRT3\|GCH1\|HTATSF1\|KCNQ2\|MGAT4C\|NDRG4\|NEBL\|NT5DC2\|PCSK1N\|PENK\|PNMT\|PPP1R1B\|PTPRN\|RAMP1\|ROBO1\|SAMD5\|SCG2\|SGPL1\|SLC18A1\|SLC24A2\|SLC31A1\|SPOCK3\|SST\|STXBP1\|SYT4\|SYT5\|TH\|TIAM1\|VAT1L | 3.69E-28 | ** |
| Chromaffin (hC4) | Cycling sympathoblast* | 22 | ANK2\|ATCAY\|CCP110\|CLASP2\|ELAVL4\|ENO2\|HAND2\|ICA1\|INPP5F\|MAP2\|MYEF2\|NBEA\|NCAM1\|PHOX2B\|RP11-457P14.4\|SESN3\|SLC4A8\|SNAP91\|TFAP2B\|TMX4\|VSTM2L\|ZNF704 | 6.11E-12 | ** |
| Endothelial (hC6) | SCPs | 11 | DHRS3\|GSN\|HSPG2\|ID1\|IGFBP4\|IGFBP5\|NOSTRIN\|PLAT\|PLPP1\|PRSS23\|SERPINB6 | 3.54E-07 | ** |
| Mesenchymal (hC7) | SCPs | 20 | BGN\|C11orf96\|C7\|CALD1\|COL1A1\|COL1A2\|COL6A2\|CSRP1\|EFEMP2\|MATN2\|MFGE8\|MMP2\|MXRA8\|NGFR\|NID1\|NR2F1\|RBPMS\|SCN7A\|TIMP1\|VCL | 1.68E-16 | ** |
| Progenitor (hC1) | Cycling sympathoblast* | 11 | ASPM\|BUB1B\|ESCO2\|KIF14\|KIF4A\|MCM10\|MMS22L\|NCAPH\|NEK2\|RTKN2\|TOP2A | 7.42E-03 | ** |
| Progenitor (hC1) | Cycling chromaffin cells* | 11 | ANLN\|ASPM\|BUB1\|BUB1B\|KIF4A\|LAMA3\|MCM10\|NEK2\|PHGDH\|RTKN2\|TOP2A | 3.85E-02 | * |
| Z. fasciculata (hC3) | SCPs | 23 | ALAS1\|ATP1B3\|AXL\|CASP9\|CYP11B1\|GALNT2\|GRAMD1B\|HOXA5\|HSPE1\|LDLR\|MC2R\|PAPSS2\|PMEPA1\|RAB20\|RP13-49I15.5\|SH3BP5\|SLC16A9\|SLC26A2\|STAR\|YBX3\|ZFAND5\|ZNF503\|ZNF536 | 5.18E-18 | ** |
| Z. fasciculata (hC3) | Non-cycling chromaffin cells* | 6 | GUK1\|MIDN\|PRMT9\|VEGFA\|ZDBF2\|ZNF331 | 1.10E-02 | * |
| Z. glomerulosa (hC5) | SCPs | 5 | ARRDC3\|IRS1\|PHACTR2\|UGCG\|VCAN | 1.14E-05 | ** |

**Human post-natal AG to mouse E13 adrenal anlagen**

| Cell cluster in human AG | Cell cluster in mouse E13 adrenal anlagen (reference) | Number of genes in the specific signature of human AG cluster shared with the specific signature of mouse E13 adrenal anlagen | Genes in the specific signature of human AG cluster shared with the specific signature of mouse E13 adrenal anlagen | FDR | Significance |
|---|---|---|---|---|---|
| Chromaffin (hC4) | Bridge cells | 5 | DPYSL3\|HDAC9\|OGDHL\|ROBO1\|TLX2 | 1.24E-02 | * |
| Chromaffin (hC4) | Chromaffin cells | 71 | ACE\|ADGRB3\|ADM\|AHI1\|ARHGAP36\|BAIAP3\|BSCL2\|CACNA1D\|CALY\|CCDC148\|CHD5\|CHGA\|CHGB\|CNKSR2\|CNTN3\|CYB561\|DBH\|DDC\|DGKB\|DPP6\|DRD2\|DUSP26\|DYNC2H1\|EML5\|ENO2\|FAIM2\|FSTL5\|GCH1\|GJD2\|GNB3\|GPRASP1\|GPRASP2\|GRIA4\|HAND2\|JPH4\|KCNK9\|KIF1A\|KSR2\|MAPRE3\|OCRL\|PCSK1N\|PER3\|PPFIA2\|PRKCE\|PTPRN\|RAB3C\|RGS4\|RGS5\|RIMBP2\|RSPH1\|RUNDC3A\|SAMD11\|SCG2\|SCG5\|SEZ6L\|SEZ6L2\|SLC18A1\|SLC18A2\|SLC24A2\|SLC31A1\|SLC4A8\|SNAP25\|SNAP91\|SYT1\|TH\|TMEM130\|TMX4\|TUB\|UNC5A\|UNC80\|ZFR2 | 1.00E-31 | ** |

| | | | | | |
|---|---|---|---|---|---|
| Chromaffin (hC4) | Sympathoblast cells | 44 | ABCC8\|ADGRA1\|AKAP6\|ANK2\|ARHGAP33\|ATCAY\|ATP1B1\|CACNA1E\|CACNA2D3\|CARTPT\|CRMP1\|FAM171B\|HMP19\|INA\|INPP5F\|KCNQ2\|KHDRBS3\|MAP7\|MARK1\|MAST1\|MPP2\|NDRG4\|NRSN1\|PCSK2\|PHOX2B\|PKIA\|PLEKHA6\|PRPH\|RET\|RTN1\|SCG3\|SCN9A\|SLC6A2\|SPOCK3\|SST\|STMN2\|SULT4A1\|SYT11\|TAGLN3\|TMEM179\|TMEM59L\|TMOD1\|UCHL1\|VSTM2L | 4.25E-10 | ** |
| Endothelial (hC6) | SCPs | 13 | APOLD1\|CASKIN2\|CLEC14A\|DHRS3\|ELK3\|EPHB4\|HSPG2\|KCNIP4\|LDB2\|LIMS2\|PLAT\|PRSS23\|TM4SF1 | 1.24E-02 | * |
| Mesenchymal (hC7) | SCPs | 31 | CALD1\|COL12A1\|COL16A1\|COL1A2\|COL27A1\|COL3A1\|COL6A1\|COL6A2\|CRISPLD2\|CSRP1\|CTGF\|CYR61\|EFEMP2\|F3\|FBLN5\|FERMT2\|GPC3\|LAMA2\|MAMDC2\|MATN2\|MFGE8\|MMP2\|NGFR\|NID1\|NR2F1\|PEAR1\|PLA2R1\|PLEKHG2\|RBPMS\|TAGLN\|VCL | 3.72E-15 | ** |
| Progenitor (hC1) | Sympathoblast cells | 22 | ANLN\|ARPP21\|ASPM\|ASXL3\|BRCA1\|BUB1\|CENPI\|CLDN11\|DIAPH3\|ESCO2\|KIF14\|KIF15\|KIF4A\|KNTC1\|MCM10\|MMS22L\|MTUS2\|NCAPH\|NEK2\|POLQ\|RTKN2\|TOP2A | 1.61E-02 | * |
