## Supplementary Table 4 for "Single-nuclei transcriptomes from human adrenal gland reveals distinct cellular identities of low and high-risk neuroblastoma tumors"

#### Human NB to van Groningen et al. 2017 (Figure 3)

<table>
<thead>
<tr><th>Cell cluster in human NB</th><th>Cell type in van Groningen et al. 2017 (reference)</th><th>Number of genes significantly (FDR <0.01) upregulated in human NB overlapping genes in van Groningen et al. 2017</th><th>Genes significantly (FDR <0.01) upregulated in human NB overlapping genes in van Groningen et al. 2017</th><th>FDR</th><th>Significance</th></tr>
</thead>
<tbody>
<tr><td>Endothelial (nC4)</td><td>MES</td><td>211</td><td>A2M|ACADVL|ACTN1|ADAMTS5|AEBP1|ANXA1|ANXA2|ANXA5|APP|ARL4A|ARPC1B|ASPH|ATP2B1|ATP2B4|ATP6V0E1|B2M|BAG3|BGN|C19orf10|C1orf54|C6orf120|CALD1|CAPN2|CBLB|CD59|CD63|CETN2|CFI|CKAP4|CLIC4|CMTM3|CNN3|COL4A1|COL4A2|CRABP2|CRTAP|CSRP1|CTNNA1|CTSB|CTSO|CYR61|DLC1|DLX1|DUSP14|DUSP5|DUSP6|EGR1|EGR3|EHD2|ELF1|ELK3|EMP1|ETS1|EXT1|F2R|FAM120A|FAM43A|FGFR1|FLNA|FN1|FNDC3B|FSTL1|FZD1|FZD2|GJA1|GNG12|GRN|GSN|HES1|HIPK3|HIST1H2AC|HLA-A|HLA-B|HLA-C|HLA-F|HLX|HSP90B1|HSPA5|HSPB1|HTRA1|ID1|ID3|IFI16|IFITM2|IFITM3|IGFBP6|IL13RA1|IL6ST|INSIG1|ITGA10|ITGAV|ITGB1|ITM2B|ITPRIPL2|JAK1|JAM3|KCTD12|KIAA1462|KLF10|KLF4|KLF6|LAMB1|LAMC1|LAMP1|LAPTM4A|LATS2|LEPROT|LGALS1|LITAF|LMNA|LOXL2|LRP10|LRRC8C|LUZP1|MAGT1|MANF|MBNL1|MEOX1|MEOX2|MGP|MMP2|MYADM|MYL12A|MYL12B|MYLIP|NANS|NFIA|NFIC|NPC2|NPTN|PDIA3|PDLIM1|PEA15|PEAK1|PLK2|PLOD3|PLPP1|PLS3|PLSCR1|PLSCR4|PON2|POSTN|PPIC|PRCP|PRDX4|PRDX6|PTBP1|PTPN14|PTRF|PXDC1|QKI|QSOX1|RAB13|RAP1A|RAP1B|RBMS1|RCN1|RGS3|RHOC|RHOJ|RNH1|RRBP1|SASH1|SDCBP|SDF4|SEC14L1|SEMA3F|SEPT10|SERPINH1|SGK1|SH3BGRL|SHC1|SKIL|SLC30A1|SLC39A14|SMAD3|SOX9|SPARC|SPARCL1|SPATA20|SPCS3|SPRED1|SPRY1|SPRY4|SSBP4|STAT3|SURF4|SVIL|SYPL1|TFPI|TGFB1I1|TGFBR2|TIMP1|TJP1|TM4SF1|TMED9|TMEM50A|TNFRSF12A|TNFRSF1A|TNS1|TPBG|TRAM1|TSC22D2|TSC22D3|TSPAN4|TUBB6|VCL|VIM|WLS|WWTR1|ZFP36L1</td><td>6.56E-39</td><td>**</td></tr>
<tr><td>Macrophages (nC6)</td><td>MES</td><td>76</td><td>ACADVL|ACAP2|ADAM9|ADGRE5|ANXA1|ANXA2|ANXA5|APOE|ATP2B1|ATP6V0E1|B2M|CD164|CD44|CMTM3|CREG1|CTSB|CYFIP1|EDEM1|ELF1|F2RL2|FAM102B|FLNA|FNDC3B|GNS|GPR137B|GRN|GSN|HEXB|HLA-A|HLA-B|HLA-C|HLA-F|HNMT|IFI16|IGF2R|IL13RA1|INSIG1|IQGAP2|KLF6|LAMP1|LGALS1|LITAF|LMNA|MAML2|MAN2A1|MBNL1|NANS|NOTCH2|NOTCH2NL|NPC2|P4HA1|PAPSS2|PLEKHA2|PLSCR1|PLXDC2|PTGER4|PYGL|QKI|RAB31|RGL1|RIT1|RNH1|RRBP1|SDCBP|SEC14L1|SGK1|SH3BGRL|SLC38A6|STAT1|STAT3|TNFRSF1A|TSC22D3|TSPAN4|VIM|WIPI1|ZFP36L1</td><td>1.86E-19</td><td>**</td></tr>
<tr><td>NOR (nC5)</td><td>ADRN</td><td>154</td><td>ABCA3|ACTL6B|ADCYAP1R1|ADGRB3|ADRBK2|ANK2|ARHGEF7|ARL6IP1|ATCAY|ATP6V0E2|AUTS2|BEX1|BMPR1B|C14orf132|CACNA1B|CADM1|CAMSAP1|CCDC167|CCND1|CCNI|CDKN2C|CENPU|CENPV|CERK|CHGA|CHGB|CHRNA3|CKB|CLASP2|CRMP1|DAPK1|DBH|DCX|DDC|DPYSL2|DPYSL3|DPYSL5|EIF1B|ELAVL2|ELAVL3|ELAVL4|ENO2|EVL|EYA1|FAM163A|FEV|FOXO3|FSD1|GAP43|GATA2|GATA3|GDI1|GLCCI1|GMNN|GNB1|GNG4|GPR22|GRIA2|H1FX|HAND1|HAND2-AS1|HMGA1|HN1|HNRNPA0|ICA1|IGFBPL1|INA|INSM2|ISL1|KDM1A|KIAA1211|KIDINS220|KIF1A|KIF21A|KIF5C|KLC1|KLF7|KLHL23|L1CAM|LSM4|MAGI3|MAP1B|MAP2|MAP6|MAPT|MCM7|MIAT|MSH6|MSI2|MYEF2|NBEA|NCAM1|NCOA7|NELFCD|NNAT|NPY|NRCAM|NRSN1|NUSAP1|PEG3|PHOX2A|PHOX2B|PKIA|PPP1R9A|PRCD|PRPH|RAB6B|RALGDS|RANBP1|RBMS3|RBP1|REEP1|RGS5|RIMBP2|RIMS3|RNF144A|RNF165|RTN1|RUNDC3A|SBK1|SCAMP5|SCG2|SCG3|SCN3A|SEPT3|SEPT6|SHD|SIX3|SLIT1|SLIT3|SNAP91|SOX11|STMN2|STMN4|STXBP1|SV2C|SYT1|TAGLN3|TFAP2B|TH|THSD7A|TMOD1|TMTC4|TTC8|TUB|TUBB2A|TUBB2B|TUBB3|TUBB4B|UBE2C|UBE2T|ZNF536|ZNF704|ZNF711</td><td><1E-50</td><td>**</td></tr>
<tr><td>NOR (nC5)</td><td>MES</td><td>30</td><td>ACADVL|ACTA2|ACTN1|C1orf198|CBFB|COL6A1|CTDSP2|DLC1|ELAVL1|ENAH|HIBADH|HSPB1|ITM2B|ITM2C|KDELR2|KDELR3|KDM5B|MEST|NES|PTN|PXDN|RECK|ROBO1|SCPEP1|SERPINE2|SHC1|TMEFF2|TPM1|TRIL|TUBB6</td><td>6.46E-03</td><td>**</td></tr>
<tr><td>NOR (nC7)</td><td>ADRN</td><td>163</td><td>ABCA3|ACTL6B|ACVR1B|ADGRB3|ADRBK2|AGTPBP1|AKAP12|ANK2|ARHGEF7|ARL6IP1|ATCAY|AUTS2|BEX1|BEX2|BMPR1B|C11orf95|C14orf132|C20orf100|CACNA1B|CADM1|CAMSAP1|CCND1|CCNI|CENPV|CHGA|CHGB|CHRNA3|CKB|CLASP2|CRMP1|CYFIP2|CYGB|DAPK1|DBH|DCX|DDC|DIABLO|DNER|DPYSL2|DPYSL3|DPYSL5|EIF1B|ELAVL2|ELAVL3|ELAVL4|ENO2|EPB41L4A-AS1|EVL|FAM163A|FEV|FOXO3|FSD1|GAP43|GATA2|GATA3|GCH1|GDAP1|GDAP1L1|GDI1|GDPD1|GLCCI1|GNB1|GRIA2|H1FX|HAND2-AS1|HMP19|HN1|HNRNPA0|ICA1|INA|INSM2|IRS2|ISL1|KDM1A|KIDINS220|KIF1A|KIF21A|KIF2A|KIF5C|KLC1|KLF7|KLHL23|L1CAM|MAGI3|MAP1B|MAP2|MAP6|MAPK8|MAPT|MARCH11|MCM7|MIAT|MSH6|MXI1|MYEF2|NBEA|NCAM1|NCOA7|NCS1|NELFCD|NELL2|NNAT|NPY|NRCAM|NRSN1|NSG1|PARP6|PEG3|PHOX2A|PHOX2B|PIK3R1|PKIA|PRIM1|PRPH|RAB6B|RALGDS|RBMS3|RBP1|REC8|REEP1|RGS17|RGS5|RIMBP2|RIMS3|RNF157|RNF165|RTN1|RUFY3|RUNDC3A|RUNDC3B|SATB1|SBK1|SCAMP5|SCG2|SCN3A|SHD|SLIT3|SNAP25|SNAP91|SOX11|STMN2|STMN4|STXBP1|SYT1|TAGLN3|TFAP2B|TH|THSD7A|TIAM1|TMOD1|TMOD2|TTC8|TUB|TUBB2A|TUBB2B|TUBB4B|UNC79|ZNF195|ZNF512|ZNF704|ZNF711|ZNF738|ZNF91</td><td><1E-50</td><td>**</td></tr>
</tbody>
</table>

| | | | | | |
|---|---|---|---|---|---|
| NOR (nC8) | ADRN | 183 | ACOT7\|ACTL6B\|ACVR1B\|AKAP12\|ANK2\|ANKRD46\|AP1S2\|ARHGEF7\|ARL6IP1\|ATCAY\|ATP6V0E2\|ATP6V1B2\|AUTS2\|BEND4\|BEX1\|BMPR1B\|C11orf95\|C14orf132\|CACNA1B\|CADM1\|CAMSAP1\|CCDC167\|CCND1\|CCNI\|CCSAP\|CDC42EP3\|CENPV\|CERK\|CHGA\|CHRNA3\|CKB\|CLASP2\|CRMP1\|CSE1L\|CXADR\|CXCR4\|CYGB\|DACH1\|DAPK1\|DBH\|DCX\|DDC\|DLK1\|DNAJC6\|DNER\|DPYSL2\|DPYSL3\|DPYSL5\|DTD1\|ELAVL3\|ELAVL4\|ENO2\|EVL\|EYA1\|FAM171B\|FAXC\|FOXO3\|GABRB3\|GAL\|GAP43\|GATA3\|GDAP1\|GDAP1L1\|GDI1\|GLCCI1\|GNB1\|GNG4\|HAND1\|HAND2-AS1\|HMP19\|HN1\|HNRNPA0\|HS6ST2\|ICA1\|IGSF3\|INA\|IRS2\|ISL1\|KIDINS220\|KIF1A\|KIF21A\|KIF5C\|KLC1\|KLF7\|KLHL13\|KLHL23\|L1CAM\|LEPROTL1\|MAP1B\|MAP2\|MAP6\|MAPK8\|MAPT\|MARCH11\|MCM7\|MIAT\|MMD\|MSI2\|MTCL1\|MYEF2\|NAP1L5\|NAPB\|NCAM1\|NCOA7\|NCS1\|NEFL\|NEFM\|NELFCD\|NELL2\|NGRN\|NNAT\|NPTX2\|NPY\|NRCAM\|NRSN1\|NSG1\|ODZ4\|OLFM1\|PARP6\|PBX3\|PHOX2A\|PHOX2B\|PHPT1\|PIK3R1\|PKIA\|PNMA2\|PRPH\|PTS\|QDPR\|RAB6B\|RALGDS\|RANBP1\|RBMS3\|RBP1\|REC8\|REEP1\|RGS17\|RIMBP2\|RIMS3\|RNF165\|RPS6KA2\|RTN1\|RUFY3\|RUNDC3A\|RUNDC3B\|SATB1\|SCAMP5\|SCG3\|SCN3A\|SEC11C\|SEPT3\|SEPT6\|SIX3\|SLC10A4\|SLIT1\|SNAP25\|SNAP91\|SOX11\|STMN2\|STMN4\|STXBP1\|SYT1\|TACC2\|TAGLN3\|TBPL1\|TDG\|TFAP2B\|TH\|THSD7A\|TMOD1\|TMOD2\|TMTC4\|TSPAN13\|TSPAN7\|TUB\|TUBB2A\|TUBB2B\|TUBB4B\|ZNF195\|ZNF24\|ZNF512\|ZNF704\|ZNF711 | <1E-50 | ** |
| NOR (nC8) | MES | 76 | ACADVL\|ACTN1\|AEBP1\|ALDH1A3\|ANXA5\|ANXA6\|APP\|ARHGAP1\|ATP10D\|ATP1B1\|ATP2B4\|B2M\|CALD1\|CD59\|CD63\|CNN3\|COL6A1\|COL6A2\|COPA\|CPED1\|CRABP2\|CTSB\|CYFIP1\|CYR61\|DKK3\|DUSP5\|EGR1\|ENAH\|FAM120A\|GPC6\|HLA-A\|ID1\|ITM2B\|ITM2C\|JAM3\|KDELR2\|KLF10\|KLF4\|LAMB1\|LIX1L\|LMAN1\|MEST\|MYL12A\|MYL12B\|NPTN\|OGFRL1\|OSTC\|PDE3A\|PPIB\|PTN\|PXDN\|RECK\|RGS3\|RHOC\|RIN2\|SCPEP1\|SDCBP\|SDF4\|SEL1L3\|SERPINE2\|SFT2D1\|SHC1\|SLC38A2\|SNAP23\|SPARC\|SURF4\|SVIL\|TJP1\|TM9SF2\|TMBIM4\|TMEFF2\|TMEM50A\|TNS1\|TPM1\|TRIL\|TUBB6 | 1.16E-17 | ** |
| NOR (nC9) | ADRN | 196 | ABCA3\|ABCB1\|ABLIM1\|ACOT7\|ACVR1B\|ADGRB3\|ADRBK2\|AGTPBP1\|AHSA1\|AKAP12\|ANK2\|ANP32A\|ARHGEF7\|ARL6IP1\|ATCAY\|BEX1\|BEX2\|BMP7\|C11orf95\|C14orf132\|C3orf14\|CADM1\|CCDC167\|CCND1\|CCNI\|CCP110\|CD200\|CENPV\|CERK\|CHGB\|CHML\|CHRNA3\|CKB\|CLASP2\|CLGN\|CRMP1\|CYFIP2\|CYGB\|DBH\|DCX\|DDX39A\|DNER\|DPYSL2\|DPYSL3\|DPYSL5\|DTD1\|EIF1B\|ELAVL3\|ELAVL4\|EML4\|EML6\|ENO2\|EPB41L4A-AS1\|FAM163A\|FAM171B\|FKBP1B\|FOXO3\|FSD1\|GABRB3\|GAP43\|GATA2\|GATA3\|GCH1\|GDAP1\|GDAP1L1\|GDI1\|GLCCI1\|GNB1\|GNG4\|GPR22\|GRIA2\|H1FX\|HAND1\|HAND2-AS1\|HMGA1\|HMP19\|HN1\|HNRNPA0\|ICA1\|IGSF3\|INA\|INSM1\|IRS2\|ISL1\|KDM1A\|KIAA1211\|KIDINS220\|KIF1A\|KIF21A\|KIF2A\|KIF5C\|KLC1\|KLF13\|KLF7\|KLHL13\|KLHL23\|L1CAM\|LRRTM2\|LSM3\|LSM4\|MAGI3\|MANEAL\|MAP1B\|MAP2\|MAP6\|MAPK8\|MAPT\|MIAT\|MMD\|MSH6\|MSI2\|MXI1\|MYEF2\|MYO5A\|NAP1L5\|NBEA\|NCAM1\|NCOA7\|NCS1\|NELFCD\|NELL2\|NMNAT2\|NNAT\|NPY\|NRCAM\|NRSN1\|NSG1\|OLA1\|PBX3\|PEG3\|PHOX2A\|PHOX2B\|PHPT1\|PHYHIPL\|PIK3R1\|PLPPR5\|PRCD\|PRPH\|QDPR\|RAB6B\|RALGDS\|RANBP1\|RBMS3\|RBP1\|REEP1\|RGS5\|RIMBP2\|RIMS3\|RNF144A\|RNF150\|RNF157\|RPS6KA2\|RTN1\|RTN2\|RUFY3\|RUNDC3A\|SATB1\|SBK1\|SCAMP5\|SCG2\|SCG3\|SCN3A\|SEPT3\|SEPT6\|SETD7\|SHD\|SLIT3\|SNAP25\|SNAP91\|SOX11\|ST3GAL6\|STMN2\|STMN4\|STXBP1\|SV2C\|SYNPO2\|SYT1\|SYT4\|TACC2\|TAGLN3\|TFAP2B\|TH\|TMEM178B\|TMOD1\|TMOD2\|TMTC4\|TSPAN7\|TTC8\|TUBB2A\|TUBB2B\|TUBB4B\|UBE2C\|ZNF512\|ZNF704\|ZNF711\|ZNF91 | <1E-50 | ** |
| NOR (nC9) | MES | 94 | ACAP2\|AEBP1\|ANXA2\|ANXA5\|APOE\|ARMCX2\|ATP1B1\|ATP2B1\|ATP8B2\|ATXN1\|B2M\|BGN\|CBFB\|CD44\|CD63\|CLIC4\|CNN3\|COL1A1\|COL3A1\|CTNNA1\|CTSB\|CYR61\|DLC1\|DMD\|DSE\|EGR1\|ENAH\|ERRFI1\|FILIP1L\|FLNA\|GNAI1\|HLA-A\|HLA-B\|HLA-C\|HSP90B1\|HSPA5\|HSPB1\|IFITM2\|IFITM3\|IGFBP5\|ITGAV\|ITGB1\|ITM2B\|JAK1\|KCTD12\|KDM5B\|KLF6\|LAMP1\|LAPTM4A\|LGALS1\|LIFR\|MANF\|MBTPS1\|MGP\|MMP2\|MOB1A\|MYADM\|NBR1\|NES\|NPC2\|NPTN\|NRP1\|OSTC\|PALLD\|PDIA3\|PDIA6\|PEA15\|PPIB\|PTBP1\|PXDN\|RBMS1\|RCN1\|RHOC\|SDF4\|SEPT10\|SERPINE2\|SHC1\|SKIL\|SLC35F5\|SLC38A2\|SPARC\|SPCS3\|SSR3\|STAT3\|THBS1\|TIMP1\|TM9SF2\|TMEM50A\|TPM1\|TRIL\|TSC22D2\|TSC22D3\|TUBB6\|VIM | 8.47E-29 | ** |
| T-cells (nC10) | MES | 8 | APOE\|B2M\|HLA-A\|HLA-B\|HLA-C\|HLA-F\|LGALS1\|STAT3 | 3.89E-02 | * |
| Undifferentiated (nC3) | MES | 196 | A2M\|ADAM19\|ADAM9\|ADAMTS5\|ADGRG6\|ANTXR1\|ANXA1\|ASPH\|ATP10D\|ATP2B1\|ATP2B4\|BMP5\|BNC2\|BTN3A2\|C4orf32\|CALU\|CAPN2\|CBLB\|CCDC80\|CD164\|CDH11\|CFH\|CFI\|COL11A1\|COL12A1\|COL4A1\|COL4A2\|COL5A2\|COL6A3\|CPED1\|CPS1\|CREB3L2\|CREG1\|CRISPLD1\|CTSC\|CYR61\|DCAF6\|DDR2\|DMD\|DNAJC1\|DNAJC10\|DNAJC3\|DPY19L1\|DUSP5\|EDEM1\|EDNRA\|EGFR\|ELF1\|ELK4\|EMP1\|EPHA3\|EPS8\|ERBIN\|ERLIN1\|ETS1\|EXT1\|F2R\|F2RL2\|FAM129A\|FAM3C\|FAM46A\|FAT1\|FBN1\|FBN2\|FILIP1L\|FKBP14\|FLRT2\|FN1\|FNDC3B\|GALNT10\|GAS2\|GNS\|GORAB\|GPX8\|HEXB\|HIPK3\|HNMT\|HOMER1\|IFI16\|IGF2R\|IL6ST\|IQGAP2\|ITGA4\|ITGAV\|ITGB1\|ITPR1\|KCNK2\|KDELC2\|KDM5B\|KIF13A\|KLF4\|LAMC1\|LAPTM4A\|LHFP\|LHX8\|LIFR\|LIPA\|LPP\|LRRC8C\|LTBP1\|LUZP1\|MAGT1\|MAML2\|MAN2A1\|MBD2\|MBNL1\|MGP\|MGST1\|MICAL2\|MXRA5\|NEK7\|NFIA\|NID1\|NID2\|NOTCH2\|NOTCH2NL\|NR3C1\|P4HA1\|PALLD\|PCOLCE2\|PCSK5\|PDE3A\|PDE7B\|PDGFC\|PDLIM1\|PEAK1\|PHLDB2\|PHTF2\|PLEKHA2\|PLEKHH2\|PLOD2\|PLPP1\|PLS3\|PLSCR1\|PLSCR4\|PLXDC2\|PON2\|POSTN\|PRCP\|PRDM6\|PROM1\|PRRX1\|PTPN14\|PTPRG\|PTPRK\|PYGL\|QKI\|RAB13\|RAB29\|RAB31\|RAP1A\|RAP1B\|REST\|RGL1\|RHOJ\|RNFT1\|ROBO1\|ROR1\|SASH1\|SCRG1\|SDCBP\|SEC14L1\|SEMA3C\|SFT2D2\|SHROOM3\|SIX4\|SLC16A4\|SLC30A7\|SLC38A6\|SMAD3\|SOSTDC1\|SPARCL1\|SPRED1\|SPRY1\|SRPX\|SSR1\|STEAP1\|STK38L\|SUCLG2\|SVIL\|SYNJ2\|TFPI\|TGFBR2\|THBS1\|TJP1\|TM4SF1\|TMEM87B\|TNC\|TOR1AIP1\|TRIM5\|UAP1\|VCL\|WLS\|WNT5A\|WWTR1\|YAP1 | <1E-50 | ** |

**Human NB to Boeva et al. 2017 (Figure 3)**

<table>
<tr><th>Cell cluster in human NB</th><th>Cell type in Boeva et al. 2017 (reference)</th><th>Number of genes significantly (FDR <0.01) upregulated in human NB overlapping genes in Boeva et al. 2017</th><th>Genes significantly (FDR <0.01) upregulated in human NB overlapping genes in Boeva et al. 2017</th><th>FDR</th><th>Significance</th></tr>
<tr><td>NOR (nC5)</td><td>Group 1 NB CRC</td><td>7</td><td>GATA3|HAND1|HAND2|ISL1|KLF7|PHOX2A|PHOX2B</td><td>5.48E-09</td><td>**</td></tr>
<tr><td>NOR (nC7)</td><td>Group 1 NB CRC</td><td>6</td><td>GATA3|HAND2|ISL1|KLF7|PHOX2A|PHOX2B</td><td>1.21E-06</td><td>**</td></tr>
<tr><td>NOR (nC8)</td><td>Group 1 NB CRC</td><td>7</td><td>GATA3|HAND1|HAND2|ISL1|KLF7|PHOX2A|PHOX2B</td><td>1.95E-08</td><td>**</td></tr>
<tr><td>NOR (nC9)</td><td>Group 1 NB CRC</td><td>7</td><td>GATA3|HAND1|HAND2|ISL1|KLF7|PHOX2A|PHOX2B</td><td>1.95E-08</td><td>**</td></tr>
<tr><td>Undifferentiated (nC3)</td><td>Group 2 NCC CRC</td><td>6</td><td>FLI1|GLIS3|NR3C1|PRRX1|RUNX1|RUNX2</td><td>1.29E-02</td><td>*</td></tr>
</table>

### Human NB to Kildisiute et al. 2021 (Figure 3)

<table>
<tr><th>Cell cluster in human NB</th><th>Cell cluster in GOSH (10X) neuroblastoma Kildisiute et al. 2021</th><th>Number of genes in the specific signature of human NB shared with GOSH markers in Kildisiute et al. 2021</th><th>Genes in the specific signature of human NB shared with GOSH markers in Kildisiute et al. 2021</th><th>FDR</th><th>Significance</th></tr>
<tr><td>Endothelial (nC4)</td><td>Mesenchyme</td><td>159</td><td>ACKR1|ACKR3|ACVRL1|ADAM15|ADAMTS1|ADAMTS4|ADAMTS9|AHNAK|ANGPTL4|ANXA2|APP|AQP1|ARL2|ATP11C|ATP1A1|BAG3|BCAR1|BCL3|BHLHE40|BMP6|CALCRL|CAV1|CBLB|CCDC85B|CD34|CD55|CD59|CD99|CDC42EP2|CDKN1A|CLIC4|CNKSR3|COL15A1|COL4A1|CRIM1|CRIP2|CSF3|DDX21|DOCK6|DUSP5|EFNB1|EGFL7|EHD2|EIF5B|ELK3|EMP1|EMP2|ENG|EPHB4|EVA1C|F2R|FAM129B|FBL|FKBP1A|FLNB|FOSL1|FOXC1|FOXO1|FUT11|FZD4|GABRE|GALNT1|GJA1|GNG11|GRB10|HAPLN3|HES1|HIPK3|HSPG2|HYAL2|ID1|ID3|IER3|IFI27|IFITM2|IFITM3|IGFBP3|IGFBP4|IGFBP7|IL33|IL6|IL6ST|INSR|ITGA5|ITPKC|JMJD1C|KCTD12|KIAA0355|KLF2|LDB2|LIMS2|LMNA|LRP10|LRRC32|LUZP1|MALL|MAN1A1|MAST4|MIDN|MYADM|MYH9|NDRG1|NFIC|NFKBIA|NPR1|PCDH17|PDLIM1|PDLIM4|PIEZO1|PKIG|PKP4|PLAT|PLEC|PLEKHG2|PLPP3|PLS3|PRNP|PXN|RAB13|RAI14|RAMP2|RASIP1|RBPMS|RCAN1|RHOC|RHOJ|S100A10|S100A13|S100A16|S100A6|SASH1|SERTAD1|SKI|SLC39A14|SLC9A3R2|SLK|SMAD3|SOCS3|SPARCL1|SPRY1|STOM|SWAP70|SYNPO|TGFBR2|THBD|TM4SF1|TMEM173|TMEM204|TMOD3|TNFRSF10B|TNFRSF10D|TNKS1BP1|TRIB1|VEGFC|VWA1|VWF|WWTR1|YBX3|ZBTB7A</td><td><1E-50</td><td>**</td></tr>
<tr><td>Endothelial (nC4)</td><td>Vascular endothelium</td><td>232</td><td>AC011526.1|ACKR1|ACKR3|ACVRL1|ADAM15|ADAMTS1|ADAMTS4|ADAMTS9|ADCY4|ADGRG1|ADGRL4|AHNAK|ANGPTL4|ANXA2|APLNR|APOLD1|APP|AQP1|ARHGEF15|ARL2|BAG3|BCAR1|BCL6B|BHLHE40|BMP2|BMP6|BST2|C2CD4B|CA2|CALCRL|CAV1|CCDC85B|CD34|CD59|CD93|CD99|CDC37|CDC42EP3|CDH5|CDK17|CDKN1A|CLEC14A|CLIC4|CNKSR3|COL15A1|COL4A1|CPLX1|CRIM1|CRIP2|CSF3|DLL4|DOCK6|DUSP5|DUSP6|ECE1|EFNA1|EFNB1|EGFL7|EHD2|EHD4|ELK3|EML3|EMP1|EMP2|ENG|EPAS1|EPHA2|EPHB4|ERG|ESAM|ETS1|ETS2|EVA1C|F2R|FAM110D|FAM129B|FKBP1A|FKBP1C|FLNB|FLT1|FLT4|FOSL1|FOXC1|FOXO1|FUT11|FZD4|FZD6|GABRE|GALNT1|GJA1|GNAI2|GNG11|GPR4|GRASP|GRB10|HAPLN3|HES1|HIPK3|HLA-E|HSPG2|HYAL2|ID1|ID3|IFI27|IFITM2|IFITM3|IGFBP3|IGFBP4|IGFBP7|IL33|IL6|IL6ST|INHBB|INSR|ITGA5|ITPKC|ITPRIP|KCNJ2|KIAA0355|KLF2|LAMA5|LDB2|LDLR|LIMS2|LMNA|LMO2|LRP10|LRRC32|LUZP1|MALL|MAN1A1|MAST4|MMRN1|MMRN2|MPZL2|MSN|MYCT1|MYH9|NDRG1|NEDD9|NFIC|NOS3|NOTCH1|NPR1|NRGN|PALMD|PCDH12|PCDH17|PDGFB|PDLIM1|PECAM1|PIEZO1|PKIG|PKP4|PLAT|PLEC|PLEKHG2|PLPP3|PLS3|PLVAP|PLXNA2|PNP|PODXL|PPM1F|PVR|PXN|RAB13|RAI14|RALGAPA2|RAMP2|RAMP3|RAPGEF3|RASIP1|RBPMS|RCAN1|RHOC|RHOJ|RNASE1|RND1|ROBO4|RPGR|S100A10|S100A13|S100A16|S100A6|S1PR1|SASH1|SCARF1|SELE|SELP|SEMA3F|SERTAD1|SHANK3|SHROOM4|SKI|SLC9A3R2|SLCO2A1|SLK|SNAI1|SNRK|SOCS2|SOCS3|SOX17|SOX18|SOX7|SPARCL1|SPRY1|SSH1|STC1|STC2|STOM|SWAP70|SYNPO|TEAD4|TGFBR2|TGM2|THBD|TIE1|TINAGL1|TM4SF1|TMEM173|TMEM204|TMEM255B|TMOD3|TNFRSF10B|TNFRSF10D|TNKS1BP1|VEGFC|VWA1|VWF|WWTR1|YBX3</td><td><1E-50</td><td>**</td></tr>
<tr><td>Endothelial (nC4)</td><td>Leukocytes</td><td>44</td><td>ADA|BAZ1A|BCL3|BHLHE40|BST2|CA2|CBLB|CCDC69|CD55|CD93|CD99|CDC42EP2|DIAPH1|DUSP5|DUSP6|EFHD2|EHD4|ETS1|FXYD5|GNAI2|GRASP|HES1|HLA-E|IFITM2|IFITM3|ITPRIP|KLF2|LRP10|MSN|MYH9|NFKBIA|NRGN|PABPC1|PDLIM1|PECAM1|S100A10|SERTAD1|SLCO4A1|SMAD3|TGFB1|TGFBR2|THBD|TMEM173|YBX3</td><td>2.80E-17</td><td>**</td></tr>
<tr><td>Endothelial (nC4)</td><td>Tumour cluster 3</td><td>87</td><td>AC011526.1|ADAM15|ADAMTS9|APOLD1|APP|ARL2|ATG9B|ATP11C|BAG5|BAZ1A|BCAR1|C2CD4B|C6orf208|CCDC85B|CD59|CDC37|CDC42EP3|CDK17|CEACAM19|CENPB|CHD7|CMIP|CNKSR3|CPLX1|DDX21|EDARADD|EFNA1|EIF5B|ENO1|EPAS1|EVA1C|FBL|FJX1|FKBP1A|FKBP1C|GALNT1|GAR1|GRPEL1|HYAL2|IFI27|IL6ST|INSR|KIAA0355|LDB2|MIDN|MRTO4|MTA2|NFIC|NIP7|PABPC1|PABPC3|PARVB|PCDH17|PDLIM4|PEX5|PIM3|PKIG|PKP4|PODXL|PRNP|PVR|RAB13|RAI14|RALGAPA2|RAMP2|RCAN1|RND1|RP11-314N13.4|RRP12|SBNO2|SEMA3F|SHB|SLC25A37|SLC35E4|SLC39A14|SLC4A7|SLC9A3R2|SLK|SNRK|SOCS2|SSH1|STC2|SWAP70|TEAD4|TMEM204|TNKS1BP1|ZBTB7A</td><td>1.28E-06</td><td>**</td></tr>
</table>

<table>
<tr><td>Endothelial (nC4)</td><td>Tumour cluster 2</td><td>72</td><td>ADAM15|ADAMTS4|ANXA2|APOLD1|APP|ARL2|ATP11C|ATP1A1|BAG5|BCAR1|C2CD4B|CD59|CDC37|CDC42EP3|CDKN1A|CENPB|CHD7|CLIC4|CMIP|CNKSR3|CRIP2|DDX21|ECE1|EDARADD|EIF5B|EMP2|ENO1|EPAS1|F2R|FAM129B|FJX1|FKBP1A|GALNT1|GRPEL1|HYAL2|IGFBP4|IL6ST|INSR|JMJD1C|KIAA0355|LAMA5|LMNA|MIDN|MRTO4|MYADM|NDRG1|NIP7|NRGN|PARVB|PDLIM4|PKIG|PKP4|PLAT|PLXNA2|PODXL|RAB13|RAMP2|RAMP3|RHOC|RNF19B|SKI|SLC35E4|SLC39A14|SLC4A7|SOCS2|SSH1|STC1|TMEM70|TMOD3|TPM3|UPP1|VWA1</td><td>3.77E-04</td><td>**</td></tr>
<tr><td>Endothelial (nC4)</td><td>Tumour cluster 1</td><td>76</td><td>ADAM15|ANXA2|APP|ARL2|ATG9B|ATP11C|ATP1A1|BAG5|BCAR1|C2CD4B|C6orf208|CD59|CDC37|CDC42EP3|CENPB|CHD7|CLIC4|CMIP|CNKSR3|CPLX1|CRIP2|ECE1|EDARADD|EIF5B|ENDOD1|ENO1|EPAS1|F2R|FAM129B|FJX1|FKBP1A|FKBP1C|GALNT1|GAR1|GRB10|GRPEL1|HYAL2|IER3|IFI27|IL6ST|INSR|JMJD1C|KIAA0355|LAMA5|LDB2|LMNA|MRTO4|MYADM|NDRG1|NIP7|PCDH17|PDLIM4|PEX5|PKIG|PKP4|PLAT|PLEC|PLXNA2|PRNP|RAMP2|RAMP3|RCAN1|RHOC|RP11-314N13.4|SHB|SLC25A37|SLC35E4|SLC39A14|SLC9A3R2|SNRK|SOCS2|STC1|TMEM70|TMOD3|UPP1|VWA1</td><td>3.35E-03</td><td>**</td></tr>
<tr><td>Macrophages (nC6)</td><td>Leukocytes</td><td>226</td><td>ACP5|ACSL1|ADAM8|ADAP2|ALOX5|AMPD3|ANPEP|APBB1IP|APOE|AQP9|ARHGAP18|ARHGAP4|ATG16L2|BCL2A1|BTK|C15orf48|C1orf162|C5AR1|CD14|CD163|CD300A|CD300LF|CD72|CD74|CD83|CD84|CD86|CDCP1|CELF2|CHST15|CIITA|CLEC7A|CPVL|CSF1R|CSF2RA|CSF3R|CTSH|CTSS|CTSZ|CXCL16|CXCL2|CXCL3|CXCL8|CYBB|CYTH4|DENND3|DOK3|DPP7|ELL|ELL2|EMILIN2|FCER1G|FCGRT|FERMT3|FGD2|FGR|FOSL2|FPR3|GAB3|GK|GLIPR2|GNA15|GPCPD1|GPR137B|GPR183|GSAP|GSTO1|HAVCR2|HCK|HCLS1|HK2|HK3|HLA-DMA|HLA-DMB|HLA-DOA|HLA-DPA1|HLA-DPB1|HLA-DQA1|HLA-DQA2|HLA-DQB1|HLA-DQB2|HLA-DRA|HLA-DRB1|HLA-DRB5|HMOX1|HPS1|HTRA4|IFI30|IFNGR1|IL18|IL2RA|IL4I1|IL6R|IRF5|IRF8|ITGAD|ITGAM|ITGAX|KCNAB2|KLHL6|KMO|LAIR1|LAPTM5|LCP2|LGALS3|LGALS9|LILRA6|LILRB1|LILRB2|LILRB3|LILRB4|LILRB5|LITAF|MAFB|MANBA|MAP3K8|MCL1|MERTK|METRNL|MGAT1|MIR181A1HG|MPEG1|MPP1|MS4A4A|MS4A6A|MS4A7|MSR1|MYO1F|NABP1|NAMPT|NBPF19|NCF2|NCF4|NEAT1|NFKBID|NPL|NR4A2|ODF3B|OLR1|OSCAR|OSM|PADI2|PARP14|PARVG|PDE4A|PFKFB3|PIK3R5|PLA2G7|PLAUR|PLCB2|PLEK|PLIN2|PLSCR1|PPIF|PREX1|PTK2B|PTPN6|RAB20|RAB31|RAP2B|RASGEF1B|RASSF2|RASSF4|RASSF5|RBM47|REL|RGS1|RGS2|RHBDF2|RIN3|RNASET2|RNF135|RNF144B|RNF149|RP11-213H15.3|RREB1|RXRA|SAMHD1|SAMSN1|SAT1|SDS|SEMA4A|SGK1|SIGLEC1|SLC11A1|SLC15A3|SLC16A3|SLC16A6|SLC17A9|SLC18B1|SLC1A3|SLC31A2|SLC37A2|SLC8B1|SLCO2B1|SMAP2|SPI1|SRGN|STK10|STXBP2|SYNGR2|TCIRG1|TFEC|TGFBI|TGIF1|THEMIS2|TIPARP|TLR2|TMEM52B|TMEM63A|TNFAIP2|TNFRSF14|TNFRSF1B|TNS3|TPP1|TREM1|TRIM38|TRPM2|TYMP|TYROBP|VAMP8|WIPI1|ZFP36|ZFP36L1|ZMYND15|ZNF331</td><td><1E-50</td><td>**</td></tr>
<tr><td>Macrophages (nC6)</td><td>Mesenchyme</td><td>86</td><td>ABCA1|ACOT9|ADAM9|ANKRD9|ANPEP|APOE|ARAP1|ARHGAP18|ARL8B|AVPI1|CD58|CELF2|CHST15|CSTB|CTSB|CTSL|CTSZ|CXCL1|CXCL2|CXCL3|CYP27A1|DAB2|ELL|ELL2|EMILIN2|FAM20A|FCGRT|FCHO2|FNIP2|FOSL2|GABARAPL1|GLIPR2|GLUL|GPCPD1|GPNMB|GRN|GSTO1|HEXA|HEXB|IFFO1|IFNGR1|LGALS3|LGMN|LRP1|MAP3K8|MCL1|METRNL|MGAT1|MGLL|NABP1|NAMPT|NBPF10|NEAT1|NR1H3|NR4A2|PAPSS2|PLSCR1|PLXDC2|PSAP|RAB31|RAB7B|RASSF2|RIPK2|RNF13|RNF135|RXRA|SAMHD1|SAT1|SDC3|SEMA4A|SGK1|SLC18B1|SLC8B1|STX4|STX7|TBC1D12|TGFBI|TGIF1|TIPARP|TNFAIP2|TNS3|TPCN1|WIPI1|ZEB2|ZFP36|ZFP36L1</td><td>2.93E-25</td><td>**</td></tr>
<tr><td>Macrophages (nC6)</td><td>Vascular endothelium</td><td>19</td><td>ARHGAP18|CTSL|CXCL1|CXCL2|DAB2|FCGRT|FCHO2|FNIP2|FOSL2|LGALS3|MCL1|MGLL|NEAT1|PARP14|PLXDC2|SAT1|SGK1|ZFP36|ZFP36L1</td><td>4.29E-04</td><td>**</td></tr>
<tr><td>MSC (nC1)</td><td>Mesenchyme</td><td>52</td><td>AC068491.1|AC116035.2|ADAM33|AEBP1|ALDH1A3|ARL2BP|AXL|C11orf96|C1R|C1S|CALD1|CD248|CFHR1|COL16A1|COL1A2|COL27A1|COL5A1|COL6A2|COL6A3|CRISPLD2|ERRFI1|F3|FGFR1|FN1|FSTL3|GEM|GFPT2|ID4|INHBA|LIMA1|LMCD1|MICALL2|MT1E|MYL9|NFATC4|NOTCH3|NR4A1|PDGFRB|PKDCC|PLEKHA4|PRRX1|PTGIR|RAB34|SERPINE1|SNAI2|SPRY2|THBS1|THBS2|TPM2|UAP1|VASN|VCAN</td><td>4.23E-22</td><td>**</td></tr>
<tr><td>MSC (nC1)</td><td>Vascular endothelium</td><td>21</td><td>AC068491.1|AC116035.2|C11orf96|CALD1|COL1A2|COL5A1|COL6A2|FGFR1|FN1|LIMA1|LMCD1|MICALL2|MT1E|MYL9|NOTCH3|NR4A1|PLEKHA4|RAB34|SERPINE1|SPRY2|TPM2</td><td>5.90E-11</td><td>**</td></tr>
</table>

| | | | | | |
|---|---|---|---|---|---|
| NOR (nC7) | Tumour cluster 1 | 287 | ACAD10\|ACAP3\|ACTG1\|ADCY1\|ADGRB2\|ADGRL1\|AGAP2\|AGAP3\|AGRN\|AHI1\|AKAP6\|AKAP9\|AMIGO2\|ANAPC2\|ANKRD12\|ANKRD13B\|ANKRD36\|ANKRD36B\|ANKRD36C\|ANKS1A\|APC2\|ARHGEF10L\|ARID1B\|ASIC3\|ASIC4\|ATAD5\|ATF6B\|ATG14\|B4GALNT4\|BICD1\|BMPR1B\|BPTF\|BRSK2\|BRWD1\|BTBD3\|C14orf132\|C1QTNF2\|C1orf226\|C2CD4A\|CALCOCO1\|CAMK2B\|CASP8AP2\|CASZ1\|CC2D1A\|CCDC136\|CCDC144NL\|CCNL2\|CDH24\|CDK10\|CDK5RAP3\|CEP162\|CEP95\|CHD5\|CHD9\|CHGA\|CIC\|CNTFR\|CORO6\|COX11\|CRLF3\|CYB561\|DAAM1\|DBH\|DBN1\|DCHS1\|DDC\|DDX17\|DDX39B\|DIRAS3\|DLC1\|DMTF1\|DNM1\|DOC2B\|DOCK4\|DPP6\|DST\|EBF1\|ECEL1\|EHMT2\|ELAVL2\|EPB41L4A\|EPHB3\|ERC2\|ERV3-1\|EXOC4\|EZH2\|FAM120B\|FAM184A\|FAM193B\|FASTK\|FNDC5\|FTX\|GABPB1-AS1\|GAS8\|GATA2\|GATA3\|GCGR\|GFRA3\|GIGYF1\|GINM1\|GLG1\|GNAS\|GNB3\|GPATCH8\|GPR137C\|GPRASP1\|GRIK2\|GSE1\|GTF2I\|GTF2IRD1\|GTF2IRD2\|GTF2IRD2B\|HECTD4\|HID1\|HIP1R\|HMX1\|HS3ST5\|ICE1\|ING4\|INSRR\|ISYNA1\|KAT8\|KCNAB3\|KCNQ1OT1\|KCNQ2\|KCNQ5\|KDM1A\|KDM5B\|KLC4\|KMT2E\|KRIT1\|LAMA4\|LINC00340\|LMBR1L\|LMO1\|LUC7L\|LUC7L2\|LUC7L3\|MAP1B\|MAPK8IP3\|MAPT\|MAST1\|MBTD1\|MC1R\|MCM7\|MDGA1\|MED23\|MEG3\|MEIS3\|MGAT4B\|MLH1\|MLXIP\|MTA1\|MTRNR2L8\|NBPF20\|NCAM1\|NCRNA00107\|NDRG4\|NDUFA4L2\|NINL\|NISCH\|NKTR\|NOP56\|NOS1AP\|NPY\|NRCAM\|NRDE2\|NTRK1\|NXPH4\|OBSL1\|OCLN\|PABPN1\|PACS2\|PAXBP1\|PCLO\|PDE9A\|PEX6\|PHC1\|PHF14\|PHIP\|PI4KA\|PIGQ\|PKD1\|PLAGL1\|PLXNA4\|PLXNB1\|PNISR\|PNN\|POLR2J2\|POU2F2\|PPIP5K2\|PRPF4B\|PRPSAP2\|PRRC2B\|PRRT2\|PSD2\|PSMG4\|PTBP2\|PTPRF\|PXDN\|R3HDM2\|RABL6\|RADIL\|RAI1\|RBBP6\|RBFOX1\|RBM33\|RBM6\|REC8\|RERE\|RGMB\|RGS7\|RHOT2\|RIMKLB\|RNF44\|RP11-349A22.5\|RPS6KL1\|RSBN1L\|RUNDC3A\|SDCCAG8\|SEMA6C\|SFPQ\|SFSWAP\|SGSM2\|SHC2\|SHPRH\|SLC18A1\|SLC25A29\|SLC38A1\|SMARCD1\|SMPD3\|SNAP91\|SOBP\|SOGA3\|SOX4\|SPG7\|SPPL2B\|SPTAN1\|SRGAP3\|SRRM1\|SRRM3\|SRRM4\|SRSF11\|STMN1\|STMN2\|STXBP1\|TAF1C\|TBC1D24\|TBKBP1\|TCERG1\|TFAP2B\|TH\|TIA1\|TLE4\|TMEM191C\|TRIM33\|TRIP11\|TSC2\|TSTD2\|TTC3\|TUBA1A\|TXNDC16\|UBAP1L\|UCHL1\|ULK1\|UNC5A\|VARS2\|VPS9D1\|WDR6\|WDR86\|WSB1\|YJEFN3\|YLPM1\|ZC3H14\|ZCCHC14\|ZFHX3\|ZFP90\|ZKSCAN1\|ZNF117\|ZNF138\|ZNF253\|ZNF292\|ZNF33A\|ZNF429\|ZNF512B\|ZNF608\|ZRANB2 | <1E-50 | ** |
| NOR (nC7) | Tumour cluster 2 | 270 | ACAD10\|ACAP3\|ACTG1\|ADCY1\|ADGRB2\|ADGRL1\|AGAP2\|AGAP3\|AGRN\|AHI1\|AKAP6\|AKAP9\|AMIGO2\|ANKRD12\|ANKRD13B\|ANKRD36\|ANKRD36B\|ANKRD36C\|ANKS1A\|APC2\|ARHGEF10L\|ARID1B\|ASIC4\|ATAD5\|ATF6B\|B4GALNT4\|BICD1\|BMPR1B\|BPTF\|BRSK2\|BRWD1\|BTBD3\|C14orf132\|C1QTNF2\|C1orf226\|CAMK2B\|CASP8AP2\|CASZ1\|CC2D1A\|CCDC136\|CCNL2\|CDH24\|CDK10\|CDK5RAP3\|CEP162\|CEP95\|CHD5\|CHD9\|CHGA\|CIC\|CNTFR\|CORO6\|COX11\|CRAMP1\|CRLF3\|CYB561\|DAAM1\|DBH\|DBN1\|DCHS1\|DDC\|DDX17\|DDX39B\|DLC1\|DMTF1\|DNM1\|DOC2B\|DOCK4\|DPP6\|DST\|EBF1\|ECEL1\|EHMT2\|ELAVL2\|EPB41L4A\|EPHB3\|ERV3-1\|EXOC4\|EZH2\|FAM120B\|FAM184A\|FAM193B\|FASTK\|FNDC5\|GABPB1-AS1\|GAS8\|GATA2\|GATA3\|GFRA3\|GIGYF1\|GINM1\|GLG1\|GNAS\|GNB3\|GPATCH8\|GPR137C\|GPRASP1\|GSE1\|GTF2I\|GTF2IRD1\|GTF2IRD2\|GTF2IRD2B\|HECTD4\|HID1\|HIP1R\|HMX1\|HS3ST5\|ICE1\|ING4\|INSRR\|ISYNA1\|KAT8\|KCNQ1OT1\|KCNQ2\|KCNQ5\|KDM1A\|KLC4\|KMT2E\|KRIT1\|LAMA4\|LINC00340\|LMBR1L\|LMO1\|LUC7L\|LUC7L2\|LUC7L3\|MAP1B\|MAPK8IP3\|MAPT\|MAST1\|MBTD1\|MC1R\|MCM7\|MDGA1\|MED23\|MEG3\|MEIS3\|MGAT4B\|MLH1\|MLXIP\|MNAT1\|MTA1\|MTRNR2L12\|MTRNR2L8\|NBPF20\|NCAM1\|NDRG4\|NDUFA4L2\|NINL\|NISCH\|NKTR\|NOP56\|NOS1AP\|NPY\|NRCAM\|NRDE2\|NTRK1\|OBSL1\|PABPN1\|PACS2\|PAXBP1\|PCLO\|PDE9A\|PHC1\|PHF14\|PHF19\|PHIP\|PI4KA\|PIGQ\|PKD1\|PLAGL1\|PLXNA4\|PLXNB1\|PNISR\|PNN\|POLR2J2\|POU2F2\|PPIP5K2\|PRPF4B\|PRPSAP2\|PRRC2B\|PRRT2\|PSD2\|PSMG4\|PTBP2\|PTPRF\|PXDN\|R3HDM2\|RABL6\|RADIL\|RAI1\|RBBP6\|RBFOX1\|RBM33\|RBM6\|REC8\|RERE\|RGMB\|RGS7\|RHOT2\|RIMKLB\|RP11-349A22.5\|RPS6KL1\|RSBN1L\|RUNDC3A\|SDCCAG8\|SEMA6C\|SFPQ\|SFSWAP\|SGSM2\|SHC2\|SHPRH\|SLC18A1\|SLC25A29\|SLC38A1\|SMARCD1\|SMPD3\|SNAP91\|SOBP\|SOGA3\|SOX4\|SPG7\|SPTAN1\|SRGAP3\|SRRM1\|SRRM3\|SRRM4\|SRSF11\|STK38\|STMN1\|STMN2\|STXBP1\|TAF1C\|TBC1D24\|TCERG1\|TFAP2B\|TH\|TIA1\|TLE4\|TMEM191C\|TRIM33\|TRIP11\|TSC2\|TSTD2\|TTC3\|TUBA1A\|TXNDC16\|UCHL1\|ULK1\|UNC5A\|VPS9D1\|WDR27\|WDR6\|WDR86\|WSB1\|YJEFN3\|YLPM1\|ZC3H14\|ZCCHC14\|ZFHX3\|ZFP90\|ZKSCAN1\|ZNF117\|ZNF138\|ZNF292\|ZNF33A\|ZNF429\|ZNF512B\|ZNF608\|ZRANB2 | <1E-50 | ** |
| NOR (nC7) | Tumour cluster 3 | 272 | ACAD10\|ACAP3\|ACTG1\|ADCY1\|ADGRB2\|ADGRL1\|AGAP3\|AGER\|AHI1\|AKAP6\|AKAP9\|AMIGO2\|ANAPC2\|ANKRD12\|ANKRD13B\|ANKRD36\|ANKRD36B\|ANKRD36C\|ANKS1A\|APC2\|ARID1B\|ASIC3\|ASIC4\|ATAD5\|ATF6B\|ATG14\|B4GALNT4\|BICD1\|BMPR1B\|BPTF\|BRSK2\|BRWD1\|BTBD3\|C14orf132\|C1orf226\|C2CD4A\|CALCOCO1\|CAMK2B\|CASP8AP2\|CC2D1A\|CCDC136\|CCDC144NL\|CDH24\|CDK10\|CDK5RAP3\|CEP162\|CEP95\|CHD9\|CHGA\|CIC\|CNTFR\|COX11\|CRAMP1\|CRLF3\|DAAM1\|DBH\|DBN1\|DCHS1\|DDC\|DDX17\|DDX39B\|DMTF1\|DNM1\|DOC2B\|DOCK4\|DST\|EHMT2\|EIF1AY\|ELAVL2\|ERV3-1\|EVL\|EXOC4\|EYA4\|EZH2\|FAM120B\|FAM184A\|FAM193B\|FASTK\|FNDC5\|FTX\|GABPB1-AS1\|GAS8\|GATA2\|GATA3\|GCGR\|GFRA3\|GIGYF1\|GINM1\|GLG1\|GNAS\|GNB3\|GPATCH8\|GPR137C\|GRIK2\|GSE1\|GTF2I\|GTF2IRD1\|GTF2IRD2\|GTF2IRD2B\|HBA1\|HECTD4\|HID1\|HIP1R\|HMX1\|ICE1\|ING4\|ISYNA1\|KAT8\|KCNAB3\|KCNQ1OT1\|KCNQ2\|KCNQ5\|KDM1A\|KDM5B\|KDM5D\|KLC4\|KMT2E\|KRIT1\|LINC00340\|LMBR1L\|LMO1\|LUC7L\|LUC7L2\|LUC7L3\|MAP1B\|MAP3K1\|MAPK8IP3\|MAPT\|MAST1\|MBTD1\|MC1R\|MCM7\|MED23\|MEG3\|MEG8\|MEIS3\|MGAT4B\|MLH1\|MLXIP\|MNAT1\|MTA1\|MTRNR2L1\|MTRNR2L12\|MTRNR2L8\|MUC20\|NBPF20\|NCAM1\|NCRNA00185\|NDRG4\|NDUFA4L2\|NINL\|NISCH\|NKTR\|NOP56\|NOS1AP\|NPIPB15\|NRDE2\|OBSL1\|OCLN\|PABPN1\|PACS2\|PAXBP1\|PCDHA9\|PCLO\|PDE9A\|PEX6\|PHC1\|PHF14\|PHIP\|PI4KA\|PKD1\|PLXNA4\|PLXNB1\|PNISR\|PNN\|POLR2J2\|POU2F2\|PPIP5K2\|PRPF4B\|PRPSAP2\|PRRC2B\|PRRT2\|PSMG4\|PTBP2\|PTPRF\|R3HDM2\|RABL6\|RADIL\|RAI1\|RBBP6\|RBCK1\|RBFOX1\|RBM33\|RBM6\|REC8\|RERE\|RGMB\|RGS7\|RHOT2\|RIMKLB\|RNF44\|RP11-349A22.5\|RPS6KL1\|RSBN1L\|RUNDC3A\|SDCCAG8\|SEMA6C\|SFPQ\|SFSWAP\|SHC2\|SHPRH\|SLC19A1\|SLC25A29\|SLC38A1\|SMARCD1\|SMPD3\|SNAP91\|SOBP\|SOGA3\|SOX4\|SPG7\|SPPL2B\|SPTAN1\|SRGAP3\|SRRM1\|SRRM3\|SRRM4\|SRSF11\|STAG3\|STMN1\|STMN2\|STXBP1\|TAF1C\|TBC1D24\|TBKBP1\|TCERG1\|TFAP2B\|TH\|TIA1\|TRIM33\|TRIP11\|TSC2\|TSTD2\|TTC3\|TUBA1A\|TXNDC16\|UBAP1L\|UCHL1\|ULK1\|UNC5A\|USP9Y\|UTY\|VARS2\|WDR27\|WDR6\|WDR86\|YJEFN3\|YLPM1\|ZC3H14\|ZCCHC14\|ZFHX3\|ZFP90\|ZFY\|ZKSCAN1\|ZNF117\|ZNF138\|ZNF253\|ZNF292\|ZNF33A\|ZNF429\|ZNF512B\|ZRANB2 | <1E-50 | ** |

<table>
<tr><th></th><th></th><th></th><th></th><th></th><th></th></tr>
<tr><td>NOR (nC7)</td><td>Mesenchyme</td><td>49</td><td>ACTG1|AHI1|AKAP9|C1QTNF2|CALCOCO1|CCNL2|CHD9|DAAM1|DBN1|DCHS1|DDX17|DLC1|DOCK4|DST|EBF1|GINM1|GLG1|GNAS|GTF2I|KCNQ1OT1|KDM5B|LAMA4|MAP1B|MEG3|MEG8|MLXIP|MNAT1|MTRNR2L1|MTRNR2L12|MTRNR2L8|NDUFA4L2|PACS2|PKD1|PLAGL1|PLXNB1|PNISR|PRRC2B|PXDN|RBBP6|RIMKLB|SOBP|SPPL2B|SPTAN1|SRSF11|TRIP11|TTC3|TUBA1A|TXNIP|WSB1</td><td>4.49E-06</td><td>**</td></tr>
<tr><td>NOR (nC7)</td><td>Vascular endothelium</td><td>14</td><td>AGRN|DCHS1|DLC1|DOCK4|EPB41L4A|LAMA4|MTRNR2L1|MTRNR2L12|MTRNR2L8|NKTR|PXDN|TAF1C|TXNIP|WSB1</td><td>3.52E-02</td><td>*</td></tr>
<tr><td>NOR (nC8)</td><td>Tumour cluster 2</td><td>175</td><td>ACLY|AKR1B1|ANKRD54|ANKRD6|ARF5|ARL16|ASB6|ASIC1|ATP1B1|ATP6V0E2|ATP6V1G1|AZIN1|BASP1|C9orf16|C9orf78|CARD19|CBX6|CDYL2|CERK|CHMP5|CHRFAM7A|CHRNA3|CLTA|CNOT7|COPS6|COX6C|CREB5|CRNDE|CS|CUTA|CXADR|CYC1|CYCS|DCLK1|DCLK3|DKK3|DPYSL4|DUSP26|EBP|EDF1|ELMO2|ENC1|EPB41L1|EYA1|FAM219A|FECH|FRMD3|FRRS1L|FSD1L|GAL|GNAQ|GNB1|GPC6|HACD3|HDAC9|HMGN3|HNRNPDL|HTR3A|IFT20|ISCA1|ISCU|ITGB8|ITM2C|JUP|KBTBD6|KHDRBS3|KIF3B|LRRC42|MAPRE2|MMD|MRFAP1L1|NDRG3|NDUFA12|NDUFB9|NEFL|NEFM|NELL1|NFIB|NGRN|NHLH2|NPTX2|NXPH1|OGFRL1|OLFM1|ONECUT2|OPRM1|PCMT1|PCSK1N|PCSK2|PDE10A|PHF1|PNMA1|POLR3F|PPP4R1|PSMA7|PSMC2|PSMD11|PTN|PTPN3|PTS|QPCT|RAB14|RAB15|RAB2A|RBM18|RGS3|RGS4|RHBDD2|RIN2|RNF10|RNF170|RPRD1A|RRAGA|RUNX1T1|RUSC2|SAR1A|SCD5|SCOC|SCP2|SCYL2|SEC61B|SERINC3|SERPINE2|SETBP1|SH3GLB2|SLC10A4|SLC25A4|SMAD9|SMU1|SNRPN|SNX18|SORCS1|SPIN1|SPTLC1|SRI|SSBP1|ST8SIA3|STMN3|STRAP|SURF4|SYBU|THRA|THSD7A|TM2D3|TMED2|TMED4|TMEFF2|TMEM163|TMEM176A|TMX2|TPI1|TRPA1|TSPAN11|TSPAN13|TUB|TUBA1B|TUBA1C|TUBB2A|TUBB4A|TUBB4B|TUBG1|TXN|UBQLN1|ULK3|VDAC3|VLDLR|VPS4B|VSNL1|WDR47|WDR61|WRB|WSB2|YWHAZ|ZBTB18|ZNF770</td><td><1E-50</td><td>**</td></tr>
<tr><td>NOR (nC8)</td><td>Tumour cluster 1</td><td>181</td><td>ACLY|AKR1B1|ANKRD54|ANKRD6|ARF5|ARL16|ASB6|ASIC1|ATP1B1|ATP6V0E2|ATP6V1G1|AZIN1|BASP1|C11orf87|C9orf16|C9orf78|CARD19|CBX6|CCDC68|CDYL2|CERK|CHMP5|CHRFAM7A|CHRNA3|CLTA|CNOT7|COPS6|COX6C|CRABP1|CREB5|CRNDE|CS|CUTA|CXADR|CYC1|CYCS|DACH1|DCLK1|DCLK3|DKK3|DPYSL4|DUSP26|EBP|EDF1|ELMO2|EPB41L1|EPB41L4B|EYA1|FAM219A|FECH|FRMD3|FRRS1L|FSD1L|GAL|GNAQ|GNB1|GPC6|HACD3|HDAC9|HMGN3|HNRNPDL|HTR3A|IFT20|ISCA1|ISCU|ITGB8|ITM2C|JUP|KBTBD6|KHDRBS3|KIF3B|KLHL35|LRRC42|MAPRE2|MMD|MRFAP1L1|NDRG3|NDUFA12|NDUFB9|NEFL|NEFM|NELL1|NFIB|NGRN|NPTX2|NXPH1|OGFRL1|OLFM1|ONECUT2|OPRM1|PCMT1|PCSK1N|PCSK2|PDE10A|PDZD8|PHF1|PNMA1|POLR3F|PPP4R1|PSMA7|PSMC2|PSMD11|PTN|PTPN3|PTS|QPCT|RAB14|RAB15|RAB2A|RBM18|RGS3|RGS4|RHBDD2|RIN2|RNF10|RNF170|RPRD1A|RRAGA|RUNX1T1|RUSC2|SAR1A|SCD5|SCOC|SCP2|SCYL2|SEC61B|SERINC3|SERPINE2|SETBP1|SH3GLB2|SLC10A4|SLC25A4|SMAD9|SMU1|SNRPN|SNX18|SORCS1|SPIN1|SPTLC1|SRI|SSBP1|ST8SIA3|STMN3|STRAP|SURF4|SV2B|SYBU|THRA|THSD7A|TM2D3|TMBIM4|TMED2|TMED4|TMEFF2|TMEM163|TMEM176A|TMX2|TPI1|TRPA1|TSPAN11|TSPAN13|TUB|TUBA1B|TUBA1C|TUBB2A|TUBB4A|TUBB4B|TUBG1|TXN|UBQLN1|ULK3|VDAC3|VLDLR|VPS4B|VSNL1|WDR47|WDR61|WRB|WSB2|YWHAZ|ZNF770</td><td><1E-50</td><td>**</td></tr>
<tr><td>NOR (nC8)</td><td>Tumour cluster 3</td><td>142</td><td>ACLY|AKR1B1|ANKRD54|ARF5|ARL16|ASB6|ATP1B1|ATP6V0E2|ATP6V1G1|AZIN1|BASP1|C9orf78|CAT|CBX6|CCDC68|CERK|CHMP5|CHRNA3|CLTA|CNOT7|COPS6|COX6C|CREB5|CRNDE|CS|CUTA|CXADR|CYC1|CYCS|DACH1|DCLK1|DCLK3|DLK1|DPYSL4|DUSP26|EBP|EDF1|ELMO2|EPB41L1|EYA1|FAM219A|FECH|FRMD3|FRRS1L|FSD1L|GAL|GNAQ|HACD3|HDAC9|HMGN3|HNRNPDL|IFT20|ISCA1|ISCU|ITGB8|KBTBD6|KHDRBS3|KIF3B|KIT|KLHL35|LRRC42|MAPRE2|MMD|MRFAP1L1|NDRG3|NDUFA12|NDUFB9|NEFL|NEFM|NELL1|NFIB|NGRN|NXPH1|ONECUT2|PCMT1|PCSK1N|PCSK2|PDE10A|PDZD8|PNMA1|POLR3F|PPP4R1|PSMA7|PSMC2|PTS|RAB2A|RBM18|RGS4|RHBDD2|RNF10|RNF170|RPRD1A|RRAGA|RUNX1T1|RUSC2|SCD5|SCOC|SCYL2|SEC61B|SERINC3|SERPINE2|SETBP1|SLC10A4|SLC25A4|SMAD9|SMU1|SNRPN|SNX18|SPIN1|SRI|SSBP1|ST8SIA3|STMN3|STRAP|SURF4|SYBU|THRA|THSD7A|TM2D3|TMED2|TMED4|TMEFF2|TMEM163|TMEM176A|TMX2|TPI1|TSPAN11|TSPAN13|TUB|TUBA1B|TUBB2A|TUBB4A|TXN|UBQLN1|ULK3|VDAC3|VLDLR|VPS4B|WDR61|WRB|WSB2|ZNF770</td><td><1E-50</td><td>**</td></tr>
<tr><td>NOR (nC8)</td><td>Mesenchyme</td><td>57</td><td>ANKRD6|BASP1|CAT|CBX6|CCDC28A|CHMP5|CLTA|COX6C|CREB5|CUTA|DCLK1|DKK3|DLK1|EBF3|EDF1|ENC1|GNAQ|GNB1|GPC6|HMGN3|IFT20|ISCA1|ISCU|ITGB8|LRRC42|NFIB|PDE10A|PSMA7|PTS|RAB14|RAB2A|RIN2|RRAGA|RUNX1T1|RUSC2|SAR1A|SCD5|SCP2|SEC61B|SERINC3|SERPINE2|SETBP1|SMAD9|SNX18|SPIN1|SRI|SURF4|TMBIM4|TMED2|TPI1|TUBA1B|TUBA1C|TUBB4B|TXN|VLDLR|WRB|YWHAZ</td><td>3.49E-21</td><td>**</td></tr>
</table>

| | | | | | |
|---|---|---|---|---|---|
| NOR (nC9) | Tumour cluster 2 | 254 | AASDHPPT\|ABHD2\|ACVR2B\|ADD2\|AFAP1\|ALCAM\|ALKBH7\|ANKRD17\|ARHGAP35\|ARL6IP5\|ASTN1\|ATF2\|ATP11A\|ATP2C1\|ATRX\|BCL2\|BEX2\|BRINP2\|BRK1\|BSG\|C19orf12\|C1orf21\|C4orf33\|CAMK4\|CAMLG\|CAMSAP2\|CANX\|CCDC82\|CD47\|CENPJ\|CHGB\|CHN1\|CHST1\|CKS1B\|CLCN3\|CLGN\|CLIP3\|CNRIP1\|CNTN1\|COMMD1\|COPE\|COX6B1\|COX8A\|CPE\|CPNE8\|CREG2\|CSRNP3\|CXXC4\|CYGB\|DCX\|DDAH1\|DDX6\|DLEU2\|DPY30\|DPYSL3\|DSEL\|DUSP8\|DUT\|EEF1B2\|EEF2\|EHBP1\|EIF1B\|EIF2AK2\|EIF3L\|EPM2AIP1\|EPRS\|ESD\|FAM171A1\|FANCL\|FBXO30\|FIGN\|FMN1\|FOXP1\|FSTL5\|FXR1\|GAB2\|GABARAPL2\|GCSH\|GIGYF2\|GNAO1\|GPR22\|GSK3B\|GSTP1\|H1F0\|H2AFZ\|HADHA\|HDGF\|HDLBP\|HIBCH\|HINT1\|HIST1H4C\|HLTF\|HMGA1\|HMGB1\|HMGB2\|HNRNPA3\|HNRNPL\|HSD17B11\|HSD17B12\|HTATSF1\|INO80D\|INPP5F\|IRS2\|JARID2\|JPH3\|JTB\|JUN\|KCNG1\|KCNK3\|KCNQ3\|KCTD16\|KIAA1549L\|KIDINS220\|KIF26A\|KIF26B\|KIF2A\|KLHL23\|LANCL1\|LDHB\|LIFR\|LONRF2\|LRRN3\|LRRTM2\|MAP4\|MBNL2\|MEIS2\|METTL9\|MOB1B\|MST1\|MTHFD2\|MTMR6\|MXRA7\|N4BP2\|NAALAD2\|NCK2\|NDUFA1\|NDUFB11\|NDUFS1\|NMNAT2\|NRP1\|OPTN\|OXCT1\|PARD3\|PBRM1\|PCDH7\|PCDHB10\|PCDHB16\|PCDHB6\|PCDHB9\|PCNP\|PFN2\|PGAP1\|PGM2L1\|PHYHIPL\|PITPNB\|PJA2\|PKIB\|PLCXD3\|PMP22\|PODXL2\|POLR1D\|PPP2R1A\|PRDX2\|PREPL\|PROX1\|RAB11B\|RAB3C\|RAMP1\|RBMS1\|RBMS3\|RGS5\|RMND5A\|RND3\|RNF144A\|RPSA\|RSF1\|RSL24D1\|RUFY3\|SAE1\|SCARB2\|SCG2\|SCG3\|SCG5\|SCN7A\|SCRN1\|SEC62\|SEMA6D\|SEPT11\|SEPT6\|SESTD1\|SETD2\|SEZ6\|SEZ6L\|SF3B2\|SFRP1\|SLC1A4\|SLC25A12\|SLC39A10\|SLC6A2\|SLIT3\|SMARCA1\|SMARCA5\|SMARCC1\|SMC1A\|SNRNP27\|SNRPD2\|SPATS2L\|SSB\|ST3GAL6\|ST6GALNAC5\|STAC\|STMN4\|SUMO1\|SV2C\|SYNPO2\|SYP\|SYT1\|SYT13\|SYT4\|TAGLN3\|TARS\|TBCB\|TCF12\|TEAD1\|TET3\|TFDP2\|TIAL1\|TKT\|TMA7\|TMCO3\|TMEM255A\|TMOD2\|TMSB15A\|TNRC6B\|TOMM22\|TOP2B\|TRIM13\|TSHZ2\|TSPAN7\|TTBK2\|U2SURP\|UBE2E3\|UQCRC1\|VAT1L\|VPS36\|WDR82\|XRCC5\|XRN1\|YBX1\|ZBTB38\|ZC3H13\|ZC3H15\|ZDBF2\|ZNF148 | <1E-50 | ** |
| NOR (nC9) | Tumour cluster 1 | 256 | AASDHPPT\|ACVR2B\|ADD2\|AFAP1\|AL022344.4\|ALCAM\|ALKBH7\|ANKRD17\|ARHGAP35\|ARL6IP5\|ASTN1\|ATF2\|ATP11A\|ATP2C1\|ATRX\|BCL2\|BEX2\|BRINP2\|BRK1\|BSG\|BTF3\|C19orf12\|C1orf21\|C4orf33\|CAMK2N1\|CAMK4\|CAMLG\|CAMSAP2\|CANX\|CCDC186\|CCDC82\|CD47\|CENPJ\|CHGB\|CHN1\|CHST1\|CKS1B\|CLCN3\|CLGN\|CLIP3\|CNRIP1\|CNTN1\|CNTNAP4\|COMMD1\|COPE\|COX7C\|COX8A\|CPE\|CPNE8\|CREG2\|CSRNP3\|CXXC4\|CYGB\|DCX\|DDAH1\|DDX6\|DLEU2\|DPY30\|DPYSL3\|DSEL\|DUSP8\|DUT\|EEF2\|EHBP1\|EIF1B\|EIF2AK2\|EIF3L\|EML6\|EPM2AIP1\|EPRS\|ESD\|FAM171A1\|FANCL\|FASTKD1\|FBXO30\|FIGN\|FMN1\|FOXP1\|FSTL5\|FXR1\|GAB2\|GABARAPL2\|GCSH\|GIGYF2\|GNAO1\|GPR22\|GSK3B\|GSTP1\|H1F0\|H2AFZ\|HADHA\|HDGF\|HDLBP\|HIBCH\|HINT1\|HIST1H4C\|HLTF\|HMGA1\|HMGB1\|HNRNPA3\|HSD17B11\|HSD17B12\|HTATSF1\|INO80D\|INPP5F\|IRS2\|ITGA1\|JARID2\|JPH3\|JTB\|JUN\|KCNG1\|KCNK3\|KCNQ3\|KCTD16\|KIAA1549L\|KIDINS220\|KIF26A\|KIF26B\|KIF2A\|KLHL23\|LANCL1\|LDHB\|LIFR\|LONRF2\|LRRN3\|LRRTM2\|MAP4\|MBNL2\|MEIS2\|METTL9\|MOB1B\|MST1\|MTHFD2\|MTMR6\|MXRA7\|N4BP2\|NAALAD2\|NCK2\|NDUFA1\|NDUFB11\|NDUFS1\|NMNAT2\|NRP1\|NTNG1\|OPTN\|OST4\|OXCT1\|PARD3\|PBRM1\|PCDH10\|PCDH7\|PCDHB10\|PCDHB16\|PCDHB6\|PCDHB9\|PCNP\|PFN2\|PGAP1\|PGM2L1\|PHYHIPL\|PITPNB\|PJA2\|PKIB\|PLCXD3\|PMP22\|PODXL2\|POLR1D\|PPP2R1A\|PRDX2\|PREPL\|PROX1\|RAB11B\|RAB3C\|RAMP1\|RBMS1\|RBMS3\|RGS5\|RMND5A\|RND3\|RNF144A\|RSF1\|RSL24D1\|RUFY3\|SAE1\|SCARB2\|SCG2\|SCG3\|SCG5\|SCN7A\|SCRN1\|SEC62\|SEMA6D\|SEPT11\|SEPT6\|SESTD1\|SETD2\|SEZ6\|SEZ6L\|SF3B2\|SFRP1\|SLC1A4\|SLC25A12\|SLC39A10\|SLC6A2\|SLIT3\|SMARCA1\|SMARCA5\|SMARCC1\|SMC1A\|SNRNP27\|SNRPD2\|SPATS2L\|SSB\|ST3GAL6\|ST6GALNAC5\|STAC\|STMN4\|SUMO1\|SV2C\|SYNPO2\|SYP\|SYT1\|SYT13\|SYT4\|TAGLN3\|TARS\|TBCB\|TCF12\|TEAD1\|TET3\|TIAL1\|TMCO3\|TMEM255A\|TMOD2\|TMSB15A\|TNRC6B\|TOMM22\|TOP2B\|TRIM13\|TSHZ2\|TSPAN7\|TTBK2\|U2SURP\|UBE2E3\|UQCRC1\|VAT1L\|VPS36\|WDR82\|XRCC5\|YBX1\|ZBTB38\|ZC3H13\|ZC3H15\|ZDBF2\|ZNF148 | <1E-50 | ** |
| NOR (nC9) | Tumour cluster 3 | 225 | AASDHPPT\|ACVR2B\|ADD2\|AFAP1\|ALCAM\|ALKBH7\|ANKRD17\|ARHGAP35\|ASTN1\|ATF2\|ATP2C1\|ATRX\|BCL2\|BEX2\|BMP7\|BRINP2\|BRK1\|BSG\|BTF3\|C19orf12\|C1orf21\|CAMLG\|CAMSAP2\|CANX\|CENPJ\|CHGB\|CHN1\|CKS1B\|CLCN3\|CLGN\|CLIP3\|CNRIP1\|CNTNAP4\|COMMD1\|COPE\|COX6B1\|COX7C\|COX8A\|CPE\|CPNE8\|CSRNP3\|CXXC4\|CYGB\|DDAH1\|DDX6\|DLEU2\|DPY30\|DUSP8\|DUT\|EEF1B2\|EEF2\|EHBP1\|EIF1B\|EIF2AK2\|EIF3L\|EML6\|EPM2AIP1\|EPRS\|ESD\|FAM171A1\|FANCL\|FASTKD1\|FIGN\|FOXP1\|FXR1\|GAB2\|GABARAPL2\|GCSH\|GIGYF2\|GNAO1\|GPR22\|GSK3B\|GSTP1\|H1F0\|H2AFZ\|HADHA\|HDGF\|HDLBP\|HIBCH\|HINT1\|HIST1H4C\|HIST1H4E\|HLTF\|HMGA1\|HMGB1\|HMGB2\|HNRNPA3\|HNRNPL\|HSD17B11\|HSD17B12\|HTATSF1\|IGFBP5\|INO80D\|IRS2\|ITGA1\|JARID2\|JPH3\|JTB\|JUN\|KCNG1\|KCNK3\|KCTD16\|KIAA1549L\|KIDINS220\|KIF26A\|KIF26B\|KIF2A\|KLHL23\|LANCL1\|LDHB\|LIFR\|LONRF2\|LRRN3\|LRRTM2\|MAP4\|MEIS2\|METTL9\|MOB1B\|MTHFD2\|MXRA7\|N4BP2\|NAALAD2\|NCK2\|NDUFB11\|NDUFS1\|NMNAT2\|NTNG1\|OST4\|OXCT1\|PARD3\|PBRM1\|PCDH7\|PCDHB9\|PCNP\|PFN2\|PGAP1\|PGM2L1\|PHYHIPL\|PITPNB\|PJA2\|PMP22\|PODXL2\|POLR1D\|PPP2R1A\|PRDX2\|PREPL\|PROX1\|RAB11B\|RAB3C\|RAMP1\|RBMS1\|RBMS3\|RMND5A\|RNF144A\|RPL18\|RPSA\|RSF1\|RSL24D1\|RUFY3\|SAE1\|SCARB2\|SCG2\|SCG3\|SCG5\|SCRN1\|SEC62\|SEMA6D\|SEPT11\|SEPT6\|SESTD1\|SETD2\|SEZ6L\|SF3B2\|SFRP1\|SLC25A12\|SLC39A10\|SLC6A2\|SLIT3\|SMARCA1\|SMARCA5\|SMARCC1\|SMC1A\|SNRNP27\|SNRPD2\|SSB\|ST3GAL6\|ST6GALNAC5\|STAC\|STMN4\|SUMO1\|SYP\|SYT1\|SYT4\|TAGLN3\|TARS\|TBCB\|TCF12\|TEAD1\|TET3\|TFDP2\|TKT\|TLE3\|TMCO3\|TMOD2\|TMSB15A\|TNRC6B\|TOMM22\|TOP2B\|TRIM13\|TSHZ2\|TSPAN7\|TTBK2\|U2SURP\|UBE2E3\|UQCRC1\|VPS36\|WDR82\|XRCC5\|XRN1\|YBX1\|ZBTB38\|ZC3H13\|ZC3H15\|ZDBF2\|ZNF148 | <1E-50 | ** |
| NOR (nC9) | Mesenchyme | 81 | ABHD2\|AFAP1\|ALKBH7\|ANKRD17\|ARL6IP5\|ATP11A\|BRK1\|BSG\|BTF3\|C1orf21\|CAMK2N1\|CAMLG\|CAMSAP2\|CANX\|CCDC186\|CKS1B\|CNTN1\|COPE\|CPE\|CYGB\|DPY30\|DPYSL3\|DSEL\|DUT\|EEF1B2\|EEF2\|ESD\|FIGN\|FMO1\|FOXP1\|GABARAPL2\|GCSH\|GSTP1\|H1F0\|HDLBP\|HNRNPL\|HSD17B12\|IGFBP5\|ITGA1\|JTB\|JUN\|LIFR\|LONRF2\|MAP4\|MBNL2\|MEIS2\|METTL9\|MXRA7\|NDUFA1\|NRP1\|OPTN\|OST4\|PARD3\|PCDH7\|PCNP\|PFN2\|PMP22\|PRDX2\|RBMS1\|RBMS3\|RND3\|RNF144A\|SCARB2\|SCN7A\|SCRN1\|SEC62\|SEPT11\|SFRP1\|SLIT3\|SMARCA1\|SNRPD2\|SPATS2L\|SYNPO2\|TCF12\|TEAD1\|TIAL1\|TMCO3\|TNRC6B\|TOP2B\|TSHZ2\|WDR82 | 1.48E-24 | ** |
| T-cells (nC10) | Leukocytes | 14 | CD2\|CD69\|ICOS\|IGKC\|IGKV1-12\|IGLC2\|IGLC3\|IGLC7\|IGLL1\|IGLV2-14\|LTB\|TRAC\|TRBC1\|TRBC2 | 5.54E-06 | ** |

| | | | | | |
|---|---|---|---|---|---|
| Undifferentiated (nC3) | Tumour cluster 3 | 323 | ABCA12\|ABCB5\|ABCC4\|AC009264.1\|AC013400.2\|AC092594.1\|ACSS3\|ACTR3B\|ACTR3C\|ADAM29\|ADAMTS20\|ADGRL2\|AK5\|AL449209.1\|ALDH1L2\|ALG10B\|ALG14\|ANAPC1\|ANKEF1\|ANKRD20A3\|ANKRD31\|ANKRD45\|ANKS1B\|ANO3\|ANO5\|AP5M1\|ARHGAP28\|ARHGEF12\|ARHGEF28\|ARMC10\|ASAH2\|ASPM\|ATAD2\|ATRNL1\|BBOF1\|BMS1\|BRCA1\|BRIP1\|BRMS1L\|C12orf45\|C12orf48\|C14orf145\|C18orf25\|C1orf112\|C2CD3\|C8orf34\|C9orf85\|CACNB4\|CBWD5\|CC2D2A\|CCDC122\|CCDC150\|CCDC30\|CCSER1\|CDC27\|CDH18\|CDON\|CEP112\|CEP152\|CEPT1\|CFAP54\|CFAP61\|CHEK2\|CHPT1\|CISD2\|CLN5\|CLSPN\|CMC2\|CNTNAP5\|COG6\|COL11A1\|COL9A1\|CPS1\|CPSF2\|CPT2\|CRB1\|CSRP2\|CTC-228N24.3\|CTD-2230D16.1\|CYP3A5\|DBT\|DCBLD1\|DCHS2\|DDX20\|DEPDC1\|DET1\|DGKE\|DGKI\|DIAPH3\|DLEU1\|DLGAP1\|DLX6-AS1\|DNAH10\|DNAH12\|DNAH14\|DNAH6\|DNAH9\|DOCK7\|DPY19L3\|DR1\|DSG3\|E2F6\|ELOVL7\|ELP4\|EML1\|ENTPD5\|EPM2A\|ERBB4\|ERVW-1\|ESCO2\|ESRRG\|EXOC5\|EXPH5\|FAM114A2\|FAM135A\|FAM135B\|FAM13A\|FAM153B\|FAM227A\|FANCD2\|FANCI\|FANCM\|FAT3\|FBXO3\|FGF7\|FGGY\|FSIP2\|FUT9\|G2E3\|GABRA2\|GABRG3\|GALK2\|GEMIN5\|GEN1\|GK5\|GLIDR\|GNAI3\|GPAM\|GREB1L\|GRID2\|GRIP1\|GRM5\|GTPBP10\|GUCY1A2\|GULP1\|HEATR4\|HGF\|HSD17B4\|HYDIN\|INTS2\|INTS6\|INTS7\|INTU\|IPO11\|KAT7\|KDM7A\|KIAA0586\|KIAA1257\|KIAA1328\|KIF14\|KIF6\|KNTC1\|L2HGDH\|LEPR\|LMBR1\|LMO3\|LMO7\|LRP1B\|LRRC7\|LRRIQ3\|LRRK1\|MACROD2\|MCCC2\|MDN1\|MED12L\|MED17\|MGAM\|MIPOL1\|MIR99AHG\|MKLN1\|MMS22L\|MROH8\|MTHFD2L\|MYO16\|MYO5B\|NAALADL2\|NCAPG2\|NEB\|NEDD1\|NEK5\|NF1\|NIPAL3\|NLGN1\|NNT\|NOX4\|NPAS3\|NT5DC3\|NTRK2\|NUBPL\|NUP133\|NUP155\|NUP37\|OSBPL1A\|PARP11\|PART1\|PCAT1\|PCCA\|PCDHB1\|PDE4DIP\|PEX5L\|PGGT1B\|PHTF2\|PIAS2\|PIGK\|PIGN\|PIWIL1\|PIWIL2\|PLCE1\|POLQ\|POLR3B\|POLR3G\|POU2F1\|POU5F2\|POU6F2\|PPARGC1A\|PPAT\|PRDM5\|PRIM2\|PRKAA2\|PRTG\|PTCD2\|PTPN20\|RANBP17\|RBBP8\|RGS22\|RNASEH2B\|RP11-373D17.1\|RP11-3B12.1\|RP11-458B24.2\|RP11-550E22.3\|RP11-58E21.3\|RP11-735B13.1\|RPAP2\|RPS6KA5\|RPS6KB1\|SAR1B\|SCIN\|SCN11A\|SCO1\|SEC22B\|SEC23A\|SH3TC2\|SHQ1\|SLC12A2\|SLC1A2\|SLC2A13\|SLC35A3\|SLC35G1\|SLC38A9\|SLC44A5\|SLC9B1\|SLCO1A2\|SLMAP\|SMC6\|SNAP25-AS1\|SOCS7\|SOX5\|SPATA17\|SPINK5\|SRGAP2B\|SRGAP2C\|SSX2IP\|STK31\|SYNE1\|TAF4B\|TAOK1\|TBC1D8B\|TCEANC2\|TF\|TMEM132B\|TMEM213\|TMEM232\|TPH1\|TPTE2\|TRDMT1\|TRPM3\|TRPM6\|TTC26\|TTC5\|TTLL5\|TYW5\|USP32\|USP37\|USP54\|UTP20\|VWA8\|WDHD1\|WDR3\|WDR35\|WDR66\|WNT2B\|WRN\|XPNPEP3\|ZBED3-AS1\|ZDHHC13\|ZFAND4\|ZGRF1\|ZNF169\|ZNF277\|ZNF285\|ZNF426\|ZNF479\|ZNF519\|ZNF527\|ZNF546\|ZNF578\|ZNF595\|ZNF678\|ZNF679\|ZNF716\|ZNF720\|ZNF736\|ZRANB3 | 2.87E-36 | ** |
| Undifferentiated (nC3) | Mesenchyme | 155 | AASS\|ABCA10\|ABCA6\|ABCA8\|ABCA9\|ABCC9\|AC074093.1\|ACSS3\|ADAMTS12\|ADAMTSL3\|ADGRL2\|ALDH1L2\|ANO4\|ANTXR2\|AOX1\|ARHGAP24\|ARHGAP28\|ARHGAP42\|ARHGEF12\|ASPA\|ATL3\|B3GLCT\|BCO2\|BMS1\|CACHD1\|CACNB4\|CADPS2\|CD109\|CDC14B\|CDH13\|CDK14\|CDON\|CEP112\|CFAP69\|CLDN11\|CLN5\|CMYA5\|CNTN4\|CNTN6\|COL14A1\|COL21A1\|COL4A4\|COL4A5\|COL6A6\|COLGALT2\|CRISPLD1\|CSRP2\|CTD-2536I1.1\|DIO2\|DOCK9\|DPY19L3\|DSC3\|DYNC2H1\|ELK4\|EML1\|ERCC6\|ETV1\|EXOC5\|EXPH5\|FAM13A\|FAM153A\|FAM153B\|FBN2\|FGF7\|FLRT2\|FNDC1\|FOXP2\|FREM1\|FRK\|GLI3\|GUCY1A2\|GULP1\|HGF\|HHIP\|HMCN1\|IDE\|IGFN1\|INTS6\|KIF13A\|L3MBTL3\|LAMA2\|LEPR\|LGI1\|LGR4\|LGR5\|LINC00960\|LMO7\|LNPEP\|LRRK1\|LRRK2\|LTBP1\|MATN3\|MEGF10\|METTL15\|MGST1\|MIR99AHG\|MKLN1\|MR1\|MUC12\|NAALADL2\|NAV3\|NF1\|NLGN1\|NOX4\|NPHP1\|NSF\|NT5DC3\|NTRK2\|OSBPL10\|OSBPL1A\|PCSK5\|PDE1A\|PDE4DIP\|PDZD2\|PGM5\|PGM5P4-AS1\|PHLPP2\|PIEZO2\|PLA2G4A\|PLA2R1\|PLCE1\|PRDM5\|PRKD1\|PRRC1\|PTPN13\|PTPRG\|RNF217\|RP11-143M1.2\|RP11-327J17.3\|RXFP2\|SAR1B\|SEC22B\|SEC23A\|SEMA3D\|SEMA5A\|SHQ1\|SLC12A2\|SLC35G1\|SLC4A4\|SLCO6A1\|SLMAP\|SPTLC3\|SRGAP2B\|SRGAP2C\|STEAP4\|STK3\|STXBP6\|SUMF1\|SVEP1\|TBC1D8B\|THRB\|TOR1AIP1\|TPD52L1\|VGLL3\|WNT2B | 3.13E-22 | ** |
| Undifferentiated (nC3) | Tumour cluster 1 | 291 | ACTR3B\|ADCY2\|ADGRL2\|AFF2\|AGBL3\|AK5\|AKAP7\|AL449209.1\|ALDH1L2\|ALG10B\|ALG14\|ANAPC1\|ANKEF1\|ANKRD20A3\|ANKRD30BL\|ANKRD31\|ANKRD7\|ANKS1B\|ANO4\|ANO5\|AP5M1\|ARHGAP28\|ARHGEF12\|ARHGEF28\|ARMC10\|ASAH2\|ATRNL1\|BBOF1\|BMS1\|BRMS1L\|C12orf45\|C2CD3\|C5\|C8orf34\|C9orf85\|CACHD1\|CACNB4\|CBWD5\|CC2D2A\|CCDC122\|CCDC169\|CCDC178\|CDC27\|CDH10\|CDH13\|CDH18\|CDH7\|CDH9\|CDK14\|CDON\|CEPT1\|CFAP43\|CFAP61\|CFAP70\|CHPT1\|CISD2\|CLN5\|CLSPN\|CMC2\|CNTN3\|CNTN5\|CNTNAP3\|CNTNAP5\|COG6\|COL28A1\|CORIN\|CPD\|CPS1\|CPT2\|CRISPLD1\|CSMD1\|CSRP2\|CTC-228N24.3\|CTNNA3\|DAB1\|DAW1\|DBT\|DCBLD1\|DDX20\|DEPDC5\|DET1\|DGKE\|DGKI\|DLGAP1\|DNAH5\|DNAH6\|DOCK7\|DR1\|DYNC1I1\|DYNC2H1\|E2F6\|EFCAB5\|EML1\|ENPP1\|ENPP2\|EPHA6\|EPM2A\|ERBB4\|ESRRG\|ETV1\|EXOC5\|EXOG\|FAM114A2\|FAM135A\|FAM135B\|FAM227A\|FAT3\|FBXO15\|FBXO3\|FGF7\|FLRT2\|FRAS1\|FREM1\|FUT9\|G2E3\|GABRA1\|GABRA2\|GABRG2\|GALK2\|GDA\|GEMIN5\|GLIDR\|GNAI3\|GPAM\|GREB1L\|GRID2\|GRIN2A\|GRIP1\|GTPBP10\|HCN1\|HGF\|HSD17B4\|HULC\|HYDIN\|INTS2\|INTS6\|INTS7\|INTU\|IQUB\|ITGA8\|KAT7\|KIAA0586\|KIAA0825\|KIAA1328\|KIF13A\|KLHL3\|L2HGDH\|L3MBTL3\|LAMA1\|LAMA3\|LINC00467\|LINC01138\|LMBR1\|LRP1B\|LRRC7\|LRRIQ3\|MAP2K6\|MATN3\|MDGA2\|MDN1\|MED17\|MKLN1\|MMS22L\|MROH8\|MTHFD2L\|MUC12\|MYH15\|NAV3\|NCAPG2\|NECAB1\|NEK5\|NF1\|NLGN1\|NNT\|NPAS3\|NPHP1\|NRXN3\|NSF\|NUBPL\|NUP133\|NUP155\|NUP37\|NUP93\|OLMALINC\|OSBPL1A\|PART1\|PCCA\|PCDH15\|PDE4DIP\|PDZD2\|PGGT1B\|PHLPP1\|PHLPP2\|PHTF2\|PIAS2\|PIEZO2\|PIGK\|PKHD1\|PLCE1\|PLD5\|POLR3B\|POU2F1\|PPARGC1A\|PPAT\|PRKAA2\|PRRC1\|PRTG\|PTCD2\|PTGFR\|PTPRG\|PTPRT\|RANBP17\|RBBP8\|RELN\|RGPD5\|RGS22\|RNASEH2B\|RNF217\|RORB\|RP11-195E11.2\|RP11-21F16.1\|RP11-3B12.1\|RP11-550E22.3\|RP11-82L2.1\|RP5-916L7.1\|RPAP2\|RPS6KA5\|RPS6KB1\|RTTN\|SAR1B\|SCO1\|SEC22B\|SEC23A\|SEMA3A\|SEMA3D\|SEMA3E\|SEMA5A\|SHC4\|SLC12A1\|SLC12A2\|SLC1A2\|SLC35F4\|SLC38A9\|SLC44A5\|SLC9B1\|SMC6\|SNAP25-AS1\|SOCS7\|SOX5\|SPATA17\|SPHKAP\|SSX2IP\|STK31\|STRIP2\|SYNE1\|TAF4B\|TAOK1\|TCEANC2\|TEX14\|THSD7B\|TMEM232\|TRHDE\|TRPM3\|TSIX\|TTC26\|TTC5\|TTLL5\|TYW5\|USP32\|USP37\|VPS13D\|WDHD1\|WDR3\|WDR35\|WDR7\|WNT2B\|WRN\|XKR4\|XPNPEP3\|ZBED3-AS1\|ZDHHC13\|ZNF277\|ZNF285\|ZNF385B\|ZNF426\|ZNF519\|ZNF527\|ZNF678\|ZNF720\|ZNF736\|ZRANB3 | 7.34E-21 | ** |

<table>
<tr><td>Undifferentiated (nC3)</td><td>Tumour cluster 2</td><td>233</td><td>ACTR3B|ADGRL2|ADGRV1|AGBL3|AK5|ALDH1L2|ALG10B|ALG14|ANAPC1|ANKRD7|ANKS1B|ANO5|AP5M1|APAF1|ARHGAP28|ARHGEF12|ARMC10|ASPM|ATAD2|ATL3|ATRNL1|BARD1|BRCA1|BRCA2|BRIP1|BRMS1L|BUB1|C12orf45|C12orf48|C14orf145|C1orf112|C9orf85|CACNB4|CBWD5|CC2D2A|CCDC15|CCDC150|CCDC178|CDC27|CDH10|CDH18|CDH7|CDK14|CDON|CENPE|CEP152|CEPT1|CHEK2|CHPT1|CISD2|CLSPN|CMC2|CNTNAP5|COG6|CPD|CRISPLD1|CSMD3|CSRP2|DAB1|DBT|DDX20|DEPDC1|DEPDC5|DGKE|DGKI|DIAPH3|DLEU1|DOCK7|DR1|DYNC1I1|E2F6|EFCAB5|EML1|ENPP1|ENPP2|EPHA6|EPM2A|ERBB4|ESCO2|EXOC5|EXOG|FAM114A2|FAM135A|FANCD2|FANCI|FANCM|FAT3|FBXO3|FGF7|FGGY|FLRT2|FUT9|G2E3|GABRB2|GABRG2|GEMIN5|GEN1|GNAI3|GPAM|GREB1L|GRID2|GRIN2A|GRIP1|GTPBP10|HCN1|HSD17B4|HYDIN|INTS2|INTS6|INTS7|INTU|ITGA8|KAT7|KIAA0586|KIAA1328|KIF13A|KIF14|KIF15|KLHL3|KNTC1|L2HGDH|LAMA1|LAMA3|LGR5|LINC00467|LINC01138|LMBR1|LMO7|LRRIQ3|MCM9|MDN1|MED17|MKLN1|MMS22L|MROH8|MTHFD2L|MYH15|NAV3|NCAPG2|NECAB1|NEDD1|NF1|NNT|NSF|NUP133|NUP155|NUP37|NUP93|OSBPL1A|PART1|PCCA|PCDH15|PDE4DIP|PGGT1B|PHLPP1|PHTF2|PIAS2|PIEZO2|PIGK|PLD5|POLQ|POLR3B|POLR3G|POU2F1|PPARGC1A|PPAT|PRIM2|PRKAA2|PRRC1|PTGFR|PTPRG|PTPRT|RANBP17|RBBP8|RNASEH2B|RNF217|RORB|RP11-3B12.1|RP11-82L2.1|RPAP2|RPS6KA5|RPS6KB1|RTTN|SAR1B|SCO1|SEC22B|SEC23A|SEMA3A|SEMA3E|SEMA5A|SEPSECS-AS1|SLC1A2|SLC35F4|SLC44A5|SLMAP|SMC6|SOCS7|SOX5|SRGAP2B|SRGAP2C|SSX2IP|STIL|STRIP2|TAF4B|TAOK1|TBC1D31|TCEANC2|THSD7B|TRPM3|TSIX|TTC26|TTC5|TTLL5|TYW5|USP32|USP37|VPS13D|WDHD1|WDR35|WDR7|WRN|XKR4|XPNPEP3|ZDHHC13|ZGRF1|ZNF277|ZNF426|ZNF519|ZNF546|ZNF678|ZNF720|ZNF736|ZRANB3</td><td>1.16E-12</td><td>**</td></tr>
<tr><td>Undifferentiated (nC3)</td><td>Vascular endothelium</td><td>40</td><td>AASS|ABHD12B|AC074093.1|ADGRL2|ARHGEF12|CADPS2|CD109|CDH13|CECR2|CEP112|CNTNAP3|COL21A1|CPD|CSRP2|CTD-2230D16.1|CTD-2536I1.1|DENND2C|DOCK9|ETV1|FLRT2|GRAMD1C|HMCN1|IPO11|LAMA3|LEPR|LGR4|LTBP1|MET|MIR99AHG|NTRK2|PCSK5|PDZD2|PGM5|PIEZO2|PTPRG|RNF217|RP11-327J17.3|SRGAP2C|STK3|TPD52L1</td><td>1.80E-04</td><td>**</td></tr>
<tr><td>Undifferentiated (nC3)</td><td>Leukocytes</td><td>41</td><td>ACSM3|ARHGAP24|ATP8B4|BEST3|C12orf42|CCDC26|CD226|COL19A1|CR1|CR2|CTC-454M9.1|DDX60|DSC2|DTHD1|F5|FCRL5|FLT3|HERC5|IL1RAP|IPCEF1|KLRD1|LRMP|LRRK2|MBOAT1|MCTP2|MGST1|MME|MR1|MYB|OTOA|PDE4D|PIP5K1B|PKHD1L1|RP11-58E21.3|RP11-90D4.2|SPECC1|SPTA1|SULT1B1|SYTL2|THEMIS|TTC39B</td><td>2.30E-02</td><td>*</td></tr>
</table>
