## Supplementary Table 5 for "Single-nuclei transcriptomes from human adrenal gland reveals distinct cellular identities of low and high-risk neuroblastoma tumors"

### Human NB to Mouse AG (Figure 5)

| Cell cluster in human NB | Cell cluster in mouse AG (reference) | Number of genes in the specific signature of NB shared with the specific signature of mouse AG | Genes in the specific signature of NB shared with the specific signature of mouse AG | FDR | Significance |
|---|---|---|---|---|---|
| Endothelial (nC4) | Endothelial (mC4) | 24 | APLNR\|ARHGEF15\|BCL6B\|C10orf10\|DLL4\|DOCK6\|EFNA1\|EGFL7\|FLT1\|FLT4\|GNG11\|HSPG2\|LMO2\|NOS3\|PLAT\|PLVAP\|RAPGEF3\|RASIP1\|SCARF1\|SHANK3\|SLC9A3R2\|SOX17\|STC1\|TMEM204 | 1.21E-21 | ** |
| Endothelial (nC4) | Macrophages (mC8) | 4 | BHLHE40\|NFKBIA\|PVR\|SOCS3 | 5.00E-02 | * |
| Macrophages (nC6) | Macrophages (mC3) | 29 | ABCC3\|ACP5\|APOE\|C5AR1\|CD84\|CLEC7A\|CLN8\|CSAR2\|CSF1R\|CSF3R\|CTSB\|CTSS\|CTSZ\|DOK3\|FYB1\|HEXB\|LAIR1\|LGMN\|MAFB\|MERTK\|MS4A7\|MSR1\|NPL\|PLA2G7\|RAB20\|SAT1\|SIGLEC1\|SLC11A1\|ZMYND15 | 3.12E-26 | ** |
| Macrophages (nC6) | Macrophages (mC14) | 15 | ADAM8\|ANPEP\|FGD2\|GNA15\|GSAP\|HCLS1\|IRF5\|IRF8\|LAPTM5\|MYO1F\|PARVG\|PSAP\|RAB7B\|SLC8B1\|TRPM2 | 4.58E-08 | ** |
| Macrophages (nC6) | Macrophages (mC8) | 7 | ALOX5\|C15orf48\|CD163\|EMILIN2\|FGR\|MCL1\|TREM1 | 6.23E-05 | ** |
| MSC (nC1) | Mesenchymal (mC6) | 4 | CRISPLD2\|ERRFI1\|PDGFRB\|WISP1 | 1.46E-03 | ** |
| MSC (nC1) | Capsule (mC13) | 4 | INHBA\|MYL9\|PKDCC\|RP3-412A9.11 | 1.50E-03 | ** |
| NOR (nC7) | Chromaffin nAChR $\alpha7^{high}$ (mC15) | 26 | ADCY1\|ASIC4\|CAMK2B\|CHGA\|DBH\|DNM1\|EML5\|ERC2\|GATA3\|GCGR\|GNAS\|GPRASP1\|HID1\|KCNQ2\|MAPT\|NDUFA4L2\|NXPH4\|PCLO\|PTPRF\|RPS6KL1\|SLC18A1\|SMPD3\|SNAP91\|SRRM3\|STXBP1\|UNC5A | 3.07E-12 | ** |
| NOR (nC7) | Chromaffin nAChR $\alpha7^{low}$ (mC11) | 3 | CYB561\|MAP1B\|UCHL1 | 1.66E-02 | * |
| NOR (nC9) | Chromaffin nAChR $\alpha7^{high}$ (mC15) | 15 | ASTN1\|CHGB\|CXXC4\|DPYSL3\|EML6\|FSTL5\|GNAO1\|GPR22\|JPH3\|PHYHIPL\|SCG2\|SLIT3\|SYT1\|SYT4\|ZDBF2 | 4.23E-04 | ** |

### Human NB to Mouse anlagen E13 (Figure 5)

| Cell cluster in human NB | Cell cluster in mouse anlagen E13 (reference) | Number of genes in the specific signature of NB shared with the specific signature of mouse adrenal anlagen at E13 | Genes in the specific signature of NB shared with the specific signature of mouse adrenal anlagen at E13 | FDR | Significance |
|---|---|---|---|---|---|
| Endothelial (nC4) | SCP | 64 | ADAMTS1\|ADAMTS4\|AHNAK\|ANXA2\|APOLD1\|CLEC14A\|COL15A1\|COL4A1\|CRIP2\|DUSP6\|EFNB1\|ELK3\|ENDOD1\|ENG\|EPHA2\|EPHB4\|ETS1\|FAM129B\|FAM198B\|FOXO1\|GJA1\|GRASP\|GRB10\|HES1\|HSPG2\|ID3\|IGFBP3\|IL6ST\|ITGA5\|KCTD12\|KIAA1462\|LDB2\|LIMS2\|LMNA\|LUZP1\|MCAM\|MYH9\|NEDD9\|NOTCH1\|PDGFB\|PDLIM1\|PDLIM4\|PIEZO1\|PLAT\|PLEC\|PLEKHG2\|PLS3\|PRAG1\|PVR\|PXN\|RBPMS\|RHOC\|RHOJ\|S100A16\|SASH1\|SKI\|SMAD3\|SNAI1\|SPARCL1\|SYNPO\|TGFBR2\|TM4SF1\|TMEM2\|WWTR1 | 2.18E-21 | ** |
| Macrophages (nC6) | SCP | 27 | ABCA1\|APBB1IP\|BMF\|CELF2\|DAB2\|GRN\|LCP2\|LITAF\|NR4A2\|P2RX7\|PARP14\|PLCB2\|PLXDC2\|PREX1\|PTPN6\|RAB31\|RAP2B\|RXRA\|SAT1\|STK10\|SYNGR2\|TCIRG1\|TGIF1\|TNS3\|WIPI1\|ZEB2\|ZFP36L1 | 1.53E-02 | * |
| MSC (nC1) | SCP | 21 | ALDH1A3\|AXL\|CALD1\|COL16A1\|COL1A2\|COL27A1\|COL5A1\|COL6A2\|CRISPLD2\|F3\|FN1\|LIMA1\|MICALL2\|MYL9\|NOTCH3\|NR4A1\|PALM2\|PLEKHA4\|RP3-412A9.11\|THBS2\|VCAN | 1.75E-10 | ** |
| NOR (nC7) | Chromaffin | 36 | ADCY1\|ADGRB2\|AHI1\|AMIGO2\|ANKRD12\|ASIC4\|CHD5\|CHGA\|CNTFR\|CYB561\|DBH\|DDC\|DNM1\|DOC2B\|DOCK4\|DPP6\|EML5\|ERC2\|GINM1\|GNAS\|GNB3\|GPRASP1\|HID1\|LRRC16B\|LUC7L3\|NDUFA4L2\|NPY\|NXPH4\|RUNDC3A\|SDCCAG8\|SLC18A1\|SMPD3\|SNAP91\|TH\|TXNIP\|UNC5A | 1.53E-08 | ** |
| NOR (nC7) | Sympathoblast | 28 | AKAP6\|ATAD5\|BMPR1B\|CASP8AP2\|EBF1\|ELAVL2\|GFRA3\|GPR137C\|HMX1\|KCNQ2\|MAP1B\|MAST1\|MCM7\|MLLT4\|NDRG4\|PHF19\|PLXNA4\|POU2F2\|PTBP2\|RBFOX1\|RGS7\|RPS6KL1\|SFPQ\|STMN1\|STMN2\|TUBA1A\|UCHL1\|ZFHX3 | 4.73E-04 | ** |
| NOR (nC8) | Sympathoblast | 22 | ATP1B1\|BASP1\|CRABP1\|CREB5\|EYA1\|FRMD3\|KHDRBS3\|KIT\|NEFL\|OLFM1\|ONECUT2\|PCSK2\|RUNX1T1\|RUSC2\|SNRPN\|STMN3\|TMEFF2\|TSPAN13\|TUBA1B\|TUBB2A\|TUBB4A\|TUBB4B | 1.31E-04 | ** |
| NOR (nC8) | Chromaffin | 16 | CBX6\|CHMP5\|DKK3\|DLK1\|DUSP26\|GAL\|NPTX2\|PCSK1N\|PNMA1\|RGS4\|RHBDD2\|SH3GLB2\|THRA\|TMEM176A\|TPI1\|TUB | 1.35E-02 | * |
| NOR (nC9) | Sympathoblast | 32 | ADD2\|CCT7\|CENPJ\|CHN1\|CLCN3\|CPE\|CSRNP3\|DCX\|DUSP8\|DUT\|H2AFZ\|HINT1\|HMGB2\|INPP5F\|JPH3\|KCNQ3\|KIAA1549L\|KIAA1715\|KIF26A\|KIF26B\|PHYHIPL\|PROX1\|SCG3\|SEPT11\|SEPT6\|SLC1A4\|SLC6A2\|SMC1A\|STMN4\|SV2C\|TAGLN3\|TMEM59L | 1.42E-05 | ** |
| NOR (nC9) | Chromaffin | 28 | CAMK2N1\|CAMK4\|CD47\|CHGB\|CNRIP1\|CYGB\|EPM2AIP1\|FAM63B\|FSTL5\|GNAO1\|IRS2\|KCNK3\|LRRN3\|N4BP2\|PKIB\|RAB3C\|RGS5\|SCG2\|SCG5\|SEZ6\|SEZ6L\|SMARCA1\|ST6GALNAC5\|STAC\|SYP\|SYT1\|TMOD2\|ZDBF2 | 1.13E-04 | ** |

**Human NB to Human AG (Figure 5)**

<table>
<tr><th>Cell cluster in human NB</th><th>Cell cluster in human AG (reference)</th><th>Number of genes in the specific signature of NB shared with the specific signature of human AG</th><th>Genes in the specific signature of NB shared with the specific signature of human AG</th><th>FDR</th><th>Significance</th></tr>
<tr><td>Endothelial (nC4)</td><td>Endothelial (hC6)</td><td>55</td><td>ADAM15|ADCY4|ADGRL4|APOLD1|ARHGEF15|CALCRL|CD34|CDH5|CLEC14A|CPLX1|CXorf36|DOCK6|EFNA1|EGFL7|ELK3|EPHB4|ESAM|FLT1|FLT4|FZD4|HSPG2|HYAL2|ID1|IFI27|IGFBP4|ITPRIP|LDB2|LIMS2|LMO2|MMRN2|MYCT1|NOS3|PCDH12|PCDH17|PECAM1|PLAT|PLPP3|PLVAP|PODXL|RAMP2|RAMP3|RASIP1|ROBO4|S1PR1|SCARF1|SDPR|SEPN1|SHANK3|SLC9A3R2|SLCO2A1|SOX18|TGM2|TIE1|TM4SF1|VWA1</td><td><1E-50</td><td>**</td></tr>
<tr><td>Endothelial (nC4)</td><td>Z. fasciculata (hC3)</td><td>5</td><td>ADAMTS9|FLNB|LDLR|MIDN|YBX3</td><td>2.35E-02</td><td>*</td></tr>
<tr><td>Macrophages (nC6)</td><td>Macrophages (hC2)</td><td>87</td><td>ACSL1|ADAP2|ALOX5|ARRB2|ASAH1|BMP2K|BTK|C1orf162|C5AR1|CD14|CD163|CD86|CLEC7A|CSF1R|CSF2RA|CSF3R|CTSB|CTSH|CYBB|DOK3|EMILIN2|FCER1G|FCGR2C|FCGRT|FCHO2|FGD2|FGR|FNIP2|FPR3|GPR183|GRN|GSAP|HAVCR2|HCK|HCLS1|HLA-DMB|HSPA6|IFI30|IL18|IRF5|ITGAX|LAIR1|LILRA6|LILRB1|LILRB2|LILRB3|LILRB4|LILRB5|MAFB|MIR181A1HG|MKNK1|MPEG1|MS4A4A|MS4A6A|MS4A7|MSR1|NCF4|NPL|PLXDC2|PRKAG2|RASSF2|RASSF4|REL|RHBDF2|RIN3|RN7SL368P|RNA5SP498|RNASET2|RNF144B|RNF149|SAT1|SIGLEC1|SLC11A1|SLC1A3|SLC43A2|SLCO2B1|SMAP2|SPI1|TBC1D12|TFEC|TGFBI|THEMIS2|TLR2|TNFAIP2|TRPM2|TYMP|TYROBP</td><td><1E-50</td><td>**</td></tr>
<tr><td>Macrophages (nC6)</td><td>Z. fasciculata (hC3)</td><td>8</td><td>ATP1B3|ELL2|GRAMD4|PAPSS2|RAB20|RGS2|TIPARP|ZNF331</td><td>1.33E-04</td><td>**</td></tr>
<tr><td>MSC (nC1)</td><td>Mesenchymal (hC7)</td><td>20</td><td>AEBP1|C11orf96|C1R|C1S|CALD1|CFHR1|COL16A1|COL1A2|COL27A1|COL6A2|COL6A3|CRISPLD2|CTD-2033D15.2|CTD-2033D15.3|ERRFI1|F3|PDGFRB|SERPINE1|THBS1|WISP1</td><td>4.59E-28</td><td>**</td></tr>
<tr><td>NOR (nC7)</td><td>Chromaffin (hC4)</td><td>50</td><td>AC006019.4|AC138969.4|AHI1|AKAP6|APC2|ASIC3|BICD1|C1orf226|CAMK2B|CHD5|CHGA|CYB561|DBH|DDC|DPP6|EML5|GCGR|GNB3|GPRASP1|GRIK2|GSE1|KCNQ2|MAPT|MAST1|MDGA1|NCAM1|NDRG4|NOS1AP|PCLO|PHC1|PKD1|RP11-1212A22.4|RP11-216L13.19|RP11-416N2.4|RP11-490G2.2|RP11-54O7.3|RP11-728K20.2|RUNDC3A|SGK494|SLC18A1|SNAP91|SRRM3|SRRM4|STMN2|STXBP1|TFAP2B|TH|UCHL1|UNC5A|ZC3H14</td><td>1.40E-45</td><td>**</td></tr>
<tr><td>NOR (nC8)</td><td>Chromaffin (hC4)</td><td>14</td><td>ATP1B1|CHRFAM7A|CHRNA3|DUSP26|FRRS1L|HDAC9|KHDRBS3|NEFM|PCSK1N|PCSK2|RGS4|RP11-650L12.2|ST8SIA3|TUB</td><td>1.11E-08</td><td>**</td></tr>
<tr><td>NOR (nC8)</td><td>Z. glomerulosa (hC5)</td><td>2</td><td>DACH1|VSNL1</td><td>2.25E-02</td><td>*</td></tr>
<tr><td>NOR (nC9)</td><td>Chromaffin (hC4)</td><td>25</td><td>ALCAM|CHGB|CNTN1|DPYSL3|FSTL5|HTATSF1|INPP5F|NTNG1|PREPL|RAB3C|RAMP1|RGS5|RP11-346C4.2|SCG2|SCG3|SCG5|SEZ6L|SLC25A12|SLC6A2|SYT1|SYT4|TAGLN3|TEAD1|TMEM59L|VAT1L</td><td>3.33E-16</td><td>**</td></tr>
<tr><td>T-cells (nC10)</td><td>T-cells (hC10)</td><td>4</td><td>CD2|TRAC|TRBC1|TRBC2</td><td>1.28E-05</td><td>**</td></tr>
<tr><td>Undifferentiated (nC2)</td><td>Progenitor (hC1)</td><td>4</td><td>CH507-513H4.1|CH507-528H12.1|ELF3|RP13-262C2.4</td><td>7.69E-04</td><td>**</td></tr>
<tr><td>Undifferentiated (nC3)</td><td>Progenitor (hC1)</td><td>239</td><td>A1CF|A2ML1|ABCA13|ABCB5|AC007064.24|AC007405.4|AC007731.1|AC008079.9|AC008103.3|AC008103.4|AC008132.15|AC008697.1|AC009264.1|AC011718.2|AC074093.1|AC099057.8|AC108025.2|AC144521.1|ACOXL|ACSM2A|ACSM2B|ADAM32|ADAM7|ADAMTS20|ADAMTS6|ADCY10|ADGRF1|AFF2|AJ006998.2|AL035610.2|ANKRD30A|ANKRD30BL|AP000432.3|ARHGAP28|ARMC3|ASPM|B4GALNT2|BRCA1|BUB1|C12orf42|C12orf45|C1orf141|C1orf87|CCDC150|CCDC168|CCDC26|CECR2|CFAP54|CFAP58|CFTR|CH17-264L24.1|CH507-154B10.1|CLDN11|CLNK|CMYA5|CNTN5|COL19A1|COL21A1|COL24A1|COL6A5|COL9A1|CORIN|CR2|CSMD3|CTD-2144E22.9|CTD-2311B13.5|CTNNA3|CUBN|DCDC1|DDX4|DIAPH3|DLX6-AS1|DNAH10|DNAH11|DNAH3|DNAH9|DOCK7|DSC1|DSG2|DSG3|EFCAB5|EGF|EPHA6|EPM2A|ERVW-1|ESCO2|FAM230A|FAM230B|FREM2|FRK|GABRA2|GBP6|GOLGA6L22|GYPA|HCG23|HELLPAR|HMCN1|HMGA2|HULC|IGFN1|KB-1183D5.13|KCNU1|KIAA1257|KIF14|KIF15|KIF6|KNTC1|LA16c-23H5.4|LA16c-3G11.5|LA16c-60D12.1|LA16c-60D12.2|LAMA1|LAMA3|LINC00466|LINC00923|LPA|LRP1B|LVRN|MALRD1|MGAM|MME|MMS22L|MUC16|MUC19|MUC3A|MYH15|MYO1H|MYO3A|MYPN|NAALADL2|NANOG|NEB|NEK10|NF1|OR10K1|OR1A1|OR2M3|OR2T12|OR4F17|OR4F4|OR4F5|OR56A4|OR5AS1|OR7C1|OTOA|OTOGL|PCAT1|PCSK5|PIEZO2|PKHD1|PLCH1|POLQ|PRAMEF13|PRAMEF25|PSG11|PTPRQ|PTPRT|PTPRZ1|RASSF6|RF00012|RF00017|ROS1|RP1-101G11.3|RP1-14N1.2|RP11-114G22.1|RP11-115D19.1|RP11-139E24.1|RP11-142I20.1|RP11-146E13.4|RP11-15E1.5|RP11-168K9.1|RP11-169F17.1|RP11-215N21.1|RP11-21F16.1|RP11-244H18.1|RP11-255B23.3|RP11-267C16.1|RP11-341D18.3|RP11-34P13.13|RP11-374A4.1|RP11-395L14.18|RP11-416O18.1|RP11-430C1.1|RP11-458B24.2|RP11-459O16.8|RP11-461F11.2|RP11-482G13.1|RP11-4L24.3|RP11-532N4.2|RP11-536C10.7|RP11-541P9.3|RP11-573D15.8|RP11-586K12.1|RP11-597A11.2|RP11-673E1.1|RP11-735B13.1|RP11-774D14.1|RP11-776H12.1|RP4-760C5.5|RP4-809F18.1|RP5-916L7.1|RP5-947P14.1|SCN11A|SEMA3E|SI|SLC13A1|SLC44A5|SLC4A4|SLC9B1|SLC9C1|SLC9C2|SOCS7|SOX2-OT|SOX9-AS1|SPATA17|SPINK5|SPTA1|STK31|SULT1B1|TAT|TF|TMEM132B|TP63|TPH2|TRIM49|TRIM49B|TRIM49L2|TRIM51|TRIM64|TRIM64B|TRPM3|TSIX|TYR|USH2A|UTS2B|XKR4|XXbac-BPG55C20.7|ZNF705B|ZPLD1</td><td><1E-50</td><td>**</td></tr>
</table>

| | | | | | |
|---|---|---|---|---|---|
| Undifferentiated (nC3) | Mesenchymal (hC7) | 10 | ABCA6\|ABCA9\|FAM153A\|FAM153C\|HGF\|LAMA2\|PLA2R1\|RP11-327J17.3\|RP11-826N14.7\|STEAP4 | 1.07E-03 | ** |

**Human NB to Human fetal AG (Dong et al. 2020, Figure 5)**

| Cell cluster in human NB | Cell cluster in human fetal AG (Dong et al. 2020) | Number of genes in the specific signature of human NB clusters shared with genes differentially expressed in human fetal AG (Dong et al. 2020) | Genes in the specific signature of human NB clusters shared with genes differentially expressed in human fetal AG (Dong et al. 2020) | FDR | Significance |
|---|---|---|---|---|---|
| Endothelial (nC4) | SCPs | 59 | AHNAK\|ANGPTL4\|ANXA2\|APP\|BST2\|CD59\|CD99\|COL15A1\|COL4A1\|DUSP5\|DUSP6\|EIF5B\|EMP2\|ENDOD1\|ENO1\|ETS1\|FBL\|GJA1\|GNAI2\|GNG11\|GRPEL1\|HES1\|HLA-E\|HSPG2\|ID1\|ID3\|IFITM2\|IFITM3\|IGFBP4\|IGFBP7\|IL6ST\|INSR\|KCTD12\|KLF2\|LDLR\|LMNA\|LUZP1\|MAN1A1\|NFKBIA\|NOTCH1\|PDLIM1\|PLAT\|PLS3\|RAB13\|RBPMS\|RHOC\|RHOJ\|RNASE1\|S100A10\|S100A16\|S100A6\|SASH1\|SERTAD1\|SLC25A37\|SOCS3\|SPRY1\|TGFBR2\|WWTR1\|YBX3 | <1E-50 | ** |
| Endothelial (nC4) | Non-cycling chromaffin cells* | 15 | ARL2\|BAG3\|C2CD4B\|CCDC85B\|EGFL7\|ETS2\|FXYD5\|HSPA5\|IGFBP7\|LMNA\|MIDN\|PDLIM4\|PRNP\|S100A6\|STC1 | 8.80E-07 | ** |
| Endothelial (nC4) | Cycling chromaffin cells* | 10 | APOLD1\|ARL2\|BAG3\|CDKN1A\|CLIC4\|GNAI2\|HIPK3\|LMNA\|MIDN\|NRGN | 2.68E-03 | ** |
| Macrophages (nC6) | SCPs | 29 | ABCA1\|APOE\|ARHGAP18\|ATP1B3\|CD83\|CELF2\|DAB2\|FCGRT\|GLIPR2\|GSTO1\|LITAF\|MCL1\|PAPSS2\|PLXDC2\|PPIF\|RAB20\|RAB31\|RASSF4\|RBM47\|SAMHD1\|SAT1\|SCARB1\|SLC15A3\|SOAT1\|TGIF1\|TNS3\|ZEB2\|ZFP36\|ZFP36L1 | 5.56E-15 | ** |
| Macrophages (nC6) | Non-cycling chromaffin cells* | 15 | AMPD2\|CSTB\|CTSL\|FOSL2\|GPNMB\|GRN\|HLA-DQB1\|HLA-DRB1\|HLA-DRB5\|HSPA6\|NR4A2\|PSAP\|RNASET2\|SLC31A2\|ZNF331 | 2.81E-06 | ** |
| MSC (nC1) | SCPs | 17 | AXL\|C11orf96\|CALD1\|COL1A2\|COL6A2\|FN1\|ID4\|LIMA1\|LMCD1\|MT1E\|MYL9\|NR4A1\|PLEKHA4\|RAB34\|SPRY2\|TPM2\|VCAN | 5.36E-11 | ** |
| NOR (nC7) | Non-cycling sympathoblasts* | 36 | ACTG1\|C14orf132\|C1QTNF2\|CAMK2B\|CNTFR\|ECEL1\|ELAVL2\|EVL\|EXOC4\|GABPB1-AS1\|GATA2\|GATA3\|GFRA3\|HMX1\|ING4\|LINC00340\|LMO1\|LUC7L3\|MAP1B\|MAPT\|MEIS3\|NOP56\|NPY\|NTRK1\|PHF14\|POU2F2\|RBFOX1\|RGS7\|SOX4\|STMN1\|STMN2\|TFAP2B\|TTC3\|TUBA1A\|UCHL1\|ZFHX3 | 4.47E-33 | ** |
| NOR (nC7) | Cycling sympathoblasts* | 18 | DAAM1\|EBF1\|EVL\|EZH2\|GTF2I\|MAP1B\|MCM7\|NCAM1\|PHF19\|PLXNA4\|PTBP2\|RBFOX1\|SFPQ\|SLC38A1\|SNAP91\|STMN1\|TFAP2B\|ZFHX3 | 1.70E-09 | ** |
| NOR (nC7) | Non-cycling chromaffin cells* | 18 | C2CD4A\|CHGA\|CYB561\|DBH\|DDC\|DPP6\|GCGR\|GINM1\|GNAS\|KCNQ2\|MEG3\|MTRNR2L8\|NDRG4\|NDUFA4L2\|SLC18A1\|STXBP1\|TH\|TXNIP | 2.40E-08 | ** |
| NOR (nC7) | Cycling chromaffin cells* | 14 | C2CD4A\|CDK10\|CHGA\|EZH2\|GINM1\|KCNQ2\|MCM7\|NDRG4\|PAXBP1\|PHF19\|SLC18A1\|SOBP\|STXBP1\|TH | 2.96E-05 | ** |
| NOR (nC8) | Non-cycling sympathoblasts* | 20 | ATP6V0E2\|BASP1\|C9orf16\|DPYSL4\|DUSP26\|GAL\|HTR3A\|NEFL\|NEFM\|OLFM1\|RGS4\|SNRPN\|STMN3\|STRAP\|THRA\|TMEFF2\|TUBB2A\|TUBB4A\|WRB\|YWHAZ | 1.08E-18 | ** |
| NOR (nC8) | Non-cycling chromaffin cells* | 23 | ATP1B1\|CBX6\|CHMP5\|CHRNA3\|CUTA\|DKK3\|DLK1\|HACD3\|HMGN3\|ITM2C\|KHDRBS3\|NPTX2\|PCSK1N\|PCSK2\|PTN\|QPCT\|RGS4\|RHBDD2\|SERPINE2\|SURF4\|TMBIM4\|TMED4\|TXN | 5.45E-18 | ** |
| NOR (nC8) | Cycling sympathoblasts* | 12 | BASP1\|CREB5\|CRNDE\|GNB1\|NFIB\|ONECUT2\|RUNX1T1\|THSD7A\|TUBA1B\|TUBA1C\|TUBB4B\|VDAC3 | 1.36E-07 | ** |
| NOR (nC8) | Cycling chromaffin cells* | 13 | CRNDE\|DKK3\|DLK1\|HACD3\|HMGN3\|NPTX2\|PCSK1N\|PTN\|TMBIM4\|TUBA1B\|TUBA1C\|TUBB4B\|TUBG1 | 2.26E-07 | ** |
| NOR (nC8) | SCPs | 7 | CRABP1\|GPC6\|MAPRE2\|PTN\|SCD5\|SCP2\|TMEM176A | 2.20E-02 | * |
| NOR (nC9) | Non-cycling sympathoblasts* | 34 | ALCAM\|BRK1\|CAMLG\|CHST1\|CLIP3\|CNRIP1\|CYGB\|DCX\|DPYSL3\|EEF2\|EIF1B\|EIF3L\|GNAO1\|HMGA1\|MAP4\|OPTN\|PGM2L1\|PHYHIPL\|PKIB\|PMP22\|PODXL2\|RBMS3\|RGS5\|RUFY3\|SCG3\|SCG5\|SLC6A2\|STMN4\|TAGLN3\|TBCB\|TKT\|TMSB15A\|TSPAN7\|UBE2E3 | 5.38E-32 | ** |
| NOR (nC9) | Cycling sympathoblasts* | 29 | CHN1\|CKS1B\|COX8A\|DLEU2\|DUT\|FOXP1\|H2AFZ\|HDGF\|HIST1H4C\|HLTF\|HMGA1\|HMGB1\|HMGB2\|INPP5F\|KLHL23\|MEIS2\|MTHFD2\|PBRM1\|PCDH7\|PROX1\|RBMS3\|SAE1\|SESTD1\|SMC1A\|SYNPO2\|TMSB15A\|TOP2B\|TSHZ2\|XRCC5 | 2.79E-21 | ** |
| NOR (nC9) | Non-cycling chromaffin cells* | 23 | BEX2\|BSG\|CAMK2N1\|CAMK4\|CHGB\|COX7C\|CPE\|GSTP1\|HTATSF1\|NDUFA1\|RAB3C\|RAMP1\|RBMS1\|RSL24D1\|SCG2\|SCG5\|SEC62\|SYP\|SYT1\|SYT4\|TMA7\|VAT1L\|ZDBF2 | 2.49E-13 | ** |
| NOR (nC9) | Cycling chromaffin cells* | 21 | CHGB\|CKS1B\|CLGN\|CNTN1\|COX8A\|DLEU2\|DUT\|FANCL\|H2AFZ\|HINT1\|HIST1H4C\|HMGB2\|HTATSF1\|MTHFD2\|RAMP1\|SAE1\|SCG2\|SMC1A\|SYT4\|VAT1L\|ZDBF2 | 5.16E-11 | ** |
| NOR (nC9) | SCPs | 13 | ABHD2\|FIGN\|IGFBP5\|ITGA1\|JUN\|LDHB\|MBNL2\|PMP22\|RND3\|SCARB2\|SCN7A\|SFRP1\|SLC39A10 | 4.61E-04 | ** |
| Undifferentiated (nC3) | Cycling chromaffin cells* | 24 | ASPM\|ATAD2\|BARD1\|BUB1\|C12orf48\|CDC27\|CDH7\|CENPE\|CEP152\|CFAP61\|CLSPN\|CMC2\|DLEU1\|DPY19L3\|FANCD2\|FANCI\|GULP1\|LAMA3\|MROH8\|NCAPG2\|NUP155\|RNASEH2B\|SAR1B\|ST18 | 9.14E-05 | ** |

| | | | | | |
|---|---|---|---|---|---|
| Undifferentiated (nC3) | Cycling sympathoblasts* | 15 | ASPM\|ATAD2\|C12orf48\|CENPE\|CLSPN\|CMC2\|CSRP2\|DEPDC1\|ESCO2\|FANCI\|G2E3\|KIF14\|MMS22L\|RBBP8\|RNASEH2B | 1.84E-02 | * |
