## Supplementary Table 6 for "Single-nuclei transcriptomes from human adrenal gland reveals distinct cellular identities of low and high-risk neuroblastoma tumors"

### Human NB to Risk in 498 NBs (Fig 6)

<table>
<thead>
<tr><th>Cell cluster in human NB</th><th>Risk in 498 SEQC NB samples</th><th>Number of genes in the specific signature of NB significantly up-regulated in Risk group(s) in 498 SEQC NB samples</th><th>Genes in the specific signature of NB significantly up-regulated in Risk group(s) in 498 SEQC NB samples</th><th>FDR</th><th>Significance</th></tr>
</thead>
<tbody>
<tr><td>Endothelial (nC4)</td><td>Low/Intermediate risk</td><td>165</td><td>ACVRL1|ADAM15|ADAMTS1|ADCY4|AHNAK|AKAP2|ANXA2|APLNR|APOLD1|APP|AQP1|ARHGEF15|ATP1A1|BCL3|BCL6B|BHLHE40|BST2|C2CD4B|C6orf208|CA2|CALCRL|CAV1|CCDC69|CD34|CD93|CDH5|CDK17|CLEC14A|CLIC4|CMIP|CNKSR3|COL15A1|COL4A1|CRIP2|DLL4|DOCK6|DUSP5|DUSP6|ECE1|EFHD2|EGFL7|EHD2|EHD4|ELK3|EML3|EMP1|EMP2|ENDOD1|ENG|EPAS1|EPHA2|EPHB4|ERG|ESAM|ETS1|ETS2|FAM129B|FLT1|FLT4|FOXO1|FUT11|FZD4|GABRE|GNAI2|GRASP|GRB10|HES1|HIPK3|HLA-E|HSPG2|IER3|IFITM2|IGFBP4|IGFBP7|IL33|IL6ST|ITGA5|ITPKC|ITPRIP|KCTD12|KIAA0355|KLF2|LAMA5|LDB2|LIMS2|LMNA|LRP10|LRRC32|LUZP1|MAN1A1|MARCH3|MAST4|MMRN2|MSN|MYADM|MYH9|NDRG1|NEDD9|NFIC|NFKBIA|NOTCH1|NPR1|PALMD|PCDH12|PCDH17|PDGFB|PECAM1|PIEZO1|PLAT|PLEC|PLEKHG2|PLS3|PLVAP|PODXL|PPM1F|PXN|RAMP3|RAPGEF3|RASIP1|RBPMS|RCAN1|RHOC|RHOJ|RNF19B|ROBO4|RP11-314N13.4|S100A10|S100A13|S100A16|S100A6|S1PR1|SASH1|SCARF1|SELE|SHANK3|SHROOM4|SKI|SLC35E4|SLC39A14|SLC9A3R2|SLCO2A1|SLK|SNRK|SOX17|SOX7|SPARCL1|SPRY1|SSH1|STOM|SWAP70|SYNPO|TGFB1|TGFBR2|THBD|TIE1|TINAGL1|TM4SF1|TMEM173|TMEM204|TMOD3|UPP1|VSIG2|VWA1|WWTR1|ZBTB7A</td><td><1E-50</td><td>**</td></tr>
<tr><td>Endothelial (nC4)</td><td>High risk</td><td>56</td><td>ARL2|ATP11C|BAG5|BAZ1A|BMP6|CCDC85B|CDC37|CDC42EP2|CDC42EP3|CEACAM19|CENPB|CHD7|CPLX1|DDX21|EDARADD|EFNB1|EIF5B|ENO1|FBL|FJX1|FKBP1A|FLNB|FOXC1|FZD6|GAR1|GAS5|GRPEL1|HAPLN3|HYAL2|MIDN|MTA2|NIP7|NRGN|PABPC1|PABPC3|PARVB|PDLIM4|PIM3|PKN3|PNP|PVR|RAI14|RAMP2|RPGR|RPL36A|RRP12|SEMA3F|SLC4A7|SLCO4A1|SMAD3|SNAI1|SOCS2|STC2|TEAD4|TMEM70|TNFRSF10B</td><td>1.02E-04</td><td>**</td></tr>
<tr><td>Macrophages (nC6)</td><td>Low/Intermediate risk</td><td>226</td><td>ABCA1|ABCC3|ACCS|ACP5|ADAM9|ADAP2|ALOX5|AMDHD2|AMPD2|AMPD3|ANPEP|APBB1IP|APBB3|APOE|ARHGAP18|ARHGAP4|ARL4C|ARL8B|ARRB2|ASAH1|ATG16L2|BHLHE41|BMF|BTK|CD14|CD163|CD300LF|CD58|CD74|CD83|CD84|CD86|CELF2|CIITA|CLEC7A|CPVL|CSF1R|CSF2RA|CSF3R|CTSH|CTSS|CTSZ|CXCL16|CXCL2|CYBB|CYP27A1|CYTH4|DAB2|DENND3|DMXL2|DOK3|EEPD1|ELL2|FCER1G|FCGRT|FCHO2|FERMT3|FGD2|FGR|FKBP15|FNIP2|FOSL2|FPR3|GAB3|GABARAPL1|GNA15|GPCPD1|GPNMB|GPR183|GRAMD4|GRINA|GRN|HAVCR2|HCK|HCLS1|HEXA|HK3|HLA-DMA|HLA-DMB|HLA-DOA|HLA-DPA1|HLA-DPB1|HLA-DQA1|HLA-DQA2|HLA-DQB1|HLA-DQB2|HLA-DRA|HLA-DRB1|HLA-DRB5|HTRA4|IFFO1|IFNGR1|IL18|IL6R|IRF8|ITGAD|ITGAM|ITGAX|KCNAB2|KLHL6|KMO|LACTB|LAIR1|LAPTM5|LCP2|LGALS3|LGALS9|LILRB1|LILRB2|LILRB3|LILRB4|LRP1|MAFB|MAP3K8|MERTK|METRNL|MFSD1|MFSD12|MGLL|MKNK1|MPEG1|MS4A4A|MS4A6A|MS4A7|MXD1|MYO1F|MYO9B|NABP1|NBPF10|NCF2|NCF4|NEAT1|NFKBID|NPL|NR4A2|OSCAR|P2RX7|PADI2|PAPSS2|PARP14|PARVG|PIK3R5|PLA2G7|PLCB2|PLEK|PLXDC2|PPARD|PREX1|PRKAG2|PSAP|PTK2B|PTPN6|RAB20|RAB31|RAB7B|RAPGEF1|RASGEF1B|RASSF2|RASSF4|RASSF5|RBM47|RGS1|RHBDF2|RIN3|RNF135|RNF144B|RNF149|RREB1|RXRA|SAMHD1|SAMSN1|SAT1|SDC3|SEMA4A|SETDB2|SGK1|SLC11A2|SLC15A3|SLC16A3|SLC17A9|SLC18B1|SLC1A3|SLC2A14|SLC31A2|SLC37A2|SLC43A2|SLCO2B1|SMAP2|SMPDL3A|SNHG12|SNORD99|SNX29|SOAT1|SPI1|STK10|STX4|STX7|STXBP2|SYNGR2|TBC1D12|TBC1D2|TCIRG1|TFEC|TGFBI|THEMIS2|TLR2|TMEM63A|TNFAIP2|TNFRSF14|TNFRSF1B|TNS3|TOM1|TPCN1|TPP1|TRIM38|TRPM2|TYROBP|UAP1L1|UBXN11|VAMP8|WDR91|YTHDF3|ZEB2|ZFP36|ZFP36L1|ZMYND15</td><td><1E-50</td><td>**</td></tr>
<tr><td>MSC (nC1)</td><td>Low/Intermediate risk</td><td>31</td><td>ADAM33|AEBP1|AXL|C1R|C1S|CALD1|COL16A1|COL1A2|COL27A1|COL5A1|COL6A2|COL6A3|CRISPLD2|ERRFI1|F3|FSTL3|GEM|ID4|LIMA1|LMCD1|MICALL2|MYL9|NOTCH3|NR4A1|PDGFRB|PLEKHA4|PRRX1|PTGIR|SNAI2|THBS2|VPS37D</td><td>1.40E-02</td><td>*</td></tr>
</tbody>
</table>

<table>
<tr><td>NOR (nC7)</td><td>Low/Intermediate risk</td><td>208</td><td>ACAD10|ACAP3|ADCY1|AGAP2|AGAP3|AGER|AGRN|AHI1|AKAP6|AKAP9|AMIGO2|ANKRD12|ANKRD13B|ANKRD36|ANKRD36B|ANKRD36C|ANKS1A|APC2|ARHGEF10L|ARID1B|ASIC3|ATF6B|ATG14|BMPR1B|BRSK2|BTBD3|C1QTNF2|C2CD4A|CALCOCO1|CAMK2B|CASZ1|CC2D1A|CCDC136|CCNL2|CDC42SE2|CHD5|CIC|CNTFR|CORO6|CRLF3|CYB561|DBH|DDC|DDX17|DIRAS3|DLC1|DMTF1|DNAJB2|DNM1|DOC2B|DOCK4|DPP6|DST|EBF1|ECEL1|ELAVL2|EPB41L4A|EPHB3|ERC2|ERV3-1|EXOC4|EYA4|FAM120B|FAM184A|FASTK|FNDC5|GAS8|GATA2|GCGR|GFRA3|GIGYF1|GINM1|GLG1|GNAS|GNB3|GPATCH8|GPR137C|GPRASP1|GRIK2|GTF2I|GTF2IRD1|GTF2IRD2|GTF2IRD2B|HECTD4|HID1|HIP1R|HMX1|HS3ST5|ING4|INSRR|KB-1572G7.1|KCNQ2|KCNQ5|KLC4|KLRK1|KRIT1|LAMA4|LMBR1L|LUC7L|LUC7L2|LUC7L3|MAP1B|MAPK8IP3|MAPT|MAST1|MDGA1|MED23|MGAT4B|MLXIP|MUC20|NCAM1|NCRNA00107|NDRG4|NDUFA4L2|NINL|NISCH|NKTR|NOS1AP|NRCAM|NTRK1|NXPH4|OBSL1|OCLN|PACS2|PCDHA9|PCLO|PDE9A|PEX6|PHC1|PI4KA|PIGQ|PKD1|PLAGL1|PLXNA4|PLXNB1|PNISR|POLR2J2|POU2F2|PPIP5K2|PRPF4B|PRPSAP2|PRRC2B|PRRT2|PSD2|PTBP2|PTPRF|PXDN|R3HDM2|RADIL|RAI1|RBCK1|RBFOX1|RBM33|RBM6|RERE|RGMB|RGS7|RIMKLB|RSBN1L|RUNDC3A|SDCCAG8|SEMA6C|SFSWAP|SGSM2|SHC2|SLC18A1|SNAP91|SOBP|SOGA3|SOX4|SPG7|SPTAN1|SRGAP3|SRRM1|SRRM3|SRRM4|SRSF11|STAG3|STK38|STMN1|STMN2|STXBP1|TAF1C|TBC1D24|TFAP2B|TH|TIA1|TLE4|TMEM191C|TSC2|TSTD2|TUBA1A|TXNIP|UBAP1L|ULK1|UNC5A|VPS9D1|WDR6|WDR86|WSB1|YJEFN3|ZCCHC14|ZNF117|ZNF33A|ZNF429|ZNF512B|ZNF608|ZRANB2</td><td><1E-50</td><td>**</td></tr>
<tr><td>NOR (nC8)</td><td>Low/Intermediate risk</td><td>102</td><td>ACLY|ANKRD6|ASIC1|ATP1B1|ATP6V0E2|BASP1|CBX6|CCDC28A|CCDC68|CDYL2|CHRFAM7A|CHRNA3|CREB5|DCLK1|DCLK3|DKK3|DUSP26|ELMO2|EPB41L1|EPB41L4B|EYA1|FAM219A|FECH|FRMD3|FRRS1L|FSD1L|GNAQ|GNB1|GPC6|HMGN3|HTR3A|IFT20|ISCA1|ISCU|ITGB8|ITM2C|JUP|KBTBD6|KIF3B|KLHL35|MAPRE2|MMD|NDRG3|NXPH1|OGFRL1|OLFM1|PCMT1|PCSK2|PDE10A|PHF1|PPP4R1|PSMD11|PTN|PTS|QPCT|RAB14|RAB15|RAB2A|RGS3|RGS4|RHBDD2|RIN2|RNF10|RNF170|RUNX1T1|RUSC2|SCP2|SCYL2|SEL1L3|SERINC3|SERPINE2|SH3GLB2|SLC25A4|SMAD9|SNRPN|SNX18|SORCS1|SPIN1|SRI|STMN3|SV2B|SYBU|THRA|THSD7A|TM2D3|TMBIM4|TMED4|TMEFF2|TMEM163|TMEM176A|TRPA1|TSPAN13|TUB|TUBB2A|TUBB4A|ULK3|VPS4B|VSNL1|WDR47|WRB|WSB2|YWHAZ</td><td>2.20E-39</td><td>**</td></tr>
<tr><td>NOR (nC8)</td><td>High risk</td><td>42</td><td>AKR1B1|ARL16|CERK|CLTA|CNOT7|COX6C|CRABP1|CRNDE|CS|CUTA|CYC1|EBF3|EBP|EDF1|ENC1|GAL|KIT|MRFAP1L1|MRPS21|MRPS26|NDUFA12|NDUFB9|NFIB|NHLH2|NPTX2|PNMA1|PSMA7|RPRD1A|SEC61B|SLC10A4|SMU1|SSBP1|ST8SIA3|STRAP|SURF4|TMED2|TMX2|TPI1|TUBB4B|TXN|VDAC3|WDR61</td><td>6.71E-06</td><td>**</td></tr>
<tr><td>NOR (nC9)</td><td>High risk</td><td>121</td><td>ALKBH7|ANKRD17|ATRX|BEX2|BMP7|BRK1|BSG|BTF3|CAMLG|CCT4|CCT7|CENPJ|CKS1B|COPE|COX6B1|COX7C|COX8A|DCX|DPY30|DUT|EEF1B2|EEF2|EIF2AK2|EIF3L|EPRS|ESD|FAU|FIGN|FXR1|GCSH|GSTP1|H2AFZ|HADHA|HDGF|HINT1|HIST1H4C|HIST1H4E|HLTF|HMGA1|HMGB1|HMGB2|HNRNPA3|HNRNPL|HSD17B11|HSD17B12|HSPA8|HTATSF1|JTB|KLHL23|LDHB|METTL9|MIR3654|MTHFD2|N4BP2|NDUFA1|NDUFB11|OIP5-AS1|OST4|PFN2|POLR1D|PRDX2|RMND5A|RPL14|RPL15|RPL18|RPL18A|RPL23A|RPL24|RPL26|RPL27|RPL29|RPL31|RPL32|RPL34|RPL35A|RPL39L|RPL4|RPS10|RPS11|RPS13|RPS15|RPS15A|RPS19|RPS21|RPS24|RPS27A|RPS3|RPS5|RPS7|RPS9|RPSA|RSF1|RSL24D1|SACS|SAE1|SEPT11|SF3B2|SLC39A10|SLIT3|SMARCA5|SMARCC1|SMC1A|SNRNP27|SNRPD2|SSB|SUMO1|TARS|TBCB|TCF12|TFDP2|TKT|TMA7|TMSB15A|TOMM22|U2SURP|UBA52|UQCRC1|XRCC5|YBX1|ZC3H15|ZNF148</td><td>1.56E-37</td><td>**</td></tr>
<tr><td>NOR (nC9)</td><td>Low/Intermediate risk</td><td>121</td><td>AASDHPPT|ABHD2|ACVR2B|AFAP1|ALCAM|ARL6IP5|ASTN1|ATF2|ATP11A|ATP2C1|BCL2|CAMK2N1|CAMK4|CCDC82|CD47|CHGB|CHST1|CLCN3|CLIP3|CNTN1|CPE|CPNE8|CREG2|CSRNP3|DDAH1|DDX6|DLEU2|DPYSL3|DSEL|EHBP1|FAM171A1|FBXO30|FMN1|FMO1|FOXP1|FSTL5|GAB2|GNAO1|GPR22|HIBCH|IGFBP5|INO80D|INPP5F|IRS2|JARID2|JPH3|JUN|KCNK3|KCNQ3|KIDINS220|LANCL1|LIFR|LONRF2|LRRN3|MAP4|MARCH4|MBNL2|MEIS2|MST1|MTMR6|MXRA7|NAALAD2|NRP1|OPTN|OXCT1|PBRM1|PCDH10|PCDH7|PCDHB10|PCDHB16|PCDHB6|PGM2L1|PHYHIPL|PITPNB|PJA2|PKIB|PLCXD3|PMP22|PREPL|PROX1|RAB3C|RBMS1|RBMS3|RGS5|RND3|RNF144A|RUFY3|SCG2|SCG5|SCN7A|SCRN1|SEMA6D|SESTD1|SEZ6|SEZ6L|SFRP1|SLC1A4|SLC25A12|SLC6A2|SMARCA1|SPATS2L|ST6GALNAC5|STAC|STMN4|SV2C|SYP|SYT1|SYT13|SYT4|TIAL1|TMCO3|TMEM255A|TMOD2|TNRC6B|TRIM13|TSHZ2|VAT1L|VPS36|ZBTB38|ZC3H13|ZDBF2</td><td>8.57E-29</td><td>**</td></tr>
</table>

<table>
<tr><td>Undifferentiated (nC3)</td><td>High risk</td><td>218</td><td>ABCA12|ABCC4|AC073464.10|ACTR3C|ADAM29|ADAMTS19|ADAMTS20|AFF2|ALG10B|ANAPC1|ANKRD20A1|ANKRD20A3|ANKRD20A4|ANKRD30B|ANKRD31|ANKRD45|ANO3|ANTXR2|AP5M1|ARHGEF38|ARMC3|ASPM|ATAD2|BMS1|BRCA1|BRCA2|BRIP1|BRMS1L|BUB1|C12orf48|C14orf145|CCDC15|CCDC150|CCDC168|CCDC169|CCSER1|CDC27|CECR2|CENPE|CEP112|CEP152|CHEK2|CISD2|CLSPN|CMC2|CPSF2|CRB1|CSMD3|CSRP2|CTCFL|DCDC1|DDX20|DEPDC1|DET1|DIAPH3|DLEU1|DLX6-AS1|DNAH11|DNAH14|DNAH3|DNAH6|DPY19L3|E2F6|EFCAB3|ELMOD1|ELP4|EML1|ESCO2|ETV1|EXOC5|EXPH5|FAM13A|FAM153A|FAM153B|FAM227A|FANCD2|FANCI|FANCM|G2E3|GABRG3|GALK2|GEMIN5|GEN1|GLDC|GNAI3|GPAM|GRM5|GULP1|HAPLN1|HEATR4|HHIP|IDE|ILDR2|INTS2|INTS7|INTU|IPO11|KIAA0586|KIAA1257|KIF14|KIF15|KIF6|KNTC1|L2HGDH|LGR4|LGR5|LMO3|LNPEP|LRRC9|MAP2K6|MCCC2|MDN1|MED12L|MET|MMS22L|MTHFD2L|MYB|MYO3A|NCAPG2|NEDD1|NEK10|NIPAL3|NMNAT3|NOX4|NT5DC3|NUBPL|NUP133|NUP155|NUP37|NUP93|OR4N2|PCAT1|PEX5L|PGGT1B|PIAS2|PIK3R4|PKHD1L1|PLCE1|PLD5|POLQ|POLR3B|POLR3G|POU6F2|PPAT|PRDM5|PRDM7|PRIM2|PRRC1|PTCD2|PTPN13|RAD51B|RANBP17|RBBP8|RBMY1A1|RBMY1B|RFX6|RMST|RNASEH2B|RNF217|RPS6KA5|RPS6KB1|RTTN|SATB2|SCIN|SEC23A|SEMA5A|SHQ1|SLC35G1|SLC38A9|SLC5A12|SLC9B1|SLCO1A2|SMC6|SNTG1|SOX2-OT|SOX5|SPATA17|SPDYA|SPECC1|SPEF2|SPINK5|STIL|STRIP2|STXBP6|STYK1|TAF4B|TMPRSS15|TRDN|TRIM64B|TTC26|TTC5|TTLL5|ULK4|USP54|UTP20|WDHD1|WDR3|WDR72|WRN|XPNPEP3|ZBBX|ZDHHC13|ZFAND4|ZNF169|ZNF215|ZNF285|ZNF385B|ZNF426|ZNF479|ZNF519|ZNF527|ZNF546|ZNF578|ZNF595|ZNF678|ZNF716|ZNF736|ZRANB3</td><td>3.46E-27</td><td>**</td></tr>
<tr><td>Undifferentiated (nC3)</td><td>Low/Intermediate risk</td><td>246</td><td>A2ML1|AASS|ABCA10|ABCA13|ABCA6|ABCA8|ABCA9|ABCB4|ABCC9|ACSS3|ADAMTSL3|ADCY10|ADCY2|AGBL3|AK5|AKAP7|ALDH1L2|AMY1A|AMY1B|AMY1C|AMY2A|ANKS1B|ANO4|APAF1|ARHGAP24|ARHGAP28|ARHGAP42|ARHGEF12|ASAH2|ASPA|ATP10B|ATP13A4|ATP8B4|BCO2|C6|CACNB4|CADPS2|CASC1|CBWD5|CC2D2B|CCDC122|CCDC146|CCDC30|CD109|CD226|CDH10|CDH13|CDH18|CDH9|CDK14|CDON|CEPT1|CLN5|CLNK|CNTN3|CNTN5|CNTNAP3|COL14A1|COL21A1|COL28A1|COL4A3|COL4A4|COL6A6|CPD|CPT2|CR1|CRISPLD1|CUBN|CYP3A5|DAB1|DBT|DDX4|DDX60|DENND2C|DEPDC5|DIO2|DMGDH|DNAJC16|DOCK7|DOCK9|DSC1|DSC2|DYNC1I1|DYNC2H1|EFCAB5|EHF|ENPP1|ENPP3|ENTPD5|EPM2A|F5|FAM135A|FAT3|FBN2|FBXO3|FGGY|FLRT2|FLT3|FNDC1|FREM1|FREM2|FRK|FSIP2|GABRA1|GABRG2|GDA|GDPD4|GK5|GLI3|GRAMD1C|GRID2|GUCY1A2|GUCY2C|HGF|HMCN1|HSFY2|HULC|HYDIN|IGFN1|IL1RAP|IMPG1|INPP4B|IPCEF1|IQUB|ITGA8|KIAA0825|KIAA1328|KIF13A|KLRD1|LAMA2|LAMA3|LAMB4|LEPR|LINC00466|LMBR1|LMO7|LRMP|LRP1B|LRP2|LRRK1|LRRK2|MBOAT1|MCM9|MR1|MROH2B|MROH8|MYH15|MYO1H|MYOM1|NAALADL2|NAV3|NEB|NECAB1|NEK5|NF1|NLGN1|NPAS3|NPHP1|NRXN3|NSF|NUP210L|OSBPL10|OTOA|PARP11|PART1|PCCA|PCDHB1|PCSK5|PDE1A|PDE4DIP|PDZD2|PGR|PHLPP1|PHLPP2|PIEZO2|PIGK|PIP5K1B|PIWIL2|PKHD1|PLA2G4A|PLA2R1|PRKAA2|PTGFR|PTPRG|PTPRT|PTPRZ1|PZP|RGPD3|RGPD4|RGPD5|RGPD6|RGPD8|RGSL1|RORB|RP1-14N1.2|RP11-217E22.2|RP11-327J17.3|RP11-478J18.1|RP11-550E22.3|RYR3|SAR1B|SCN11A|SCO1|SEMA3E|SERPINB8|SH3TC2|SLC12A1|SLC15A2|SLC2A13|SLC4A4|SLCO4C1|SLMAP|SNAP25-AS1|SOCS7|SPHKAP|SPTLC3|SSX2IP|STEAP4|STK3|STK31|SUMF1|SVEP1|SYNE1|SYTL2|TAOK1|TBC1D8B|TF|THEMIS|THRB|THSD7B|TMC1|TMEM232|TP63|TPK1|TPTE2|TRPM3|TYW5|USP32|VEPH1|VGLL3|VPS13D|WDR66|WDR7|WNT2B|XKR4|ZBED3-AS1</td><td>3.06E-25</td><td>**</td></tr>
</table>

**Human NB to Age at diagnosis in 498 NBs (Fig 6)**

<table>
<tr><th>Cell cluster in human NB</th><th>Correlation with age at diagnosis in 498 SEQC NB samples</th><th>Number of genes in the specific signature of NB significantly correlated with age at diagnosis in 498 SEQC NB samples</th><th>Genes in the specific signature of NB significantly correlated with age at diagnosis in 498 SEQC NB samples</th><th>FDR</th><th>Significance</th></tr>
<tr><td>Endothelial (nC4)</td><td>Inversely correlated</td><td>71</td><td>ADCY4|ANXA2|APLNR|APOLD1|ARHGEF15|BCL6B|C2CD4B|CA2|CALCRL|CBLB|CD34|CD93|CDH5|CLEC14A|CMIP|CNKSR3|COL15A1|COL4A1|CRIP2|ECE1|EPAS1|EPHA2|EPHB4|ESAM|ETS1|ETS2|FAM129B|FLT4|FOXO1|FZD4|GRB10|HSPG2|INSR|ITPKC|JMJD1C|KIAA0355|LAMA5|MARCH3|MSN|MYH9|NEDD9|NOTCH1|PALMD|PCDH12|PDGFB|PIEZO1|PLAT|PLEC|PLS3|PODXL|PPM1F|RASIP1|RBPMS|ROBO4|RP11-14D22.5|S100A16|SASH1|SCARF1|SHANK3|SHROOM4|SKI|SLC35E4|SNRK|SOX17|SPRY1|SSH1|TMEM173|TMEM204|TMOD3|TNFRSF10D|TNKS1BP1</td><td>5.79E-20</td><td>**</td></tr>
<tr><td>Endothelial (nC4)</td><td>Directly correlated</td><td>36</td><td>ARL2|ATG9B|BAG5|BMP6|CCDC85B|CDC37|CDC42EP2|CDC42EP3|CENPB|CPLX1|DDX21|EIF5B|EVA1C|FBL|FKBP1A|FZD6|GAR1|GAS5|GRPEL1|IFI27|LCN6|MTA2|NIP7|NRGN|PABPC1|PABPC3|PARVB|PNP|RPGR|RRP12|SLCO4A1|STC1|TEAD4|TMEM70|TNFRSF10B|TPM3</td><td>3.69E-03</td><td>**</td></tr>
</table>

| | | | | | |
|---|---|---|---|---|---|
| Macrophages (nC6) | Inversely correlated | 108 | ABCC3\|APOE\|ARL4C\|ARRB2\|ATG16L2\|BMF\|CD14\|CD300LF\|CD84\|CD86\|CELF2\|CLEC7A\|CSF1R\|CSF2RA\|CTSH\|CTSZ\|CXCL16\|CYBB\|DAB2\|DENND3\|DMXL2\|EEPD1\|ELL2\|FCHO2\|FERMT3\|FGD2\|FNIP2\|FPR3\|GABARAPL1\|GPCPD1\|GPNMB\|GRAMD4\|HAVCR2\|HCK\|HCLS1\|HLA-DMB\|HLA-DQB2\|IFNGR1\|IL18\|IL6R\|IRF8\|ITGAD\|ITGAM\|ITGAX\|KCNAB2\|KLHL6\|KMO\|LACTB\|LAIR1\|LILRB4\|LRP1\|MAFB\|MERTK\|MFSD12\|MKNK1\|MPEG1\|MS4A6A\|MXD1\|MYO1F\|MYO9B\|NBPF10\|NPL\|PAPSS2\|PLA2G7\|PLEK\|PLXDC2\|PPARD\|PREX1\|PRKAG2\|RAB20\|RAB7B\|RASGEF1B\|RASSF2\|RASSF4\|RBM47\|RIN3\|RNF135\|RNF144B\|RREB1\|SDC3\|SETDB2\|SGK1\|SLC15A3\|SLC1A3\|SLC31A2\|SLC37A2\|SLC43A2\|SLCO2B1\|SMAP2\|SMPDL3A\|SNHG12\|SPI1\|STX7\|STXBP2\|TBC1D12\|TBC1D2\|TFEC\|TGFBI\|TMEM63A\|TNFRSF14\|TNFRSF1B\|TNS3\|TOM1\|TRPM2\|WDR91\|ZEB2\|ZFP36L1\|ZMYND15 | 2.61E-43 | ** |
| NOR (nC7) | Inversely correlated | 187 | ACAD10\|ACAP3\|ADCY1\|AGAP2\|AGER\|AGRN\|AHI1\|AMIGO2\|ANKRD12\|ANKRD13B\|ANKRD36\|ANKRD36B\|ANKS1A\|APC2\|ARHGEF10L\|ARID1B\|B4GALNT4\|BMPR1B\|BRSK2\|BTBD3\|C1QTNF2\|C2CD4A\|CALCOCO1\|CAMK2B\|CASZ1\|CCNL2\|CHD5\|CHD9\|CIC\|CNTFR\|CORO6\|CRLF3\|DBH\|DDC\|DDX17\|DDX39B\|DIRAS3\|DLC1\|DMTF1\|DNAJB2\|DNM1\|DOC2B\|DOCK4\|DPP6\|DST\|EBF1\|ECEL1\|ELAVL2\|EPB41L4A\|EPHB3\|ERC2\|ERV3-1\|EVL\|EYA4\|FAM184A\|GATA2\|GCGR\|GFRA3\|GINM1\|GLG1\|GNAS\|GNB3\|GPATCH8\|GPRASP1\|GRIK2\|GTF2I\|GTF2IRD1\|GTF2IRD2\|GTF2IRD2B\|HBG1\|HECTD4\|HID1\|HIP1R\|HMX1\|HS3ST5\|ING4\|INSRR\|KB-1572G7.1\|KCNQ5\|KDM1A\|KLC4\|KLRK1\|LAMA4\|LMBR1L\|LUC7L\|LUC7L2\|LUC7L3\|MAPK8IP3\|MAPT\|MAST1\|MC1R\|MDGA1\|MED23\|MGAT4B\|MLH1\|MLXIP\|NCAM1\|NDRG4\|NDUFA4L2\|NINL\|NISCH\|NKTR\|NOS1AP\|NRCAM\|NTRK1\|NXPH4\|OBSL1\|OCLN\|PCDHA9\|PCLO\|PDE9A\|PEX6\|PHF19\|PHIP\|PI4KA\|PKD1\|PLAGL1\|PLXNA4\|PLXNB1\|PNISR\|POU2F2\|PPIP5K2\|PRPF4B\|PRPSAP2\|PRRC2B\|PRRT2\|PSD2\|PTBP2\|PTPRF\|PXDN\|R3HDM2\|RADIL\|RAI1\|RBFOX1\|RBM33\|RBM6\|RERE\|RGMB\|RGS7\|RIMKLB\|SDCCAG8\|SFSWAP\|SGSM2\|SHC2\|SLC18A1\|SMARCD1\|SMPD3\|SNAP91\|SOBP\|SOGA3\|SOX4\|SPTAN1\|SRGAP3\|SRRM1\|SRRM3\|SRRM4\|SRSF11\|STAG3\|STK38\|STMN1\|STMN2\|STXBP1\|TBC1D24\|TFAP2B\|TH\|TIA1\|TLE4\|TMEM191C\|TRIM33\|TSTD2\|TUBA1A\|TXNIP\|UBAP1L\|ULK1\|UNC5A\|WDR6\|WDR86\|WSB1\|YJEFN3\|ZCCHC14\|ZFHX3\|ZNF117\|ZNF33A\|ZNF429\|ZNF512B\|ZNF608\|ZRANB2 | <1E-50 | ** |
| NOR (nC8) | Inversely correlated | 46 | ANKRD6\|BASP1\|CCDC68\|CHRFAM7A\|CREB5\|CS\|DCLK1\|DCLK3\|DUSP26\|EPB41L1\|EPB41L4B\|EYA1\|FAM219A\|GNB1\|HMGN3\|IFT20\|ITGB8\|JUP\|KBTBD6\|MAPRE2\|NXPH1\|ONECUT2\|PDE10A\|PPP4R1\|PTN\|QPCT\|RAB15\|RGS3\|RIN2\|RNF10\|RUNX1T1\|RUSC2\|SNRPN\|SPIN1\|SV2B\|SYBU\|THRA\|THSD7A\|TM2D3\|TMBIM4\|TMEFF2\|TMEM163\|TUBB2A\|TUBB4A\|VSNL1\|WDR47 | 2.60E-14 | ** |
| NOR (nC8) | Directly correlated | 35 | ARL16\|ASB6\|ATP6V1G1\|CHMP5\|CLTA\|CNOT7\|COX6C\|CRABP1\|CUTA\|EBF3\|EBP\|GAL\|KIT\|MRFAP1L1\|MRPS21\|MRPS26\|NDUFB9\|NHLH2\|NPTX2\|OPRM1\|PDZD8\|PNMA1\|PSMA7\|PSMC2\|PTPN3\|SCOC\|SEC61B\|SPTLC1\|SSBP1\|ST8SIA3\|SURF4\|TMED2\|TMX2\|TUBB4B\|TXN | 3.41E-07 | ** |
| NOR (nC9) | Inversely correlated | 109 | AASDHPPT\|ACVR2B\|AFAP1\|ALCAM\|ARHGAP35\|ARL6IP5\|ASTN1\|ATF2\|ATP11A\|BCL2\|CCDC82\|CD47\|CHST1\|CNTN1\|CPNE8\|CREG2\|CSRNP3\|CYGB\|DDAH1\|DDX6\|DLEU2\|DPYSL3\|DSEL\|DUSP8\|EHBP1\|EIF1B\|FAM171A1\|FANCL\|FMN1\|FOXP1\|FSTL5\|GAB2\|GIGYF2\|GNAO1\|H1F0\|HIBCH\|INO80D\|IRS2\|JARID2\|JPH3\|JUN\|KCNK3\|KCNQ3\|KIAA1549L\|KIDINS220\|KIF26B\|LIFR\|LONRF2\|LRRN3\|LRRTM2\|MARCH4\|MBNL2\|MEIS2\|MRPS5\|MST1\|MTMR6\|MXRA7\|NAALAD2\|NRP1\|OPTN\|OXCT1\|PARD3\|PBRM1\|PCDH10\|PCDH7\|PCDHB16\|PGM2L1\|PITPNB\|PJA2\|PKIB\|PMP22\|PODXL2\|RBMS1\|RBMS3\|RGS5\|RND3\|RNF144A\|SCG2\|SCG5\|SCN7A\|SCRN1\|SEMA6D\|SEPT6\|SESTD1\|SETD2\|SEZ6\|SFRP1\|SLC25A12\|SPATS2L\|ST3GAL6\|ST6GALNAC5\|STAC\|STMN4\|SYT13\|TAGLN3\|TET3\|TIAL1\|TLE3\|TMCO3\|TMEM255A\|TMOD2\|TNRC6B\|TOP2B\|TRIM13\|TSHZ2\|XRN1\|ZBTB38\|ZC3H13\|ZDBF2 | <1E-50 | ** |
| NOR (nC9) | Directly correlated | 44 | ALKBH7\|BMP7\|COPE\|COX6B1\|COX7C\|COX8A\|DPY30\|EEF1B2\|FAU\|GCSH\|H2AFZ\|HDGF\|HINT1\|HIST1H4E\|HMGB2\|HNRNPL\|HSD17B11\|HSD17B12\|HTATSF1\|JTB\|MIR3654\|NDUFA1\|NDUFB11\|OST4\|RMND5A\|RPL18A\|RPL31\|RPL34\|RPL35A\|RPS10\|RPS11\|RPS13\|RPS15\|RPS15A\|RPS19\|RPS21\|RPS3\|RPS7\|SF3B2\|SMARCA5\|SNRNP27\|SNRPD2\|TOMM22\|UBA52 | 2.42E-05 | ** |
| Undifferentiated (nC2) | Directly correlated | 5 | ALOXE3\|CDC45\|OTOP2\|SPATA31A3\|VAX1 | 2.53E-02 | * |
| Undifferentiated (nC3) | Directly correlated | 175 | ABCA12\|ABCC4\|AC073464.10\|ACOXL\|ACTR3B\|ACTR3C\|ADAM29\|ADAM7\|ADAMTS19\|ADAMTS20\|AFF2\|AGBL1\|ANKRD20A1\|ANKRD20A2\|ANKRD20A3\|ANKRD20A4\|ANKRD45\|ANKRD7\|ANO3\|AP5M1\|ARMC10\|ARMC3\|BRCA1\|BRMS1L\|C14orf145\|CCDC169\|CCSER1\|CDC14B\|CECR2\|CEP112\|CEP152\|CISD2\|CMC2\|CNTN4\|CNTN6\|COL24A1\|CPSF2\|CRB1\|CRISPLD1\|CSMD3\|CTNNA3\|DCDC1\|DCHS2\|DLX6-AS1\|DNAH10\|DNAH11\|DNAH14\|DNAH3\|DNAH5\|DNAH6\|DNAH7\|DNAH8\|DPY19L3\|EFCAB6\|ELMOD1\|ELP4\|EML1\|ERBB4\|ESCO2\|EXOC5\|FAM153A\|FAM153B\|FAM197Y5\|FAM227A\|FANCI\|FANCM\|FBXO15\|G2E3\|GABRA6\|GABRG3\|GALK2\|GEMIN5\|GRIN2B\|HEATR4\|HERC5\|ILDR2\|INTS2\|INTU\|IPO11\|KIAA1257\|KIF6\|KNTC1\|L2HGDH\|LGI1\|LGR5\|LINC00467\|LMO3\|LRRC9\|LRRIQ3\|MATN3\|MCCC2\|MET\|MGAM\|MTHFD2L\|MUC12\|MUC3A\|MYH13\|MYH4\|MYO3B\|NEK10\|NLRP14\|NRXN3\|NT5DC3\|NUBPL\|NUP155\|NUP37\|NUP93\|OR4N2\|PGGT1B\|PIAS2\|PIK3R4\|PIWIL1\|PKD1L1\|PKHD1L1\|POLR3G\|POTEC\|POU6F2\|PPAT\|PRDM5\|PRDM7\|PRIM2\|PTCD2\|RAD51B\|RBBP8\|RBMY1A1\|RBMY1B\|RBMY1D\|RBMY1E\|RBMY1F\|RBMY1J\|RFX6\|RGS22\|RMST\|RP11-217E22.2\|RP11-373D17.1\|SATB2\|SEC23A\|SHQ1\|SLC13A1\|SLC35G1\|SLC5A12\|SLC5A8\|SLC9B1\|SLC9C2\|SLCO1A2\|SLCO6A1\|SMC1B\|SNTG1\|SPAG17\|SPATA17\|SPECC1\|SPINK5\|STRIP2\|TEX14\|TMEM232\|TRDN\|TSHR\|TTC26\|TTC5\|TTLL5\|TYR\|TYRP1\|ULK4\|UTP20\|WDR3\|WDR35\|WDR72\|ZBBX\|ZDHHC13\|ZFAND4\|ZNF169\|ZNF215\|ZNF479\|ZNF595\|ZNF716 | 8.12E-31 | ** |

| | | | | | |
|---|---|---|---|---|---|
| Undifferentiated (nC3) | Inversely correlated | 109 | A2ML1\|ABCA10\|ABCA6\|ABCA9\|ABCB4\|ACSS3\|ADAMTS12\|ADCY10\|AGBL3\|AK5\|AKAP7\|ALDH1L2\|AMY1A\|AMY1B\|AMY1C\|AMY2A\|ANO4\|APAF1\|ARHGAP28\|ARHGAP42\|ARHGEF12\|ATP13A4\|ATRNL1\|BCO2\|CC2D2B\|CD109\|CDH9\|CDON\|CNTN5\|COL21A1\|COL6A6\|CUBN\|DAB1\|DEPDC5\|DSC2\|EFCAB5\|EHF\|ENPP1\|FAM135A\|FAT3\|FBN2\|FLRT2\|FNDC1\|FREM1\|GDPD4\|GK5\|GLI3\|GRID2\|GUCY1A2\|HGF\|HMGA2\|IGFN1\|IL1RAP\|IMPG1\|INPP4B\|ITGA8\|KIAA1328\|KIF13A\|LAMA1\|LAMA2\|LAMA3\|LAMB4\|LMO7\|LRP1B\|MCM9\|MGST1\|MROH8\|NAALADL2\|NAV3\|NEB\|NF1\|NPAS3\|NUP210L\|PARP11\|PCSK5\|PDE1A\|PDE4DIP\|PDZD2\|PHLPP1\|PIP5K1B\|PLA2G4A\|POU2F1\|PRKAA2\|PRTG\|PTPRZ1\|RGPD3\|RGPD5\|RGPD6\|RGPD8\|RORB\|RP11-327J17.3\|SEMA3E\|SH3TC2\|SLC12A1\|SLC1A2\|SLCO4C1\|SOCS7\|STK31\|SYTL2\|TAOK1\|TBC1D8B\|TF\|THRB\|TYW5\|VGLL3\|VPS13D\|WDR7\|WNT2B\|ZBED3-AS1 | 5.69E-08 | ** |

**Human NB to Survival in 498 NBs (Fig 6)**

| Cell cluster in human NB | Survival in 498 SEQC NB samples | Number of genes in the specific signature of NB significantly up-regulated in survival group in 498 SEQC NB samples | Genes in the specific signature of NB significantly up-regulated in survival group in 498 SEQC NB samples | FDR | Significance |
|---|---|---|---|---|---|
| Endothelial (nC4) | Better survival | 70 | ADCY4\|ANXA2\|APLNR\|APP\|ARHGEF15\|BCL3\|BHLHE40\|C2CD4B\|C6orf208\|CCDC69\|CD93\|CMIP\|CNKSR3\|COL15A1\|CRIP2\|DOCK6\|DUSP6\|ECE1\|ENDOD1\|EPAS1\|EPHA2\|ETS1\|ETS2\|FAM129B\|FLT1\|FOXO1\|FXYD5\|GABRE\|GNAI2\|GRASP\|GRB10\|HES1\|HIPK3\|HLA-E\|HSPG2\|IL6ST\|KIAA0355\|KLF2\|LAMA5\|LMNA\|MARCH3\|MSN\|NDRG1\|NOTCH1\|PDGFB\|PIEZO1\|PLAT\|PLEC\|PLEKHG2\|PODXL\|PXN\|RAMP3\|RAPGEF3\|RCAN1\|RNF19B\|RP11-314N13.4\|SCARF1\|SHANK3\|SHROOM4\|SKI\|SLC35E4\|SNRK\|SOX7\|SPRY1\|SSH1\|TGFB1\|TGFBR2\|TMEM173\|UPP1\|VSIG2 | 1.16E-11 | ** |
| Macrophages (nC6) | Better survival | 169 | ABCC3\|ADAM8\|ALOX5\|AMDHD2\|AMPD2\|APBB1IP\|APBB3\|APOE\|ARHGAP18\|ARL8B\|ARRB2\|ASAH1\|ATG16L2\|BCL2A1\|BHLHE41\|BMF\|BTK\|CD14\|CD300LF\|CD58\|CD74\|CD83\|CD84\|CD86\|CELF2\|CIITA\|CLEC7A\|CSF1R\|CSF2RA\|CTSH\|CTSS\|CTSZ\|CXCL16\|CYBB\|CYTH4\|DAB2\|DENND3\|DMXL2\|DOK3\|ELL2\|FCHO2\|FERMT3\|FGD2\|FGR\|GAB3\|GABARAPL1\|GPCPD1\|GPNMB\|GPR183\|HAVCR2\|HCK\|HCLS1\|HERPUD1\|HEXA\|HK3\|HLA-DMA\|HLA-DMB\|HLA-DPA1\|HLA-DPB1\|HLA-DQA2\|HLA-DQB2\|HLA-DRA\|HLA-DRB1\|IFFO1\|IL18\|IL6R\|IRF8\|ITGAD\|ITGAM\|ITGAX\|KCNAB2\|KLHL6\|LACTB\|LAIR1\|LAPTM5\|LCP2\|LGALS9\|LILRB1\|LILRB3\|LILRB4\|LRP1\|MAFB\|MAP3K8\|MFSD1\|MFSD12\|MGLL\|MKNK1\|MPEG1\|MS4A6A\|MXD1\|MYO1F\|MYO9B\|NBPF10\|NCF2\|NCF4\|NEAT1\|NPL\|NR1H3\|NR4A2\|P2RX7\|PAPSS2\|PARVG\|PDE4A\|PIK3R5\|PLA2G7\|PLCB2\|PLEK\|PPARD\|PREX1\|PRKAG2\|PSAP\|PTK2B\|PTPN6\|RAB20\|RAB31\|RAB7B\|RAPGEF1\|RASGEF1B\|RASSF2\|RASSF4\|RASSF5\|RBM47\|RGS1\|RHBDF2\|RIN3\|RNF135\|RNF144B\|RREB1\|SCARNA9\|SDC3\|SEMA4A\|SETDB2\|SGK1\|SLC15A3\|SLC17A9\|SLC1A3\|SLC31A2\|SLC37A2\|SLC43A2\|SLCO2B1\|SMAP2\|SMPDL3A\|SNHG12\|SNORD99\|SNX29\|SPI1\|STX7\|STXBP2\|SYNGR2\|TBC1D12\|TBC1D2\|TFEC\|THEMIS2\|TLR2\|TMEM63A\|TNFAIP2\|TNFRSF14\|TNFRSF1B\|TOM1\|TPCN1\|TRIM38\|TRPM2\|UAP1L1\|VAMP8\|WDR91\|ZEB2\|ZFP36\|ZFP36L1\|ZMYND15 | <1E-50 | ** |
| NOR (nC7) | Better survival | 197 | ACAD10\|ACAP3\|ADCY1\|AGAP2\|AGAP3\|AGER\|AGRN\|AHI1\|AKAP6\|AMIGO2\|ANKRD12\|ANKRD13B\|ANKRD36\|ANKRD36B\|ANKRD36C\|ANKS1A\|APC2\|ARHGEF10L\|ARID1B\|ASIC3\|ATF6B\|ATG14\|B4GALNT4\|BMPR1B\|BRSK2\|BTBD3\|C1QTNF2\|C2CD4A\|CALCOCO1\|CAMK2B\|CASZ1\|CC2D1A\|CCDC136\|CCNL2\|CDK5RAP3\|CHD5\|CIC\|CNTFR\|CORO6\|CRLF3\|CYB561\|DBH\|DDC\|DDX17\|DIRAS3\|DLC1\|DMTF1\|DNAJB2\|DNM1\|DOC2B\|DOCK4\|DPP6\|DST\|EBF1\|ECEL1\|ELAVL2\|EPB41L4A\|EPHB3\|ERC2\|ERV3-1\|EXOC4\|EYA4\|FAM120B\|FAM184A\|FAM193B\|FASTK\|FNDC5\|FTX\|GCGR\|GFRA3\|GIGYF1\|GINM1\|GLG1\|GNAS\|GNB3\|GPATCH8\|GPRASP1\|GRIK2\|GTF2I\|GTF2IRD1\|GTF2IRD2\|GTF2IRD2B\|HECTD4\|HID1\|HIP1R\|HMX1\|HS3ST5\|ING4\|INSRR\|KB-1572G7.1\|KCNQ5\|KDM5D\|KLC4\|KLRK1\|LAMA4\|LMBR1L\|LUC7L2\|LUC7L3\|MAPK8IP3\|MAPT\|MAST1\|MC1R\|MDGA1\|MED23\|MEG3\|MEIS3\|MGAT4B\|MLXIP\|MUC20\|NCAM1\|NCRNA00107\|NDRG4\|NINL\|NISCH\|NKTR\|NOS1AP\|NRCAM\|NTRK1\|OBSL1\|OCLN\|PACS2\|PCDHA9\|PCLO\|PDE9A\|PEX6\|PHC1\|PI4KA\|PKD1\|PLAGL1\|PLXNA4\|PLXNB1\|PNISR\|POLR2J2\|POU2F2\|PPIP5K2\|PRRT2\|PSD2\|PTBP2\|PTPRF\|PXDN\|R3HDM2\|RADIL\|RAI1\|RBCK1\|RBFOX1\|RBM33\|RERE\|RGMB\|RGS7\|RUNDC3A\|SDCCAG8\|SEMA6C\|SFSWAP\|SGSM2\|SHC2\|SLC18A1\|SNAP91\|SOBP\|SOGA3\|SOX4\|SPG7\|SPTAN1\|SRGAP3\|SRRM1\|SRRM3\|SRRM4\|SRSF11\|STAG3\|STK38\|STMN2\|STXBP1\|TBC1D24\|TBKBP1\|TFAP2B\|TH\|TIA1\|TLE4\|TMEM191C\|TSC2\|TSTD2\|TUBA1A\|TXNIP\|UBAP1L\|ULK1\|UNC5A\|VPS9D1\|WDR6\|WDR86\|WSB1\|YJEFN3\|ZCCHC14\|ZFHX3\|ZNF117\|ZNF33A\|ZNF512B\|ZNF608\|ZRANB2 | <1E-50 | ** |
| NOR (nC8) | Better survival | 76 | ANKRD6\|ASIC1\|ATP1B1\|ATP6V0E2\|BASP1\|CCDC28A\|CCDC68\|CHRFAM7A\|CHRNA3\|CREB5\|DCLK1\|DCLK3\|DKK3\|DUSP26\|ELMO2\|EPB41L1\|EPB41L4B\|FAM219A\|FECH\|FRMD3\|FRRS1L\|GNB1\|HDAC9\|HMGN3\|HTR3A\|IFT20\|ISCU\|ITGB8\|ITM2C\|JUP\|KBTBD6\|KLHL35\|MAPRE2\|NDRG3\|NXPH1\|OGFRL1\|PCSK1N\|PCSK2\|PDE10A\|PHF1\|PTN\|QPCT\|RAB14\|RAB15\|RAB2A\|RGS3\|RGS4\|RHBDD2\|RIN2\|RNF10\|RUNX1T1\|RUSC2\|SERINC3\|SH3GLB2\|SLC25A4\|SMAD9\|SNRPN\|SORCS1\|SPIN1\|SV2B\|SYBU\|THRA\|THSD7A\|TM2D3\|TMBIM4\|TMED4\|TMEFF2\|TMEM163\|TRPA1\|TUBB2A\|TUBB4A\|VPS4B\|VSNL1\|WDR47\|WRB\|WSB2 | 2.44E-28 | ** |
| NOR (nC8) | Poor survival | 40 | ARL16\|CLTA\|CNOT7\|COX6C\|CRABP1\|CRNDE\|CS\|CUTA\|CYC1\|DACH1\|DLK1\|EBP\|EDF1\|GAL\|KIT\|LRRC42\|MRPS21\|MRPS26\|NDUFA12\|NDUFB9\|NFIB\|NHLH2\|NPTX2\|PDZD8\|PNMA1\|PSMA7\|SEC61B\|SLC10A4\|SSBP1\|STRAP\|SURF4\|TMED2\|TMX2\|TPI1\|TUBA1C\|TUBB4B\|TUBG1\|TXN\|WDR61\|ZNF770 | 1.16E-02 | * |

| | | | | | |
|---|---|---|---|---|---|
| NOR (nC9) | Poor survival | 114 | ALKBH7\|ATRX\|BEX2\|BMP7\|BRK1\|BSG\|BTF3\|CCT4\|CCT7\|CENPJ\|CKS1B\|COPE\|COX6B1\|COX7C\|COX8A\|DPY30\|DUT\|EEF1B2\|EEF2\|EIF2AK2\|EIF3L\|EPRS\|ESD\|FAU\|FXR1\|GCSH\|GSTP1\|H1F0\|H2AFZ\|HADHA\|HDGF\|HINT1\|HIST1H4E\|HMGA1\|HMGB1\|HMGB2\|HNRNPL\|HSD17B11\|HSD17B12\|HSPA8\|HTATSF1\|JTB\|KLHL23\|LDHB\|METTL9\|MIR3654\|MTHFD2\|N4BP2\|NDUFA1\|NDUFB11\|OIP5-AS1\|OST4\|PFN2\|POLR1D\|PRDX2\|RMND5A\|RPL14\|RPL15\|RPL18\|RPL18A\|RPL23A\|RPL24\|RPL26\|RPL27\|RPL29\|RPL31\|RPL32\|RPL34\|RPL35A\|RPL39L\|RPL4\|RPS10\|RPS11\|RPS13\|RPS15\|RPS15A\|RPS19\|RPS21\|RPS24\|RPS27A\|RPS3\|RPS5\|RPS7\|RPS9\|RPSA\|RSL24D1\|SACS\|SAE1\|SEPT11\|SF3B2\|SLC39A10\|SLIT3\|SMARCA5\|SMARCC1\|SMC1A\|SNORD114-20\|SNORD114-23\|SNRNP27\|SNRPD2\|SSB\|SYNPO2\|TARS\|TBCB\|TCF12\|TEAD1\|TKT\|TMA7\|TMSB15A\|TOMM22\|U2SURP\|UBA52\|UQCRC1\|YBX1\|ZC3H15 | 1.25E-21 | ** |
| NOR (nC9) | Better survival | 89 | AFAP1\|ALCAM\|ARL6IP5\|ASTN1\|ATF2\|ATP11A\|ATP2C1\|CCDC82\|CD47\|CHST1\|CLIP3\|CNTN1\|CPE\|CPNE8\|CREG2\|CSRNP3\|CYGB\|DDAH1\|DDX6\|DLEU2\|DPYSL3\|DSEL\|DUSP8\|EHBP1\|EIF1B\|EPM2AIP1\|FMN1\|FOXP1\|FSTL5\|GAB2\|GNAO1\|IGFBP5\|INO80D\|INPP5F\|JARID2\|JPH3\|JUN\|KCNQ3\|KIDINS220\|LIFR\|LONRF2\|LRRN3\|MARCH4\|MBNL2\|MST1\|MTMR6\|MXRA7\|NAALAD2\|NRP1\|OPTN\|PCDH10\|PCDH7\|PCDHB10\|PCDHB16\|PCDHB6\|PGM2L1\|PJA2\|PKIB\|PMP22\|PREPL\|RBMS1\|RBMS3\|RGS5\|RNF144A\|SCG2\|SCG5\|SCN7A\|SCRN1\|SEMA6D\|SEZ6\|SEZ6L\|SFRP1\|SLC1A4\|SLC25A12\|ST6GALNAC5\|STMN4\|SYP\|SYT1\|SYT13\|SYT4\|TAGLN3\|TMCO3\|TMEM255A\|TMOD2\|TNRC6B\|TSHZ2\|TSPAN7\|VAT1L\|ZBTB38 | 6.32E-21 | ** |
| Undifferentiated (nC3) | Poor survival | 252 | ABCA12\|ABCC4\|ACSM2B\|ACSM3\|ACTR3C\|ADAM29\|ADAM7\|ADAMTS19\|ADAMTS20\|AFF2\|ALG10B\|ANAPC1\|ANKRD30A\|ANKRD30B\|ANKRD45\|ANKUB1\|ANTXR2\|AP5M1\|APOH\|ARHGEF38\|ARMC10\|ARMC3\|ASPM\|ATAD2\|B4GALNT2\|BARD1\|BMS1\|BRCA1\|BRCA2\|BRIP1\|BRMS1L\|BUB1\|C12orf48\|C14orf145\|C8A\|CC2D2A\|CCDC15\|CCDC150\|CCDC169\|CCSER1\|CECR2\|CENPE\|CEP112\|CEP152\|CFTR\|CHEK2\|CISD2\|CLSPN\|CMC2\|CPS1\|CPSF2\|CRB1\|CSMD3\|CSRP2\|CTCFL\|CYP2C9\|DCDC1\|DDX20\|DEPDC1\|DET1\|DIAPH3\|DLEU1\|DLX6-AS1\|DNAH11\|DNAH14\|DNAH3\|DPY19L3\|DR1\|DSG2\|DSG3\|E2F6\|EGF\|ELMOD1\|ELP4\|EML1\|ESCO2\|ESRP1\|ETV1\|EXOC5\|EXPH5\|FAM13A\|FAM197Y5\|FAM227A\|FANCD2\|FANCI\|FANCM\|FBXO15\|FRAS1\|G2E3\|GABRA6\|GALK2\|GEMIN5\|GEN1\|GLDC\|GLYATL1\|GNAI3\|GPAM\|GRIN2B\|GULP1\|HEATR4\|IDE\|INTS2\|INTS7\|INTU\|IPO11\|IYD\|KCNU1\|KIAA1257\|KIF14\|KIF15\|KIF6\|KNG1\|KNTC1\|L2HGDH\|LGR4\|LGR5\|LMO3\|LRRC9\|MAP2K6\|MCCC2\|MDN1\|MED12L\|MED17\|MET\|MIA2\|MMS22L\|MROH9\|MTHFD2L\|MUC16\|MUC17\|MYB\|MYH1\|MYH13\|MYH4\|NCAPG2\|NEDD1\|NNT\|NOX4\|NT5DC3\|NUBPL\|NUP133\|NUP155\|NUP37\|NUP93\|OR10K1\|OR1A1\|OR2AG1\|OR2AT4\|OR2M3\|OR2T12\|OR4D9\|OR4F4\|OR4F5\|OR4N2\|OR52A1\|OR56A1\|OR56A4\|OR5AS1\|OR8D1\|PCAT1\|PEX5L\|PGGT1B\|PIK3C2G\|PIK3R4\|POLQ\|POTEC\|POU6F2\|PPAT\|PRAMEF1\|PRAMEF13\|PRAMEF14\|PRAMEF25\|PRDM5\|PRDM7\|PRIM2\|PRRC1\|PSG11\|PTCD2\|PTPN13\|RAD51B\|RANBP17\|RBBP8\|RFX6\|RGS22\|RMST\|RNASEH2B\|RNF217\|ROS1\|RP1\|RTTN\|SATB2\|SCEL\|SEC23A\|SERPINB5\|SHQ1\|SI\|SLC35G1\|SLC5A12\|SLC5A8\|SLC9B1\|SLC9C2\|SLCO6A1\|SMC6\|SNTG1\|SPAG17\|SPATA17\|SPDYA\|SPECC1\|SPINK5\|ST18\|STIL\|STXBP6\|TAF4B\|TMPRSS15\|TRDN\|TRIM43\|TRIM43B\|TRIM48\|TRIM49\|TRIM49B\|TRIM51\|TRIM64\|TRIM64B\|TTC26\|TTC5\|TTLL5\|TYR\|TYRP1\|ULK4\|UNC13C\|UTP20\|WDHD1\|WDR3\|WDR35\|WDR72\|WRN\|XPNPEP3\|ZBBX\|ZDHHC13\|ZFAND4\|ZNF169\|ZNF215\|ZNF285\|ZNF426\|ZNF479\|ZNF519\|ZNF578\|ZNF595\|ZNF678\|ZNF705G\|ZNF716\|ZRANB3 | 3.04E-22 | ** |
| Undifferentiated (nC3) | Better survival | 165 | A2ML1\|ABCA10\|ABCA6\|ABCA9\|ABCB4\|ADAMTSL3\|ADCY2\|AGBL3\|AK5\|AKAP7\|ALDH1L2\|AMY1A\|AMY1B\|AMY1C\|AMY2A\|ANKS1B\|ANO4\|ARHGAP28\|ARHGAP42\|ARHGEF12\|ATP10B\|ATP13A4\|BCO2\|BEST3\|CASC1\|CC2D2B\|CCDC146\|CCDC30\|CD226\|CDH10\|CDH18\|CDH9\|CDON\|CEPT1\|CLN5\|CLNK\|CNTN3\|CNTNAP3\|COL21A1\|COL28A1\|COL4A4\|CR1\|CUBN\|DAB1\|DBT\|DDX4\|DDX60\|DNAJC16\|DOCK7\|DOCK9\|DSC2\|DYNC1I1\|EFCAB5\|EHF\|ENPP3\|EPM2A\|F5\|FAM135A\|FAT3\|FBXO3\|FLRT2\|FLT3\|FREM1\|FREM2\|FSIP2\|GDA\|GDPD4\|GK5\|GLI3\|GRID2\|GRIN2A\|HMCN1\|HMGA2\|HULC\|HYDIN\|IGFN1\|IL1RAP\|IMPG1\|INPP4B\|IPCEF1\|IQUB\|ITGA8\|KIAA0825\|KIAA1328\|KIF13A\|KLRD1\|LAMA3\|LAMB4\|LRP1B\|MBOAT1\|MCM9\|MR1\|MROH8\|MUSK\|MYH15\|MYO16\|MYO1H\|NAALADL2\|NAV3\|NEB\|NEK5\|NF1\|NPAS3\|NPHP1\|NSF\|NUP210L\|PARP11\|PCDHB1\|PCSK5\|PDE1A\|PDE4DIP\|PDZD2\|PGR\|PHLPP1\|PIGK\|PIP5K1B\|PIWIL2\|PKHD1\|PLA2G4A\|PLA2R1\|PRKAA2\|PTGFR\|PTPRZ1\|PZP\|RBM44\|RGPD4\|RGPD5\|RGPD6\|RGPD8\|RORB\|RP11-327J17.3\|RP11-478J18.1\|RP11-550E22.3\|RYR3\|SCO1\|SEMA3E\|SERPINB8\|SH3TC2\|SLC12A1\|SLC2A13\|SNAP25-AS1\|SOCS7\|SPEF2\|SPHKAP\|SSX2IP\|STK31\|SYNE1\|SYTL2\|TAOK1\|TBC1D8B\|TEX14\|TF\|THEMIS\|THSD7B\|TMC1\|TMEM232\|TP63\|TRPM3\|USP32\|VGLL3\|VPS13D\|WDR66\|WDR7\|WNT2B\|XKR4 | 8.25E-14 | ** |

**Human NB to Survival ∩ Age at diagnosis in 498 NBs (Fig 6)**

| NB cell cluster with signature genes significantly up-regulated in different survival groups in 498 SEQC NB samples | Correlation with age at diagnosis in 498 SEQC NB samples | Number of genes in NB specific signature significantly up-regulated in survival group and correlated with age at diagnosis in 498 SEQC NB samples | Genes in NB specific signature significantly up-regulated in survival group and correlated with age at diagnosis in 498 SEQC NB samples | p-value/FDR | Significance |
|---|---|---|---|---|---|

<table>
<tr><td>NOR (nC9) ∩ Better survival</td><td>Inversely correlated</td><td>73</td><td>AFAP1|ALCAM|ARL6IP5|ASTN1|ATF2|ATP11A|CCDC82|CD47|CHST1|CNTN1|CPNE8|CREG2|CSRNP3|CYGB|DDAH1|DDX6|DLEU2|DPYSL3|DSEL|DUSP8|EHBP1|EIF1B|FMN1|FOXP1|FSTL5|GAB2|GNAO1|INO80D|JARID2|JPH3|JUN|KCNQ3|KIDINS220|LIFR|LONRF2|LRRN3|MARCH4|MBNL2|MST1|MTMR6|MXRA7|NAALAD2|NRP1|OPTN|PCDH10|PCDH7|PCDHB16|PGM2L1|PJA2|PKIB|PMP22|RBMS1|RBMS3|RGS5|RNF144A|SCG2|SCG5|SCN7A|SCRN1|SEMA6D|SEZ6|SFRP1|SLC25A12|ST6GALNAC5|STMN4|SYT13|TAGLN3|TMCO3|TMEM255A|TMOD2|TNRC6B|TSHZ2|ZBTB38</td><td>9.51E-073/<1E-50</td><td>**</td></tr>
<tr><td>Undifferentiated (nC3) ∩ Poor survival</td><td>Directly correlated</td><td>115</td><td>ABCA12|ABCC4|ACTR3C|ADAM29|ADAM7|ADAMTS19|ADAMTS20|AFF2|ANKRD45|AP5M1|ARMC10|ARMC3|BRCA1|BRMS1L|C14orf145|CCDC169|CCSER1|CECR2|CEP112|CEP152|CISD2|CMC2|CPSF2|CRB1|CSMD3|DCDC1|DLX6-AS1|DNAH11|DNAH14|DNAH3|DPY19L3|ELMOD1|ELP4|EML1|ESCO2|EXOC5|FAM197Y5|FAM227A|FANCI|FANCM|FBXO15|G2E3|GABRA6|GALK2|GEMIN5|GRIN2B|HEATR4|INTS2|INTU|IPO11|KIAA1257|KIF6|KNTC1|L2HGDH|LGR5|LMO3|LRRC9|MCCC2|MET|MTHFD2L|MYH13|MYH4|NT5DC3|NUBPL|NUP155|NUP37|NUP93|OR4N2|PGGT1B|PIK3R4|POTEC|POU6F2|PPAT|PRDM5|PRDM7|PRIM2|PTCD2|RAD51B|RBBP8|RFX6|RGS22|RMST|SATB2|SEC23A|SHQ1|SLC35G1|SLC5A12|SLC5A8|SLC9B1|SLC9C2|SLCO6A1|SNTG1|SPAG17|SPATA17|SPECC1|SPINK5|TRDN|TTC26|TTC5|TTLL5|TYR|TYRP1|ULK4|UTP20|WDR3|WDR35|WDR72|ZBBX|ZDHHC13|ZFAND4|ZNF169|ZNF215|ZNF479|ZNF595|ZNF716</td><td>1.9E-067/<1E-50</td><td>**</td></tr>
<tr><td>Undifferentiated (nC3) ∩ Better survival</td><td>Inversely correlated</td><td>83</td><td>A2ML1|ABCA10|ABCA6|ABCA9|ABCB4|AGBL3|AK5|AKAP7|ALDH1L2|AMY1A|AMY1B|AMY1C|AMY2A|ANO4|ARHGAP28|ARHGAP42|ARHGEF12|ATP13A4|BCO2|CC2D2B|CDH9|CDON|COL21A1|CUBN|DAB1|DSC2|EFCAB5|EHF|FAM135A|FAT3|FLRT2|FREM1|GDPD4|GK5|GLI3|GRID2|HMGA2|IGFN1|IL1RAP|IMPG1|INPP4B|ITGA8|KIAA1328|KIF13A|LAMA3|LAMB4|LRP1B|MCM9|MROH8|NAALADL2|NAV3|NEB|NF1|NPAS3|NUP210L|PARP11|PCSK5|PDE1A|PDE4DIP|PDZD2|PHLPP1|PIP5K1B|PLA2G4A|PRKAA2|PTPRZ1|RGPD5|RGPD6|RGPD8|RORB|RP11-327J17.3|SEMA3E|SH3TC2|SLC12A1|SOCS7|STK31|SYTL2|TAOK1|TBC1D8B|TF|VGLL3|VPS13D|WDR7|WNT2B</td><td>1.67E-055/<1E-50</td><td>**</td></tr>
<tr><td>NOR (nC9) ∩ Poor survival</td><td>Directly correlated</td><td>44</td><td>ALKBH7|BMP7|COPE|COX6B1|COX7C|COX8A|DPY30|EEF1B2|FAU|GCSH|H2AFZ|HDGF|HINT1|HIST1H4E|HMGB2|HNRNPL|HSD17B11|HSD17B12|HTATSF1|JTB|MIR3654|NDUFA1|NDUFB11|OST4|RMND5A|RPL18A|RPL31|RPL34|RPL35A|RPS10|RPS11|RPS13|RPS15|RPS15A|RPS19|RPS21|RPS3|RPS7|SF3B2|SMARCA5|SNRNP27|SNRPD2|TOMM22|UBA52</td><td>4.76E-023/9.52E-023</td><td>**</td></tr>
</table>
