## Supplementary Table 7 for "Single-nuclei transcriptomes from human adrenal gland reveals distinct cellular identities of low and high-risk neuroblastoma tumors"

**Biological processes (GO) enriched in genes human NB significantly associated with survival in 498 NBs (Supplementary Fig 6)**

| NB cell cluster with signature genes significantly up-regulated in different survival groups in 498 SEQC NB samples | Biological process (GO) | Number of genes in NB specific signature significantly up-regulated in survival group and enriched in biological process (GO) | Genes in NB specific signature significantly up-regulated in survival group and enriched in biological process (GO) | FDR | Significance |
|---|---|---|---|---|---|
| Undifferentiated (nC3) ∩ Better survival | cell adhesion | 19 | CD226\|CDH10\|CDH18\|CDH9\|CDON\|CNTN3\|CNTNAP3\|COL28A1\|DSC2\|FAT3\|FLRT2\|FREM1\|FREM2\|ITGA8\|LAMA3\|LAMB4\|PCDHB1\|PDZD2\|SSX2IP | 1.84E-10 | ** |
| Undifferentiated (nC3) ∩ Better survival | multicellular organism development | 18 | DAB1\|DDX4\|DOCK7\|EHF\|FAT3\|FLRT2\|FREM1\|FREM2\|HMGA2\|HYDIN\|ITGA8\|MUSK\|PIWIL2\|RORB\|SEMA3E\|THEMIS\|TP63\|WNT2B | 2.01E-05 | ** |
| Undifferentiated (nC3) ∩ Better survival | homophilic cell adhesion via plasma membrane adhesion molecules | 7 | CD226\|CDH10\|CDH18\|CDH9\|DSC2\|FAT3\|PCDHB1 | 7.83E-04 | ** |
| Undifferentiated (nC3) ∩ Better survival | intracellular transport | 4 | RGPD4\|RGPD5\|RGPD6\|RGPD8 | 3.06E-03 | ** |
| Undifferentiated (nC3) ∩ Better survival | lipid transport | 6 | ABCA10\|ABCA6\|ABCA9\|ABCB4\|ANO4\|ATP10B | 3.06E-03 | ** |
| Undifferentiated (nC3) ∩ Better survival | smooth muscle tissue development | 3 | ITGA8\|NF1\|TP63 | 5.83E-03 | ** |
| Undifferentiated (nC3) ∩ Better survival | signal transduction | 19 | ADCY2\|ARHGAP28\|ARHGAP42\|CD226\|CUBN\|GRID2\|GRIN2A\|HMGA2\|IL1RAP\|INPP4B\|MR1\|NF1\|NPHP1\|PDE1A\|PGR\|PKHD1\|PLA2R1\|PRKAA2\|PTGFR | 7.17E-03 | ** |
| Undifferentiated (nC3) ∩ Better survival | brain development | 7 | CLN5\|DAB1\|GLI3\|GRIN2A\|HYDIN\|ITGA8\|NF1 | 7.89E-03 | ** |
| Undifferentiated (nC3) ∩ Better survival | negative regulation of JAK-STAT cascade | 4 | DAB1\|FLRT2\|HMGA2\|SOCS7 | 9.38E-03 | ** |
| Undifferentiated (nC3) ∩ Better survival | ion transmembrane transport | 7 | ANO4\|ATP13A4\|GRID2\|GRIN2A\|RYR3\|SLC12A1\|TRPM3 | 1.38E-02 | * |
| Undifferentiated (nC3) ∩ Better survival | cell differentiation | 12 | DAB1\|DDX4\|DOCK7\|FLT3\|ITGA8\|MUSK\|NPHP1\|PARP11\|PIWIL2\|SEMA3E\|SYNE1\|TP63 | 1.41E-02 | * |
| Undifferentiated (nC3) ∩ Better survival | cell-cell adhesion | 5 | CDH10\|CDH9\|ITGA8\|NPHP1\|PKHD1 | 2.37E-02 | * |
| Undifferentiated (nC3) ∩ Better survival | phosphorylation | 10 | AK5\|AKAP7\|FLT3\|GK5\|MUSK\|NEK5\|PIP5K1B\|PRKAA2\|STK31\|TAOK1 | 3.03E-02 | * |
| Undifferentiated (nC3) ∩ Better survival | calcium ion transmembrane transport | 5 | ATP13A4\|GRIN2A\|RYR3\|TMC1\|TRPM3 | 3.32E-02 | * |
| Undifferentiated (nC3) ∩ Better survival | carbohydrate metabolic process | 6 | AMY1A\|AMY1B\|AMY1C\|AMY2A\|EPM2A\|GK5 | 4.47E-02 | * |
| NOR (nC9) ∩ Better survival | calcium-dependent cell-cell adhesion via plasma membrane cell adhesion molecules | 4 | ATP2C1\|PCDHB10\|PCDHB16\|PCDHB6 | 2.72E-04 | ** |
| NOR (nC9) ∩ Better survival | cell adhesion | 9 | ALCAM\|ASTN1\|CD47\|CNTN1\|PCDH10\|PCDH7\|PCDHB10\|PCDHB16\|PCDHB6 | 9.18E-04 | ** |
| NOR (nC9) ∩ Better survival | chemical synaptic transmission | 6 | KCNQ3\|PCDHB10\|PCDHB16\|PCDHB6\|PMP22\|SYT1 | 2.23E-03 | ** |
| NOR (nC9) ∩ Better survival | motor neuron axon guidance | 3 | ALCAM\|FOXP1\|NRP1 | 4.27E-03 | ** |
| NOR (nC9) ∩ Better survival | homophilic cell adhesion via plasma membrane adhesion molecules | 5 | PCDH10\|PCDH7\|PCDHB10\|PCDHB16\|PCDHB6 | 5.90E-03 | ** |
| NOR (nC9) ∩ Better survival | positive regulation of focal adhesion assembly | 3 | FMN1\|NRP1\|SFRP1 | 7.71E-03 | ** |

| | | | | | |
|---|---|---|---|---|---|
| NOR (nC9) ∩ Better survival | calcium ion-regulated exocytosis of neurotransmitter | 3 | SYT1\|SYT13\|SYT4 | 1.14E-02 | * |
| NOR (nC9) ∩ Better survival | nervous system development | 7 | CNTN1\|KIDINS220\|NRP1\|PCDH10\|PCDHB6\|SEMA6D\|TMOD2 | 2.04E-02 | * |
| NOR (nC9) ∩ Better survival | regulation of protein kinase C signaling | 2 | SEZ6\|SEZ6L | 3.65E-02 | * |
| NOR (nC9) ∩ Better survival | axon extension involved in axon guidance | 2 | ALCAM\|NRP1 | 4.59E-02 | * |
| Undifferentiated (nC3) ∩ Poor survival | cellular response to DNA damage stimulus | 21 | BARD1\|BRCA1\|BRCA2\|BRIP1\|C12orf48\|CHEK2\|CLSPN\|FANCD2\|FANCI\|FANCM\|GEN1\|INTS7\|MMS22L\|POLQ\|RAD51B\|RBBP8\|SMC6\|SPDYA\|TTC5\|WRN\|ZRANB3 | 3.47E-11 | ** |
| Undifferentiated (nC3) ∩ Poor survival | DNA repair | 19 | BARD1\|BRCA1\|BRCA2\|BRIP1\|C12orf48\|CHEK2\|CLSPN\|FANCD2\|FANCI\|FANCM\|GEN1\|MMS22L\|POLQ\|RAD51B\|RBBP8\|SMC6\|TTC5\|WRN\|ZRANB3 | 5.87E-11 | ** |
| Undifferentiated (nC3) ∩ Poor survival | cell cycle | 20 | ANAPC1\|ASPM\|BRCA1\|BRCA2\|BUB1\|CENPE\|CHEK2\|CLSPN\|CTCFL\|E2F6\|ESCO2\|FANCD2\|FANCI\|GNAI3\|KNTC1\|NCAPG2\|NEDD1\|NUP37\|RBBP8\|SPDYA | 9.03E-08 | ** |
| Undifferentiated (nC3) ∩ Poor survival | double-strand break repair via homologous recombination | 9 | BRCA1\|BRCA2\|GEN1\|MMS22L\|POLQ\|RAD51B\|RBBP8\|SMC6\|WRN | 3.55E-07 | ** |
| Undifferentiated (nC3) ∩ Poor survival | microtubule-based movement | 10 | CENPE\|DNAH11\|DNAH14\|DNAH3\|KIF14\|KIF15\|KIF6\|MYH1\|MYH13\|MYH4 | 4.15E-07 | ** |
| Undifferentiated (nC3) ∩ Poor survival | detection of chemical stimulus involved in sensory perception of smell | 15 | OR10K1\|OR1A1\|OR2AG1\|OR2AT4\|OR2M3\|OR2T12\|OR4D9\|OR4F4\|OR4F5\|OR4N2\|OR52A1\|OR56A1\|OR56A4\|OR5AS1\|OR8D1 | 1.06E-06 | ** |
| Undifferentiated (nC3) ∩ Poor survival | sensory perception of smell | 15 | OR10K1\|OR1A1\|OR2AG1\|OR2AT4\|OR2M3\|OR2T12\|OR4D9\|OR4F4\|OR4F5\|OR4N2\|OR52A1\|OR56A1\|OR56A4\|OR5AS1\|OR8D1 | 2.57E-06 | ** |
| Undifferentiated (nC3) ∩ Poor survival | regulation of transcription, DNA-templated | 35 | ANKRD30A\|ATAD2\|BRCA1\|BRMS1L\|CHEK2\|CTCFL\|DEPDC1\|DR1\|E2F6\|ELP4\|ETV1\|LMO3\|MAP2K6\|MED12L\|MED17\|MYB\|NUP133\|POU6F2\|PRDM5\|PRDM7\|RFX6\|SATB2\|ST18\|TAF4B\|ZNF169\|ZNF215\|ZNF285\|ZNF426\|ZNF479\|ZNF519\|ZNF578\|ZNF595\|ZNF678\|ZNF705G\|ZNF716 | 1.20E-05 | ** |
| Undifferentiated (nC3) ∩ Poor survival | response to stimulus | 16 | OR10K1\|OR1A1\|OR2AG1\|OR2AT4\|OR2M3\|OR2T12\|OR4D9\|OR4F4\|OR4F5\|OR4N2\|OR52A1\|OR56A1\|OR56A4\|OR5AS1\|OR8D1\|RP1 | 1.61E-05 | ** |
| Undifferentiated (nC3) ∩ Poor survival | transcription, DNA-templated | 33 | ATAD2\|BRCA1\|BRMS1L\|CHEK2\|CTCFL\|DEPDC1\|DR1\|E2F6\|ELP4\|ETV1\|LMO3\|MAP2K6\|MED12L\|MED17\|MYB\|POU6F2\|PRDM5\|RFX6\|SATB2\|ST18\|TAF4B\|TTLL5\|ZNF169\|ZNF215\|ZNF285\|ZNF426\|ZNF479\|ZNF519\|ZNF578\|ZNF595\|ZNF678\|ZNF705G\|ZNF716 | 2.05E-05 | ** |
| Undifferentiated (nC3) ∩ Poor survival | double-strand break repair | 7 | BRCA1\|BRCA2\|BRIP1\|CHEK2\|ESCO2\|POLQ\|WRN | 3.77E-05 | ** |
| Undifferentiated (nC3) ∩ Poor survival | replication fork processing | 5 | FANCM\|GEN1\|MMS22L\|WRN\|ZRANB3 | 3.01E-04 | ** |
| Undifferentiated (nC3) ∩ Poor survival | regulation of transcription by RNA polymerase II | 21 | ATAD2\|BRCA1\|BRIP1\|ELP4\|MED12L\|MED17\|POU6F2\|PRDM7\|RBBP8\|SATB2\|WDHD1\|ZNF169\|ZNF215\|ZNF285\|ZNF426\|ZNF479\|ZNF519\|ZNF595\|ZNF678\|ZNF705G\|ZNF716 | 3.59E-04 | ** |
| Undifferentiated (nC3) ∩ Poor survival | cell division | 12 | ANAPC1\|ASPM\|BUB1\|CENPE\|CHEK2\|GNAI3\|KIF14\|KNTC1\|NCAPG2\|NEDD1\|NUP37\|RBBP8 | 5.02E-04 | ** |
| Undifferentiated (nC3) ∩ Poor survival | metabolic process | 13 | ACSM2B\|ACSM3\|CPS1\|GALK2\|GPAM\|IDE\|MTHFD2L\|PPAT\|SI\|TYR\|TYRP1\|WRN\|ZRANB3 | 5.35E-04 | ** |
| Undifferentiated (nC3) ∩ Poor survival | DNA replication | 8 | BARD1\|BRCA1\|BRIP1\|CLSPN\|POLQ\|PRIM2\|RBBP8\|WRN | 8.80E-04 | ** |
| Undifferentiated (nC3) ∩ Poor survival | G-protein coupled receptor signaling pathway | 19 | GNAI3\|KNG1\|LGR4\|LGR5\|OR10K1\|OR1A1\|OR2AG1\|OR2AT4\|OR2M3\|OR2T12\|OR4D9\|OR4F4\|OR4F5\|OR4N2\|OR52A1\|OR56A1\|OR56A4\|OR5AS1\|OR8D1 | 1.38E-03 | ** |
| Undifferentiated (nC3) ∩ Poor survival | detection of chemical stimulus involved in sensory perception | 6 | OR10K1\|OR2AT4\|OR4D9\|OR4F4\|OR4F5\|OR4N2 | 3.06E-03 | ** |
| Undifferentiated (nC3) ∩ Poor survival | signal transduction | 25 | ANTXR2\|DEPDC1\|EGF\|FAM13A\|GABRA6\|GNAI3\|LGR4\|LGR5\|MAP2K6\|MET\|OR10K1\|OR1A1\|OR2AG1\|OR2AT4\|OR2M3\|OR2T12\|OR4D9\|OR4F4\|OR4F5\|OR4N2\|OR52A1\|OR56A1\|OR56A4\|OR5AS1\|OR8D1 | 4.59E-03 | ** |
| Undifferentiated (nC3) ∩ Poor survival | DNA damage checkpoint | 4 | BRIP1\|CHEK2\|CLSPN\|INTS7 | 1.41E-02 | * |

| | | | | | |
|---|---|---|---|---|---|
| Undifferentiated (nC3) ∩ Poor survival | DNA double-strand break processing | 3 | BARD1\|BRCA1\|RBBP8 | 2.94E-02 | * |
| Undifferentiated (nC3) ∩ Poor survival | chromosome segregation | 5 | BRCA1\|BUB1\|CENPE\|ESCO2\|NUP37 | 3.06E-02 | * |
| Undifferentiated (nC3) ∩ Poor survival | DNA damage response, signal transduction by p53 class mediator resulting in transcription of p21 class mediator | 3 | BRCA1\|BRCA2\|CHEK2 | 3.39E-02 | * |
| Undifferentiated (nC3) ∩ Poor survival | neural tube development | 4 | CECR2\|INTU\|NUP133\|STIL | 4.06E-02 | * |
| Undifferentiated (nC3) ∩ Poor survival | positive regulation of canonical Wnt signaling pathway | 5 | ASPM\|EGF\|LGR4\|LGR5\|SCEL | 4.64E-02 | * |
| NOR (nC9) ∩ Poor survival | SRP-dependent cotranslational protein targeting to membrane | 30 | FAU\|RPL14\|RPL15\|RPL18\|RPL18A\|RPL23A\|RPL24\|RPL26\|RPL27\|RPL29\|RPL31\|RPL32\|RPL34\|RPL35A\|RPL4\|RPS10\|RPS11\|RPS13\|RPS15\|RPS15A\|RPS19\|RPS21\|RPS24\|RPS27A\|RPS3\|RPS5\|RPS7\|RPS9\|RPSA\|UBA52 | <1E-50 | ** |
| NOR (nC9) ∩ Poor survival | translation | 38 | EEF1B2\|EEF2\|EIF2AK2\|EIF3L\|EPRS\|FAU\|RPL14\|RPL15\|RPL18\|RPL18A\|RPL23A\|RPL24\|RPL26\|RPL27\|RPL29\|RPL31\|RPL32\|RPL34\|RPL35A\|RPL39L\|RPL4\|RPS10\|RPS11\|RPS13\|RPS15\|RPS15A\|RPS19\|RPS21\|RPS24\|RPS27A\|RPS3\|RPS5\|RPS7\|RPS9\|RPSA\|RSL24D1\|TARS\|UBA52 | <1E-50 | ** |
| NOR (nC9) ∩ Poor survival | nuclear-transcribed mRNA catabolic process, nonsense-mediated decay | 30 | FAU\|RPL14\|RPL15\|RPL18\|RPL18A\|RPL23A\|RPL24\|RPL26\|RPL27\|RPL29\|RPL31\|RPL32\|RPL34\|RPL35A\|RPL4\|RPS10\|RPS11\|RPS13\|RPS15\|RPS15A\|RPS19\|RPS21\|RPS24\|RPS27A\|RPS3\|RPS5\|RPS7\|RPS9\|RPSA\|UBA52 | <1E-50 | ** |
| NOR (nC9) ∩ Poor survival | translational initiation | 31 | EIF3L\|FAU\|RPL14\|RPL15\|RPL18\|RPL18A\|RPL23A\|RPL24\|RPL26\|RPL27\|RPL29\|RPL31\|RPL32\|RPL34\|RPL35A\|RPL4\|RPS10\|RPS11\|RPS13\|RPS15\|RPS15A\|RPS19\|RPS21\|RPS24\|RPS27A\|RPS3\|RPS5\|RPS7\|RPS9\|RPSA\|UBA52 | <1E-50 | ** |
| NOR (nC9) ∩ Poor survival | cytoplasmic translation | 13 | RPL15\|RPL18\|RPL18A\|RPL24\|RPL26\|RPL29\|RPL31\|RPL32\|RPL35A\|RPL4\|RPS21\|RPS3\|RPS7 | 6.90E-22 | ** |
| NOR (nC9) ∩ Poor survival | rRNA processing | 9 | RPL14\|RPL26\|RPL27\|RPL35A\|RPS15\|RPS19\|RPS24\|RPS7\|RPS9 | 1.54E-07 | ** |
| NOR (nC9) ∩ Poor survival | ribosomal small subunit assembly | 5 | RPS10\|RPS15\|RPS19\|RPS5\|RPSA | 6.73E-07 | ** |
| NOR (nC9) ∩ Poor survival | mRNA splicing, via spliceosome | 8 | HNRNPL\|HSPA8\|HTATSF1\|SF3B2\|SNRNP27\|SNRPD2\|U2SURP\|YBX1 | 8.59E-05 | ** |
| NOR (nC9) ∩ Poor survival | ribosomal small subunit biogenesis | 4 | RPS15\|RPS19\|RPS24\|RPS7 | 1.60E-04 | ** |
| NOR (nC9) ∩ Poor survival | positive regulation of transcription by RNA polymerase II | 12 | ATRX\|BMP7\|H2AFZ\|HMGA1\|HMGB1\|HMGB2\|RPS27A\|SMARCC1\|TCF12\|TEAD1\|UBA52\|YBX1 | 2.27E-03 | ** |
| NOR (nC9) ∩ Poor survival | oxidation-reduction process | 10 | ALKBH7\|HADHA\|HSD17B11\|HSD17B12\|LDHB\|MTHFD2\|NDUFA1\|NDUFB11\|PRDX2\|UQCRC1 | 2.81E-03 | ** |
| NOR (nC9) ∩ Poor survival | nucleosome assembly | 5 | ATRX\|H1F0\|HIST1H4E\|HMGB2\|SMARCA5 | 3.75E-03 | ** |
| NOR (nC9) ∩ Poor survival | apoptotic DNA fragmentation | 3 | H1F0\|HMGB1\|HMGB2 | 4.27E-03 | ** |
| NOR (nC9) ∩ Poor survival | endonucleolytic cleavage to generate mature 3'-end of SSU-rRNA from (SSU-rRNA, 5.8S rRNA, LSU-rRNA) | 2 | RPS21\|RPSA | 4.59E-03 | ** |
| NOR (nC9) ∩ Poor survival | assembly of large subunit precursor of preribosome | 2 | RPL24\|RSL24D1 | 1.14E-02 | * |
| NOR (nC9) ∩ Poor survival | negative regulation of translation | 4 | EIF2AK2\|EPRS\|FXR1\|RPS3 | 1.14E-02 | * |
| NOR (nC9) ∩ Poor survival | ribosomal large subunit assembly | 3 | RPL23A\|RPL24\|RSL24D1 | 1.54E-02 | * |

| | | | | | |
|---|---|---|---|---|---|
| NOR (nC9) ∩ Poor survival | transcription, DNA-templated | 16 | ATRX\|BTF3\|DPY30\|EIF2AK2\|HDGF\|HINT1\|HMGA1\|HMGB2\|HSPA8\|HTATSF1\|POLR1D\|RPS3\|SMARCC1\|TCF12\|TEAD1\|YBX1 | 1.94E-02 | * |
| NOR (nC9) ∩ Poor survival | DNA geometric change | 2 | HMGB1\|HMGB2 | 2.02E-02 | * |
| NOR (nC9) ∩ Poor survival | ribosomal large subunit biogenesis | 3 | RPL14\|RPL26\|RPL35A | 2.04E-02 | * |
| NOR (nC9) ∩ Poor survival | positive regulation of transcription, DNA-templated | 8 | BMP7\|CKS1B\|HMGA1\|HMGB2\|SMARCA5\|SMARCC1\|TCF12\|TEAD1 | 2.75E-02 | * |
| NOR (nC9) ∩ Poor survival | DNA damage response, detection of DNA damage | 3 | RPS27A\|RPS3\|UBA52 | 3.80E-02 | * |

**Biological processes (GO) enriched in genes human NB significantly associated with age at diagnosis in 498 NBs (Supplementary Fig 6)**

| NB cell cluster with signature genes significantly correlated with age at diagnosis in 498 SEQC NB samples | Biological process (GO) | Number of genes in NB specific signature significantly correlated with age at diagnosis and enriched in biological process (GO) | Genes in NB specific signature significantly correlated with age at diagnosis and enriched in biological process (GO) | FDR | Significance |
|---|---|---|---|---|---|
| NOR (nC9) ∩ Directly correlated with age at diagnosis | SRP-dependent cotranslational protein targeting to membrane | 15 | FAU\|RPL18A\|RPL31\|RPL34\|RPL35A\|RPS10\|RPS11\|RPS13\|RPS15\|RPS15A\|RPS19\|RPS21\|RPS3\|RPS7\|UBA52 | 3.92E-27 | ** |
| NOR (nC9) ∩ Directly correlated with age at diagnosis | nuclear-transcribed mRNA catabolic process, nonsense-mediated decay | 15 | FAU\|RPL18A\|RPL31\|RPL34\|RPL35A\|RPS10\|RPS11\|RPS13\|RPS15\|RPS15A\|RPS19\|RPS21\|RPS3\|RPS7\|UBA52 | 1.60E-25 | ** |
| NOR (nC9) ∩ Directly correlated with age at diagnosis | translational initiation | 15 | FAU\|RPL18A\|RPL31\|RPL34\|RPL35A\|RPS10\|RPS11\|RPS13\|RPS15\|RPS15A\|RPS19\|RPS21\|RPS3\|RPS7\|UBA52 | 2.55E-24 | ** |
| NOR (nC9) ∩ Directly correlated with age at diagnosis | translation | 16 | EEF1B2\|FAU\|RPL18A\|RPL31\|RPL34\|RPL35A\|RPS10\|RPS11\|RPS13\|RPS15\|RPS15A\|RPS19\|RPS21\|RPS3\|RPS7\|UBA52 | 4.56E-21 | ** |
| NOR (nC9) ∩ Directly correlated with age at diagnosis | cytoplasmic translation | 6 | RPL18A\|RPL31\|RPL35A\|RPS21\|RPS3\|RPS7 | 6.20E-09 | ** |
| NOR (nC9) ∩ Directly correlated with age at diagnosis | ribosomal small subunit assembly | 3 | RPS10\|RPS15\|RPS19 | 1.17E-03 | ** |
| NOR (nC9) ∩ Directly correlated with age at diagnosis | ribosomal small subunit biogenesis | 3 | RPS15\|RPS19\|RPS7 | 1.69E-03 | ** |
| NOR (nC9) ∩ Directly correlated with age at diagnosis | mRNA splicing, via spliceosome | 5 | HNRNPL\|HTATSF1\|SF3B2\|SNRNP27\|SNRPD2 | 4.41E-03 | ** |
| NOR (nC9) ∩ Directly correlated with age at diagnosis | rRNA processing | 4 | RPL35A\|RPS15\|RPS19\|RPS7 | 1.18E-02 | * |
| NOR (nC9) ∩ Directly correlated with age at diagnosis | estrogen biosynthetic process | 2 | HSD17B11\|HSD17B12 | 4.99E-02 | * |
| NOR (nC9) ∩ Inversely correlated with age at diagnosis | forebrain development | 5 | ARHGAP35\|FOXP1\|GNAO1\|SETD2\|TOP2B | 1.17E-03 | ** |
| NOR (nC9) ∩ Inversely correlated with age at diagnosis | cell adhesion | 9 | ALCAM\|ASTN1\|CD47\|CNTN1\|PCDH10\|PCDH7\|PCDHB16\|PODXL2\|RND3 | 7.06E-03 | ** |
| NOR (nC9) ∩ Inversely correlated with age at diagnosis | response to cytokine | 4 | BCL2\|GNAO1\|JUN\|LIFR | 1.04E-02 | * |

| | | | | | |
|---|---|---|---|---|---|
| NOR (nC9) ∩ Inversely correlated with age at diagnosis | motor neuron axon guidance | 3 | ALCAM\|FOXP1\|NRP1 | 1.12E-02 | * |
| NOR (nC9) ∩ Inversely correlated with age at diagnosis | positive regulation of smooth muscle cell migration | 3 | BCL2\|NRP1\|SEMA6D | 1.12E-02 | * |
| NOR (nC9) ∩ Inversely correlated with age at diagnosis | positive regulation of cell proliferation | 8 | BCL2\|CD47\|GAB2\|IRS2\|JUN\|LIFR\|SFRP1\|TIAL1 | 1.31E-02 | * |
| NOR (nC9) ∩ Inversely correlated with age at diagnosis | positive regulation of cardiac muscle myoblast proliferation | 2 | ATF2\|MEIS2 | 1.52E-02 | * |
| NOR (nC9) ∩ Inversely correlated with age at diagnosis | ureteric bud invasion | 2 | FMN1\|KIF26B | 1.52E-02 | * |
| NOR (nC9) ∩ Inversely correlated with age at diagnosis | positive regulation of focal adhesion assembly | 3 | FMN1\|NRP1\|SFRP1 | 1.81E-02 | * |
| NOR (nC9) ∩ Inversely correlated with age at diagnosis | negative regulation of cell proliferation | 7 | BCL2\|DDAH1\|JARID2\|JUN\|PBRM1\|PMP22\|SFRP1 | 2.68E-02 | * |
| NOR (nC9) ∩ Inversely correlated with age at diagnosis | response to drug | 6 | BCL2\|GNAO1\|JUN\|KCNK3\|OXCT1\|SFRP1 | 3.55E-02 | * |
| NOR (nC9) ∩ Inversely correlated with age at diagnosis | axon guidance | 5 | ALCAM\|ARHGAP35\|GAB2\|IRS2\|NRP1 | 3.85E-02 | * |
| NOR (nC9) ∩ Inversely correlated with age at diagnosis | mammary gland development | 3 | ARHGAP35\|IRS2\|MST1 | 4.02E-02 | * |
| Undifferentiated (nC3) ∩ Directly correlated with age at diagnosis | microtubule-based movement | 12 | DNAH10\|DNAH11\|DNAH14\|DNAH3\|DNAH5\|DNAH6\|DNAH7\|DNAH8\|KIF6\|MYH13\|MYH4\|MYO3B | 1.83E-11 | ** |
| Undifferentiated (nC3) ∩ Directly correlated with age at diagnosis | cilium movement | 5 | DNAH11\|DNAH3\|DNAH5\|DNAH7\|DNAH8 | 6.37E-04 | ** |
| Undifferentiated (nC3) ∩ Directly correlated with age at diagnosis | cilium assembly | 8 | ABCC4\|CDC14B\|DNAH14\|DNAH5\|DNAH6\|INTU\|TTC26\|WDR35 | 6.55E-04 | ** |
| Undifferentiated (nC3) ∩ Directly correlated with age at diagnosis | mRNA processing | 10 | AFF2\|CPSF2\|GEMIN5\|PTCD2\|RBMY1A1\|RBMY1B\|RBMY1D\|RBMY1E\|RBMY1F\|RBMY1J | 1.17E-03 | ** |
| Undifferentiated (nC3) ∩ Directly correlated with age at diagnosis | cilium-dependent cell motility | 3 | DNAH3\|DNAH7\|DNAH8 | 7.80E-03 | ** |
| Undifferentiated (nC3) ∩ Directly correlated with age at diagnosis | RNA splicing | 8 | AFF2\|GEMIN5\|RBMY1A1\|RBMY1B\|RBMY1D\|RBMY1E\|RBMY1F\|RBMY1J | 9.29E-03 | ** |
| Undifferentiated (nC3) ∩ Directly correlated with age at diagnosis | ion transport | 11 | GABRA6\|GABRG3\|GRIN2B\|PKD1L3\|SLC13A1\|SLC5A12\|SLC5A8\|SLC9B1\|SLC9C2\|SLCO1A2\|SLCO6A1 | 1.38E-02 | * |
| Undifferentiated (nC3) ∩ Directly correlated with age at diagnosis | cilium or flagellum-dependent cell motility | 2 | DNAH14\|DNAH6 | 1.42E-02 | * |
| Undifferentiated (nC3) ∩ Directly correlated with age at diagnosis | flagellated sperm motility | 4 | DNAH11\|DNAH5\|SLC9B1\|TTLL5 | 3.85E-02 | * |
| Undifferentiated (nC3) ∩ Inversely correlated with age at diagnosis | cell adhesion | 14 | CDH9\|CDON\|CNTN5\|COL6A6\|DSC2\|FAT3\|FLRT2\|FREM1\|ITGA8\|LAMA1\|LAMA2\|LAMA3\|LAMB4\|PDZD2 | 1.14E-07 | ** |

| | | | | | |
|---|---|---|---|---|---|
| Undifferentiated (nC3) ∩ Inversely correlated with age at diagnosis | multicellular organism development | 13 | APAF1\|DAB1\|EHF\|FAT3\|FLRT2\|FREM1\|HMGA2\|ITGA8\|PRTG\|RORB\|SEMA3E\|SLCO4C1\|WNT2B | 1.17E-03 | ** |
| Undifferentiated (nC3) ∩ Inversely correlated with age at diagnosis | intracellular transport | 4 | RGPD3\|RGPD5\|RGPD6\|RGPD8 | 1.53E-03 | ** |
| Undifferentiated (nC3) ∩ Inversely correlated with age at diagnosis | regulation of cell adhesion | 4 | LAMA1\|LAMA2\|LAMA3\|LMO7 | 4.66E-03 | ** |
| Undifferentiated (nC3) ∩ Inversely correlated with age at diagnosis | negative regulation of JAK-STAT cascade | 4 | DAB1\|FLRT2\|HMGA2\|SOCS7 | 4.83E-03 | ** |
| Undifferentiated (nC3) ∩ Inversely correlated with age at diagnosis | extracellular matrix organization | 6 | FBN2\|ITGA8\|LAMA1\|LAMA2\|LAMA3\|NF1 | 6.36E-03 | ** |
| Undifferentiated (nC3) ∩ Inversely correlated with age at diagnosis | metabolic process | 8 | ACSS3\|ALDH1L2\|AMY1A\|AMY1B\|AMY1C\|AMY2A\|ENPP1\|PLA2G4A | 9.31E-03 | ** |
| Undifferentiated (nC3) ∩ Inversely correlated with age at diagnosis | lipid transport | 5 | ABCA10\|ABCA6\|ABCA9\|ABCB4\|ANO4 | 1.04E-02 | * |
| Undifferentiated (nC3) ∩ Inversely correlated with age at diagnosis | regulation of embryonic development | 3 | LAMA1\|LAMA2\|LAMA3 | 1.12E-02 | * |
| Undifferentiated (nC3) ∩ Inversely correlated with age at diagnosis | limb morphogenesis | 3 | FBN2\|GLI3\|PCSK5 | 1.81E-02 | * |
| Undifferentiated (nC3) ∩ Inversely correlated with age at diagnosis | intracellular signal transduction | 7 | ADCY10\|AKAP7\|ARHGEF12\|DEPDC5\|GUCY1A2\|PRKAA2\|SOCS7 | 3.25E-02 | * |
| Undifferentiated (nC3) ∩ Inversely correlated with age at diagnosis | radial glia guided migration of Purkinje cell | 2 | DAB1\|SOCS7 | 4.03E-02 | * |
| Undifferentiated (nC3) ∩ Inversely correlated with age at diagnosis | cytokine-mediated signaling pathway | 6 | FLRT2\|HGF\|IL1RAP\|POU2F1\|PTPRZ1\|SOCS7 | 4.03E-02 | * |
| Undifferentiated (nC3) ∩ Inversely correlated with age at diagnosis | metanephros development | 3 | GLI3\|ITGA8\|NF1 | 4.99E-02 | * |
